# A novel virus alters gene expression and vacuolar morphology in *Malassezia* cells and induces a TLR3-mediated inflammatory immune response

**DOI:** 10.1101/2019.12.17.880526

**Authors:** Minji Park, Yong-Joon Cho, Donggyu Kim, Chul-Su Yang, Shi Mun Lee, Thomas L. Dawson, Satoshi Nakamizo, Kenji Kabashima, Yang Won Lee, Won Hee Jung

**Author notes:** Address correspondence to Won Hee Jung.

## Abstract

Most fungal viruses have been identified in plant pathogens, whereas the presence of viral particles in human pathogenic fungi is less well studied. In the present study, we observed extrachromosomal double-stranded RNA (dsRNA) segments in various clinical isolates of *Malassezia* species. *Malassezia* is the most dominant fungal genus on the human skin surface, and species in this group are considered etiological factors of various skin diseases including dandruff, seborrheic dermatitis, and atopic dermatitis. We identified novel dsRNA segments, and our sequencing results revealed that the virus, named MrV40, belongs to the Totiviridae family and contains an additional satellite dsRNA segment encoding a novel protein. The transcriptome of virus-infected *Malassezia restricta* cells was compared to that of virus-cured cells, and the results showed that transcripts involved in ribosomal biosynthesis were downregulated and those involved in energy production and programmed cell death were upregulated. Moreover, transmission electron microscopy revealed significantly larger vacuoles in virus-infected *M. restricta* cells, indicating that MrV40 infection dramatically altered *M. restricta* physiology. Our analysis also revealed that viral nucleic acid from MrV40 induced a TLR3-mediated inflammatory immune response in bone marrow-derived dendritic cells, suggesting that a viral element contributes to the pathogenicity of *Malassezia*.

**Importance:** *Malassezia* is the most dominant fungal genus on the human skin surface and is associated with various skin diseases including dandruff and seborrheic dermatitis. Among *Malassezia* species, *Malassezia restricta* is the most widely observed species on the human skin. In the current study, we identified a novel dsRNA virus, named MrV40, in *M. restricta* and characterized the sequence and structure of the viral genome along with an independent satellite dsRNA viral segment. Moreover, expression of genes involved in ribosomal synthesis and programmed cell death was altered, indicating that virus infection affected the physiology of the fungal host cells. Our data also showed that the viral nucleic acid from MrV40 induces a TLR3-mediated inflammatory immune response in bone marrow-derived dendritic cells, indicating that a viral element likely contributes to the pathogenicity of *Malassezia*. This is the first study to identify and characterize a novel mycovirus in *Malassezia*.

## Introduction

Viruses have been observed in many fungal species since their first identification in mushrooms (1). Fungal viruses, also known as mycoviruses, possess different forms of viral genomes including double-stranded RNA (dsRNA), single-stranded RNA (ssRNA), and single-stranded DNA (ssDNA). It is estimated that 30–80% of all fungal species, mainly endophytic fungi, are infected with viruses. Unlike viruses that infect other organisms, the transmission of fungal virus occurs vertically by cell division or horizontally via mating or hyphal anastomosis, with no extracellular phase of the virus life cycle. dsRNA segments have predominantly been found in fungal viruses, and, taxonomically, the fungal dsRNA viruses are classified into seven families: Chrysoviridae, Endornaviridae, Megabirnaviridae, Quadriviridae, Partitiviridae, Reoviridae, and Totiviridae (2).

The model fungus *Saccharomyces cerevisiae* also carries a dsRNA virus that belongs to the Totiviridae family and is known as the L-A virus. A unique feature of fungal viruses of the Totiviridae family is their capability to produce a killer toxin that lyses susceptible neighbor strains, whereas the virus-containing strain (also known as the killer strain) is immune to the toxin. Studies of how the virus produces killer toxins in *S. cerevisiae* showed that the killer toxins are encoded by a satellite dsRNA segment, known as the M satellite, within the L-A virus. To date, four different killer toxins, K1, K2, K28, and Klus, have been described (3–6). The *S. cerevisiae* L-A virus forms icosahedral particles with a diameter of approximately 39 nm (7). The virus possesses a non-segmented 4.6-kb dsRNA genome consisting of two open reading frames (ORFs), *gag* and *pol*, which overlap by 130 base pairs (bp) (8). A major 76-kDa capsid protein is encoded by *gag*, and a 180-kDa minor protein species is encoded as a Gag-Pol fusion protein by a −1 ribosomal frame-shift event (9, 10). The C-terminus of the Gag-Pol fusion protein possesses viral RNA-dependent RNA polymerase (RDRP) activity (8).

The ribosomal frameshift is an interesting feature in a compact viral genome and has been commonly found in various viral genomes as a mechanism to allow viruses to express overlapping ORFs. Studies of *S. cerevisiae* L-A virus revealed that the mechanism of frameshifting is based on the sequence structures including a canonical slippery heptamer, 5′-X XXY YYZ-3′ (X= A, U or G; Y=A or U; Z=A, U, or C) and RNA pseudoknot (11).

Fungal viruses have also been considered biocontrol agents in the field of agriculture. For example, a virus causes hypovirulence in the chestnut blight fungus *Cryphonectria parasitica* (12, 13), and a virus mediates the biocontrol of other phytopathogenic fungi such as *Helminthosporium victoriae*, *Sclerotinia sclerotiorum*, and *Botrytis cinerea* (14).

Although viral infections in fungal cells are widespread, the interactions between the fungal virus and its host are not well understood. One of the most studied host defense mechanisms is RNA silencing. Several studies have shown that RNA silencing functions as an antiviral defense mechanism against *C. parasitica* in fungi. Disruption of the dicer pathway in *C. parasitica* increases the susceptibility to virus infections (15), and p29 was identified as a suppressor that inhibits expression of the genes required for RNA silencing-mediated viral defense in the fungus (16). Similarly, conserved RNA silencing-mediated antiviral defense systems have been identified in *Aspergillus nidulans*, *Rosellinia necatrix*, and *Fusarium graminearum* (17–19).

*Malassezia* is the most dominant fungal genus on the human skin surface and is considered an etiological factor in various skin diseases including dandruff, seborrheic dermatitis, and atopic dermatitis (20–23). Eighteen *Malassezia* species have been identified; among them, *Malassezia restricta* is the most abundant on the human skin (20). Recent studies showed an increased burden of *M. restricta* on the scalp of patients with dandruff, indicating an association between dandruff and the fungus, although its role as a pathogenic organism is still unclear, and host susceptibility should be taken into consideration (21, 24–27). Most fungal viruses are found in plant pathogenic fungi, whereas few examples of viral particles have been identified in human pathogenic fungi such as *Candida albicans* (28). In the present study, we observed extrachromosomal dsRNA segments in various *M. restricta* clinical isolates which represented a novel viral genome and its satellite. Sequence analysis revealed that the virus belongs to the Totiviridae family and that the additional satellite dsRNA segment encodes a novel protein. The interactions between the viral elements and the fungal host, and the impact of the virus on fungal interactions with immune cells were evaluated.

## Results

### Identification of extrachromosomal dsRNA segments in *Malassezia*

Extrachromosomal nucleic acid bands were observed in total nucleic acid extracts of the *M. restricta* strains isolated in our recent study (29). Among the strains, *M. restricta* KCTC 27540 was used to identify extrachromosomal segments. Total nucleic acids were extracted from the strain and digested with DNase I, RNase A, and RNase T1. The extrachromosomal segments and ribosomal RNA were resistant to DNase I, indicating that they were neither ssDNA nor dsDNA. RNase A degraded all nucleic acids except for genomic DNA, whereas RNase T1, which catalyzes the degradation of ssRNA, removed ribosomal RNA only (30). These results suggested that the extrachromosomal segments correspond to dsRNA and that *M. restricta* KCTC 27540 possesses two separate extrachromosomal segments as estimated by agarose gel electrophoresis to be approximately 4.5 and 1.5 kb (Fig. 1A).

**Fig. 1.**
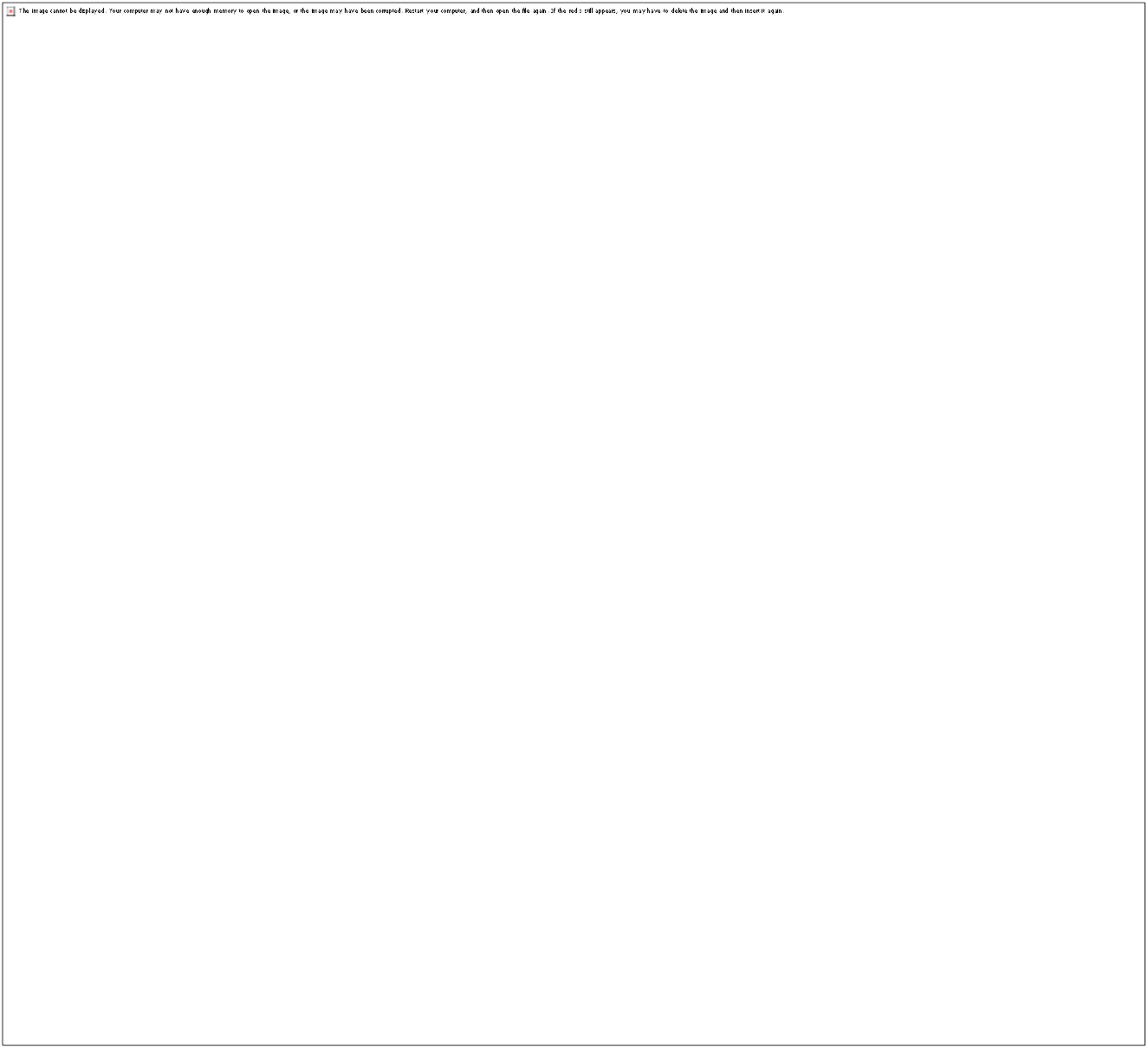
Extrachromosomal dsRNA segments in *Malassezia*. (A) Nucleic acids from *Malassezia restricta* KCTC 27540 were separated on a 0.7% agarose gel. Lane 1, total nucleic acids; lane 2, total nucleic acids treated with DNase I; lane 3, total nucleic acids treated with RNase A; lane 4, total nucleic acids treated with RNase T1. (B) Presence of dsRNA in *M. restricta* strains. Total nucleic acids were extracted and treated with DNase I and RNase T1 and were separated on a 0.7% agarose gel. (C) Total nucleic acids were extracted from different *Malassezia* species, treated with DNase I and RNase T1, and were separated on a 0.7% agarose gel.

To confirm whether the extrachromosomal segments observed in other *M. restricta* clinical isolates were also dsRNA, total nucleic acid extracts from strains other than *M. restricta* KCTC 27540 were treated with DNase I and RNase T1. The extrachromosomal segments remained unaffected after enzyme treatment, indicating that they are also dsRNA, as in *M. restricta* KCTC 27540 (Fig. 1B). Moreover, other *Malassezia* species including *M. globosa*, *M. pachydermatis*, and *M. sympodialis* showed similar extrachromosomal dsRNA segments suggesting that these segments are common characteristics of *Malassezia* species (Fig. 1C). Agarose gel electrophoresis revealed extrachromosomal segments composed of at least two separate dsRNA fragments except for *M. restricta* KCTC 27543, which showed a single dsRNA fragment. Additionally, the large fragments of dsRNA showed similar sizes (~5.0 kb), whereas the small dsRNA segments varied in size in different strains (Fig. 1B and 1C). We hypothesized that the dsRNA segments from *Malassezia* strains represent the dsRNA elements of mycoviruses, which are prevalent in all major fungal taxa (2). Indeed, a dsRNA mycovirus infected soil fungus, *Trichoderma atroviride* NFCF028, displayed similar extrachromosomal segments (Fig. S1) (31).

Sucrose gradient ultracentrifugation was conducted to purify virus particles to confirm that the dsRNA segments in the *Malassezia* strains were indeed viral elements. The separated nucleic acids and proteins in each fraction were analyzed. Two dsRNA fragments (~5.0 and ~1.7 kb) were clearly visible in fractions 1–6 following agarose gel electrophoresis (Fig. 2A). Moreover, the results of sodium dodecyl sulfate-poly acrylamide gel electrophoresis (SDS-PAGE) showed that fractions 3–6 contained protein bands with an estimated molecular weight of ~77 kDa (Fig. 2B). This molecular weight is similar to that of the known capsid protein of the *S. cerevisiae* mycovirus (9, 32, 33). Fractions 3–6 were subsequently evaluated by microscopy to visualize mycovirus particles in *M. restricta*. Transmission electron microscopy (TEM) images showed virus-like particles with an isometric shape and a diameter of 43 nm (Fig. 2C). These results support the hypothesis that the extrachromosomal dsRNA segments form the genome of mycovirus in *M. restricta* KCTC27540. We named the viral particle as MrV40 (*<u>M. r</u>estricta* KCTC 275<u>40</u> Mycovirus). The large and small dsRNA viral fragments were named as MrV40L and MrV40S, respectively. In addition to evaluating images of the purified virus particles, we examined the morphology of the virus-infected and the virus-cured *M. restricta* KCTC 27540 strain pairs by TEM. The virus-cured *M. restricta* KCTC 27540 strain was prepared by sterilizing the virus-infected *M. restricta* KCTC 27540 through numerous serial passages (Supplemental Materials and Methods, and Fig. S2). In general, there was no morphological difference between the virus-infected and the virus-cured KCTC 27540 strains. However, we noticed that the size of vacuoles in the virus-infected strain was significantly larger than that in the virus-cured strain, suggesting that the virus influences vacuole size in *M. restricta* (Fig. 3).

**Fig. 2.**
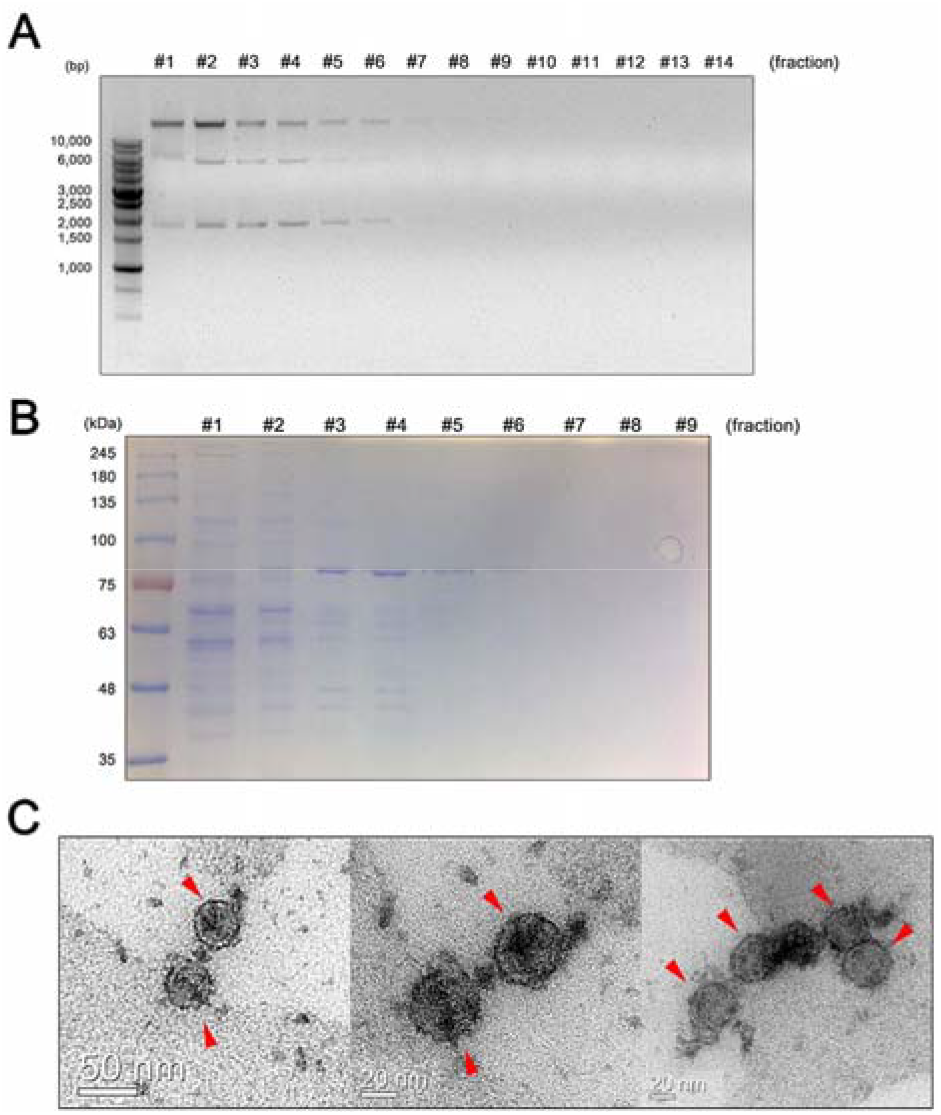
Isolation of virus particles from *Malassezia restricta* KCTC 27540, and their TEM images. The collected fractions after sucrose gradient ultracentrifugation were analyzed for their nucleic acids and proteins on an agarose gel (A) and SDS-PAGE gel (B), respectively. Images of the viral particles (red arrows) were obtained using a transmission electron microscope (C).

**Fig. 3.**
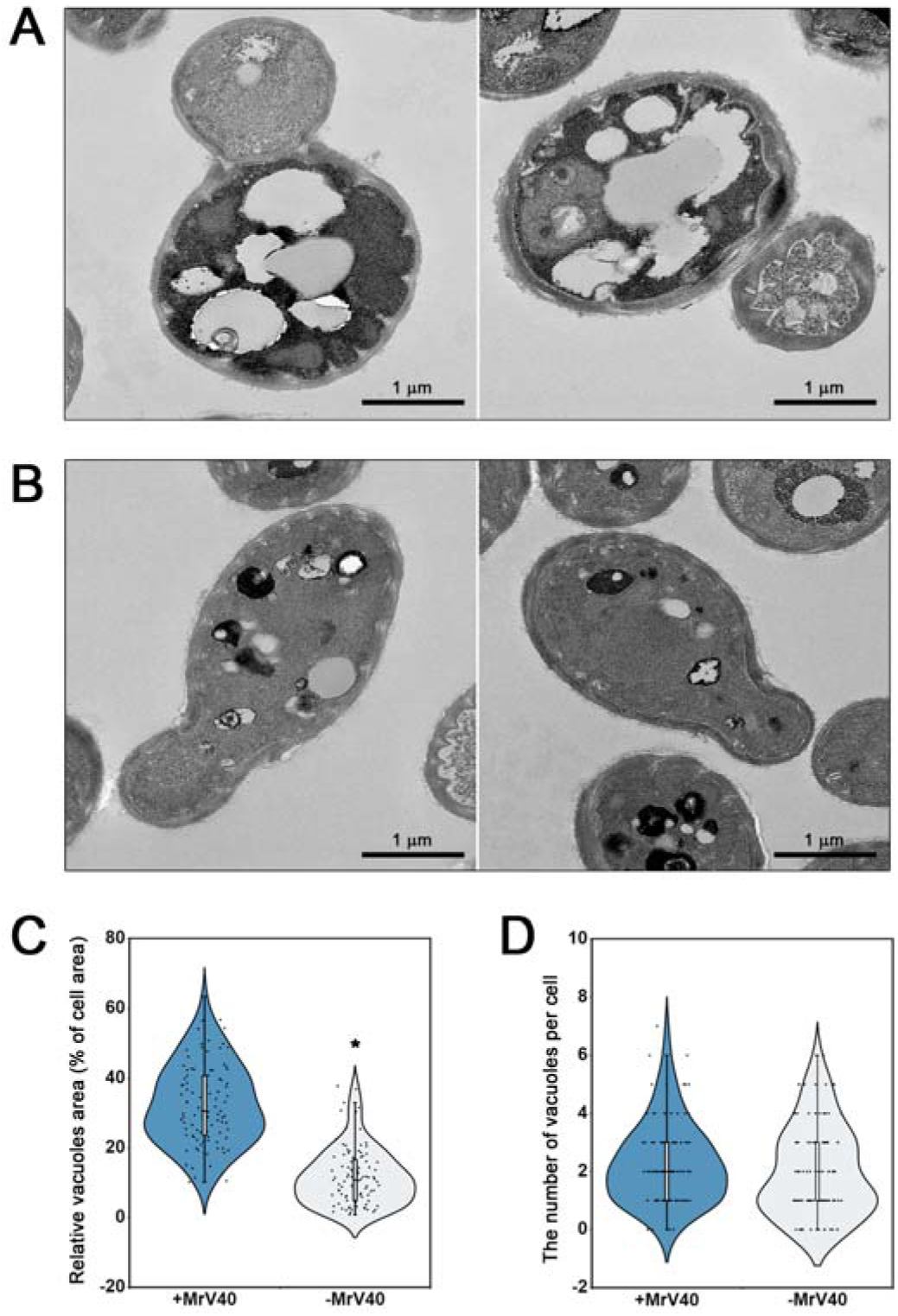
Morphology of vacuoles in virus-infected and virus-cured *Malassezia restricta* cells. Ultrathin sections of virus-infected (A) and virus-cured (B) *M. restricta* KCTC 27540 strains were observed by TEM. Cytoplasmic vacuole formation in a cell, expressed as the percentage of the area occupied by vacuoles (C) and the number of vacuoles (D) per cell, was estimated (n = 100 cells for each strain). The plot was created with BioVinci software (BioTuring Inc., San Diego, CA, USA). Statistical analysis for differences between the strains was performed with unpaired t-tests. *p < 0.001.

### Determination of dsRNA sequence of MrV40L

The complete sequence of MrV40L was determined by a combination of the Illumina MiSeq technique and the Sanger sequencing method using purified viral dsRNA. The length of the complete assembled sequence of MrV40L was 4,606 bp, and two overlapping open reading frames (ORFs), designated as ORF1 and ORF2, were identified (Fig. 4A). ORF1 corresponds to the region from nucleotides (nt) 28 to 2,097 and encodes a polypeptide of 689 amino acids with a molecular weight of 77 kDa. ORF2 corresponds to the region from nt 1,949 to 4,538 and encodes a polypeptide of 862 amino acids with a molecular weight of 98 kDa. The results of BLAST analysis showed that the protein sequences of ORF1 and ORF2 were highly similar to that of the capsid protein (the Pfam families of LA-virus_coat, PF09220) and viral RNA-directed RNA polymerase (RDRP; the Pfam family of RDRP_4, PF02123) of *S. segobiensis* virus L belonging to the genus *Totivirus* (Totiviridae family) with 53% (YP_009507830.1) and 52% (YP_009507831.1) identities, respectively (34). Eight conserved motifs, which are commonly found in totiviruses, were found within ORF2, supporting the classification of MrV40 as a totivirus (Fig. 4B) (35). Moreover, phylogenetic analysis with known amino acid sequences of RDRPs from dsRNA mycoviruses demonstrated that the RDRP encoded by the MrV40L genome was clustered with totiviruses (Fig. 4C). Thus, MrV40L is a dsRNA viral genome encoding a capsid protein and a RDRP, and we propose that MrV40 belongs to the *Totivirus* genus.

**Fig. 4.**
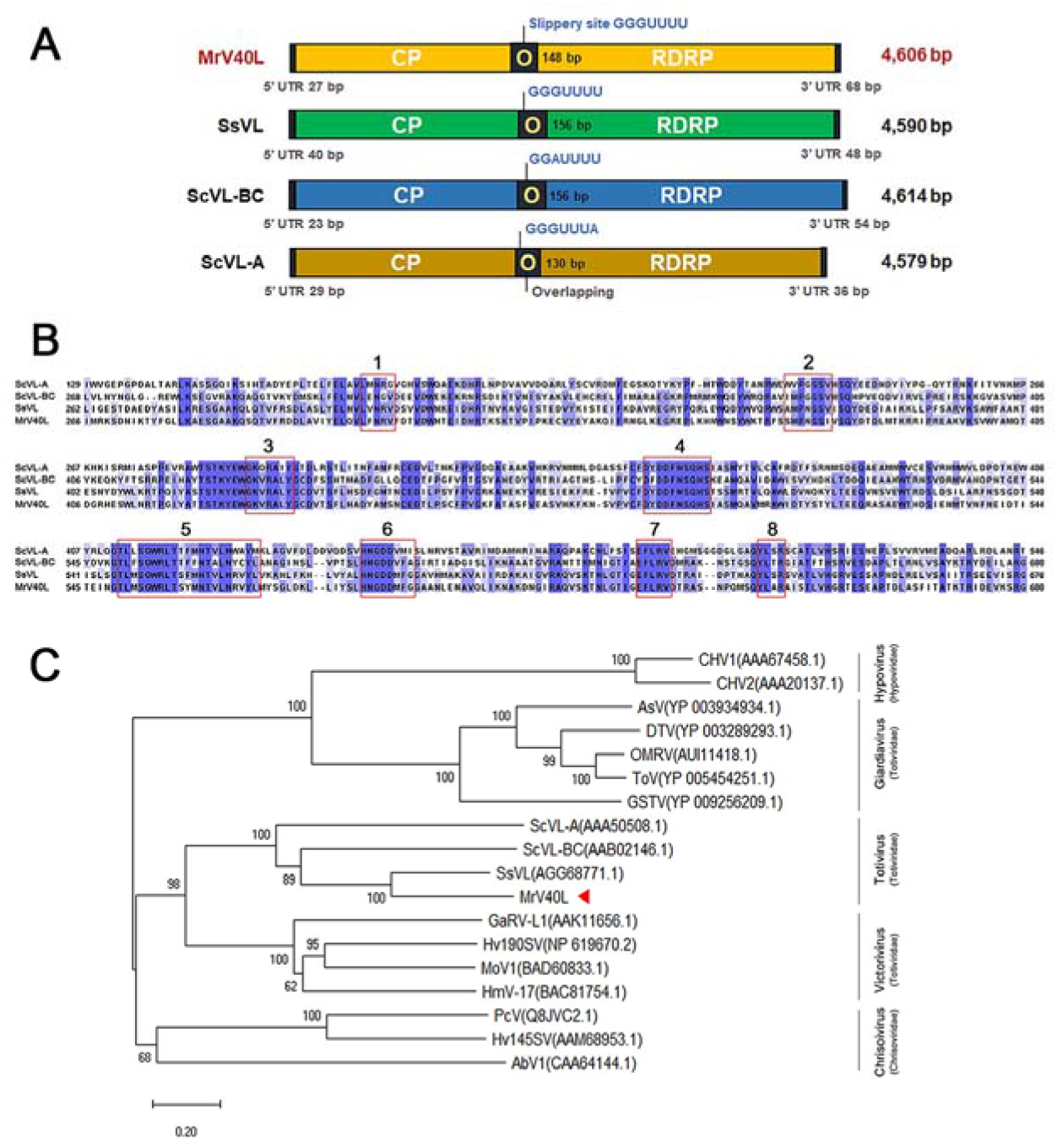
Genome and phylogenetic analysis of MrV40L. (A) Comparison of the genomic organization of MrV40L with that of other totiviruses. (B) Eight conserved motifs within the viral RDRP of MrV40L. The red boxes indicate conserved motifs. The amino acid sequences of ScVL-A, ScVL-BC, SsVL, and MrV40L were aligned using Jalview. (D) Multiple alignment of 18 amino acid sequences of RDRPs from dsRNA viruses was analyzed using the neighbor-joining method with bootstrap test (1,000 replicates) using MEGA X (82, 85, 86). The evolutionary distances are in units of the number of amino acid substitutions per site. All ambiguous positions were removed for each sequence pair. CHV1, *Cryphonectria* hypovirus 1; CHV2, *Cryphonectria* hypovirus 2; AsV, *Armigeres subalbatus* virus; DTV, *Drosophila melanogaster* totivirus; OMRV, *Omono river* virus; ToV, *Tianjin* totivirus; GSTV, *Golden shiner* totivirus; ScVL-A, *S. cerevisiae* virus L-A; ScVL-BC, *S. cerevisiae* virus L-BC; SsVL, *S. segobienesis* virus L; GaRV-L1, *Gremmeniella abietina* virus L1; Hv190SV, *Helminthosporium victoriae* virus-190S; MoV1, *Magnaporthe oryzae* virus 1; HmV-17, *Helicobasidium mompa* totivirus 1-17; PcV, *Penicillium chrysogenum* virus; Hv145SV, *Helminthosporium victoriae* 145S virus; AbV1, *Agaricus bisporus* virus 1 (8, 33, 34, 87–99).

Next, we analyzed the genomic sequences of dsRNA viruses in other clinical *M. restricta* isolates to confirm that they were similar to those of MrV40L, and thus belong to the *Totivirus* genus. RT-PCR was performed using four primer sets corresponding to the sequence of the conserved regions of the capsid protein and RDRP in the MrV40L genome (Fig. 5A). The results showed that six of 11 viruses generated RT-PCR products, suggesting the presence of viral genomes similar to MrV40Ls, while the remaining samples had dissimilar viral genomes (Fig. 5B).

**Fig. 5.**
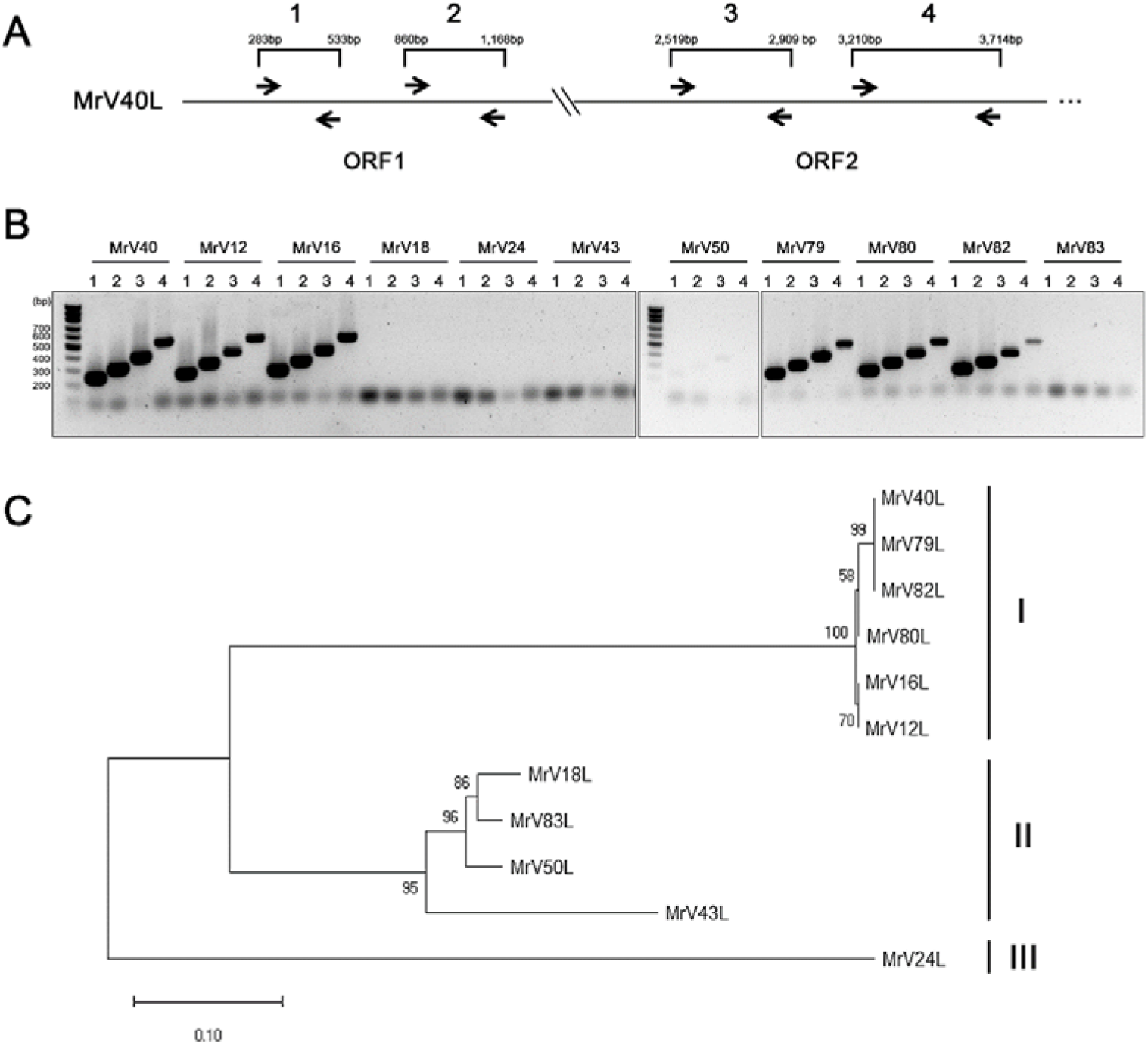
Phylogenetic classification of viruses found in *Malassezia restricta*. (A, B) The cDNAs of dsRNAs from *M. restricta* strains were amplified using primers for genes encoding CP and RDRP (ORF1 and ORF2, respectively) homologs in the viral genome. The following primers were used: lane 1, MrV40L_CP_F1 and MrV40L_CP_R1; lane 2, MrV40L_CP_F2 and MrV40L_CP_R2; lane 3, MrV40L_RDRP_F1 and MrV40L_RDRP_R1; lane 4, MrV40L_RDRP_F2 and MrV40L_RDRP_R2 (see Table S2 in supplemental material). (C) MrV-Ls were clustered into three clades (clade ◻, ◻, and ◻). Multiple alignment of nucleotide sequences of combined parts of *gag* and *pol* from 11 MrV-Ls was analyzed by the neighbor-joining method with bootstrap test (1,000 replicates) in MEGA X (82, 85, 86). The evolutionary distances are in units of the number of base substitutions per site.

Furthermore, we performed phylogenetic classification of the same viruses found in other clinical *M. restricta* isolates by multilocus sequence typing of the 1,075-bp region, 638 bp of ORF1, and 437 bp of ORF2, corresponding to *gag* and *pol*, respectively, within the viral genomes. The results revealed that the viruses were classified into three clades: clade ◻ (MrV12L, MrV16L, MrV40L, MrV79L, MrV80L, and MrV82L), clade ◻ (MrV18L, MrV43L, MrV83L, and MrV50L), and clade ◻ (MrV24L) (Fig. 5C). Additionally, the sequences of the 1,075-bp region of MrV40L, MrV79L, and MrV82L in the clade I were 100% identical, suggesting that they originated from the same lineage.

### Determination of dsRNA sequence of MrV40S

To determine the sequence of MrV40S, the dsRNA segments of MrV40S were extracted from agarose gels and then subjected to cDNA cloning and sequencing (See Materials and Methods). Using the partial sequences obtained from cDNA clones and the sequences obtained from repeated 5′ rapid amplification of cDNA ends (RACE), we successfully determined the complete sequence of MrV40S. The sequence of MrV40S was 1,355 bp, and a single ORF was identified in the 3′ region (from nt 773 to 1,097), which encoded a polypeptide of 124 amino acids with a molecular weight of 15.6 kDa. The ORF was designated as ORF3. Although we obtained the complete sequence of MrV40S ORF3, no homologous protein sequence was identified by BLAST analysis using all currently available databases.

Mycoviruses belonging to *Totivirus*, particularly the *S. cerevisiae* LA-virus, often possess a satellite dsRNA segment known as M dsRNA, which is responsible for producing a killer toxin that excludes neighboring yeast cells. Because MrV40L resembles M dsRNA, we predicted that MrV40S produces a protein that inhibits other *Malassezia* strains and/or other fungal and/or bacterial cells residing on the skin. To test the toxin-like activity of the protein produced from MrV40S, ORF3 was cloned and the protein was heterologously expressed in *Escherichia coli* and purified (Fig. 6A). The activity of the purified protein was evaluated against several pathogenic fungi and bacteria including *M. restricta*, *C. albicans*, *Cryptococcus neoformans, E. coli*, and *Staphylococcus aureus*. Unexpectedly, the purified protein displayed no growth inhibitory effect on the microbial cells tested (data not shown). Based on these results, we concluded that the novel protein produced from ORF3 likely has no toxin-like activity against the microorganisms tested, and further functional studies are required to characterize its function.

**Fig. 6.**
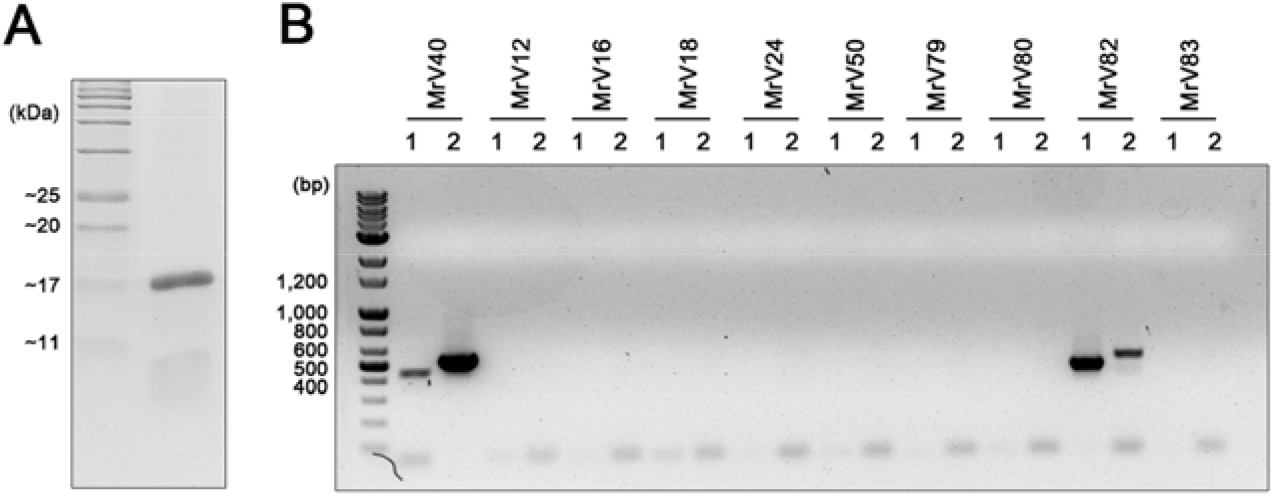
Characterization and similarity analysis of MrV40S. (A) The protein encoded by ORF3 was heterologously expressed in *E. coli* and purified using a His-tag column. (B) RT-PCR results. The primers specific to MrV40S were used to amplify satellite dsRNA from other *M. restricta* strain containing small dsRNA segments. The following primers were used: lane 1, MrV40S_F5 and V40S_SP1; lane 2, MrV40S_ORF_F1 and MrV40S_R1 (see Table S2 in supplemental material).

M dsRNA in *S. cerevisiae* produce several variants of killer toxins, namely, M1, M2, M3, and Mlus, and they showed limited homology in their protein sequences (4–6). Based on this information, we compared the sequences of satellite dsRNA genomes between MrV40S and those in other *M. restricta* strains containing the small dsRNA segments. Reverse transcription (RT)-PCR was conducted using a series of primers specific to MrV40S. The results suggested that among the nine strains tested, only the satellite dsRNA in *M. restricta* KCTC 27582 possessed a similar dsRNA segment. These data indicate that the satellite dsRNA genome sequences are highly variable (Fig. 6B).

### MrV40 altered transcriptome profiles in the host *M. restricta* strain

To understand the influence of MrV40 on the host physiology, we compared the transcriptomes of virus-infected and virus-cured *M. restricta* KCTC27540 strain pairs by RNA sequencing. Transcriptome analysis showed that 258 and 809 genes were up- and down-regulated by more than 2-fold, respectively, in the virus-infected strain compared to those in the virus-cured strain (Table 1 and Table S1). These results indicate that the presence of the virus may impact a number of physiological processes in *M. restricta*.

**Table 1.** Number of differentially expressed genes.

|  | Number of differential expressed genes (+virus/-virus) |  |  |  |  |
| --- | --- | --- | --- | --- | --- |
|  | 2-fold | 3-fold | 4-fold | >5-fold | Total |
| Upregulated | 204 | 32 | 12 | 10 | 258 |
| Downregulated | 597 | 152 | 70 | 60 | 809 |

In *S. cerevisiae*, numerous genes are known to be involved in maintaining its dsRNA virus (36) and are categorized into two groups: *MAK* (maintenance of killer) and *SKI* (superkiller). At least 25 *MAK* genes have been reported (37); among them, *MAK3*, *MAK10*, and *PET18* were shown to be required for the maintenance of both *S. cerevisiae* L-A virus and its M satellite, whereas all other *MAK* genes are responsible only for the M satellite (38–43). We could not find homologs of *MAK10* or *PET18* in the genome of *M. restricta*, suggesting that a different or modified virus maintenance mechanism may be present in the fungus compared to *S. cerevisiae* (Table 2). The results also showed that among the *MAK* homologs, *MAK1*, *MAK5*, *MAK11*, *MAK16*, *MAK21*, *MAK7*, and *MAK8* were downregulated by more than 2-fold. In *S. cerevisiae*, these genes, except for *MAK1*, were involved in 60S ribosomal subunit biosynthesis and M satellite maintenance. Thus, we predicted that MrV40 reduced ribosomal biogenesis within the host. Additionally, *M. restricta* contained a series of *SKI* homologs. However, no *SKI* homologs were differentially expressed in the virus-infected *M. restricta* KCTC27540 compared to those in the virus-cured strain, indicating that the function of *SKI* genes was not critical for maintaining the virus. In addition to the genes involved in virus maintenance, we observed upregulation of numerous genes required for the TCA cycle and the electron transport chain, including MRET_1104 (NADH dehydrogenase (ubiquinone) Fe-S protein 7), MRET_1378 (succinate dehydrogenase (ubiquinone) iron-sulfur subunit), MRET_1953 (NADH dehydrogenase (ubiquinone) Fe-S protein 1), MRET_2042 (fumarate hydratase, class II), MRET_2097 (succinate dehydrogenase (ubiquinone), MRET_2956 (2-oxoglutarate dehydrogenase E1 component), MRET_3173 (dihydrolipoamide dehydrogenase) and MRET_4117 (aconitate hydratase) (Table 3). These results suggest that virus maintenance and propagation may require higher energy production in the host cell.

**Table 2.**
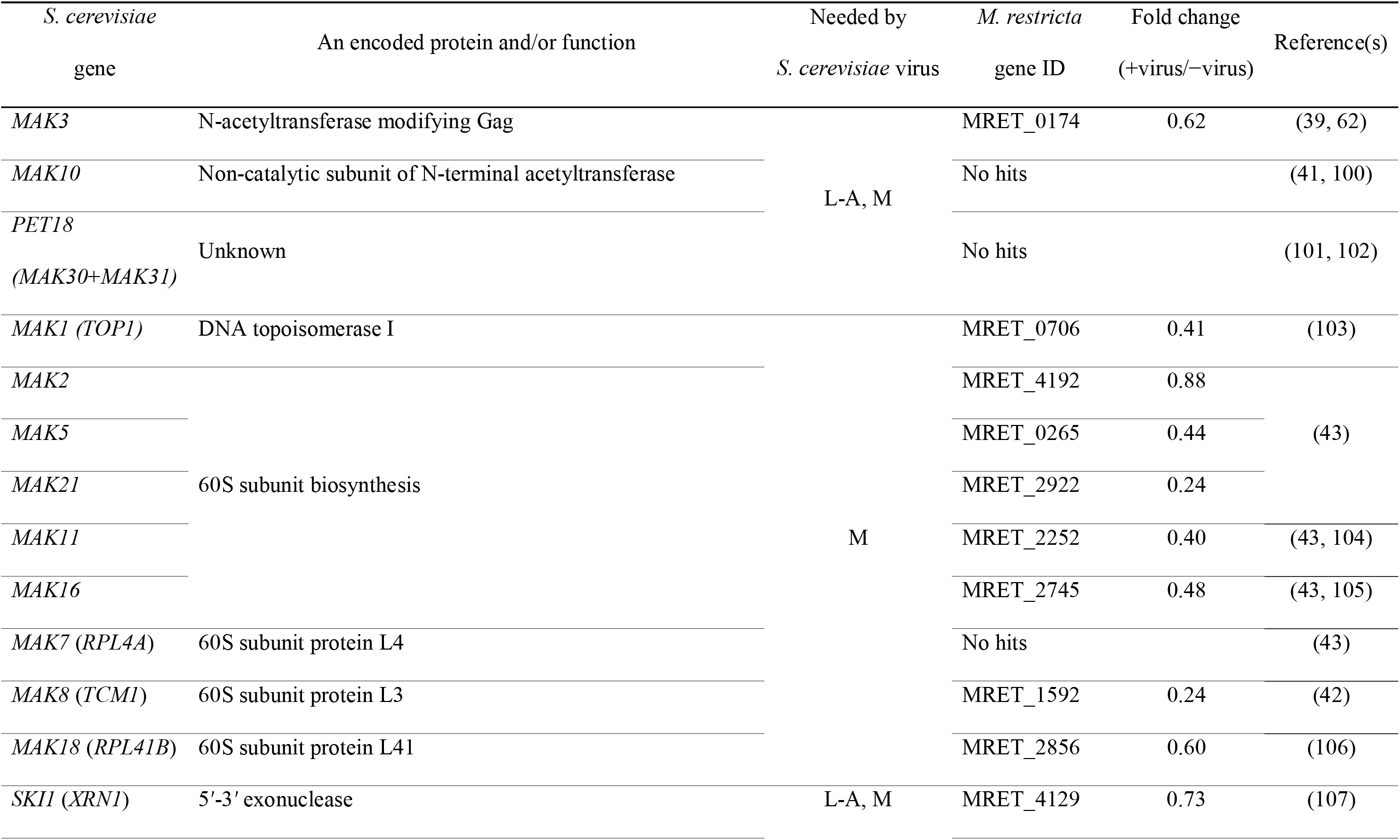
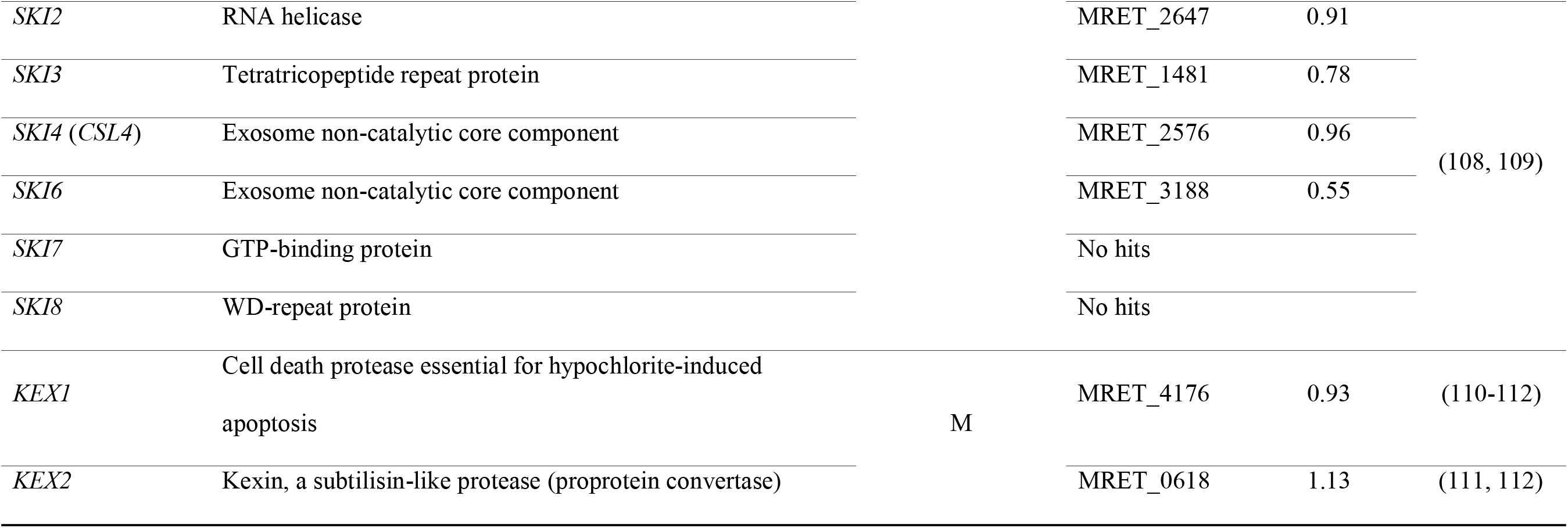
Differential expression of genes involved in maintaining dsRNA virus.

**Table 3.** Differential expression of genes involved in TCA cycle and electron transport chain.

| Gene | Annotation | Fold change<br>(+virus/−virus) |
| --- | --- | --- |
| MRET_1104 | NADH dehydrogenase (ubiquinone) Fe-S protein 7 | 2.85 |
| MRET_1378 | succinate dehydrogenase (ubiquinone) iron-sulfur subunit | 3.55 |
| MRET_1709 | cytochrome c | 2.56 |
| MRET_1953 | NADH dehydrogenase (ubiquinone) Fe-S protein 1 | 2.44 |
| MRET_2042 | fumarate hydratase, class II | 2.42 |
| MRET_2097 | succinate dehydrogenase (ubiquinone) flavoprotein subunit | 2.19 |
| MRET_2956 | 2-oxoglutarate dehydrogenase E1 component | 2.21 |
| MRET_3173 | dihydrolipoamide dehydrogenase | 2.27 |
| MRET_4117 | aconitate hydratase | 3.55 |

The overall dysregulation of primary metabolism may disturb the normal cell physiology in the *M. restricta* strain infected with MrV40. Indeed, we observed up-regulation of genes involved in programmed cell death in the fungal host cells. For example, the expression of MRET_3200 (p38 MAP kinase), MRET_1134 (programmed cell death 6-interacting protein), and MRET_2499 (autophagy-related protein, *ATG101*), which are associated with programmed cell death, was upregulated by 4.43-, 3.14-, and 2.83-fold in the virus-infected fungal cells, respectively. Moreover, MRET_0131 (FAS-associated factor), which is involved in Fas-induced apoptosis, was found to be strongly upregulated (8.80-fold) in the presence of the virus within the fungal host (44). It is well-known that programmed cell death is triggered during virus infection (45, 46), and our findings agree with this observation. To confirm that there was an increase in programmed cell death, we stained cells with Annexin V and propidium iodide, which is commonly used to quantitate the apoptosis (47, 48). Using flow cytometric analysis, we found higher fluorescence intensity in the virus-infected KCTC 27540 strain than in the virus-cured strain, confirming that programmed cell death is increased in the virus-infected *M. restricta* strain (Fig. S5). We also tested whether increased programmed cell death influence other physiological characteristics of the virus-infected cells such as susceptibility to an antifungal drug. However, antifungal susceptibility of the virus-infected KCTC 27540 was similar to that of the virus-cured strain (data not shown), suggesting that the viral infection may have a limited effect on the physiological characteristics of the host cells.

### MrV40 induced the TLR3-mediated immune system

Since it was first reported that fungal viral dsRNA induces cytokine production in rabbits (49–51), several studies have demonstrated that Toll-like receptor 3 (TLR3) plays a central role in viral dsRNA recognition and production of inflammatory cytokines in innate immune cells (52, 53). Additionally, a recent study showed that *S. cerevisiae* viral dsRNA stimulates the immune system through TLR3 in a human embryonic kidney cell line (54). We therefore investigated whether MrV40 itself or the virus-containing *Malassezia* cells alter the expression patterns of TLRs and cytokine production in mammalian cells. To this end, bone marrow-derived dendritic cells (BMDCs) obtained from C57BL/6 mice, which have been used as a model to study the interactions between fungal cells including *Malassezia* and the innate immune system in a mammalian host, were used in our study (55, 56).

Two independent strain pairs, virus-infected and virus-cured *M. restricta* KCTC 27540 strains as well as virus-infected and virus-cured *M. restricta* KCTC 27524 strains were used for the study. Live fungal cells, fungal cell lysates, purified capsid protein, purified satellite protein (produced from MrV40S ORF3), and purified MrV40 dsRNA were prepared and co-incubated with BMDCs, and the expression of TLRs and cytokines were analyzed. We found that TLR3 expression was significantly upregulated by live fungal cells and fungal cell lysates of the virus-infected *M. restricta* strains, while live cells and their lysates of the virus-cured strains did not enhance the expression of TLR3. Considering that the expression of other TLRs were unchanged upon addition of live cells and cell lysates, we concluded that host response upon treatment with virus-infected *M. restricta* cells was mainly mediated by TLR3 in murine BMDCs. Moreover, upregulation of TLR3 expression was observed in BMDCs treated with purified MrV40 dsRNA but not with purified capsid and satellite proteins, suggesting that MrV40 dsRNA is the causative agent that enhances the expression of TLR3 (Fig. 7A). Moreover, our data suggested that purified capsid and satellite proteins from MrV40 did not influence the expression of TLRs.

**Fig. 7.**
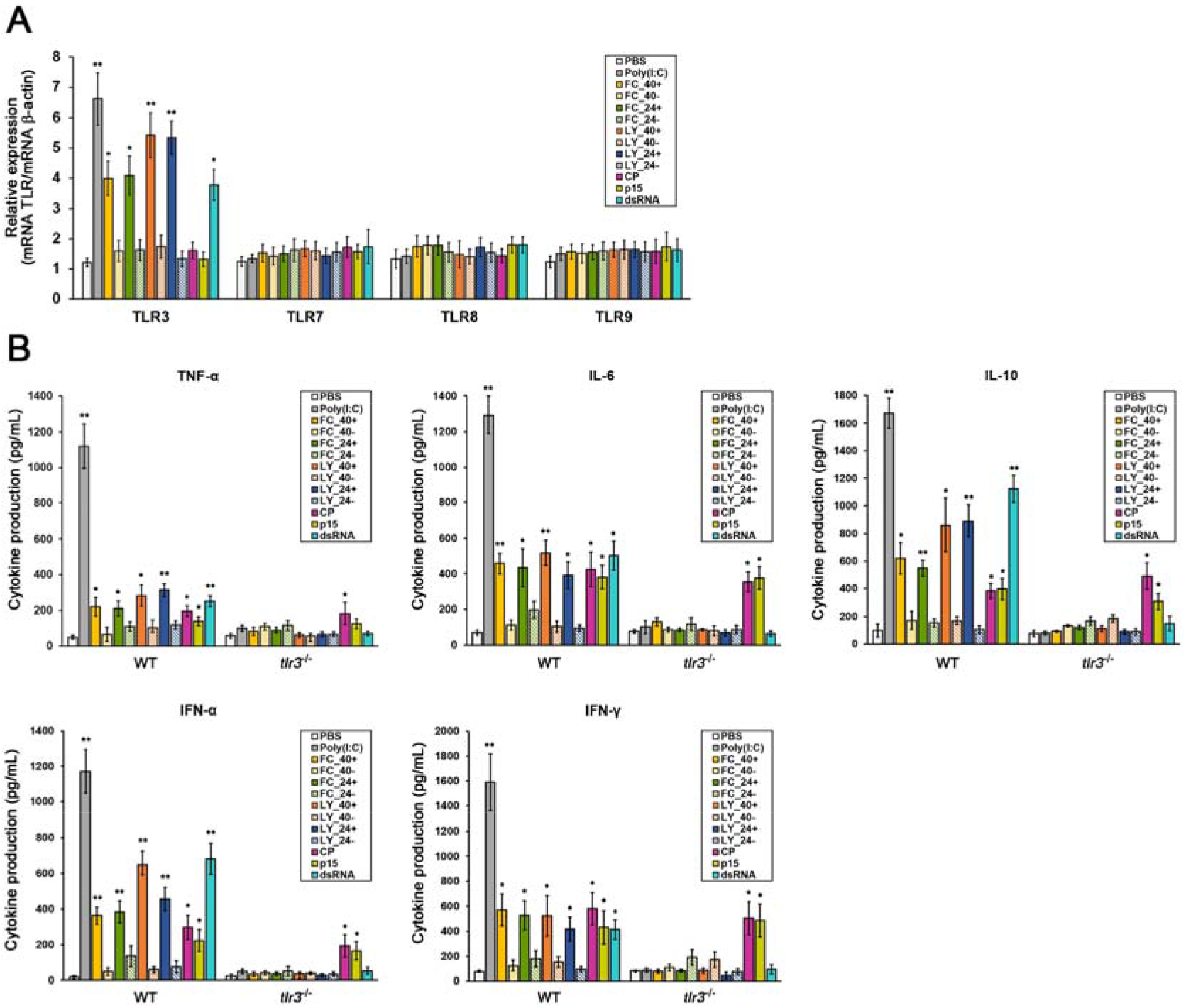
Evaluation of TLR and cytokine levels upon treatment with the viral elements. *TLR* expression (A) and cytokine production (B) in BMDCs co-incubated with live fungal cells, total fugal cell lysates, purified MrV40 capsid protein, purified satellite protein, and purified MrV40 dsRNA. PBS and poly (I:C) served as negative and positive controls, respectively. WT, wild-type mice (C57BL/6); *tlr3*^-/-^, TLR3 knock out mice (B6;129S1-Tlr3^tm1Flv^/J, 005217); FC, live fungal cells; LY, cell lysate; 40+, virus-infected KCTC 27540; 40-, virus-cured KCTC 27540; 24+, virus-infected KCTC 27524; 24-, virus-cured KCTC 27524; CP, purified capsid protein; p15, purified MrV40S satellite protein; dsRNA, purified MrV40 dsRNA. Values represent the average from quadruplicates with standard deviations. Statistical analysis for differences between PBS and each sample was performed with unpaired t-tests. *p < 0.001 and **p < 0.0001.

The expression of several inflammatory cytokines including tumor necrosis factor (TNF)-α, interleukin (IL)-6, IL-10, interferon (IFN)-α and IFN-γ, in response to live fungal cells, fungal cell lysates, purified capsid protein, purified satellite protein, and purified MrV40 dsRNA, were evaluated. Particularly, IFN-α and IFN-γ were included because of their involvement in the antiviral response in mammals (57). The results of the cytokine analysis indicated that virus-infected live *M. restricta* cells and their lysates induced the production of TNF-α, IL-6, IL-10, IFN-α and IFN-γ, while no change in cytokine expression was observed in BMDCs treated with virus-cured live *M. restricta* cells and their cell lysates. Similarly, capsid protein and MrV40 dsRNA also enhanced the expression of all cytokines tested (Fig. 7B). Cytokine profiles were also measured in homozygous TLR3 knockout mice, as our observations suggested a TLR3-mediated host response in BMDCs treated with virus-infected fungal cells. As shown in Fig. 7B, cytokine levels in BMDCs isolated from TLR3 knockout mice were not significantly altered except in those treated with the capsid and satellite proteins. These data suggest that MrV40 increased the production of the cytokines TNF-α, IL-6, IL-10, IFN-α, and IFN-γ in a TLR3-dependent manner in BMDCs. However, as suggested above, the cytokine production induced by the capsid protein seemed to be TLR3-independent. Moreover, the purified satellite protein also caused a TLR3-independent increase in cytokine production except of TNF-α. These results suggest that the capsid and the satellite proteins also contributed to upregulation of cytokine expression in a TLR3-independent manner. Overall, our data suggest that dsRNA in MrV40 triggers an increase in the production of cytokines involved in inflammation and that TLR3 plays a central role in the host response.

## Discussion

In the current study, we detected dsRNA virus in several clinical isolates of *Malassezia* species. Among them, MrV40 identified in *M. restricta* KCTC 27540 was selected, and its genome structure and effects on host gene expression and on the mammalian immune response were evaluated. Our data showed that MrV40 consists of two RNA segments, which we named MrV40L and MrV40S. The results of the genome sequence analysis suggested that these segments were 4,606 and 1,355 bp in length, respectively, and belong to the genus *Totivirus*. Typically, the genomes of the viruses belonging to the genus *Totivirus* consist of non-segmented dsRNA with sizes between 4.6 and 7.0 kb, and contain two ORFs, *gag* and *pol*. Studies have specifically examined the genome structure of *Totivirus* because of the overlapping nature of the two ORFs, where a −1 frameshift occurs, resulting translation of the fusion protein (2). The overlapping ORFs and frameshift were frequently observed in a compact viral genome to translate proteins and were found in several dsRNA and ssRNA viruses; these ORFs allow ribosomes to translate CP and RDRP continuously with a missing CP termination codon (11, 58, 59). The mechanism of frameshifting in the viral genome is based on the pseudoknot structure of the mRNA for efficient slipping via a slippery site (60). For example, in the genome of the *S. cerevisiae* L-A virus, the 5′ ORF (*gag*) encodes a 76-kDa CP and the 3′ ORF (*pol*) encodes an RDRP, which is expressed as a 180-kDa CP-RDRP fusion protein generated by a −1 ribosomal frameshift (8, 11). In our study, MrV40L (the major dsRNA segment in MrV40) contained two overlapping ORFs, ORF1 and ORF2. We identified a putative slippery site heptamer, 5′-GGGTTTT-3′, at the region from nt 1, 968 to 1,974 for the −1 ribosomal frameshift, which may be associated with production of the fusion protein of 170 kDa in MrV40L. A previous study suggested that in the *S. cerevisiae* L-A virus, the rate of ribosomal frameshifts was approximately 1.8%, giving 120 CP and 2 CP-RDRP fusion protein molecules per virus particle (11). Considering the low efficiency of producing the fusion proteins by ribosomal frameshifting, we expected to observe a significantly lower translation rate of the fusion protein; indeed, the putative 170-kDa band was not detected by SDS-PAGE. In addition to MrV40L, we determined the sequence of the satellite dsRNA segment MrV40S and found that it consists of 1,355 nt containing a single ORF, ORF3, producing a novel 15.6-kDa protein. As observed for other totiviruses, the possible toxin-like activity of the protein was investigated in our study, but no growth inhibitory activity against several bacteria and fungi was observed.

It has been estimated that 30–80% of fungal species in nature are infected with viruses, and a fungal host normally shows no specific symptoms upon infection (14). However, several genes were shown to be required for maintaining and propagating viruses in the host fungal cells. In *S. cerevisiae*, numerous chromosomal genes are known to be involved in viral propagation and expression of the viral killer toxin (36). Furthermore, several genes are known to be responsible for maintaining the L-A virus and M dsRNA in *S. cerevisiae*. The *MAK* genes are required for the propagation and maintenance of the L-A virus and M dsRNA in *S. cerevisiae* (61). Among the *MAK* genes, *MAK3*, *MAK10*, and *PET18* are required for the maintenance of both L-A virus and M dsRNA, whereas all other *MAK* genes are responsible only for M dsRNA (38–43). Particularly, *MAK3* encodes an N-acetyltransferase and is required for N-acetylation of the coat protein (38, 62). A previous study showed that the coat proteins without acetylation failed to self-assemble, resulting in the loss of all dsRNA viruses (39). *MAK10* and *PET18* (*MAK30*+*MAK31*) encode a non-catalytic subunit of N-terminal acetyltransferase and a protein of unknown function, respectively. Mutant strains lacking each gene contained unstable viral particles, indicating that the genes are involved in the structural stability of LA-virus and M dsRNA (40). *MAK1* (*MAK17*, *TOP1*) encodes DNA topoisomerase I, and other *MAK* genes including *MAK2*, *MAK5*, *MAK11*, *MAK16*, *MAK21*, *MAK7* (*RPL4A*), *MAK8* (*TCM1*), and *MAK18* (*RPL41B*) are related to 60S ribosomal subunit assembly (43, 63, 64). All mutant strains lacking the above genes showed decreased levels of free 60S ribosomal subunits and the inability to maintain M dsRNA, suggesting that stable propagation of the satellite dsRNA depends on 60S ribosome synthesis (42, 43). In addition to the *MAK* genes, the *SKI* gene family has been shown to be involved in the maintenance and propagation of virus in *S. cerevisiae*. *SKI1* (*XRN1*) encoding a 5′-3′ exonuclease is involved in the degradation of uncapped mRNA including viral mRNA, and *SKI2*, *SKI3*, *SKI6*, *SKI7*, and *SKI8* block the translation of viral mRNAs (65–67).

In the current study, homologs of most *MAK* and *SKI* genes were identified in *M. restricta*. The results of the transcriptome analysis suggested that most *MAK* genes were downregulated, which may in turn reduce ribosome synthesis in the *M. restricta* strain containing MrV40. In contrast, no *SKI* homolog showed significantly altered transcript levels in the *M. restricta* strain harboring MrV40. Moreover, we found enhanced expression of genes involved in energy metabolism and programmed cell death in the *M. restricta* strain containing the virus. *S. cerevisiae* also displayed relatively small changes in fungal host gene expression upon virus infection, possibly because of co-adaptation of the virus within the fungal host (68). Maintenance and propagation of virus within the fungal host may be involved in the post-transcriptional mechanism and may contribute to the minimal changes in host gene expression. Notably, the possibility that an RNA silencing pathway in *M. restricta* cells influences virus maintenance was excluded because of the absence of homologous genes required for the pathway in the genome of the fungus.

In addition to transcriptome analysis, we directly investigated whether the virus influences the cellular morphology of *M. restricta* and the structures of its intracellular organelles by TEM. We observed significantly larger vacuoles in virus-infected *M. restricta* cells. An increased vacuole size upon virus infection has been reported previously. The phytopathogenic fungus *Botrytis porri* infected with dsRNA virus 1 showed the formation of abundant vacuoles (69), and turnip mosaic virus induced the formation of a large central vacuole in *Nicotiana benthamiana* plant cells (70). Particularly, in *N. benthamiana*, turnip mosaic virus particles were shown to accumulate in vacuoles and be protected by the organelle membranes against the harsh host environment, particularly during xylem vessel differentiation. Furthermore, virus accumulation in vacuoles in plant-to-plant virus transmission has been suggested (70). Although the expression of genes involved in vacuole biogenesis was not altered in transcriptome analysis in virus-infected *M. restricta* cells, there may be a relationship between vacuole functions and MrV40 in *M. restricta*. Therefore, additional physiological studies are needed.

TLR3 is well-conserved in most vertebrates, localizes on the endosomal transmembrane, and plays a role in immune and non-immune cells. It has been suggested that TLR3 senses viral dsRNA taken up through endocytosis and contributes to defending the host against viral infection by regulating the expression of a range of cytokines (52, 71). A previous study demonstrated a significant decrease in cytokines (IFN-α/β, IFN-γ, and IL-12p40) and increase in viral PFU in the spleen of *Tlr3*^−/−^ mice infected with mouse cytomegalovirus (72). We found that TLR3 expression was significantly increased by the live cells of two independent virus-infected *M. restricta* strains and their cell lysates, while no change was found by the live cells and cell lysates of the virus-cured strains. The causative agent was very likely to be dsRNA of MrV40 in the virus-infected *M. restricta* strains since purified dsRNA from the MrV40 particles enhanced TLR3 expression. The virus-infected live *M. restricta* cells, their cell lysates, and purified dsRNA from the MrV40 particles also showed increased production of cytokines involved in inflammation. However, in *Tlr3*^−/−^ BMDCs, the production of all cytokines in the cells treated with the virus-infected live *M. restricta* cells, their cell lysates, and purified dsRNA from the MrV40 particles was attenuated, suggesting that TLR3 plays a central role in the host response against dsRNA of MrV40 and mediates the production of the cytokines.

Similarly, the *Leishmania* parasite *L. guyanensis* infected with the dsRNA virus belonging to *Totivirus* induced a TLR3-mediated inflammatory immune response within the vertebrate host, indicating that the *Leishmania* RNA virus (LRV1) functions as an innate immunogen. Moreover, in *Leishmania*, LRV1 was suggested to stimulate the inflammatory immune response and increase the severity and persistence of the parasite (73). However, whether MrV40 contributes to hyper-or hypo-pathogenicity of *M. restricta* remains unclear. Therefore, further studies to characterize the potential function of MrV40 as an innate immunogen are needed. In addition, since our data suggest contribution of the MrV40 capsid and the satellite proteins to host cytokine expression in a TLR3-independent manner, additional studies are needed to define a role of the capsid and the satellite proteins in interactions between these viral components and the mammalian host cells.

Overall, our study demonstrated the existence of a dsRNA virus within *M. restricta*. We also determined the sequences and structure of the viral genome along with the independent RNA segment. Moreover, we identified viruses not only from different strains of *M. restricta* but also from other *Malassezia* species, although variations were observed in the viral genomes. Evidence that the viral nucleic acid from MrV40 induces a TLR3-mediated inflammatory immune response was obtained, suggesting that a viral element contributes to the pathogenicity of *Malassezia*.

## Materials and Methods

### Strains and growth conditions

*M. restricta* KCTC 27512, KCTC 27516, KCTC 27518, KCTC 27524, KCTC 27540, KCTC 27543, KCTC 27550, KCTC 27879, KCTC 27880, KCTC 27882, and KCTC 27883, *M. globosa* CBS 7966, *M. pachydermatis* KCTC 27587, and *M. sympodialis* KCTC 27817 were obtained as previously described and cultured on Leeming and Notman agar (LNA) medium at 34°C for 3 days (29, 75–77). Among these strains, *M. restricta* KCTC 27540 was used to identify dsRNA viruses. *E. coli* DH10B and BL21 were grown in Luria-Bertani broth at 37°C.

### Purification of virus particles

The virus particles were purified as previously described with some modifications (69). Briefly, approximately 10 g of *M. restricta* cells was harvested and suspended in 10 mL of extraction buffer (0.1 M sodium phosphate buffer containing 3% (v/v) Triton X-100, pH 7.0). The suspended cells were vortexed with glass beads and centrifuged at 10,000 ×*g* at 4°C for 20 min to remove cell debris. The supernatant containing crude extracts was ultracentrifuged at 119,000 ×*g* at 4°C for 2 h to precipitate all particles. The resulting pellet was suspended in 1 mL of 0.1 M sodium phosphate buffer (pH 7.0), and the suspension was centrifuged at 16,000 ×*g* at 4°C for 30 min to remove large particles. The supernatant was overlaid on sucrose solution with a gradient concentration ranging from 10% to 40% (w/v) and centrifuged at 70,000 ×*g* at 4°C for 2 h. Fractions (700 μL/each fraction) were collected from the top of the ultracentrifuged sucrose gradient solution. Each fraction was analyzed by 1% agarose gel electrophoresis and 8% SDS-PAGE to detect dsRNA segments and protein, respectively. Methods for cell fixation and TEM are described in the Supplementary Materials and Methods.

### Determination of MrV40L and MrV40S sequences

The reference genome sequence was required to obtain the complete sequence of MrV40L by Illumina Miseq. We first obtained the partial fragment of MrV40L and determined its nucleotide sequence. Briefly, 1 μg of cDNA obtained as described above was amplified using 1 μM of the tagged oligo_1 primer to bind the tag and 1.25 units of *Taq* DNA polymerase (BIONEER, Daejeon, Korea). The amplicons were cloned into the pGEM-T vector (Promega, Madison, WI, USA), transformed into *E. coli* DH10B, and sequenced with the universal M13 primers. The resulting partial sequences of MrV40L showed high similarity with the genome of *Scheffersomyces segobiensis* virus L isolate NRRL Y-11571, which was used to assemble the reference genome in our study. To sequence the entire genome of the virus from *M. restricta* KCTC 27540, dsRNA was purified using Cellulose A (Advantec, Taipei, Taiwan) from total cell extract as described previously (78). After purification, purified dsRNA was treated with Ambion™ DNase I (Thermo Fisher Scientific, Waltham, MA, USA) to remove remaining genomic DNA remained. The sequencing library was constructed using the TruSeq Stranded Total RNA Sample Prep Kit (Illumina, San Diego, CA, USA) following the manufacturer’s instructions and excluding poly-A selection. The resulting library was sequenced on an Illumina MiSeq instrument according to the manufacturer’s instructions. The generated raw sequences, 250-bp paired-end reads, were assembled by CLC Genomics Workbench v7.5 (Qiagen, Hilden, Germany), and the resulting contigs were identified by BLASTN of the NCBI nucleotide database. Based on the reference genome, two contigs of MrV40L were assembled. The gap between the contigs was sequenced by gap-filling RT-PCR using primers Linkage1_V40 and Linkage2_V40) (see Table S2), and both termini were sequenced by RACE PCR using a 5′/3′ RACE Kit, 2^nd^ Generation (Roche, Basel, Switzerland) according to the 5′ RACE protocol described by the manufacturer.

To determine the MrV40S genome, the RACE PCR method was used because we failed to find any contig of MrV40S. Like MrV40L, the partial sequence of cDNA from MrV40S was amplified using the tagged oligo_1 primer. The amplicons were cloned into the pGEM-T vector (Promega) and sequenced. Using the partial sequence obtained, 5′ RACE PCR was performed and repeated until no additional sequence was found.

### *In silico* analysis

To identify the ORFs in MrV40L and MrV40S, the nucleotide sequences of the genomes were analyzed by ORFfinder (https://www.ncbi.nlm.nih.gov/orffinder/). To predict ribosomal frameshift signals, a putative slippery site was identified using FSFinder2 (79). The amino acid sequences of CP and RDRP of MrV40L were analyzed using the Pfam database (http://pfam.xfam.org/) and BLASTP (https://blast.ncbi.nlm.nih.gov/). For sequence alignment, Jalview2.0 was used (80). The phylogenetic tree showing the relationship of MrV40L with RDRP of other dsRNA fungal viruses was constructed using MEGA X, and phylogenetic reconstruction analysis was performed by the neighbor-joining method with 1,000 bootstrap replications (81, 82).

### Mice and cell culture

Wild-type C57BL/6 mice were purchased from Orient Bio Co. (Gyeonggi-do, Korea). *TLR3*^-/-^ (B6;129S1-Tlr3^tm1Flv^/J, 005217) mice were obtained from Jackson Laboratory (Bar Harbor, ME, USA). All animal experimental procedures were reviewed and approved by the Institutional Animal Care and Use Committee of Hanyang University (protocol 2018-0085) and performed in accordance with Korean Food and Drug Administration guidelines. All animals were maintained in a specific pathogen-free environment. Primary BMDCs were isolated from C57BL/6 mice and cultured in Dulbecco’s modified Eagle medium for 3–5 days in the presence of 20 ng/mL recombinant granulocyte-macrophage colony-stimulating factor (R&D Systems, Minneapolis, MN, USA) as described previously (83). Cell cultures were stained to detect dendritic cells with CD11c-FITC (eBiosciences, San Diego, CA, USA).

### Cytokine measurement

Mouse cytokines in the culture supernatants were measured with a BD OptEIA ELISA kit (BD Biosciences, Franklin Lakes, NJ, USA) as described previously (84). All assays were performed according to the manufacturer’s instructions. Two independent strain pairs, virus-infected and virus-cured *M. restricta* KCTC 27540 and virus-infected and virus-cured *M. restricta* KCTC 27540, were used. To treat BMDCs with live fungal cells, a 20:1 ratio (2.0×10^6^ and 1.0×10^5^ cells/mL of *M. restricta* and BMDCs, respectively) was used. A final concentration of 100 μg/mL (total protein/volume) of fungal cell lysates was used to treat BMDCs. A final concentration of 1 μM of purified MrV40 capsid and satellite proteins and 500 ng/mL of purified MrV40 dsRNA from virus-infected M. restricta KCTC 27540 strain were incubated with BMDCs. Phosphate-buffered saline (PBS) and polyinosinic-polycytidylic acid [poly (I:C), 25 μg/mL] served as negative and positive controls, respectively. Cytokines were measured 6 h after treatment. All experiments were repeated four times.

### cDNA synthesis, RNA isolation, transcriptome analysis by RNA sequencing, quantitative real-time PCR, and heterologous expression and purification of MrV40 proteins

See Supplementary Materials and Methods

## Data availability

The complete genome sequences of MrV40L and MrV40S were deposited into GenBank under accession numbers MN603497 and MN603498, respectively. The transcriptome data were deposited in Gene Express Omnibus under the accession number GSE138985.

## Acknowledgments

We thank James W. Kronstad for critical reading of the manuscript and Taegun Seo for helpful discussions. There is a companion paper from the Joe Heitman group at Duke University, and we thank Joe Heitman and Giuseppe Ianiri for communication. This work was supported by the National Research Foundation of Korea (NRF) grant funded by the Korea government (MSIT) 2019R1F1A1061930 (WHJ), 2019R1A4A1024764 (WHJ), and 2019R1I1A2A01064237 (CSY).

## References

1. Hollings M. 1962. Viruses associated with a die-back disease of cultivated mushroom. Nature 196:962.

2. Ghabrial SA, Caston JR, Jiang D, Nibert ML, Suzuki N. 2015. 50-plus years of fungal viruses. Virology 479-480:356–68.

3. Bussey H. 1991. K1 killer toxin, a pore-forming protein from yeast. Mol Microbiol 5:2339–43.

4. Hannig EM, Leibowitz MJ. 1985. Structure and expression of the M2 genomic segment of a type 2 killer virus of yeast. Nucleic Acids Res 13:4379–400.

5. Schmitt MJ, Tipper DJ. 1990. K28, a unique double-stranded RNA killer virus of Saccharomyces cerevisiae. Mol Cell Biol 10:4807–15.

6. Rodriguez-Cousino N, Maqueda M, Ambrona J, Zamora E, Esteban R, Ramirez M. 2011. A new wine Saccharomyces cerevisiae killer toxin (Klus), encoded by a double-stranded rna virus, with broad antifungal activity is evolutionarily related to a chromosomal host gene. Appl Environ Microbiol 77:1822–32.

7. Cheng RH, Caston JR, Wang G-j, Gu F, Smith TJ, Baker TS, Bozarth RF, Trus BL, Cheng N, Wickner RB. 1994. Fungal virus capsids, cytoplasmic compartments for the replication of double-stranded RNA, formed as icosahedral shells of asymmetric Gag dimers. Elsevier.

8. Icho T, Wickner RB. 1989. The double-stranded RNA genome of yeast virus L-A encodes its own putative RNA polymerase by fusing two open reading frames. J Biol Chem 264:6716–23.

9. Hopper JE, Bostian K, Rowe L, Tipper D. 1977. Translation of the L-species dsRNA genome of the killer-associated virus-like particles of Saccharomyces cerevisiae. Journal of Biological Chemistry 252:9010–9017.

10. Fujimura T, Wickner RB. 1988. Gene overlap results in a viral protein having an RNA binding domain and a major coat protein domain. Cell 55:663–671.

11. Dinman JD, Icho T, Wickner RB. 1991. A −1 ribosomal frameshift in a double-stranded RNA virus of yeast forms a gag-pol fusion protein. Proc Natl Acad Sci U S A 88:174–8.

12. Van Alfen NK, Jaynes RA, Anagnostakis SL, Day PR. 1975. Chestnut Blight: Biological Control by Transmissible Hypovirulence in Endothia parasitica. Science 189:890–1.

13. Choi GH, Nuss DL. 1992. Hypovirulence of chestnut blight fungus conferred by an infectious viral cDNA. Science 257:800–3.

14. Ghabrial SA, Suzuki N. 2009. Viruses of plant pathogenic fungi. Annu Rev Phytopathol 47:353–84.

15. Segers GC, Zhang X, Deng F, Sun Q, Nuss DL. 2007. Evidence that RNA silencing functions as an antiviral defense mechanism in fungi. Proc Natl Acad Sci U S A 104:12902–6.

16. Zhang X, Segers GC, Sun Q, Deng F, Nuss DL. 2008. Characterization of hypovirus-derived small RNAs generated in the chestnut blight fungus by an inducible DCL-2-dependent pathway. J Virol 82:2613–9.

17. Hammond TM, Andrewski MD, Roossinck MJ, Keller NP. 2008. Aspergillus mycoviruses are targets and suppressors of RNA silencing. Eukaryot Cell 7:350–7.

18. Yaegashi H, Shimizu T, Ito T, Kanematsu S. 2016. Differential Inductions of RNA Silencing among Encapsidated Double-Stranded RNA Mycoviruses in the White Root Rot Fungus Rosellinia necatrix. J Virol 90:5677–92.

19. Yu J, Lee KM, Cho WK, Park JY, Kim KH. 2018. Differential Contribution of RNA Interference Components in Response to Distinct Fusarium graminearum Virus Infections. J Virol 92.

20. Findley K, Oh J, Yang J, Conlan S, Deming C, Meyer JA, Schoenfeld D, Nomicos E, Park M, Program NIHISCCS, Kong HH, Segre JA. 2013. Topographic diversity of fungal and bacterial communities in human skin. Nature 498:367–70.

21. Clavaud C, Jourdain R, Bar-Hen A, Tichit M, Bouchier C, Pouradier F, El Rawadi C, Guillot J, Ménard-Szczebara F, Breton L. 2013. Dandruff is associated with disequilibrium in the proportion of the major bacterial and fungal populations colonizing the scalp. PloS one 8:e58203.

22. Patino-Uzcategui A, Amado Y, Cepero de Garcia M, Chaves D, Tabima J, Motta A, Cardenas M, Bernal A, Restrepo S, Celis A. 2011. Virulence gene expression in Malassezia spp from individuals with seborrheic dermatitis. J Invest Dermatol 131:2134–6.

23. Zhang E, Tanaka T, Tajima M, Tsuboi R, Nishikawa A, Sugita T. 2011. Characterization of the skin fungal microbiota in patients with atopic dermatitis and in healthy subjects. Microbiol Immunol 55:625–32.

24. Park T, Kim HJ, Myeong NR, Lee HG, Kwack I, Lee J, Kim BJ, Sul WJ, An S. 2017. Collapse of human scalp microbiome network in dandruff and seborrhoeic dermatitis. Experimental dermatology 26:835–838.

25. Wang L, Clavaud C, Bar⏻Hen A, Cui M, Gao J, Liu Y, Liu C, Shibagaki N, Guéniche A, Jourdain R. 2015. Characterization of the major bacterial–fungal populations colonizing dandruff scalps in Shanghai, China, shows microbial disequilibrium. Experimental dermatology 24:398–400.

26. Xu Z, Wang Z, Yuan C, Liu X, Yang F, Wang T, Wang J, Manabe K, Qin O, Wang X. 2016. Dandruff is associated with the conjoined interactions between host and microorganisms. Scientific reports 6:srep24877.

27. Grice EA, Dawson TL, Jr. 2017. Host-microbe interactions: Malassezia and human skin. Curr Opin Microbiol 40:81–87.

28. Sharma S, Gupta S, Shrivastava JN. 2011. Presence of Virus like Particles in Human Pathogenic Fungi: Chrysosporium sps and Candida albicans. Indian J Virol 22:104–10.

29. Park M, Cho YJ, Lee YW, Jung WH. 2017. Whole genome sequencing analysis of the cutaneous pathogenic yeast Malassezia restricta and identification of the major lipase expressed on the scalp of patients with dandruff. Mycoses 60:188–197.

30. Edy VG, Szekely M, Loviny T, Dreyer C. 1976. Action of Nucleases on Double◻Stranded RNA. European journal of biochemistry 61:563–572.

31. Lee SH, Yun S-H, Chun J, Kim D-H. 2017. Characterization of a novel dsRNA mycovirus of Trichoderma atroviride NFCF028. Archives of virology 162:1073–1077.

32. Sommer SS, Wickner RB. 1982. Yeast L dsRNA consists of at least three distinct RNAs; evidence that the non-Mendelian genes [HOK],[NEX] and [EXL] are on one of these dsRNAs. Cell 31:429–441.

33. Park CM, Lopinski JD, Masuda J, Tzeng TH, Bruenn JA. 1996. A second double-stranded RNA virus from yeast. Virology 216:451–4.

34. Taylor DJ, Ballinger MJ, Bowman SM, Bruenn JA. 2013. Virus-host co-evolution under a modified nuclear genetic code. PeerJ 1:e50.

35. Bruenn JA. 1993. A closely related group of RNA-dependent RNA polymerases from double-stranded RNA viruses. Nucleic Acids Res 21:5667–9.

36. Schmitt MJ, Breinig F. 2002. The viral killer system in yeast: from molecular biology to application. FEMS microbiology reviews 26:257–276.

37. Wickner RB, Leibowitz MJ. 1979. Mak mutants of yeast: mapping and characterization. J Bacteriol 140:154–60.

38. Tercero J, Dinman J, Wickner R. 1993. Yeast MAK3 N-acetyltransferase recognizes the N-terminal four amino acids of the major coat protein (gag) of the LA double-stranded RNA virus. Journal of bacteriology 175:3192–3194.

39. Tercero J, Wickner RB. 1992. MAK3 encodes an N-acetyltransferase whose modification of the LA gag NH2 terminus is necessary for virus particle assembly. Journal of Biological Chemistry 267:20277–20281.

40. Fujimura T, Wickner RB. 1987. LA double-stranded RNA viruslike particle replication cycle in Saccharomyces cerevisiae: particle maturation in vitro and effects of mak10 and pet18 mutations. Molecular and cellular biology 7:420–426.

41. Lee Y-J, Wickner R. 1992. MAK10, a glucose-repressible gene necessary for replication of a dsRNA virus of Saccharomyces cerevisiae, has T cell receptor alpha-subunit motifs. Genetics 132:87–96.

42. Wickner RB, Ridley SP, Fried HM, Ball SG. 1982. Ribosomal protein L3 is involved in replication or maintenance of the killer double-stranded RNA genome of Saccharomyces cerevisiae. Proceedings of the National Academy of Sciences 79:4706–4708.

43. Ohtake Y, Wickner RB. 1995. Yeast virus propagation depends critically on free 60S ribosomal subunit concentration. Molecular and Cellular Biology 15:2772–2781.

44. Ryu S-W, Lee S-J, Park M-Y, Jun J-i, Jung Y-K, Kim E. 2003. Fas-associated factor 1, FAF1, is a member of Fas death-inducing signaling complex. Journal of Biological Chemistry 278:24003–24010.

45. Fujikura D, Miyazaki T. 2018. Programmed Cell Death in the Pathogenesis of Influenza. Int J Mol Sci 19.

46. Jorgensen I, Rayamajhi M, Miao EA. 2017. Programmed cell death as a defence against infection. Nat Rev Immunol 17:151–164.

47. Carmona-Gutierrez D, Bauer MA, Zimmermann A, Aguilera A, Austriaco N, Ayscough K, Balzan R, Bar-Nun S, Barrientos A, Belenky P. 2018. Guidelines and recommendations on yeast cell death nomenclature. Microbial Cell 5:4.

48. Newbold A, Martin BP, Cullinane C, Bots M. 2014. Detection of apoptotic cells using propidium iodide staining. Cold Spring Harb Protoc 2014:1202–6.

49. Banks GT, Buck KW, Chain EB, Himmelweit F, Marks JE, Tyler JM, Hollings M, Last FT, Stone OM. 1968. Viruses in fungi and interferon stimulation. Nature 218:542–5.

50. Kleinschmidt WJ, Cline JC, Murphy EB. 1964. Interferon Production Induced by Statolon. Proc Natl Acad Sci U S A 52:741–4.

51. Lampson GP, Tytell AA, Field AK, Nemes MM, Hilleman MR. 1967. Inducers of interferon and host resistance. I. Double-stranded RNA from extracts of Penicillium funiculosum. Proc Natl Acad Sci U S A 58:782–9.

52. Perales-Linares R, Navas-Martin S. 2013. Toll-like receptor 3 in viral pathogenesis: friend or foe? Immunology 140:153–67.

53. Thompson MR, Kaminski JJ, Kurt-Jones EA, Fitzgerald KA. 2011. Pattern recognition receptors and the innate immune response to viral infection. Viruses 3:920–40.

54. Claudepierre MC, Hortelano J, Schaedler E, Kleinpeter P, Geist M, Remy-Ziller C, Brandely R, Tosch C, Laruelle L, Jawhari A, Menguy T, Marchand JB, Romby P, Schultz P, Hartmann G, Rooke R, Bonnefoy JY, Preville X, Rittner K. 2014. Yeast virus-derived stimulator of the innate immune system augments the efficacy of virus vector-based immunotherapy. J Virol 88:5242–55.

55. Ishikawa T, Itoh F, Yoshida S, Saijo S, Matsuzawa T, Gonoi T, Saito T, Okawa Y, Shibata N, Miyamoto T, Yamasaki S. 2013. Identification of distinct ligands for the C-type lectin receptors Mincle and Dectin-2 in the pathogenic fungus Malassezia. Cell Host Microbe 13:477–88.

56. Vargas G, Rocha JD, Oliveira DL, Albuquerque PC, Frases S, Santos SS, Nosanchuk JD, Gomes AM, Medeiros LC, Miranda K, Sobreira TJ, Nakayasu ES, Arigi EA, Casadevall A, Guimaraes AJ, Rodrigues ML, Freire-de-Lima CG, Almeida IC, Nimrichter L. 2015. Compositional and immunobiological analyses of extracellular vesicles released by Candida albicans. Cell Microbiol 17:389–407.

57. Samuel CE. 2001. Antiviral actions of interferons. Clinical microbiology reviews 14:778–809.

58. Brierley I, Boursnell ME, Binns MM, Bilimoria B, Blok VC, Brown TD, Inglis SC. 1987. An efficient ribosomal frame-shifting signal in the polymerase-encoding region of the coronavirus IBV. EMBO J 6:3779–85.

59. Jacks T, Varmus HE. 1985. Expression of the Rous sarcoma virus pol gene by ribosomal frameshifting. Science 230:1237–42.

60. Jacks T, Madhani HD, Masiarz FR, Varmus HE. 1988. Signals for ribosomal frameshifting in the Rous sarcoma virus gag-pol region. Cell 55:447–58.

61. Magliani W, Conti S, Gerloni M, Bertolotti D, Polonelli L. 1997. Yeast killer systems. Clinical microbiology reviews 10:369–400.

62. Tercero JC, Riles LE, Wickner RB. 1992. Localized mutagenesis and evidence for post-transcriptional regulation of MAK3. A putative N-acetyltransferase required for double-stranded RNA virus propagation in Saccharomyces cerevisiae. J Biol Chem 267:20270–6.

63. Wickner RB. 1996. Double-stranded RNA viruses of Saccharomyces cerevisiae. Microbiological reviews 60:250.

64. Thrash C, Bankier AT, Barrell BG, Sternglanz R. 1985. Cloning, characterization, and sequence of the yeast DNA topoisomerase I gene. Proceedings of the National Academy of Sciences 82:4374–4378.

65. Widner WR, Wickner RB. 1993. Evidence that the SKI antiviral system of Saccharomyces cerevisiae acts by blocking expression of viral mRNA. Molecular and cellular biology 13:4331–4341.

66. Masison DC, Blanc A, Ribas JC, Carroll K, Sonenberg N, Wickner RB. 1995. Decoying the cap-mRNA degradation system by a double-stranded RNA virus and poly (A)-mRNA surveillance by a yeast antiviral system. Molecular and Cellular Biology 15:2763–2771.

67. Benard L, Carroll K, Valle RC, Masison DC, Wickner RB. 1999. The ski7 antiviral protein is an EF1-α homolog that blocks expression of non-Poly (A) mRNA in Saccharomyces cerevisiae. Journal of virology 73:2893–2900.

68. McBride RC, Boucher N, Park DS, Turner PE, Townsend JP. 2013. Yeast response to LA virus indicates coadapted global gene expression during mycoviral infection. FEMS yeast research 13:162–179.

69. Wu M, Jin F, Zhang J, Yang L, Jiang D, Li G. 2012. Characterization of a novel bipartite double-stranded RNA mycovirus conferring hypovirulence in the phytopathogenic fungus Botrytis porri. J Virol 86:6605–19.

70. Wan J, Basu K, Mui J, Vali H, Zheng H, Laliberte JF. 2015. Ultrastructural Characterization of Turnip Mosaic Virus-Induced Cellular Rearrangements Reveals Membrane-Bound Viral Particles Accumulating in Vacuoles. J Virol 89:12441–56.

71. Matsumoto M, Oshiumi H, Seya T. 2011. Antiviral responses induced by the TLR3 pathway. Reviews in Medical Virology 21:67–77.

72. Tabeta K, Georgel P, Janssen E, Du X, Hoebe K, Crozat K, Mudd S, Shamel L, Sovath S, Goode J. 2004. Toll-like receptors 9 and 3 as essential components of innate immune defense against mouse cytomegalovirus infection. Proceedings of the National Academy of Sciences 101:3516–3521.

73. Hartley MA, Ronet C, Zangger H, Beverley SM, Fasel N. 2012. Leishmania RNA virus: when the host pays the toll. Frontiers in Cellular and Infection Microbiology 2.

74. Shelly Applen Clancey FR, Salomé LeibundGut-Landmann, Joseph Heitman, Giuseppe Ianiri. 2019. A novel mycovirus evokes transcriptional rewiring in Malassezia and provokes host inflammation and an immunological response. bioRxiv doi:https://doi.org/10.1101/2019.12.18.880518.

75. Kim M, Cho Y-J, Park M, Choi Y, Hwang SY, Jung WH. 2018. Genomic tandem quadruplication is associated with ketoconazole resistance in Malassezia pachydermatis. J Microbiol Biotechnol 28:1937–1945.

76. Park M, Jung WH, Han SH, Lee YH, Lee YW. 2015. Characterisation and Expression Analysis of MrLip1, a Class 3 Family Lipase of Malassezia restricta. Mycoses 58:671–8.

77. Leeming JP, Notman FH. 1987. Improved methods for isolation and enumeration of Malassezia furfur from human skin. J Clin Microbiol 25:2017–9.

78. Okada R, Kiyota E, Moriyama H, Fukuhara T, Natsuaki T. 2015. A simple and rapid method to purify viral dsRNA from plant and fungal tissue. Journal of General Plant Pathology 81:103–107.

79. Moon S, Byun Y, Kim HJ, Jeong S, Han K. 2004. Predicting genes expressed via −1 and +1 frameshifts. Nucleic Acids Res 32:4884–92.

80. Waterhouse AM, Procter JB, Martin DM, Clamp M, Barton GJ. 2009. Jalview Version 2--a multiple sequence alignment editor and analysis workbench. Bioinformatics 25:1189–91.

81. Kumar S, Nei M, Dudley J, Tamura K. 2008. MEGA: a biologist-centric software for evolutionary analysis of DNA and protein sequences. Brief Bioinform 9:299–306.

82. Kumar S, Stecher G, Li M, Knyaz C, Tamura K. 2018. MEGA X: Molecular Evolutionary Genetics Analysis across Computing Platforms. Mol Biol Evol 35:1547–1549.

83. Vargas G, Rocha JD, Oliveira DL, Albuquerque PC, Frases S, Santos SS, Nosanchuk JD, Gomes AMO, Medeiros LC, Miranda K. 2015. Compositional and immunobiological analyses of extracellular vesicles released by C andida albicans. Cellular microbiology 17:389–407.

84. Koh H-J, Kim Y-R, Kim J-S, Yun J-S, Kim S, Kim SY, Jang K, Yang C-S. 2018. CD82 hypomethylation is essential for tuberculosis pathogenesis via regulation of RUNX1-Rab5/22. Experimental & molecular medicine 50:62.

85. Saitou N, Nei M. 1987. The neighbor-joining method: a new method for reconstructing phylogenetic trees. Mol Biol Evol 4:406–25.

86. Felsenstein J. 1985. Confidence Limits on Phylogenies: An Approach Using the Bootstrap. Evolution 39:783–791.

87. Sanderlin RS, Ghabrial SA. 1978. Physicochemical properties of two distinct types of virus-like particles from Helminthosporium victoriae. Virology 87:142–51.

88. Shapira R, Choi GH, Nuss DL. 1991. Virus-like genetic organization and expression strategy for a double-stranded RNA genetic element associated with biological control of chestnut blight. EMBO J 10:731–9.

89. Hillman BI, Halpern BT, Brown MP. 1994. A viral dsRNA element of the chestnut blight fungus with a distinct genetic organization. Virology 201:241–50.

90. Van der Lende TR, Duitman EH, Gunnewijk MG, Yu L, Wessels JG. 1996. Functional analysis of dsRNAs (L1, L3, L5, and M2) associated with isometric 34-nm virions of Agaricus bisporus (white button mushroom). Virology 217:88–96.

91. Nomura K, Osaki H, Iwanami T, Matsumoto N, Ohtsu Y. 2003. Cloning and characterization of a totivirus double-stranded RNA from the plant pathogenic fungus, Helicobasidium mompa Tanaka. Virus Genes 26:219–26.

92. Tuomivirta TT, Hantula J. 2003. Two unrelated double-stranded RNA molecule patterns in Gremmeniella abietina type A code for putative viruses of the families Totiviridae and Partitiviridae. Arch Virol 148:2293–305.

93. Jiang D, Ghabrial SA. 2004. Molecular characterization of Penicillium chrysogenum virus: reconsideration of the taxonomy of the genus Chrysovirus. J Gen Virol 85:2111–21.

94. Urayama S, Ohta T, Onozuka N, Sakoda H, Fukuhara T, Arie T, Teraoka T, Moriyama H. 2012. Characterization of Magnaporthe oryzae chrysovirus 1 structural proteins and their expression in Saccharomyces cerevisiae. J Virol 86:8287–95.

95. Wu Q, Luo Y, Lu R, Lau N, Lai EC, Li WX, Ding SW. 2010. Virus discovery by deep sequencing and assembly of virus-derived small silencing RNAs. Proc Natl Acad Sci U S A 107:1606–11.

96. Isawa H, Kuwata R, Hoshino K, Tsuda Y, Sakai K, Watanabe S, Nishimura M, Satho T, Kataoka M, Nagata N, Hasegawa H, Bando H, Yano K, Sasaki T, Kobayashi M, Mizutani T, Sawabe K. 2011. Identification and molecular characterization of a new nonsegmented double-stranded RNA virus isolated from Culex mosquitoes in Japan. Virus Res 155:147–55.

97. Mor SK, Phelps NB. 2016. Molecular detection of a novel totivirus from golden shiner (Notemigonus crysoleucas) baitfish in the USA. Arch Virol 161:2227–34.

98. Yang X, Zhang Y, Ge X, Yuan J, Shi Z. 2012. A novel totivirus-like virus isolated from bat guano. Arch Virol 157:1093–9.

99. Zhai Y, Attoui H, Mohd Jaafar F, Wang HQ, Cao YX, Fan SP, Sun YX, Liu LD, Mertens PP, Meng WS, Wang D, Liang G. 2010. Isolation and full-length sequence analysis of Armigeres subalbatus totivirus, the first totivirus isolate from mosquitoes representing a proposed novel genus (Artivirus) of the family Totiviridae. J Gen Virol 91:2836–45.

100. Dihanich M, Van Tuinen E, Lambris J, Marshallsay B. 1989. Accumulation of viruslike particles in a yeast mutant lacking a mitochondrial pore protein. Molecular and cellular biology 9:1100–1108.

101. Leibowitz MJ, Wickner RB. 1978. pet 18: A chromosomal gene required for cell growth and for the maintenance of mitochondrial DNA and the killer plasmid of yeast. Molecular and General Genetics MGG 165:115–121.

102. Toh◻e A, Sahashi Y. 1985. The PET18 locus of Saccharomyces cerevisiae: a complex locus containing multiple genes. Yeast 1:159–171.

103. Thrash C, Voelkel K, DiNardo S, Sternglanz R. 1984. Identification of Saccharomyces cerevisiae mutants deficient in DNA topoisomerase I activity. Journal of Biological Chemistry 259:1375–1377.

104. Icho T, Wickner RB. 1988. The MAK11 protein is essential for cell growth and replication of M double-stranded RNA and is apparently a membrane-associated protein. Journal of Biological Chemistry 263:1467–1475.

105. Wickner RB. 1988. Host function of MAK16: G1 arrest by a mak16 mutant of Saccharomyces cerevisiae. Proceedings of the National Academy of Sciences 85:6007–6011.

106. Carroll K, Wickner RB. 1995. Translation and M1 double-stranded RNA propagation: MAK18= RPL41B and cycloheximide curing. Journal of bacteriology 177:2887–2891.

107. Toh-E A, Guerry P, Wickner RB. 1978. Chromosomal superkiller mutants of Saccharomyces cerevisiae. Journal of bacteriology 136:1002–1007.

108. Ball SG, Tirtiaux C, Wickner RB. 1984. Genetic control of LA and L-(BC) dsRNA copy number in killer systems of Saccharomyces cerevisiae. Genetics 107:199–217.

109. Ridley SP, Sommer SS, Wickner RB. 1984. Superkiller mutations in Saccharomyces cerevisiae suppress exclusion of M2 double-stranded RNA by LA-HN and confer cold sensitivity in the presence of M and LA-HN. Molecular and Cellular Biology 4:761–770.

110. Dmochowska A, Dignard D, Henning D, Thomas DY, Bussey H. 1987. Yeast KEX1 gene encodes a putative protease with a carboxypeptidase B-like function involved in killer toxin and α-factor precursor processing. Cell 50:573–584.

111. Wickner RB. 1974. Chromosomal and nonchromosomal mutations affecting the “killer character” of Saccharomyces cerevisiae. Genetics 76:423–432.

112. Wickner RB, Leibowitz MJ. 1976. Two chromosomal genes required for killing expression in killer strains of Saccharomyces cerevisiae. Genetics 82:429–442.

