## Supplemental Materials and Methods for "A novel virus alters gene expression and vacuolar morphology in *Malassezia* cells and induces a TLR3-mediated inflammatory immune response"

### Cell fixation

For TEM analysis of *M. restricta* cells, a previously reported method was used with slight modifications (1). Briefly, cells ($1 \times 10^8$ cells) of virus-infected and virus-cured strains KCTC 27540 and KCTC 27527 cultured for 3 days were fixed in 2.5% glutaraldehyde in 0.1 M sodium phosphate buffer (pH 7.4) at 4°C overnight. After washing three times for 5 min each time in 0.1 M sodium phosphate buffer (pH 7.4) at room temperature (RT), the samples were post-fixed in 1% osmium tetroxide in 0.1 M sodium phosphate buffer (pH 7.4) at 4°C for 1 h. The samples were washed with distilled water three times at RT for 5 min and stained with 0.5% uranyl acetate at 4°C overnight. After washing three times for 5 min in distilled water at RT, the samples were gradually dehydrated with 30%, 50%, 70%, 80%, 90%, and 100% ethanol for 10 min each at RT. Washing with 100% ethanol was repeated twice. The samples were embedded with fresh Spurr's resin at 4°C overnight and subsequently at RT for 3 h and then embedded with the resin and polymerized at 70°C overnight. After polymerization, ultrathin (70-nm-thick) sections were prepared.

### Transmission electron microscopy (TEM)

Purified virus particles and fixed 70-nm-thick sections were observed by TEM. The samples were mounted on a 200-mesh copper grid, and the grid was stained with 2% uranyl acetate solution and examined using Talos L120C (FEI, Hillsboro, OR, USA).

### cDNA synthesis

The total nucleic acids from *M. restricta* were extracted as described previously (2). To isolate dsRNA, the total nucleic acids were digested with RNase T1 (Sigma, St. Louis, MO, USA) and DNase I (Thermo Fisher Scientific, Waltham, MA, USA). DsRNA was extracted from the agarose gel and used as a template for cDNA synthesis. The cDNA was synthesized according to the random RT-PCR method described by Darissa *et al.* with slight modifications (3). One hundred nanograms of dsRNA was mixed with 2 μM of the Tagged $dN_6$_1 primer and 12.5% (v/v) dimethyl sulfoxide, incubated at 98°C for 5 min, and chilled on ice for 5 min. Two hundred units of RevertAid™ Reverse Transcriptase (Thermo Fisher Scientific, Waltham, MA, USA), 20 units of RiboLock™ RNase Inhibitor (Thermo Fisher Scientific, Waltham, MA, USA), 1 mM dNTPs, 50 mM Tris–HCl (pH 8.3), 50 mM KCl, 4 mM $MgCl_2$, and 10 mM dithiothreitol were added. The mixture was incubated at 45°C for 1 h.

**RNA isolation and transcriptome analysis by RNA sequencing**

*M. restricta* KCTC 27540 cells containing virus were cultured in LNA medium at 34°C for 4 days and harvested for RNA extraction (see Fig. S2 in supplemental material). The virus in the same strain was cured by serially passaging the fungal cells eight times in LNA medium (each culture was incubated at 34°C for 4 days). The obtained virus-cured *M. restricta* cells were cultured in LNA medium at 34°C for 4 days, and total RNA was extracted using a RNeasy Mini Kit (Qiagen, Hilden, Germany). The quality of RNA was evaluated using a TapeStation RNA ScreenTape (Agilent Technologies, Santa Clara, CA, USA). Libraries for RNA sequencing were prepared using high-quality RNA with RNA integrity values greater than 7.0 and a TruSeq Stranded mRNA Sample Prep Kit (Illumina, San Diego, CA, USA) following the manufacturer's protocol. The libraries were evaluated on an Illumina HiSeq2500 platform, and

100-bp paired-end reads were generated and sequenced. Generated raw reads that passed quality filters were mapped to the reference genome of *M. restricta* KCTC 27527 using bowtie2 v2.2.1 (4, 5). Mapped reads were counted by featureCounts in Subread package v1.4.3 and the relative transcript abundance was TPM-normalized (6). Differential expression of the selected genes was confirmed by quantitative real time PCR (Q-RT PCR; Fig. S3).

**Q-RT PCR**

To validate the transcriptome data, total RNA was extracted from the virus-infected and the virus-cured *M. restricta* KCTC 27540 cells using an RNeasy Mini Kit (Qiagen, Hilden, Germany) and cDNA was synthesized using the RevertAid™ First Strand cDNA synthesis Kit (Fermentas, Waltham, MA, USA). Relative gene expression was quantified using primers listed in Table S3 and a 7500 system (Applied Biosystems, Foster City, CA, USA) based on the $2^{-\triangle\triangle C_T}$ method. The actin gene (MRET_1518) was used as a reference.

To evaluate TLR expression, total RNA was extracted from the BMDCs using an RNeasy Mini Kit (Qiagen) and used for cDNA synthesis. Q-RT PCR was performed using gene-specific primer sets and a QuantStudio™ 3 (Applied Biosystems, Foster City, CA, USA) according to the manufacturer's instructions (see Table S3 in supplemental material). Relative expression was calculated using by the $2^{-\Delta\Delta CT}$ method and normalized against β-actin expression (7). PBS and lipopolysaccharide served as negative and positive controls, respectively.

**Flow cytometric analysis**

The virus-infected and the virus-cured *M. restricta* KCTC 27540 cells were grown at 34 °C for

4 days, and $5\times10^7$ cells/mL of each strain was suspended in 3 mL of mDixon's medium followed by incubation at 34℃ for 6 h. Cells were stained with Annexin V and propidium iodide (PI) using an EzWay Annexin V-FITC Apoptosis Detection Kit (Komabiotech, Korea) following the manufacturer's protocol. Samples were diluted 20-fold in cold PBS, then directly subjected to flow cytometry using CyFlow® Cube6 (Sysmex, Japan) at a wave-length of 488 nm. The data from 50,000 cells of each strain were collected at the FL1 channel for Annexin V and at the FL3 channel for PI, and analyzed.

**Heterologous expression and purification of the capsid protein**

For heterologous expression of the capsid protein, the cDNA of ORF1 was amplified using the primers MrV40L.CP.F_BamHI and MrV40L.CP.R_HindIII (see Table S2 in supplemental materials). The amplified PCR products were ligated into the BamHI and HindIII sites of the plasmid pET28a(+) (Novagen, Madison, WI, USA). To express the proteins, the constructed plasmids were transformed into *E. coli* BL21 cells, which were cultured at 10°C overnight in the presence of 0.5 mM isopropyl β-D-1-thiogalactopyranoside. The cells were lysed by sonication, and protein was purified using a His GraviTrap™ column (GE Healthcare, Little Chalfont, UK) (see Fig. S5 in supplemental materials). The purified protein was used for co-culture with BMDCs.

**Heterologous expression and purification of the MrV40S satellite proteins**

For heterologous expression of the protein produced from MrV40S, the cDNA of ORF3 was amplified using the primers and MrV40S.F_BamHI and MrV40S.R_XbaI (see Table S2 in supplemental material). The amplified PCR product was ligated into the BamHI and XbaI sites of the plasmid pET28a(+) (Novagen, Madison, WI, USA). To express the proteins, the constructed plasmids were transformed into *E. coli* BL21, which were cultured at 10°C overnight in the presence of 0.5 mM isopropyl β-D-1-thiogalactopyranoside. The cells were lysed by sonication and the protein was purified using His GraviTrap™ columns (GE Healthcare, Little Chalfont, UK).

**Preparation of fungal cell lysates**

The *M. restricta* strains were grown on LNA plates for 3 days, harvested, washed twice with PBS, and resuspended in PBS. Glass beads were added into the cell suspension, and total fungal cell lysates were prepared by vortexing. The resulting fungal cell lysates were used for co-culture with BMDCs.

**References**

1. Byers B, Goetsch L. 1991. [41] Preparation of yeast cells for thin-section electron microscopy, p 602-608, Methods in enzymology, vol 194. Elsevier.
2. van Burik JA, Schreckhise RW, White TC, Bowden RA, Myerson D. 1998. Comparison of six extraction techniques for isolation of DNA from filamentous fungi. Med Mycol 36:299-303.
3. Darissa O, Willingmann P, Adam G. 2010. Optimized approaches for the sequence determination of double-stranded RNA templates. J Virol Methods 169:397-403.
4. Cho Y-J, Park M, Jung WH. 2019. Resequencing the genome of Malassezia restricta

strain KCTC 27527. Microbiol Resour Announc 8:e00213-19.

5. Langmead B, Salzberg SL. 2012. Fast gapped-read alignment with Bowtie 2. Nat Methods 9:357-9.

6. Liao Y, Smyth GK, Shi W. 2013. The Subread aligner: fast, accurate and scalable read mapping by seed-and-vote. Nucleic Acids Res 41:e108.

7. Livak KJ, Schmittgen TD. 2001. Analysis of relative gene expression data using real-time quantitative PCR and the 2− ΔΔCT method. methods 25:402-408.
