## Supplementary material for "A novel virus alters gene expression and vacuolar morphology in *Malassezia* cells and induces a TLR3-mediated inflammatory immune response": table S1

**Table S1. Expression of genes calculated from RNA-seq data**

| Gene ID | Annotation | Reads (TPM) | |
|---|---|---|---|
| | | +Virus | -Virus |
| MRET_0001 | uncharacterized protein | 0 | 0 |
| MRET_0002 | iron transport multicopper oxidase | 0 | 0 |
| MRET_0003 | carboxypeptidase D | 41.14 | 75.11 |
| MRET_0004 | Csr1-phosphatidylinositol transfer protein | 640.09 | 634.52 |
| MRET_0005 | Bromodomain associated protein | 25.39 | 48.84 |
| MRET_0006 | ATP-dependent RNA helicase SUPV3L1/SUV3 | 31.01 | 52.81 |
| MRET_0007 | diphthine-ammonia ligase | 8.97 | 16.23 |
| MRET_0008 | oligosaccharyltransferase complex subunit beta | 42.48 | 77.19 |
| MRET_0009 | pheromone-dependent cell cycle arrest protein Far11 | 27.31 | 59.25 |
| MRET_0010 | mitochondrial distribution and morphology protein 10 | 53.53 | 60.18 |
| MRET_0011 | pre-mRNA-processing factor 8 | 317.53 | 248.21 |
| MRET_0012 | DUF410 domain protein | 116.97 | 363.53 |
| MRET_0013 | acetyltransferase (GNAT) family | 886.71 | 746.57 |
| MRET_0014 | small nuclear ribonucleoprotein G | 136.09 | 136.61 |
| MRET_0015 | metal homeostatis protein BSD2 | 97.2 | 363.97 |
| MRET_0016 | uncharacterized protein | 205.34 | 226.76 |
| MRET_0017 | DnaJ domain protein | 595.35 | 439.68 |
| MRET_0018 | 6-phosphofructokinase 1 | 107.33 | 105.28 |
| MRET_0019 | lipase | 3678.04 | 10249.18 |
| MRET_0020 | uncharacterized protein | 73.69 | 139.31 |
| MRET_0021 | cytoplasmic GTPase-activating protein | 18.59 | 58.32 |
| MRET_0022 | uncharacterized protein | 15.86 | 36.13 |
| MRET_0023 | U4/U6 small nuclear ribonucleoprotein SNU13 | 154.25 | 540.65 |
| MRET_0024 | DnaJ homolog subfamily B member 4 | 1343.75 | 1798.91 |
| MRET_0025 | AAA family ATPase | 47.17 | 88.2 |
| MRET_0026 | PHD finger domain protein | 10.69 | 48.08 |
| MRET_0027 | pre-mRNA-splicing factor ATP-dependent RNA helicase DHX16 | 12.58 | 17.89 |
| MRET_0028 | RING-H2 domain core subunit of multiple ubiquitin ligase complexes | 1039.03 | 886.43 |
| MRET_0029 | elongator complex protein 2 | 51.24 | 54.29 |
| MRET_0030 | leucine-rich repeat protein | 22.33 | 30.33 |
| MRET_0031 | helicase associated domain (HA2) containing protein | 30.68 | 70.09 |
| MRET_0032 | kinetochore protein Mis13/DSN1 | 15.49 | 41.56 |
| MRET_0033 | THO complex subunit 3 | 365.31 | 429.76 |
| MRET_0034 | microtubule-associated protein, RP/EB family | 39.38 | 163.24 |

| | | | |
|---|---|---|---|
| MRET_0035 | cysteine desulfurase | 1059.79 | 1001.13 |
| MRET_0036 | optic atrophy 3 protein (OPA3) | 107.16 | 253.76 |
| MRET_0037 | polyadenylate-binding protein 2 | 139.53 | 357.88 |
| MRET_0038 | 26 proteasome complex subunit DSS1 | 235.2 | 452.65 |
| MRET_0039 | oligosaccharyltransferase complex subunit alpha (ribophorin I) | 77.09 | 74.43 |
| MRET_0040 | Tic20-like protein | 42.86 | 60.56 |
| MRET_0041 | rRNA-processing protein CGR1 | 82.16 | 248.54 |
| MRET_0042 | NADH dehydrogenase (ubiquinone) Fe-S protein 6 | 226.47 | 534.7 |
| MRET_0043 | pheromone-regulated membrane protein | 318.41 | 171.24 |
| MRET_0044 | transcription initiation factor TFIIH subunit 2 | 90.72 | 138.64 |
| MRET_0045 | protein transport protein SEC31 | 375.03 | 165.41 |
| MRET_0046 | vacuolar protein sorting-associated protein 18 | 58.11 | 50.48 |
| MRET_0047 | triose/dihydroxyacetone kinase/FAD-AMP lyase (cyclizing) | 1609.71 | 1305.02 |
| MRET_0048 | mitochondrial DNA replication protein | 54.54 | 99.03 |
| MRET_0049 | ESCRT-II complex subunit VPS22 | 132.98 | 218.96 |
| MRET_0050 | homeobox domain protein | 249.46 | 400.45 |
| MRET_0051 | ubiquinone biosynthesis monooxygenase Coq7 | 434.37 | 294.57 |
| MRET_0052 | uncharacterized protein | 78.9 | 127.91 |
| MRET_0053 | 2-(3-amino-3-carboxypropyl)histidine synthase | 127.44 | 124.66 |
| MRET_0054 | pyruvate dehydrogenase protein X component, mitochondrial precursor | 609.73 | 442.64 |
| MRET_0055 | palmitoyltransferase ZDHHC13/17 | 107.28 | 68.46 |
| MRET_0056 | uncharacterized protein | 24.84 | 48.55 |
| MRET_0057 | large subunit ribosomal protein L34 | 148.95 | 126.91 |
| MRET_0058 | DNA-binding protein HGH1 | 40.62 | 65.52 |
| MRET_0059 | NTF2-related export protein 1/2 | 467.57 | 417.02 |
| MRET_0060 | uncharacterized protein | 325.15 | 197.08 |
| MRET_0061 | solute carrier family 36 (proton-coupled amino acid transporter) | 57.58 | 114.74 |
| MRET_0062 | C-4 methylsterol oxidase | 564.39 | 498.99 |
| MRET_0063 | TBC1 domain family member 8/9 | 346.67 | 205.3 |
| MRET_0064 | uncharacterized protein | 56.26 | 49.76 |
| MRET_0065 | transcription elongation factor | 39.11 | 66.91 |
| MRET_0066 | SCY1-like protein 2 | 47.52 | 104.7 |
| MRET_0067 | protein YIH1 | 102.06 | 210.81 |
| MRET_0068 | dehydrogenase | 168.04 | 375.42 |
| MRET_0069 | dehydrogenase | 62.07 | 119.21 |
| MRET_0070 | golgi reassembly stacking protein | 63.84 | 132.79 |
| MRET_0071 | glycoside hydrolase family 5 protein | 20.64 | 45.03 |

| | | | |
|---|---|---|---|
| MRET_0072 | protein of unknown function (DUF1769) | 93.34 | 141.99 |
| MRET_0073 | Bsp1 protein | 17.88 | 60.7 |
| MRET_0074 | uncharacterized protein | 31.79 | 152.64 |
| MRET_0075 | protein of unknown function (DUF2422) | 21.48 | 90.94 |
| MRET_0076 | uncharacterized protein | 108.32 | 93.63 |
| MRET_0077 | 4-coumarate-coA ligase | 201.03 | 163.55 |
| MRET_0078 | pyridoxamine 5'-phosphate oxidase | 308.01 | 345.85 |
| MRET_0079 | enhancer of yellow 2 transcription factor | 408.33 | 463.06 |
| MRET_0080 | LCCL domain protein | 16 | 26.94 |
| MRET_0081 | syntaxin-binding protein 5 | 76.69 | 53.03 |
| MRET_0082 | C2 domain protein | 247.29 | 88.97 |
| MRET_0083 | fungal Zn(2)-Cys(6) binuclear cluster domain protein | 274.99 | 269.52 |
| MRET_0084 | ataxin-3 | 1148.37 | 829.45 |
| MRET_0085 | RuvB-like protein 2 | 78.2 | 188.97 |
| MRET_0086 | uncharacterized protein | 3.92 | 11 |
| MRET_0087 | glutathione synthase | 33.82 | 33.08 |
| MRET_0088 | sphingolipid 4-desaturase/C4-monooxygenase | 90.22 | 58.85 |
| MRET_0089 | uncharacterized protein | 16.19 | 46.05 |
| MRET_0090 | NADH dehydrogenase (ubiquinone) flavoprotein 1 | 162.13 | 141.94 |
| MRET_0091 | SURF1-like protein | 22.26 | 41.84 |
| MRET_0092 | exocyst complex component 2 | 21.69 | 36.03 |
| MRET_0093 | 26S proteasome regulatory subunit N5 | 71.94 | 112.16 |
| MRET_0094 | adenosine kinase | 181.06 | 183.68 |
| MRET_0095 | transporter | 33.17 | 20.91 |
| MRET_0096 | uncharacterized protein | 128.65 | 95.02 |
| MRET_0097 | transcriptional regulation protein GF11 | 25.84 | 106.14 |
| MRET_0098 | aarF domain kinase | 18.97 | 45.26 |
| MRET_0099 | nucleoside-diphosphate kinase | 473.29 | 784.92 |
| MRET_0100 | uncharacterized protein | 35.77 | 65.25 |
| MRET_0101 | uncharacterized protein | 44.41 | 103.1 |
| MRET_0102 | glutathione peroxidase | 15418.96 | 13664.52 |
| MRET_0103 | ornithine carbamoyltransferase | 101.7 | 153.1 |
| MRET_0104 | small EDRK-rich factor | 244.58 | 241.61 |
| MRET_0105 | uncharacterized protein | 50.15 | 64.47 |
| MRET_0106 | protein FRG1 | 334.53 | 161.8 |
| MRET_0107 | 6-phosphofructo-2-kinase/fructose-2,6-biphosphatase 4 | 276.53 | 142.81 |
| MRET_0108 | uncharacterized protein | 78.53 | 109.79 |

| | | | |
|---|---|---|---|
| MRET_0109 | uncharacterized protein | 45.12 | 84.15 |
| MRET_0110 | DNA repair protein | 431.89 | 327.87 |
| MRET_0111 | 2-dehydropantoate 2-reductase | 44.28 | 45.64 |
| MRET_0112 | subunit of the DSC ubiquitin ligase complex | 40.33 | 54.21 |
| MRET_0113 | transcription initiation factor TFIID subunit 1 | 26.29 | 49.52 |
| MRET_0114 | TatD DNase family protein | 25.73 | 42.17 |
| MRET_0115 | uncharacterized protein | 118.34 | 832.44 |
| MRET_0116 | COP9 signalosome complex subunit 2 | 84.92 | 216.53 |
| MRET_0117 | mitochondrial organizing structure protein 2 | 211.68 | 332.09 |
| MRET_0118 | coiled-coil domain protein 130 | 37.43 | 95.66 |
| MRET_0119 | protein farnesyltransferase/geranylgeranyltransferase type-1 subunit alpha | 34.57 | 96.13 |
| MRET_0120 | uncharacterized protein | 568.87 | 1244.42 |
| MRET_0121 | uncharacterized protein | 658.24 | 1145.85 |
| MRET_0122 | uncharacterized protein | 709.81 | 1125.89 |
| MRET_0123 | cystathionine | 851.59 | 577.53 |
| MRET_0124 | membrane-associated progesterone receptor component | 1072.3 | 1128.67 |
| MRET_0125 | DUF1992 domain protein | 96.86 | 368.42 |
| MRET_0126 | SMR domain protein | 291.66 | 365.67 |
| MRET_0127 | ATP-dependent helicase IRC3 | 172.54 | 149.91 |
| MRET_0128 | MEMO1 family protein | 273.51 | 128.45 |
| MRET_0129 | uncharacterized protein | 105.51 | 100.82 |
| MRET_0130 | uncharacterized protein | 101.88 | 97.34 |
| MRET_0131 | FAS-associated factor 2 | 494.03 | 56.17 |
| MRET_0132 | cortical ER protein involved in ER-plasma membrane tethering | 34.04 | 21.15 |
| MRET_0133 | Na+/H+ antiporter | 80.67 | 13.62 |
| MRET_0134 | methyltransferase | 104.93 | 290.32 |
| MRET_0135 | alpha/beta-hydrolase | 42.24 | 45.9 |
| MRET_0136 | pseudouridylate synthase/pseudouridine kinase | 75.49 | 63.44 |
| MRET_0137 | sulfhydryl oxidase | 329.29 | 137.55 |
| MRET_0138 | uncharacterized protein | 1450.01 | 419.49 |
| MRET_0139 | uncharacterized protein | 71.52 | 145.21 |
| MRET_0140 | protein-L-isoaspartate(D-aspartate) O-methyltransferase | 190.61 | 117.54 |
| MRET_0141 | protein OS-9 | 519.38 | 253.6 |
| MRET_0142 | DUF1183 domain protein | 222.37 | 212.53 |
| MRET_0143 | peptide-methionine (R)-S-oxide reductase | 83.73 | 243.16 |
| MRET_0144 | uncharacterized protein | 776.79 | 2537.67 |
| MRET_0145 | adenylosuccinate lyase | 89.98 | 109.12 |

| | | | |
|---|---|---|---|
| MRET_0146 | DNA cross-link repair 1A protein | 114.3 | 42.18 |
| MRET_0147 | cyclin-C | 36.27 | 57.28 |
| MRET_0148 | uncharacterized protein | 70.01 | 62.48 |
| MRET_0149 | uncharacterized protein | 131.96 | 78.86 |
| MRET_0150 | uncharacterized protein | 23.36 | 18.89 |
| MRET_0151 | 2,5-diamino-6-(ribosylamino)-4(3H)-pyrimidinone 5'-phosphate reductase | 57.92 | 38.22 |
| MRET_0152 | transcription factor C subunit 7 | 119.64 | 116.16 |
| MRET_0153 | HAP4 transcription factor | 140.65 | 136.3 |
| MRET_0154 | nicotinamide N-methyltransferase | 71.69 | 121.26 |
| MRET_0155 | poly(A) RNA-binding protein | 181.94 | 147.34 |
| MRET_0156 | small subunit ribosomal protein S5 | 164.67 | 219.46 |
| MRET_0157 | exocyst complex component 4 | 44.15 | 45.67 |
| MRET_0158 | protein N-lysine methyltransferase METTL21D | 55.66 | 43.2 |
| MRET_0159 | protein MPE1 | 114.16 | 114.51 |
| MRET_0160 | vacuolar protein sorting-associated protein VTA1 | 42.13 | 63.85 |
| MRET_0161 | succinyl-CoA synthetase alpha subunit | 643.8 | 775.7 |
| MRET_0162 | deoxyhypusine monooxygenase | 236.33 | 195.07 |
| MRET_0163 | mRNA turnover protein 4 | 55.44 | 80.84 |
| MRET_0164 | mitochondrial import inner membrane translocase subunit TIM21 | 87.43 | 134.29 |
| MRET_0165 | uncharacterized protein | 3896.55 | 2379.36 |
| MRET_0166 | ubiquitin carboxyl-terminal hydrolase 12/46 | 21.1 | 39.57 |
| MRET_0167 | meiosis induction protein kinase IME2/SME1 | 10.15 | 40.64 |
| MRET_0168 | centractin | 54.32 | 119.42 |
| MRET_0169 | alpha-1,3-glucosyltransferase | 25.04 | 43.09 |
| MRET_0170 | translocation protein SEC62 | 48.22 | 102.08 |
| MRET_0171 | condensin complex subunit 1 | 31.69 | 46.45 |
| MRET_0172 | cwfJ domain protein | 188.93 | 82.23 |
| MRET_0173 | vacuolar ATPase assembly integral membrane protein VMA21 | 131.9 | 246.11 |
| MRET_0174 | N-alpha-acetyltransferase 30 | 40.03 | 65.02 |
| MRET_0175 | alpha/beta-hydrolase | 390.61 | 285.57 |
| MRET_0176 | alpha/beta-hydrolase | 46.19 | 95.91 |
| MRET_0177 | alpha-1,2-glucosyltransferase | 57.46 | 119.64 |
| MRET_0178 | tyrosine-protein kinase srms | 67.18 | 94.95 |
| MRET_0179 | uncharacterized protein | 11.1 | 28.1 |
| MRET_0180 | mannose-6-phosphate isomerase | 29.93 | 67.69 |
| MRET_0181 | 5'-3' exoribonuclease 2 | 39.35 | 80.67 |
| MRET_0182 | PITH domain protein | 246.56 | 247.53 |

| | | | |
|---|---|---|---|
| MRET_0183 | NADH dehydrogenase (ubiquinone) 1 beta subcomplex subunit 7 | 104.17 | 240.07 |
| MRET_0184 | small subunit ribosomal protein YMR-31 | 100.74 | 191.54 |
| MRET_0185 | syntaxin 16 | 53.74 | 47.9 |
| MRET_0186 | protein NAR1 | 246.34 | 133.39 |
| MRET_0187 | U3 small nucleolar ribonucleoprotein protein IMP3 | 24.2 | 32.8 |
| MRET_0188 | intermediate cleaving peptidase 55 | 22.01 | 61.97 |
| MRET_0189 | kinesin family member 20 | 12.19 | 39.33 |
| MRET_0190 | RNA recognition motif domain protein | 57.84 | 71.23 |
| MRET_0191 | pentatricopeptide repeat protein | 20.94 | 33.39 |
| MRET_0192 | alpha 1,2-mannosyltransferase | 86.32 | 144.83 |
| MRET_0193 | transcription initiation factor TFIID component TAF4 family | 34.59 | 68.37 |
| MRET_0194 | inosine-5'-monophosphate dehydrogenase | 58.74 | 221.7 |
| MRET_0195 | RuvB-like protein 1 (pontin 52) | 80.91 | 120.08 |
| MRET_0196 | uroporphyrinogen decarboxylase | 215.18 | 271.28 |
| MRET_0197 | protein of unknown function (DUF2456) | 53.96 | 81.29 |
| MRET_0198 | uncharacterized protein | 30.36 | 96.54 |
| MRET_0199 | DNA-directed RNA polymerase II subunit RPB3 | 39.62 | 129.21 |
| MRET_0200 | alpha 1,2-mannosyltransferase | 16.41 | 59.86 |
| MRET_0201 | alpha 1,2-mannosyltransferase | 69.54 | 110.08 |
| MRET_0202 | component of the NuA4 histone acetyltransferase complex | 42.51 | 57.5 |
| MRET_0203 | nuclear cap-binding protein subunit 1 | 54.22 | 52.83 |
| MRET_0204 | transcription elongation factor SPT4 | 43.77 | 103.99 |
| MRET_0205 | type II pantothenate kinase | 24.1 | 79.8 |
| MRET_0206 | translocation protein SEC63 | 123.3 | 160.65 |
| MRET_0207 | tRNA (guanine-N(7)-)-methyltransferase subunit TRM82 | 328.55 | 232.84 |
| MRET_0208 | elongation factor 3 | 33.43 | 53.32 |
| MRET_0209 | peroxin-3 | 41.88 | 39.71 |
| MRET_0210 | acyl CoA binding protein | 72.49 | 104.85 |
| MRET_0211 | 5-formyltetrahydrofolate cyclo-ligase | 40.57 | 60.47 |
| MRET_0212 | dolichyl-phosphate-mannose-protein mannosyltransferase | 96.83 | 94.64 |
| MRET_0213 | solute carrier family 25 (mitochondrial folate transporter), member 32 | 150.39 | 94.78 |
| MRET_0214 | polynucleotide 5'-hydroxyl-kinase GRC3/NOL9 | 30.1 | 30.64 |
| MRET_0215 | transcription factor | 72.58 | 68.4 |
| MRET_0216 | rRNA small subunit pseudouridine methyltransferase Nep1 | 201.79 | 221.08 |
| MRET_0217 | cell division cycle 14 | 10.13 | 39.83 |
| MRET_0218 | uncharacterized protein | 168 | 129.74 |
| MRET_0219 | protein CMC4 | 345.19 | 279.84 |

| | | | |
|---|---|---|---|
| MRET_0220 | folylpolyglutamate synthase | 23.8 | 18.1 |
| MRET_0221 | telomere length regulation protein | 19.54 | 26.74 |
| MRET_0222 | transcription factor | 77.04 | 85.69 |
| MRET_0223 | variant SH3 domain protein | 159.24 | 158.61 |
| MRET_0224 | cytochrome c oxidase subunit 15 | 156.99 | 241.54 |
| MRET_0225 | vacuolar transporter chaperone | 92.76 | 143.1 |
| MRET_0226 | Lhp1-RNA binding protein | 31.23 | 84.36 |
| MRET_0227 | ATP-dependent RNA helicase DDX31/DBP7 | 10.77 | 20.27 |
| MRET_0228 | protein SDA1 | 28.44 | 55.48 |
| MRET_0229 | predicted membrane protein required for lumenal ER protein retention | 8.76 | 14.63 |
| MRET_0230 | large subunit ribosomal protein L8e | 221.35 | 1382.14 |
| MRET_0231 | A1 cistron-splicing factor AAR2 | 97.63 | 125.91 |
| MRET_0232 | translation initiation factor IF-3 | 86.98 | 81.22 |
| MRET_0233 | golgi-specific brefeldin A-resistance guanine nucleotide exchange factor 1 | 105.75 | 51.91 |
| MRET_0234 | Ras-related protein Rab-1A | 584.62 | 536.99 |
| MRET_0235 | solute carrier family 25 (peroxisomal adenine nucleotide transporter), member 17 | 90.87 | 77.21 |
| MRET_0236 | transcriptional repressor NF-X1 | 67.42 | 54.82 |
| MRET_0237 | uncharacterized protein | 66.67 | 49.21 |
| MRET_0238 | DUF250 domain membrane protein | 183.36 | 178.46 |
| MRET_0239 | DNA damage-responsive transcriptional repressor | 78.51 | 55.21 |
| MRET_0240 | uncharacterized protein | 23.98 | 38.5 |
| MRET_0241 | seryl-tRNA synthetase | 89.64 | 50.06 |
| MRET_0242 | dCMP deaminase | 20.16 | 21.7 |
| MRET_0243 | solute carrier family 31 (copper transporter), member 1 | 20.73 | 33.22 |
| MRET_0244 | UDP-glucose--hexose-1-phosphate uridylyltransferase | 187.72 | 81.44 |
| MRET_0245 | uncharacterized protein | 122.07 | 92.11 |
| MRET_0246 | NET1-associated nuclear protein 1 (U3 small nucleolar RNA-associated protein 17) | 112.8 | 53.13 |
| MRET_0247 | RNA polymerase II subunit A C-terminal domain phosphatase | 50.91 | 78.75 |
| MRET_0248 | anaphase-promoting complex subunit 10 | 39.21 | 127.39 |
| MRET_0249 | DNA replication and checkpoint protein | 30.32 | 72.27 |
| MRET_0250 | DNA repair protein RAD5 | 356.76 | 380.79 |
| MRET_0251 | alpha-1,6-mannosyltransferase | 134 | 162.98 |
| MRET_0252 | very-long-chain (3R)-3-hydroxyacyl-CoA dehydratase | 391.4 | 220.86 |
| MRET_0253 | alkyl hydroperoxide reductase Thiol specific antioxidant Mal allergen | 269.29 | 385.09 |
| MRET_0254 | antiviral helicase SLH1 | 41.8 | 37.06 |
| MRET_0255 | glutaminyl-peptide cyclotransferase | 163.92 | 106.47 |
| MRET_0256 | nitrogen permease regulator 2-like protein | 79.58 | 61.25 |

| | | | |
|---|---|---|---|
| MRET_0257 | mitochondrial inner membrane carnitine transporter | 33.8 | 71.24 |
| MRET_0258 | CDP-diacylglycerol--inositol 3-phosphatidyltransferase | 76.51 | 206.65 |
| MRET_0259 | origin recognition complex subunit 3 | 31.76 | 35.02 |
| MRET_0260 | ATP-dependent RNA helicase MSS116, mitochondrial | 97.03 | 77.54 |
| MRET_0261 | uncharacterized protein | 4520.94 | 4076.95 |
| MRET_0262 | phosphodiesterase | 117.19 | 255.28 |
| MRET_0263 | nicotinamide mononucleotide adenylyltransferase | 210.09 | 428.5 |
| MRET_0264 | homeobox transcription factor | 24.41 | 47.31 |
| MRET_0265 | ATP-dependent RNA helicase DDX24/MAK5 | 16.76 | 38.38 |
| MRET_0266 | U3 small nucleolar RNA-associated protein 4 | 14.97 | 31.39 |
| MRET_0267 | glutamate 5-kinase | 44.79 | 82.16 |
| MRET_0268 | small subunit ribosomal protein S6 | 72.56 | 137.48 |
| MRET_0269 | replication factor C subunit 2/4 | 77.41 | 146.39 |
| MRET_0270 | chromatin assembly factor 1 subunit A | 110.17 | 52.18 |
| MRET_0271 | kinetochore protein Spc24, fungi type | 29.12 | 67.66 |
| MRET_0272 | C-8 sterol isomerase | 41.53 | 86.73 |
| MRET_0273 | MFS family protein | 98.83 | 270.16 |
| MRET_0274 | E3 ubiquitin-protein ligase RAD18 | 80.3 | 100.32 |
| MRET_0275 | zinc finger protein, C2H2 type | 27.65 | 51.16 |
| MRET_0276 | RNA recognition motif domain protein | 12.84 | 29.39 |
| MRET_0277 | protein JSN1 | 259.3 | 141.63 |
| MRET_0278 | uncharacterized protein | 670.87 | 595.85 |
| MRET_0279 | M-phase inducer tyrosine phosphatase | 14.98 | 31.38 |
| MRET_0280 | DNA polymerase phi | 12.68 | 41.36 |
| MRET_0281 | mitochondrial import inner membrane translocase subunit TIM8 | 82.22 | 211.29 |
| MRET_0282 | casein kinase I | 88.5 | 177.3 |
| MRET_0283 | monooxygenase | 66.32 | 78.8 |
| MRET_0284 | mitogen-activated protein kinase organizer 1 | 59.22 | 44.07 |
| MRET_0285 | metallo-beta-lactamase superfamily protein | 79.04 | 77.83 |
| MRET_0286 | Ran GTPase-activating protein 1 | 66.71 | 112.12 |
| MRET_0287 | myotubularin-related protein 6/7/8 | 38.44 | 101.93 |
| MRET_0288 | elongator complex protein 3 | 35.92 | 50.5 |
| MRET_0289 | fatty acid synthetase | 120.84 | 110.5 |
| MRET_0290 | glycolipid transfer protein HET-C2 | 277.4 | 1430.08 |
| MRET_0291 | fructose-bisphosphate aldolase, class II | 1343.96 | 1207.62 |
| MRET_0292 | Ca2+ transporting ATPase, sarcoplasmic/endoplasmic reticulum | 74.34 | 72 |
| MRET_0293 | uncharacterized protein | 116.92 | 115.85 |

| | | | |
|---|---|---|---|
| MRET_0294 | uncharacterized protein | 10.81 | 25.5 |
| MRET_0295 | cytochrome c peroxidase | 1230.63 | 504.72 |
| MRET_0296 | cytoplasmic inorganic pyrophosphatase (PPase) | 198.29 | 246.72 |
| MRET_0297 | pyruvate dehydrogenase E1 component beta subunit | 407.56 | 407.37 |
| MRET_0298 | casein kinase II subunit beta | 114.27 | 193.5 |
| MRET_0299 | translocation protein SEC66 | 45.28 | 81.51 |
| MRET_0300 | serine/threonine-protein phosphatase 2A regulatory subunit B' | 116.37 | 246.34 |
| MRET_0301 | prefoldin subunit 1 | 33.49 | 101.53 |
| MRET_0302 | cleavage stimulation factor subunit 2 | 41.93 | 75.91 |
| MRET_0303 | dTMP kinase | 29.02 | 36.86 |
| MRET_0304 | U1 snRNP splicing complex subunit Luc7 | 92.55 | 203.12 |
| MRET_0305 | uncharacterized protein | 52.55 | 103.41 |
| MRET_0306 | inositol polyphosphate phosphatase | 763.58 | 344.21 |
| MRET_0307 | protein N-lysine methyltransferase METTL21A | 50.91 | 29.02 |
| MRET_0308 | cysteinyl-tRNA synthetase | 122.98 | 77.74 |
| MRET_0309 | vacuolar protein sorting-associated protein 33 | 39.1 | 54.74 |
| MRET_0310 | component of cytosolic iron-sulfur protein assembly | 347.5 | 278.59 |
| MRET_0311 | transcriptional enhancer factor | 37.13 | 19.04 |
| MRET_0312 | ribonucleases P/MRP protein subunit RPP40 | 101.99 | 59.34 |
| MRET_0313 | sphingolipid long chain base-responsive protein | 53.76 | 49.2 |
| MRET_0314 | precorrin-2 dehydrogenase/sirohydrochlorin ferrochelatase | 229.06 | 165.49 |
| MRET_0315 | telomere length regulation protein | 75.93 | 38.23 |
| MRET_0316 | transcriptional regulator CBF1 | 186.87 | 567.35 |
| MRET_0317 | translation initiation factor 5A | 256.57 | 641.85 |
| MRET_0318 | chitin synthase | 23.48 | 25.12 |
| MRET_0319 | mitogen-activated protein kinase 1/3 | 423.26 | 164.91 |
| MRET_0320 | chitin synthase | 280.44 | 126.31 |
| MRET_0321 | zinc finger protein | 220.47 | 184.15 |
| MRET_0322 | serine/threonine-protein kinase receptor-associated protein | 89.59 | 163.93 |
| MRET_0323 | cell cycle checkpoint protein | 88.1 | 154.38 |
| MRET_0324 | SNARE domain protein | 199.5 | 118.97 |
| MRET_0325 | DNA repair protein RAD51 | 1418.28 | 1095.21 |
| MRET_0326 | amino acid transporter | 76.44 | 91.07 |
| MRET_0327 | ATP adenylyltransferase | 133.89 | 190.93 |
| MRET_0328 | ribonuclease P/MRP protein subunit RPP1 | 8.86 | 35.26 |
| MRET_0329 | uncharacterized protein | 241.89 | 196.38 |
| MRET_0330 | DASH complex subunit SPC19 | 39.33 | 182.11 |

| | | | |
|---|---|---|---|
| MRET_0331 | acylpyruvate hydrolase | 119 | 176.49 |
| MRET_0332 | mitochondrial pyruvate carrier 2 | 190.53 | 431.34 |
| MRET_0333 | pre-mRNA polyadenylation factor fip-1 | 37.7 | 78.42 |
| MRET_0334 | protein ATG11 | 32.36 | 60.84 |
| MRET_0335 | large subunit ribosomal protein L35 | 28.14 | 66.11 |
| MRET_0336 | mitochondrial transcription factor MTF1 | 43.26 | 84.07 |
| MRET_0337 | DnaJ homolog subfamily C member 17 | 24.57 | 66.28 |
| MRET_0338 | mitochondrial import inner membrane translocase subunit TIM23 | 32.15 | 62.89 |
| MRET_0339 | ATP-dependent RNA helicase DDX6/DHH1 | 48.42 | 65.45 |
| MRET_0340 | small plasma membrane protein | 61.7 | 152.46 |
| MRET_0341 | histone-binding protein RBBP4 | 66.2 | 124.24 |
| MRET_0342 | TELO2-interacting protein 1 | 54.47 | 51.45 |
| MRET_0343 | DNA repair protein RAD16 | 72.73 | 128.24 |
| MRET_0344 | uncharacterized protein | 83.83 | 71.08 |
| MRET_0345 | T-complex protein 1 subunit gamma | 80.8 | 242.52 |
| MRET_0346 | H+-transporting ATPase | 472.6 | 393.57 |
| MRET_0347 | mitochondrial import inner membrane translocase subunit TIM22 | 29.09 | 75.73 |
| MRET_0348 | 26S proteasome regulatory subunit T2 | 75.21 | 180.28 |
| MRET_0349 | histone demethylase JARID1 | 142.58 | 96.44 |
| MRET_0350 | uncharacterized protein | 118.33 | 101.14 |
| MRET_0351 | replication factor A2 | 1094.71 | 958.53 |
| MRET_0352 | V-type H+-transporting ATPase subunit e | 309.01 | 535.95 |
| MRET_0353 | dihydrofolate synthase | 23.78 | 72.12 |
| MRET_0354 | MYND domain protein (SamB) | 650.62 | 491.53 |
| MRET_0355 | mediator of RNA polymerase II transcription subunit 22 | 7.95 | 38.4 |
| MRET_0356 | uncharacterized protein | 350.64 | 521.96 |
| MRET_0357 | thymidylate synthase | 168.14 | 230.85 |
| MRET_0358 | protein transport protein SEC24 | 56.75 | 77.92 |
| MRET_0359 | solute carrier family 25 (mitochondrial phosphate transporter), member 23/24/25/41 | 25.41 | 95.68 |
| MRET_0360 | glyoxal/methylglyoxal oxidase | 186.62 | 142.09 |
| MRET_0361 | glycine-rich RNA binding protein | 1055.43 | 1349.56 |
| MRET_0362 | guanine nucleotide-binding protein subunit beta-2-like 1 protein | 189.61 | 653.62 |
| MRET_0363 | large subunit ribosomal protein L12e | 297.51 | 997.05 |
| MRET_0364 | uncharacterized protein | 19.46 | 37.85 |
| MRET_0365 | uncharacterized protein | 62.02 | 81.58 |
| MRET_0366 | uncharacterized protein | 80.3 | 46.84 |
| MRET_0367 | UDP-glucose 4-epimerase | 86.66 | 89.38 |

| | | | |
|---|---|---|---|
| MRET_0368 | DUF453 domain protein | 598.36 | 211.38 |
| MRET_0369 | uncharacterized protein | 10.84 | 25.97 |
| MRET_0370 | centromeric protein E | 51.5 | 46.5 |
| MRET_0371 | GTPase binding protein Rid1 | 7.03 | 16.11 |
| MRET_0372 | nitrogen permease regulator 3-like protein | 32.5 | 43.43 |
| MRET_0373 | ATP-dependent DNA helicase MPH1 | 149.25 | 167.38 |
| MRET_0374 | amidase | 50.8 | 76.68 |
| MRET_0375 | protein involved in negative regulation of iron regulon transcription | 522.5 | 506.39 |
| MRET_0376 | F-box protein, helicase, 18 | 131.29 | 87.89 |
| MRET_0377 | COP9 signalosome complex subunit 3 | 33 | 62.74 |
| MRET_0378 | sister chromatid cohesion protein PDS5 | 33.52 | 41.83 |
| MRET_0379 | PHD finger and SET domain protein | 90.62 | 104.04 |
| MRET_0380 | pre-mRNA-splicing helicase BRR2 | 37.91 | 92.79 |
| MRET_0381 | protein DJ-1 | 340.99 | 459.14 |
| MRET_0382 | long-chain acyl-CoA synthetase | 1352.44 | 2435.51 |
| MRET_0383 | serine arginine repetitive matrix 2 | 26.08 | 55.86 |
| MRET_0384 | hydroxymethylglutaryl-CoA reductase (NADPH) | 52.82 | 150.83 |
| MRET_0385 | protein TIF31 | 36.35 | 39.07 |
| MRET_0386 | ESCRT-II complex subunit VPS36 | 22.78 | 35.27 |
| MRET_0387 | 2-polyprenyl-6-hydroxyphenyl methylase/3-demethylubiquinone-9 3-methyltransferase | 28.12 | 32.85 |
| MRET_0388 | glycosylphosphatidylinositol transamidase | 416.25 | 523.79 |
| MRET_0389 | polyadenylation factor subunit 2 | 399.35 | 295.04 |
| MRET_0390 | phosphoinositide PI4,5P | 1426.66 | 874.42 |
| MRET_0391 | uncharacterized protein | 28.83 | 37.04 |
| MRET_0392 | ribonuclease Z | 416.41 | 213.87 |
| MRET_0393 | 4-hydroxysphinganine ceramide fatty acyl 2-hydroxylase | 839.31 | 794.93 |
| MRET_0394 | COX assembly mitochondrial protein 1 | 19.96 | 90.12 |
| MRET_0395 | ATP-dependent RNA helicase DDX51/DBP6 | 105.91 | 90.39 |
| MRET_0396 | DnaJ homolog subfamily C member 8 | 12.06 | 73.13 |
| MRET_0397 | acyl-CoA-dependent ceramide synthase | 33.12 | 83.48 |
| MRET_0398 | acetyltransferase (GNAT) family | 37.25 | 50.1 |
| MRET_0399 | peptidyl-prolyl isomerase | 3348.11 | 1779.06 |
| MRET_0400 | actin-related protein 6 | 36.73 | 30.11 |
| MRET_0401 | cytochrome c oxidase subunit 19 | 688.29 | 351.76 |
| MRET_0402 | SET and MYND domain protein | 90.87 | 82.65 |
| MRET_0403 | dolichyl-phosphate mannosyltransferase polypeptide 3 | 191.28 | 256.89 |
| MRET_0404 | uncharacterized protein | 54.44 | 121.43 |

| | | | |
|---|---|---|---|
| MRET_0405 | myosin V | 34.7 | 43.59 |
| MRET_0406 | solute carrier family 12 (potassium/chloride transporters), member 9 | 29.03 | 43.98 |
| MRET_0407 | pre-mRNA-splicing factor 38A | 23.81 | 39.64 |
| MRET_0408 | centromeric protein E | 16.84 | 39.67 |
| MRET_0409 | 26S proteasome regulatory subunit T5 | 129.53 | 150.17 |
| MRET_0410 | uncharacterized protein | 138.27 | 125.33 |
| MRET_0411 | G-patch domain protein | 119.42 | 87.5 |
| MRET_0412 | mediator of RNA polymerase II transcription subunit 4 | 30.87 | 54.25 |
| MRET_0413 | dual specificity tyrosine-phosphorylation-regulated kinase 2/3/4 | 36.35 | 52.21 |
| MRET_0414 | importin-4 | 22.78 | 60.68 |
| MRET_0415 | DnaJ homolog subfamily B member 6 | 364.1 | 423.41 |
| MRET_0416 | uncharacterized protein | 18 | 27.86 |
| MRET_0417 | TBC1 domain family member 5 | 42.71 | 47.88 |
| MRET_0418 | uncharacterized protein | 18.66 | 18.98 |
| MRET_0419 | amphiphysin | 668.6 | 492.97 |
| MRET_0420 | translation initiation factor eIF-2B subunit delta | 216.46 | 144.31 |
| MRET_0421 | dynein light chain LC8-type | 101.11 | 156.21 |
| MRET_0422 | conserved hypothetical protein | 112.45 | 208.46 |
| MRET_0423 | ATP-binding protein required for mismatch repair | 37.5 | 41.64 |
| MRET_0424 | dienelactone hydrolase | 148.28 | 130 |
| MRET_0425 | uncharacterized protein | 32.56 | 60.29 |
| MRET_0426 | V-type H+-transporting ATPase subunit G | 1174.92 | 829.94 |
| MRET_0427 | symplekin | 83.95 | 78.45 |
| MRET_0428 | tryptophan synthase | 95.23 | 93.03 |
| MRET_0429 | dienelactone hydrolase | 682.85 | 657.21 |
| MRET_0430 | mitochondrial protein import protein ZIM17 | 505.29 | 386.17 |
| MRET_0431 | solute carrier family 30 (zinc transporter), member 5/7 | 78.94 | 166.4 |
| MRET_0432 | uncharacterized protein | 29.05 | 30.95 |
| MRET_0433 | uncharacterized protein | 95.48 | 100.72 |
| MRET_0434 | WD repeat protein 68 | 156.58 | 169.75 |
| MRET_0435 | DNA replication licensing factor MCM5 | 36.16 | 44.88 |
| MRET_0436 | prohibitin 2 | 895.92 | 650.39 |
| MRET_0437 | uncharacterized protein | 108.93 | 78.71 |
| MRET_0438 | pyruvate carboxylase | 139.34 | 112.58 |
| MRET_0439 | small subunit ribosomal protein S18e | 413.52 | 1056.32 |
| MRET_0440 | solute carrier family 25 (mitochondrial iron transporter), member 28/37 | 212.18 | 170.24 |
| MRET_0441 | uncharacterized protein | 253.15 | 159.38 |

| | | | |
|---|---|---|---|
| MRET_0442 | uncharacterized protein | 56.12 | 125.67 |
| MRET_0443 | translation initiation factor 3 subunit G | 39.33 | 85.41 |
| MRET_0444 | V-type H+-transporting ATPase subunit E | 358.4 | 515.33 |
| MRET_0445 | large subunit ribosomal protein L54 | 29.03 | 77.56 |
| MRET_0446 | small G protein signaling modulator 3 | 9.42 | 14.66 |
| MRET_0447 | cleavage and polyadenylation specificity factor subunit 3 | 30.87 | 39.29 |
| MRET_0448 | DNA replication licensing factor MCM4 | 57.18 | 76.16 |
| MRET_0449 | mitotic spindle assembly checkpoint protein MAD2 | 21.35 | 66.08 |
| MRET_0450 | tyrosine-protein phosphatase SIW14 | 54.74 | 107.46 |
| MRET_0451 | F-type H+-transporting ATPase subunit g | 423.32 | 372.26 |
| MRET_0452 | protein of unknown function (DUF2781) | 24.69 | 35.41 |
| MRET_0453 | 4a-hydroxytetrahydrobiopterin dehydratase | 16.21 | 29.24 |
| MRET_0454 | U5 snRNP protein, DIM1 family | 16.7 | 43.24 |
| MRET_0455 | protein DML1 | 48.81 | 77.57 |
| MRET_0456 | vacuolar protein sorting-associated protein 16 | 48.78 | 38.38 |
| MRET_0457 | glutamate--cysteine ligase catalytic subunit | 70.65 | 84.72 |
| MRET_0458 | ubiquitin carboxyl-terminal hydrolase 1 | 22.71 | 33.42 |
| MRET_0459 | proteasome maturation protein | 134.2 | 106.22 |
| MRET_0460 | ATP-dependent RNA helicase DOB1 | 9.85 | 31.69 |
| MRET_0461 | nonsense-mediated mRNA decay protein 3 | 130.35 | 152.65 |
| MRET_0462 | zinc finger protein (RING finger) | 383.34 | 327.59 |
| MRET_0463 | transcription initiation factor TFIIE subunit alpha | 16.72 | 37.95 |
| MRET_0464 | MFS family protein | 71.69 | 164.09 |
| MRET_0465 | ubiquitin-like 1-activating enzyme E1 B | 219.81 | 448.25 |
| MRET_0466 | uncharacterized protein | 28.15 | 50.59 |
| MRET_0467 | PAPA-1-like conserved region containing protein | 29.23 | 87.65 |
| MRET_0468 | ATP-dependent RNA helicase DHX8/PRP22 | 31.65 | 54.69 |
| MRET_0469 | uncharacterized protein | 195.89 | 114.51 |
| MRET_0470 | chitin synthase activator | 461.92 | 360.5 |
| MRET_0471 | calmodulin | 682.64 | 902.4 |
| MRET_0472 | conserved hypothetical protein | 179.62 | 220.11 |
| MRET_0473 | uncharacterized protein | 53.75 | 96.22 |
| MRET_0474 | transcription regulator Maf1 | 204.71 | 82.28 |
| MRET_0475 | serine carboxypeptidase | 29.41 | 101.93 |
| MRET_0476 | phosphoribosyl-ATP pyrophosphohydrolase/phosphoribosyl-AMP cyclohydrolase/histidinol dehydrogenase | 231.24 | 179.61 |
| MRET_0477 | uncharacterized protein | 103.09 | 46.03 |
| MRET_0478 | carnitine O-acetyltransferase | 92.14 | 63.16 |

| | | | |
|---|---|---|---|
| MRET_0479 | SNF1-activating kinase 1 | 228.02 | 119.26 |
| MRET_0480 | XPG I-region protein | 100.44 | 102.04 |
| MRET_0481 | large subunit ribosomal protein L11e | 189.97 | 837.75 |
| MRET_0482 | uncharacterized protein | 203.98 | 152.9 |
| MRET_0483 | ATP-dependent RNA helicase DDX3X | 157.43 | 131.04 |
| MRET_0484 | deaminated glutathione amidase | 388.79 | 121.69 |
| MRET_0485 | nucleoside hydrolase | 62.05 | 82.18 |
| MRET_0486 | small nuclear ribonucleoprotein B and B' | 657.63 | 691.94 |
| MRET_0487 | tryptophan aminotransferase | 390.39 | 322.5 |
| MRET_0488 | DASH complex subunit Hsk3 like protein | 468.64 | 324.72 |
| MRET_0489 | syntaxin-binding protein 1 | 60.18 | 75.02 |
| MRET_0490 | DNA topoisomerase 2-associated protein PAT1 | 61.28 | 98.37 |
| MRET_0491 | BRCA1 C terminus (BRCT) domain protein | 64.19 | 73.3 |
| MRET_0492 | solute carrier family 26 (sodium-independent sulfate anion transporter), member 11 | 144.03 | 143.95 |
| MRET_0493 | nucleotide exchange factor | 22.5 | 45.51 |
| MRET_0494 | ethanolaminephosphotransferase | 29.86 | 56.29 |
| MRET_0495 | condensin complex subunit 2 | 17.52 | 50.2 |
| MRET_0496 | 2-acylglycerol O-acyltransferase 2 | 54.79 | 68.34 |
| MRET_0497 | histidine phosphatase | 19.85 | 33.96 |
| MRET_0498 | lysophospholipase | 1443.06 | 1052.97 |
| MRET_0499 | translation factor GUF1, mitochondrial | 34.84 | 48.08 |
| MRET_0500 | mRNA export factor | 269.6 | 371.53 |
| MRET_0501 | large subunit ribosomal protein L24e | 424.31 | 1072.06 |
| MRET_0502 | serine palmitoyltransferase | 105.96 | 138.47 |
| MRET_0503 | aldehyde dehydrogenase | 114.56 | 104.32 |
| MRET_0504 | COP9 signalosome complex subunit 5 | 24.65 | 37.22 |
| MRET_0505 | G2/mitotic-specific cyclin 1/2 | 96.43 | 116.47 |
| MRET_0506 | MIS18 kinetochore protein homolog A | 12.07 | 18.05 |
| MRET_0507 | mRNA-decapping enzyme 1B | 48.26 | 63.97 |
| MRET_0508 | replication fork protection complex subunit Tof1/Swi1 | 124.94 | 66.98 |
| MRET_0509 | DNA polymerase subunit cdc27 | 28.57 | 42.9 |
| MRET_0510 | ATP-dependent RNA helicase DDX19/DBP5 | 70.5 | 71.53 |
| MRET_0511 | uncharacterized protein | 36.31 | 40.47 |
| MRET_0512 | uncharacterized protein | 373.74 | 449.29 |
| MRET_0513 | CRAL/TRIO domain protein | 236.31 | 182.17 |
| MRET_0514 | F-type H+-transporting ATPase subunit k | 189.82 | 266.06 |
| MRET_0515 | uncharacterized protein | 106.78 | 108.1 |

| | | | |
|---|---|---|---|
| MRET_0516 | large subunit ribosomal protein L32e | 257.93 | 878.62 |
| MRET_0517 | large subunit ribosomal protein L13Ae | 107.09 | 438.12 |
| MRET_0518 | uncharacterized protein | 143.22 | 102.44 |
| MRET_0519 | DUF803 domain protein | 18.98 | 30.02 |
| MRET_0520 | phosphatidylinositol phospholipase C, delta | 192.26 | 118.9 |
| MRET_0521 | plasminogen activator inhibitor 1 RNA-binding protein | 104.36 | 237.61 |
| MRET_0522 | small subunit ribosomal protein S3e | 206.87 | 364.35 |
| MRET_0523 | dynamin-like GTPase MGM1, mitochondrial | 907.65 | 571.84 |
| MRET_0524 | 3-keto steroid reductase | 121.49 | 60.19 |
| MRET_0525 | RNA exonuclease 1 | 38.09 | 35.75 |
| MRET_0526 | uncharacterized protein | 136.22 | 112.67 |
| MRET_0527 | MAGE family protein | 24.79 | 55.29 |
| MRET_0528 | phospholipase carboxylesterase | 18.99 | 40.14 |
| MRET_0529 | HMG (high mobility group) box protein | 257.97 | 397.19 |
| MRET_0530 | 26S proteasome regulatory subunit N12 | 31.85 | 59.46 |
| MRET_0531 | uncharacterized protein | 11.79 | 32.11 |
| MRET_0532 | cytochrome b5 | 140.66 | 192.44 |
| MRET_0533 | pyruvate dehydrogenase phosphatase | 84.75 | 94.3 |
| MRET_0534 | vesicle-fusing ATPase | 239.62 | 130.4 |
| MRET_0535 | vacuolar import and degradation protein | 475.44 | 383.27 |
| MRET_0536 | translation initiation factor 3 subunit D | 52.09 | 100.22 |
| MRET_0537 | uncharacterized protein | 51.36 | 47.62 |
| MRET_0538 | CDP-diacylglycerol---glycerol-3-phosphate 3-phosphatidyltransferase | 105.16 | 61.77 |
| MRET_0539 | uncharacterized protein | 189.37 | 149.06 |
| MRET_0540 | uncharacterized protein | 173.53 | 166.04 |
| MRET_0541 | tRNA (adenine-N(1)-)-methyltransferase non-catalytic subunit | 143.44 | 249.37 |
| MRET_0542 | uncharacterized protein | 128.62 | 119.24 |
| MRET_0543 | U3 small nucleolar RNA-associated protein MPP10 | 1237.71 | 5947.52 |
| MRET_0544 | bifunctional dethiobiotin synthetase/adenosylmethionine---8-amino-7-oxononanoate aminotransferase | 114.62 | 77.84 |
| MRET_0545 | AHNAK nucleoprotein | 44.23 | 63.99 |
| MRET_0546 | anaphase-promoting complex subunit 8 | 9.39 | 28.53 |
| MRET_0547 | cytoplasm to vacuole targeting protein | 56.19 | 124.67 |
| MRET_0548 | large subunit ribosomal protein L38e | 71.91 | 384.42 |
| MRET_0549 | small subunit ribosomal protein S14e | 117.47 | 510.61 |
| MRET_0550 | nuclear movement protein NUDC | 50.44 | 155.75 |
| MRET_0551 | DNA repair and recombination protein RAD54B | 110.27 | 95.94 |
| MRET_0552 | outer membrane protein insertion porin family | 148 | 148.28 |

| | | | |
|---|---|---|---|
| MRET_0553 | DNA-directed RNA polymerase III subunit RPC8 | 19.42 | 53.65 |
| MRET_0554 | GTPase-activating protein SAC7 | 245.93 | 315.71 |
| MRET_0555 | vacuolar protein sorting-associated protein 8 | 103.39 | 94.52 |
| MRET_0556 | AFG3 family protein | 468.92 | 312.24 |
| MRET_0557 | SWI/SNF chromatin-remodeling complex subunit SWI1 | 48.25 | 53.73 |
| MRET_0558 | 26S proteasome regulatory subunit N1 | 67.13 | 113.97 |
| MRET_0559 | DNA-3-methyladenine glycosylase II | 87.45 | 113.67 |
| MRET_0560 | AHNAK nucleoprotein | 112.66 | 119.6 |
| MRET_0561 | solute carrier family 32 (vesicular inhibitory amino acid transporter) | 250.67 | 129.41 |
| MRET_0562 | nuclear pore complex protein Nup62 | 76.24 | 131.56 |
| MRET_0563 | 20S proteasome subunit beta 7 | 295.2 | 409.93 |
| MRET_0564 | dipeptidylpeptidase | 50.85 | 57.36 |
| MRET_0565 | replication factor C subunit 3/5 | 53.14 | 103.69 |
| MRET_0566 | alanine transaminase | 122.59 | 236.81 |
| MRET_0567 | uncharacterized protein | 32.77 | 39.12 |
| MRET_0568 | small subunit ribosomal protein S11 | 329.42 | 290.54 |
| MRET_0569 | small subunit ribosomal protein S9 | 121.03 | 161.49 |
| MRET_0570 | uncharacterized protein | 265.3 | 193.01 |
| MRET_0571 | phosphatidylinositol 4-kinase A | 82.34 | 47.52 |
| MRET_0572 | coiled-coil domain protein 12 | 64.3 | 65.07 |
| MRET_0573 | mitochondrial splicing suppressor protein 51 | 703.39 | 391.93 |
| MRET_0574 | urease accessory protein | 835.56 | 530.45 |
| MRET_0575 | endoplasmic reticulum-based factor for assembly of V-ATPase | 143.53 | 88.7 |
| MRET_0576 | peroxisome-assembly ATPase | 81.62 | 60.19 |
| MRET_0577 | checkpoint serine/threonine-protein kinase | 24.19 | 58.91 |
| MRET_0578 | ribosome biogenesis protein SSF1/2 | 260.3 | 358.3 |
| MRET_0579 | T-complex protein 1 subunit alpha | 52.65 | 95.91 |
| MRET_0580 | calcium/calmodulin-dependent protein kinase I | 1459.68 | 716.77 |
| MRET_0581 | protein of unknown function (DUF2423) | 38.94 | 38.25 |
| MRET_0582 | CDK inhibitor PHO81 | 201.8 | 172.51 |
| MRET_0583 | exportin-1 | 324.34 | 235.74 |
| MRET_0584 | golgi SNAP receptor complex member 1 | 187.46 | 105.34 |
| MRET_0585 | DNA binding regulatory protein AmdX | 83.79 | 99.15 |
| MRET_0586 | bZIP transcription factor | 126.32 | 34.01 |
| MRET_0587 | monothiol glutaredoxin | 975.59 | 906.18 |
| MRET_0588 | pre-rRNA-processing protein TSR4 | 50.22 | 89.77 |
| MRET_0589 | methylenetetrahydrofolate reductase (NADPH) | 58.84 | 70.99 |

| | | | |
|---|---|---|---|
| MRET_0590 | uncharacterized protein | 80.46 | 75.33 |
| MRET_0591 | origin recognition complex subunit 1 | 38.28 | 50.42 |
| MRET_0592 | alpha-1,2-mannosyltransferase | 54.87 | 61.07 |
| MRET_0593 | phospholipase C | 104.27 | 116.39 |
| MRET_0594 | uncharacterized protein | 248.98 | 184.11 |
| MRET_0595 | IPT/TIG domain protein | 365.73 | 220.47 |
| MRET_0596 | uncharacterized protein | 305.2 | 238.45 |
| MRET_0597 | protein SEY1 | 157.31 | 141.12 |
| MRET_0598 | NTF2 and RRM domain protein | 46.73 | 43.19 |
| MRET_0599 | dynamin 1-like protein | 680.26 | 419.21 |
| MRET_0600 | Cut9 interacting protein Scn1 | 83.95 | 121.02 |
| MRET_0601 | G2 mitotic-specific protein | 30.49 | 33.53 |
| MRET_0602 | zinc finger HIT domain protein 1 | 5.16 | 16.5 |
| MRET_0603 | uncharacterized protein | 24.49 | 30.2 |
| MRET_0604 | uncharacterized protein | 106.21 | 47.9 |
| MRET_0605 | protein farnesyltransferase subunit beta | 81.84 | 48.85 |
| MRET_0606 | protein disulfide-isomerase A6 | 33.92 | 47.91 |
| MRET_0607 | uncharacterized protein | 13.54 | 21.67 |
| MRET_0608 | protein of unknown function (DUF3405) | 480.71 | 227.25 |
| MRET_0609 | dolichyl-diphosphooligosaccharide---protein glycosyltransferase | 87.17 | 104.22 |
| MRET_0610 | peroxin-6 | 192.64 | 92.2 |
| MRET_0611 | malate dehydrogenase | 445.03 | 348.13 |
| MRET_0612 | conserved oligomeric golgi complex subunit 7 | 90.1 | 60.4 |
| MRET_0613 | 3-deoxy-7-phosphoheptulonate synthase | 179.23 | 210.36 |
| MRET_0614 | importin subunit beta-1 | 68.88 | 49.79 |
| MRET_0615 | SWI/SNF-related matrix-associated actin-dependent regulator of chromatin subfamily A member 5 | 147.63 | 101.86 |
| MRET_0616 | MICOS complex subunit MIC12 | 143.03 | 154.77 |
| MRET_0617 | DUF2407 ubiquitin-like domain protein | 77.38 | 74.4 |
| MRET_0618 | kexin | 94.64 | 83.72 |
| MRET_0619 | calnexin | 211.41 | 181.2 |
| MRET_0620 | argininosuccinate lyase | 290.57 | 139.49 |
| MRET_0621 | DnaJ homolog subfamily A member 2 | 310.55 | 262.69 |
| MRET_0622 | 25S rRNA (cytosine2278-C5)-methyltransferase | 11.59 | 12.31 |
| MRET_0623 | mannosyl-glycoprotein endo-beta-N-acetylglucosaminidase | 55.44 | 42.48 |
| MRET_0624 | SNARE domain protein | 109.66 | 94.65 |
| MRET_0625 | transcription initiation factor TFIID subunit 6 | 55.61 | 70.3 |
| MRET_0626 | methylenetetrahydrofolate dehydrogenase (NAD+) | 28.92 | 34.11 |

| | | | |
|---|---|---|---|
| MRET_0627 | translation initiation factor 4B | 212.5 | 103.52 |
| MRET_0628 | ubiquitin-protein ligase involved in ER-associated protein degradation | 142 | 108.46 |
| MRET_0629 | phospholipid-translocating ATPase | 80.73 | 45.76 |
| MRET_0630 | vacuolar protein sorting-associated protein 35 | 151.86 | 79.89 |
| MRET_0631 | large subunit ribosomal protein L18Ae | 712.09 | 1386.04 |
| MRET_0632 | RNA-binding protein 26 | 36.99 | 70.59 |
| MRET_0633 | large subunit ribosomal protein L40e | 1102.26 | 2167.89 |
| MRET_0634 | pre-60S factor REI1 | 241.64 | 292.62 |
| MRET_0635 | CUE domain protein | 51.17 | 27.35 |
| MRET_0636 | stress response protein NST1 | 119.53 | 112.16 |
| MRET_0637 | mitochondrial FAD-linked sulfhydryl oxidase | 419.58 | 168.86 |
| MRET_0638 | ubiquitin carboxyl-terminal hydrolase 8 | 16.29 | 23.66 |
| MRET_0639 | chromatin structure-remodeling complex protein RSC7 | 53.08 | 60.02 |
| MRET_0640 | uncharacterized protein | 181.97 | 108.04 |
| MRET_0641 | translation initiation factor 2 subunit 3 | 98.6 | 174.97 |
| MRET_0642 | golgi traffic protein SFT2 | 64.64 | 75.7 |
| MRET_0643 | F-type H+-transporting ATPase subunit h | 617.26 | 946.61 |
| MRET_0644 | ATPase inhibitor, mitochondrial | 2746.03 | 2255.54 |
| MRET_0645 | regulator of nonsense transcripts | 100.05 | 76.56 |
| MRET_0646 | OTU domain protein 3 | 14.22 | 27.89 |
| MRET_0647 | anaphase-promoting complex subunit 1 | 7.15 | 8.54 |
| MRET_0648 | 18S rRNA (adenine1779-N6/adenine1780-N6)-dimethyltransferase | 14.11 | 29.76 |
| MRET_0649 | AP complex subunit beta | 44.28 | 57.16 |
| MRET_0650 | protein of unknown function (DUF3074) | 234.99 | 217.19 |
| MRET_0651 | uncharacterized protein | 345.88 | 282.93 |
| MRET_0652 | aldose 1-epimerase | 20.82 | 23.58 |
| MRET_0653 | dolichyl-phosphate-mannose-protein mannosyltransferase | 32.62 | 47.23 |
| MRET_0654 | uncharacterized protein | 42.19 | 29.92 |
| MRET_0655 | long-chain-alcohol oxidase | 580.62 | 512.15 |
| MRET_0656 | 7SK snRNA methylphosphate capping enzyme | 510.19 | 457.97 |
| MRET_0657 | DNA-directed RNA polymerase, mitochondrial | 79.39 | 81.4 |
| MRET_0658 | uncharacterized protein | 25.76 | 20.58 |
| MRET_0659 | uncharacterized protein | 20.5 | 68.14 |
| MRET_0660 | superoxide dismutase, Fe-Mn family | 236.57 | 272.63 |
| MRET_0661 | exportin-4 | 120.6 | 70.65 |
| MRET_0662 | histone-lysine N-methyltransferase SETD2 | 49.58 | 60.09 |
| MRET_0663 | DNA-repair protein complementing XP-A cells | 14.76 | 27.88 |

| | | | |
|---|---|---|---|
| MRET_0664 | uncharacterized protein | 17.85 | 19.27 |
| MRET_0665 | prolactin regulatory element-binding protein | 56.17 | 74.6 |
| MRET_0666 | sphingolipid 8-(E)-desaturase | 285.95 | 117.27 |
| MRET_0667 | sphingomyelin phosphodiesterase | 115.85 | 80.71 |
| MRET_0668 | sphingomyelin phosphodiesterase | 26.72 | 59.33 |
| MRET_0669 | glutaredoxin-like protein | 3.6 | 7.03 |
| MRET_0670 | SUR7/PalI family protein | 3.89 | 31.44 |
| MRET_0671 | leucyl-tRNA synthetase | 48.46 | 56.72 |
| MRET_0672 | mediator of RNA polymerase II transcription subunit 12, fungi type | 49.29 | 32.17 |
| MRET_0673 | uncharacterized protein | 76.93 | 63.23 |
| MRET_0674 | actin related protein 2/3 complex, subunit 4 | 357.25 | 307.03 |
| MRET_0675 | chromodomain-helicase-DNA-binding protein 1 | 37.51 | 39.05 |
| MRET_0676 | nuclear GTP-binding protein | 40.74 | 51.84 |
| MRET_0677 | threonine dehydratase | 281.52 | 108.93 |
| MRET_0678 | ABC multidrug transporter | 46.31 | 68.12 |
| MRET_0679 | ferricrocin synthase | 31.1 | 157.42 |
| MRET_0680 | ATP-dependent RNA helicase DDX54/DBP10 | 69.89 | 61.48 |
| MRET_0681 | uncharacterized protein | 50.26 | 45.52 |
| MRET_0682 | serine/threonine-protein phosphatase PPG1 | 25.36 | 48.98 |
| MRET_0683 | pyridoxal phosphate phosphatase PHOSPHO2 | 43.52 | 71.07 |
| MRET_0684 | RhoGEF domain protein | 16.99 | 29.05 |
| MRET_0685 | peptidoglycan-binding domain protein | 38.53 | 43.49 |
| MRET_0686 | uncharacterized protein | 33.11 | 68.82 |
| MRET_0687 | protein of unknown function (DUF1168) | 36.16 | 82.79 |
| MRET_0688 | 2-methylcitrate dehydratase | 110.67 | 63.42 |
| MRET_0689 | cysteine synthase A | 69.62 | 122.13 |
| MRET_0690 | dual specificity protein kinase YAK1 | 192.8 | 115.06 |
| MRET_0691 | nuclear distribution protein nudE homolog 1 | 66.62 | 68.04 |
| MRET_0692 | ER to golgi transport-related protein | 48.9 | 75.75 |
| MRET_0693 | uncharacterized protein | 12.46 | 14.22 |
| MRET_0694 | nuclear GTP-binding protein | 162.86 | 899.77 |
| MRET_0695 | 20S proteasome subunit beta 1 | 74.73 | 134.27 |
| MRET_0696 | serine/threonine-protein phosphatase 2B regulatory subunit | 100.56 | 210.98 |
| MRET_0697 | small subunit ribosomal protein S13 | 35.66 | 107.59 |
| MRET_0698 | serine/threonine-protein kinase | 50.69 | 47.55 |
| MRET_0699 | acetoacetyl-CoA synthetase | 55.58 | 55.23 |
| MRET_0700 | H/ACA ribonucleoprotein complex subunit 1 | 180.77 | 260.92 |

| | | | |
|---|---|---|---|
| MRET_0701 | urea-proton symporter | 119.04 | 93.77 |
| MRET_0702 | vacuolar protein sorting-associated protein 26 | 40.44 | 54.07 |
| MRET_0703 | autophagy-related protein 27 | 128.23 | 102.41 |
| MRET_0704 | di- and tripeptidase | 18.35 | 30.87 |
| MRET_0705 | DNA-directed RNA polymerase III subunit RPC6 | 29.45 | 47.78 |
| MRET_0706 | DNA topoisomerase I | 29.52 | 72.45 |
| MRET_0707 | transcription activator of gluconeogenesis ERT1 | 4.86 | 23.57 |
| MRET_0708 | multifunctional methyltransferase subunit TRM112 | 30.52 | 106.73 |
| MRET_0709 | phosphoenolpyruvate carboxykinase (ATP) | 325.68 | 892.72 |
| MRET_0710 | mitochondrial pyruvate carrier 1 | 101.83 | 602.41 |
| MRET_0711 | DUF803 domain membrane protein | 1.67 | 26.25 |
| MRET_0712 | adenylate cyclase | 3.38 | 24.72 |
| MRET_0713 | solute carrier family 25 (mitochondrial phosphate transporter), member 3 | 187.77 | 189.07 |
| MRET_0714 | sulfate adenylyltransferase | 49.39 | 81.68 |
| MRET_0715 | histone-like transcription factor (CBF/NF-Y) | 238.81 | 258.91 |
| MRET_0716 | ATP-dependent DNA helicase 2 subunit 1 | 77.12 | 40.26 |
| MRET_0717 | peroxisomal 2,4-dienoyl-CoA reductase | 571.83 | 266.27 |
| MRET_0718 | large subunit ribosomal protein L39e | 188.88 | 667.18 |
| MRET_0719 | thiamin pyrophosphokinase-related protein | 312.93 | 284.42 |
| MRET_0720 | actin cortical patch component | 270.44 | 166.16 |
| MRET_0721 | peroxin-5 | 443.49 | 119.81 |
| MRET_0722 | HATPase_c domain protein | 17.49 | 19.34 |
| MRET_0723 | diphthamide biosynthesis protein 2 | 16.11 | 31.89 |
| MRET_0724 | MRC1-like domain protein | 23.76 | 38.12 |
| MRET_0725 | uncharacterized protein | 12.39 | 31.57 |
| MRET_0726 | phosphatidylinositol glycan, class P | 268.8 | 236.92 |
| MRET_0727 | cell division control protein | 38.3 | 73.45 |
| MRET_0728 | tyrosyl-tRNA synthetase | 171.68 | 166.98 |
| MRET_0729 | uncharacterized protein | 72.13 | 84.4 |
| MRET_0730 | mitochondrial import inner membrane translocase subunit TIM17 | 236.33 | 302.28 |
| MRET_0731 | beclin | 55.02 | 133.97 |
| MRET_0732 | F-type H+-transporting ATPase subunit delta | 402.03 | 524.07 |
| MRET_0733 | ATP-binding cassette, subfamily D (ALD), peroxisomal long-chain fatty acid import protein | 246.2 | 167.12 |
| MRET_0734 | ornithine decarboxylase antizyme | 342.66 | 365.34 |
| MRET_0735 | protein transport protein SEC61 subunit alpha | 152.7 | 321.71 |
| MRET_0736 | DNA-directed RNA polymerases I, II, and III subunit RPABC5 | 104.63 | 265.92 |
| MRET_0737 | uncharacterized protein | 90.26 | 122.19 |

| | | | |
|---|---|---|---|
| MRET_0738 | E3 ubiquitin-protein ligase BRE1 | 16.24 | 38.85 |
| MRET_0739 | sphinganine-1-phosphate aldolase | 408.63 | 288.13 |
| MRET_0740 | transcription initiation factor TFIIA large subunit | 793.21 | 714.76 |
| MRET_0741 | chromatin remodeling | 104.33 | 120.74 |
| MRET_0742 | Hamartin protein | 93.5 | 59.37 |
| MRET_0743 | glutaminyl-tRNA synthetase | 48.73 | 45.91 |
| MRET_0744 | transmembrane 9 superfamily member 2/4 | 93.13 | 109.7 |
| MRET_0745 | trafficking protein particle complex subunit 1 | 307.71 | 159.35 |
| MRET_0746 | diphosphomevalonate decarboxylase | 81.79 | 77.78 |
| MRET_0747 | large subunit ribosomal protein L27e | 119.36 | 492.46 |
| MRET_0748 | small subunit ribosomal protein S23e | 572.37 | 1437.31 |
| MRET_0749 | mitochondrial protein required for assembly of cytochrome bc1 complex | 384.21 | 361.49 |
| MRET_0750 | ribosome biogenesis protein BRX1 | 157.54 | 267.59 |
| MRET_0751 | uncharacterized protein | 6.3 | 7.34 |
| MRET_0752 | ergosterol biosynthesis protein | 639.75 | 442.54 |
| MRET_0753 | uncharacterized protein | 30.48 | 23.67 |
| MRET_0754 | uncharacterized protein | 9.86 | 13.09 |
| MRET_0755 | lysyl-tRNA synthetase, class II | 40.26 | 70.31 |
| MRET_0756 | endonuclease LCL3 | 188.23 | 152.84 |
| MRET_0757 | zinc finger protein (RING finger) | 289.98 | 194.29 |
| MRET_0758 | phosphoribosylformylglycinamidine synthase | 342.74 | 115.13 |
| MRET_0759 | transcriptional adapter 2-alpha | 29.55 | 65.7 |
| MRET_0760 | splicing factor U2AF 35 kDa subunit | 67.54 | 168.99 |
| MRET_0761 | Rab family GTPase | 127.71 | 204.55 |
| MRET_0762 | palmitoyltransferase | 44.98 | 71.89 |
| MRET_0763 | uncharacterized protein | 43.04 | 117 |
| MRET_0764 | uncharacterized protein | 28.79 | 43.95 |
| MRET_0765 | pentatricopeptide repeat protein | 7.88 | 19.89 |
| MRET_0766 | phosphatidylinositol 4-kinase type 2 | 37.23 | 36.99 |
| MRET_0767 | RNA polymerase I specific initiation factor | 290.37 | 113.57 |
| MRET_0768 | protein phosphatase PTC1 | 713.67 | 658.99 |
| MRET_0769 | required for meiotic nuclear division 5 homolog | 320.78 | 197.28 |
| MRET_0770 | ATP synthase mitochondrial F1 complex assembly factor 1 | 29.6 | 35.35 |
| MRET_0771 | phosphatidylinositol glycan, class W | 18.15 | 30.7 |
| MRET_0772 | F-type H+-transporting ATPase subunit O | 499.14 | 705.96 |
| MRET_0773 | translation initiation factor 3 subunit C | 73.05 | 101.7 |
| MRET_0774 | regulatory subunit for Cdc7p protein kinase | 285.99 | 127.18 |

| | | | |
|---|---|---|---|
| MRET_0775 | ribonuclease HI | 23.23 | 25.69 |
| MRET_0776 | NADPH-ferrihemoprotein reductase | 80.42 | 113.79 |
| MRET_0777 | DNA excision repair protein ERCC-1 | 19.91 | 44.86 |
| MRET_0778 | ribosome biogenesis protein NSA2 | 44.34 | 131.25 |
| MRET_0779 | protein transport protein YIP1 | 35.48 | 65.74 |
| MRET_0780 | DNA mismatch repair protein MLH1 | 56.05 | 82.56 |
| MRET_0781 | uncharacterized protein | 43.15 | 52.56 |
| MRET_0782 | cyclin H | 408.01 | 186.71 |
| MRET_0783 | exosome complex component RRP45 | 318.17 | 154.06 |
| MRET_0784 | uncharacterized protein | 2841.3 | 1910.96 |
| MRET_0785 | serine/threonine-protein phosphatase 2A activator | 164.47 | 92.05 |
| MRET_0786 | WD domain, G-beta repeat protein | 15.58 | 24.52 |
| MRET_0787 | TBC1 domain family member 10 | 69.1 | 120.36 |
| MRET_0788 | NIMA-related kinase 2 | 18.3 | 62.31 |
| MRET_0789 | protein FMP21 | 324.18 | 292.87 |
| MRET_0790 | SRP40, C-terminal domain protein | 38.17 | 89.2 |
| MRET_0791 | signal peptidase complex subunit 3 | 56.42 | 123.83 |
| MRET_0792 | uncharacterized protein | 237.98 | 223 |
| MRET_0793 | oxysterol-binding protein 1 | 125.27 | 122.07 |
| MRET_0794 | DASH complex subunit Duo1 | 87.44 | 170.31 |
| MRET_0795 | golgi pH regulator | 22.99 | 59.6 |
| MRET_0796 | protein EFR3 | 48.09 | 53.35 |
| MRET_0797 | serine hydrolase (FSH1) | 27.39 | 39.93 |
| MRET_0798 | UV excision repair protein RAD23 | 25.06 | 59.97 |
| MRET_0799 | homeobox domain protein | 29.9 | 65 |
| MRET_0800 | carbamoyl-phosphate synthase small subunit | 75.44 | 174.67 |
| MRET_0801 | uncharacterized protein | 15.05 | 39.72 |
| MRET_0802 | U4/U6.U5 tri-snRNP component SNU23 | 21.22 | 38.83 |
| MRET_0803 | isoleucyl-tRNA synthetase | 59.39 | 75.97 |
| MRET_0804 | F-type H+-transporting ATPase subunit gamma | 434.08 | 489.5 |
| MRET_0805 | translation initiation factor 2 subunit 2 | 248.14 | 434.74 |
| MRET_0806 | HIT finger domain protein | 23.13 | 40.99 |
| MRET_0807 | SAP domain ribonucleoprotein | 46.76 | 131.14 |
| MRET_0808 | uncharacterized protein | 62.82 | 146.81 |
| MRET_0809 | uncharacterized protein | 93.4 | 156.01 |
| MRET_0810 | ADP-ribosylation factor | 22.38 | 44.07 |
| MRET_0811 | SWI/SNF-related matrix-associated actin-dependent regulator of chromatin subfamily A | 19.55 | 36.22 |

| | | | |
|---|---|---|---|
| MRET_0812 | geranylgeranyl transferase type-2 subunit alpha | 30.13 | 98.47 |
| MRET_0813 | protein Yae1 | 32.11 | 144.53 |
| MRET_0814 | emopamil binding protein | 61.46 | 148.37 |
| MRET_0815 | uncharacterized protein | 451.68 | 689.26 |
| MRET_0816 | uncharacterized protein | 198.57 | 191.26 |
| MRET_0817 | histone acetyltransferase | 25.16 | 101.84 |
| MRET_0818 | phosphatidylinositol-binding clathrin assembly protein | 43.03 | 38.02 |
| MRET_0819 | uncharacterized protein | 11.03 | 46.35 |
| MRET_0820 | 25S rRNA (uracil2634-N3)-methyltransferase | 29.93 | 40.17 |
| MRET_0821 | DnaJ homolog subfamily C member 3 | 54.97 | 99.99 |
| MRET_0822 | uncharacterized protein | 453.73 | 409.24 |
| MRET_0823 | iron-sulfur cluster assembly protein ISA1 | 410.2 | 241.46 |
| MRET_0824 | GTPase | 34.58 | 33.85 |
| MRET_0825 | membrane protein TMS1 | 64.92 | 55.46 |
| MRET_0826 | folic acid synthesis protein | 37.51 | 58.75 |
| MRET_0827 | uncharacterized protein | 104.31 | 142.88 |
| MRET_0828 | DNA polymerase epsilon subunit 2 | 317 | 286.61 |
| MRET_0829 | phosphatidylinositol glycan, class A | 103.67 | 141.65 |
| MRET_0830 | alanyl-tRNA synthetase | 34.11 | 59.72 |
| MRET_0831 | zinc finger protein, C2H2 type | 517.82 | 206.12 |
| MRET_0832 | 3-dehydrosphinganine reductase | 40.92 | 20.39 |
| MRET_0833 | uncharacterized protein | 43.33 | 69.06 |
| MRET_0834 | uncharacterized protein | 685.46 | 543.67 |
| MRET_0835 | cathepsin D | 217.94 | 143.11 |
| MRET_0836 | mRNA guanylyltransferase | 23.88 | 62.44 |
| MRET_0837 | tRNA-splicing endonuclease subunit Sen2 | 29.66 | 42.87 |
| MRET_0838 | dentin sialophosphoprotein | 1044.49 | 747.63 |
| MRET_0839 | programmed cell death protein 5 | 27.38 | 153.21 |
| MRET_0840 | conserved hypothetical protein | 31.1 | 85.07 |
| MRET_0841 | RanBD | 134 | 125.37 |
| MRET_0842 | translation initiation factor eIF-2B subunit epsilon | 86.14 | 104.34 |
| MRET_0843 | phosphopantothenoylcysteine decarboxylase | 51.53 | 38.76 |
| MRET_0844 | L-galactose dehydrogenase | 193.03 | 116.14 |
| MRET_0845 | phosphoglycerate kinase | 1515.02 | 839.74 |
| MRET_0846 | AMP deaminase | 27.3 | 27.17 |
| MRET_0847 | merozoite surface protein msp-1 | 247.67 | 117.84 |
| MRET_0848 | zinc finger protein | 108.51 | 61.25 |

| | | | |
|---|---|---|---|
| MRET_0849 | pyridoxine kinase | 67.26 | 76.51 |
| MRET_0850 | uncharacterized protein | 7.5 | 32.14 |
| MRET_0851 | DNA polymerase alpha subunit A | 59.5 | 41.79 |
| MRET_0852 | centromere protein Scm3 | 57.96 | 67.6 |
| MRET_0853 | uncharacterized protein | 194.5 | 181.33 |
| MRET_0854 | large subunit ribosomal protein L35Ae | 359.24 | 891.7 |
| MRET_0855 | bud site selection protein 20 | 35.47 | 57.4 |
| MRET_0856 | FAD dependent oxidoreductase | 170.71 | 213.57 |
| MRET_0857 | DUF1014 domain protein | 1039.27 | 718.56 |
| MRET_0858 | uncharacterized protein | 118.08 | 80.42 |
| MRET_0859 | large subunit ribosomal protein L27 | 215.08 | 215.75 |
| MRET_0860 | mitochondrial import inner membrane translocase subunit TIM23 | 83.38 | 152.94 |
| MRET_0861 | uncharacterized protein | 630.76 | 388.9 |
| MRET_0862 | multi-transmembrane subunit of the DSC ubiquitin ligase complex | 24.5 | 56.8 |
| MRET_0863 | fungal domain of unknown function (DUF1712) | 61.25 | 53.11 |
| MRET_0864 | uncharacterized protein | 546.93 | 232.93 |
| MRET_0865 | prephenate dehydrogenase (NADP+) | 94.47 | 54.58 |
| MRET_0866 | PX domain protein | 77.71 | 49.69 |
| MRET_0867 | beta-1,4-N-acetylglucosaminyltransferase | 33.7 | 34.48 |
| MRET_0868 | uncharacterized protein | 4.04 | 31.78 |
| MRET_0869 | syntaxin 1B/2/3 | 207.16 | 217.75 |
| MRET_0870 | large subunit ribosomal protein L15 | 49.85 | 75.22 |
| MRET_0871 | ATP-dependent RNA helicase DDX23/PRP28 | 245.81 | 152.35 |
| MRET_0872 | acetyl-CoA carboxylase, biotin containing enzyme | 77.99 | 40.93 |
| MRET_0873 | ubiquinol-cytochrome c reductase cytochrome c1 subunit | 353.55 | 248.38 |
| MRET_0874 | 4-amino-4-deoxychorismate lyase | 65.14 | 67.9 |
| MRET_0875 | crossover junction endonuclease MUS81 | 91.13 | 94.29 |
| MRET_0876 | small subunit ribosomal protein S15e | 121.11 | 586.72 |
| MRET_0877 | large subunit ribosomal protein LP2 | 939.91 | 1753.94 |
| MRET_0878 | Rho GTPase-activating protein RGD1 | 114.56 | 120.05 |
| MRET_0879 | cryptococcal mannosyltransferase 1 | 95.06 | 91.97 |
| MRET_0880 | uncharacterized protein | 22.66 | 34.39 |
| MRET_0881 | DUF803 domain membrane protein | 146.06 | 71.29 |
| MRET_0882 | PPR repeat containing protein | 93.96 | 33.92 |
| MRET_0883 | uncharacterized protein | 498.6 | 810.14 |
| MRET_0884 | urea transporter | 414.23 | 895.59 |
| MRET_0885 | transcriptional adapter 3 | 25.4 | 50.83 |

| | | | |
|---|---|---|---|
| MRET_0886 | nucleotide exchange factor for Gsp1p | 20.28 | 23.85 |
| MRET_0887 | fatty acid elongase 3 | 158.67 | 64.57 |
| MRET_0888 | ubiquitin-conjugating enzyme E2 D | 3274.93 | 3163.48 |
| MRET_0889 | SANT domain protein | 93.1 | 66.14 |
| MRET_0890 | FMN binding oxidoreductase | 1138.25 | 663.24 |
| MRET_0891 | origin recognition complex subunit 5 | 53.84 | 52.76 |
| MRET_0892 | uncharacterized protein | 18.09 | 25.54 |
| MRET_0893 | member of the NineTeen Complex | 162.66 | 119.38 |
| MRET_0894 | glyoxylate/hydroxypyruvate reductase | 211.13 | 205.72 |
| MRET_0895 | ubiquinol-cytochrome c reductase core subunit 2 | 565.64 | 366.4 |
| MRET_0896 | DNA/RNA-binding protein KIN17 | 1810.19 | 723.3 |
| MRET_0897 | glycine hydroxymethyltransferase | 179.36 | 554.4 |
| MRET_0898 | mitochondrial exoribonuclease Cyt-4 | 97.47 | 103.54 |
| MRET_0899 | DNA damage-inducible protein 1 | 109.39 | 146.32 |
| MRET_0900 | pre-mRNA-splicing factor ATP-dependent RNA helicase DHX38/PRP16 | 138.3 | 92.07 |
| MRET_0901 | nuclear pore complex protein Nup37 | 32.62 | 33.26 |
| MRET_0902 | E3 ubiquitin-protein ligase TRIP12 | 370.09 | 188.73 |
| MRET_0903 | CLIP-associating protein 1/2 | 252.72 | 158.99 |
| MRET_0904 | replication factor C subunit 3/5 | 67.26 | 76.52 |
| MRET_0905 | uncharacterized protein | 54.47 | 50.96 |
| MRET_0906 | sorting nexin-41/42 | 163.65 | 131.33 |
| MRET_0907 | ubiquitin | 262.02 | 104.81 |
| MRET_0908 | aspartate aminotransferase, cytoplasmic | 1083.88 | 731.69 |
| MRET_0909 | non-structural maintenance of chromosomes element 1 | 53.49 | 38.63 |
| MRET_0910 | methylenetetrahydrofolate reductase (NADPH) | 32.66 | 28.28 |
| MRET_0911 | actin-related protein 8 | 91.55 | 70.04 |
| MRET_0912 | SAP domain protein | 367.15 | 236.02 |
| MRET_0913 | imidazoleglycerol-phosphate dehydratase | 181.93 | 129.35 |
| MRET_0914 | dynein intermediate chain, cytosolic | 137.21 | 121.52 |
| MRET_0915 | uncharacterized protein | 154.84 | 141.16 |
| MRET_0916 | SAC3 GANP domain protein | 109.14 | 87.27 |
| MRET_0917 | homoserine kinase | 116.65 | 138.87 |
| MRET_0918 | kinetochore protein Nuf2 | 48.47 | 116.19 |
| MRET_0919 | glutathione-dependent oxidoreductase | 619.81 | 433.09 |
| MRET_0920 | ubiquitin carboxyl-terminal hydrolase 4/11 | 66.17 | 56.32 |
| MRET_0921 | peptidase inhibitor activity protein | 618.34 | 4126.24 |
| MRET_0922 | glycosyl transferase family protein | 93.58 | 60.31 |

| | | | |
|---|---|---|---|
| MRET_0923 | acylglycerol lipase | 142.31 | 141.33 |
| MRET_0924 | histone H3 | 218.28 | 125.65 |
| MRET_0925 | histone H4 | 4208.07 | 2174.05 |
| MRET_0926 | acetyl-CoA acyltransferase 1 | 716.55 | 394.34 |
| MRET_0927 | cyclin-dependent protein kinase complex component | 12.19 | 34.03 |
| MRET_0928 | uncharacterized protein | 31.49 | 87.3 |
| MRET_0929 | HMG (high mobility group) box protein | 59.1 | 121.23 |
| MRET_0930 | secretory lipase | 499.03 | 824.46 |
| MRET_0931 | annexin A7 | 512.01 | 288.68 |
| MRET_0932 | uncharacterized protein | 186.01 | 96.26 |
| MRET_0933 | glutamate N-acetyltransferase/amino-acid N-acetyltransferase | 32.1 | 40.76 |
| MRET_0934 | solute carrier family 39 (zinc transporter), member 1/2/3 | 65.93 | 63.67 |
| MRET_0935 | solute carrier family 39 (zinc transporter), member 1/2/3 | 59.53 | 11.95 |
| MRET_0936 | cell cycle control protein | 36.4 | 67.76 |
| MRET_0937 | small subunit ribosomal protein S4e | 119.97 | 531.01 |
| MRET_0938 | DnaJ domain protein | 457.77 | 551.68 |
| MRET_0939 | nucleus protein | 96.19 | 69.57 |
| MRET_0940 | SAC3 GANP domain protein | 127.96 | 85.06 |
| MRET_0941 | solute carrier family 35, member E1 | 16.8 | 23.99 |
| MRET_0942 | uncharacterized protein | 203.49 | 114.98 |
| MRET_0943 | ATP-dependent RNA helicase DDX10/DBP4 | 158.18 | 125.83 |
| MRET_0944 | mitochondrial carrier protein | 87.5 | 138.8 |
| MRET_0945 | multidrug resistance protein, MATE family | 33.11 | 36.71 |
| MRET_0946 | CTD kinase subunit beta | 37.4 | 33 |
| MRET_0947 | DnaJ homolog subfamily C member 9 | 68 | 79.74 |
| MRET_0948 | uncharacterized protein | 1692.05 | 1124.25 |
| MRET_0949 | uncharacterized protein | 1679.59 | 2251.19 |
| MRET_0950 | AHA1 family protein | 346.95 | 448.37 |
| MRET_0951 | 26S proteasome regulatory subunit N6 | 76.36 | 142.91 |
| MRET_0952 | 26S proteasome regulatory subunit N7 | 287.48 | 215.4 |
| MRET_0953 | UBA TS-N domain protein | 144.31 | 85.95 |
| MRET_0954 | signal transducing adaptor molecule | 180.07 | 181.86 |
| MRET_0955 | queuosine salvage protein | 27.35 | 28.32 |
| MRET_0956 | DEAD/DEAH box helicase | 28.99 | 39.3 |
| MRET_0957 | trafficking protein particle complex subunit 11 | 20.06 | 25.04 |
| MRET_0958 | midasin | 173.91 | 137.42 |
| MRET_0959 | U3 small nucleolar RNA-associated protein 21 | 260.38 | 117.23 |

| | | | |
|---|---|---|---|
| MRET_0960 | ATP-dependent RNA helicase DDX18/HAS1 | 22.91 | 51.82 |
| MRET_0961 | periodic tryptophan protein 2 | 108.15 | 85.23 |
| MRET_0962 | Sec7 domain protein | 11 | 33.89 |
| MRET_0963 | cyclin (Pcl1) | 11.66 | 67.51 |
| MRET_0964 | VanZ domain protein | 126.98 | 291.39 |
| MRET_0965 | uncharacterized protein | 140.55 | 222.15 |
| MRET_0966 | fructose-1,6-bisphosphatase I | 473.71 | 336.68 |
| MRET_0967 | protoporphyrinogen/coproporphyrinogen III oxidase | 190.65 | 89.46 |
| MRET_0968 | ubiquinol-cytochrome c reductase subunit 7 | 325.26 | 378.43 |
| MRET_0969 | DUF300 domain protein | 31.6 | 53.06 |
| MRET_0970 | RAD50-interacting protein 1 | 54.35 | 68.73 |
| MRET_0971 | uncharacterized protein | 635.25 | 660.87 |
| MRET_0972 | translin family protein | 109.01 | 126.38 |
| MRET_0973 | uncharacterized protein | 11.79 | 14.32 |
| MRET_0974 | uncharacterized protein | 357.21 | 252.37 |
| MRET_0975 | COP9 signalosome complex subunit 6 | 110.75 | 83.55 |
| MRET_0976 | DNA replication licensing factor MCM3 | 156.15 | 150.39 |
| MRET_0977 | Ras-related protein Rab-11A | 179.74 | 286.91 |
| MRET_0978 | peroxin-4 | 64.98 | 63.34 |
| MRET_0979 | mitogen-activated protein kinase kinase kinase | 98.63 | 87.58 |
| MRET_0980 | transportin-3 | 55.32 | 83.52 |
| MRET_0981 | vacuolar protein-sorting-associated protein 4 | 257.07 | 325.94 |
| MRET_0982 | phospholipase D1/2 | 15.5 | 29.86 |
| MRET_0983 | citrate synthase | 168.01 | 175.27 |
| MRET_0984 | uroporphyrinogen-III synthase | 32.81 | 57.83 |
| MRET_0985 | uncharacterized protein | 43.29 | 52.97 |
| MRET_0986 | golgi vesicular membrane trafficking protein | 91.26 | 102.68 |
| MRET_0987 | rhomboid family membrane protein | 100.91 | 107.71 |
| MRET_0988 | uncharacterized protein | 36.34 | 51.95 |
| MRET_0989 | LIM domain protein | 11.75 | 24.38 |
| MRET_0990 | enhanced filamentous growth protein 1 | 98.17 | 103.36 |
| MRET_0991 | uncharacterized protein | 30.93 | 53.26 |
| MRET_0992 | uncharacterized protein | 53.76 | 78.42 |
| MRET_0993 | 1,3-beta-glucan synthase | 54.75 | 89.61 |
| MRET_0994 | sterol 22-desaturase | 55.13 | 92.22 |
| MRET_0995 | serine/threonine-protein kinase | 117.06 | 93.91 |
| MRET_0996 | transcription initiation factor TFIIH subunit 3 | 30.34 | 42.67 |

| | | | |
|---|---|---|---|
| MRET_0997 | cyclin | 19.84 | 26.39 |
| MRET_0998 | charged multivesicular body protein 3 | 15.43 | 20.32 |
| MRET_0999 | histone H2B | 279.61 | 315.19 |
| MRET_1000 | histone H2A | 529 | 370.9 |
| MRET_1001 | lysine-specific demethylase 8 | 27.24 | 24.82 |
| MRET_1002 | isocitrate dehydrogenase (NAD+) IDH1 | 267.38 | 345.42 |
| MRET_1003 | isocitrate dehydrogenase (NAD+) IDH2 | 373.87 | 455.25 |
| MRET_1004 | component of the cleavage and polyadenylation factor I | 442.45 | 472.92 |
| MRET_1005 | large subunit ribosomal protein L18e | 223.31 | 488.5 |
| MRET_1006 | LisH domain protein | 233.71 | 121.21 |
| MRET_1007 | AP-1 complex subunit mu | 69.24 | 112.6 |
| MRET_1008 | RhoGEF domain protein | 181.41 | 90.5 |
| MRET_1009 | BadF/BadG/BcrA/BcrD ATPase family | 17.26 | 13.19 |
| MRET_1010 | tRNA (cytosine34-C5)-methyltransferase | 54.7 | 31.26 |
| MRET_1011 | 2-dehydropantoate 2-reductase | 235.14 | 181.46 |
| MRET_1012 | uncharacterized protein | 756.82 | 579.39 |
| MRET_1013 | uncharacterized protein | 65.81 | 33.56 |
| MRET_1014 | conserved hypothetical protein | 23.36 | 36.2 |
| MRET_1015 | uncharacterized protein | 85.21 | 199.1 |
| MRET_1016 | peptide-methionine (S)-S-oxide reductase | 1472.67 | 1319.02 |
| MRET_1017 | uncharacterized protein | 24.37 | 54.15 |
| MRET_1018 | ubiquitin-conjugating enzyme E2 J1 | 8.74 | 25.92 |
| MRET_1019 | ER membrane protein | 643.66 | 481.75 |
| MRET_1020 | fatty acid hydroxylase | 89.28 | 151.26 |
| MRET_1021 | uncharacterized protein | 25.44 | 57.42 |
| MRET_1022 | adenylyltransferase and sulfurtransferase | 54.99 | 57.57 |
| MRET_1023 | Importin-11 | 54.54 | 69.59 |
| MRET_1024 | vacuolar carboxypeptidase | 1121.27 | 939.77 |
| MRET_1025 | SNF2 family helicase ATPase | 27.47 | 16.83 |
| MRET_1026 | alkylated DNA repair protein alkB homolog 6 | 34.09 | 17.84 |
| MRET_1027 | mitosis inhibitor protein kinase SWE1 | 53.3 | 86.51 |
| MRET_1028 | uncharacterized protein | 97.4 | 213.26 |
| MRET_1029 | uncharacterized protein | 300.03 | 424.42 |
| MRET_1030 | small subunit ribosomal protein S6e | 138.15 | 629.32 |
| MRET_1031 | small subunit ribosomal protein S13e | 192.33 | 981.01 |
| MRET_1032 | lipase | 31.13 | 29.77 |
| MRET_1033 | NADPH-dependent medium chain alcohol dehydrogenase | 126.28 | 140.29 |

| | | | |
|---|---|---|---|
| MRET_1034 | inositol-polyphosphate multikinase | 2.32 | 2.69 |
| MRET_1035 | vacuolar protein sorting-associated protein 52 | 48.28 | 47.67 |
| MRET_1036 | protein regulator of cytokinesis 1 | 62.67 | 100.67 |
| MRET_1037 | protein NUD1 | 20.88 | 35.72 |
| MRET_1038 | riboflavin synthase | 60.34 | 102 |
| MRET_1039 | bud site selection protein 31 | 36.5 | 50.81 |
| MRET_1040 | GTPase-activating protein SST2 | 72.5 | 79.24 |
| MRET_1041 | histone deacetylase 6 | 39.57 | 41.8 |
| MRET_1042 | uncharacterized protein | 49.04 | 35.89 |
| MRET_1043 | uncharacterized protein | 78.5 | 68.68 |
| MRET_1044 | uncharacterized protein | 25.27 | 32.64 |
| MRET_1045 | 26S proteasome regulatory subunit N11 | 154.1 | 153.18 |
| MRET_1046 | nuclear pore complex protein Nup160 | 34.23 | 25.22 |
| MRET_1047 | kinesin family member 18/19 | 17.3 | 32.34 |
| MRET_1048 | protein-S-isoprenylcysteine O-methyltransferase | 13.45 | 22.72 |
| MRET_1049 | dihydroorotase | 17.35 | 38.21 |
| MRET_1050 | zinc finger protein HUA1 | 130.25 | 100.26 |
| MRET_1051 | cactin protein | 220.91 | 140 |
| MRET_1052 | anticodon-binding domain protein | 55.44 | 122.62 |
| MRET_1053 | NADH dehydrogenase (ubiquinone) 1 alpha subcomplex subunit 2 | 107.23 | 116.44 |
| MRET_1054 | protein pelota | 55.93 | 59.77 |
| MRET_1055 | uncharacterized protein | 28.6 | 54.66 |
| MRET_1056 | uncharacterized protein | 11.39 | 26.47 |
| MRET_1057 | uncharacterized protein | 41.06 | 100.24 |
| MRET_1058 | Gly-Xaa carboxypeptidase | 62.55 | 39.41 |
| MRET_1059 | Anion exchange family protein | 36.95 | 68.88 |
| MRET_1060 | RNA polymerase Rpb4 | 22.58 | 54.21 |
| MRET_1061 | small subunit ribosomal protein S18 | 56.15 | 132.01 |
| MRET_1062 | thiamine-phosphate diphosphorylase/hydroxyethylthiazole kinase | 118.5 | 105.98 |
| MRET_1063 | GINS complex subunit 3 | 97.22 | 132.08 |
| MRET_1064 | exocyst complex component 5 | 99.37 | 102.48 |
| MRET_1065 | uncharacterized protein | 18.02 | 22.95 |
| MRET_1066 | peroxin-2 | 38.85 | 35.25 |
| MRET_1067 | magnesium-dependent phosphatase 1 | 513.22 | 146.69 |
| MRET_1068 | histone deacetylase 1/2 | 205.24 | 148.92 |
| MRET_1069 | general transcription factor 3C polypeptide 3 (transcription factor C subunit 4) | 186.81 | 81.97 |
| MRET_1070 | uncharacterized protein | 63.43 | 110.21 |

| | | | |
|---|---|---|---|
| MRET_1071 | 18S rRNA (guanine1575-N7)-methyltransferase | 29.55 | 56.02 |
| MRET_1072 | syntaxin 18 | 91.06 | 136.42 |
| MRET_1073 | pyruvate dehydrogenase kinase | 26.59 | 49.24 |
| MRET_1074 | uncharacterized protein | 51.58 | 67.22 |
| MRET_1075 | calcium calmodulin-dependent protein kinase | 54.99 | 109.46 |
| MRET_1076 | L-threonylcarbamoyladenylate synthase | 189.1 | 161.76 |
| MRET_1077 | cytochrome c oxidase assembly factor 6 | 29.52 | 105.01 |
| MRET_1078 | uncharacterized protein | 19.18 | 53.79 |
| MRET_1079 | L-2-aminoadipate reductase | 32.95 | 45.01 |
| MRET_1080 | cell morphogenesis protein (PAG1) | 33.76 | 29.85 |
| MRET_1081 | poly(A) RNA binding protein involved in nuclear mRNA export | 15.72 | 33.73 |
| MRET_1082 | phosphatidylinositol glycan, class F | 49.24 | 39.2 |
| MRET_1083 | AHNAK nucleoprotein | 20.23 | 27.94 |
| MRET_1084 | mitochondrial distribution and morphology protein 12 | 45.89 | 38.89 |
| MRET_1085 | uncharacterized protein | 62.98 | 123.64 |
| MRET_1086 | large subunit ribosomal protein L29e | 288.06 | 556.37 |
| MRET_1087 | FYVE zinc finger protein | 63.36 | 41.1 |
| MRET_1088 | nucleoporin NUP82 | 114.26 | 79.55 |
| MRET_1089 | phosphatidylserine decarboxylase | 510.95 | 232.63 |
| MRET_1090 | uncharacterized protein | 1542.46 | 3133.36 |
| MRET_1091 | Rhodanese-like domain protein | 52.69 | 67.11 |
| MRET_1092 | nucleolar protein 58 | 150.67 | 188.21 |
| MRET_1093 | DNA polymerase alpha subunit B | 95.04 | 104.42 |
| MRET_1094 | general transcription factor 3C polypeptide 5 (transcription factor C subunit 1) | 14.08 | 27.9 |
| MRET_1095 | Ran-binding protein 1 | 151.5 | 363.35 |
| MRET_1096 | p24 family protein beta-1 | 65.35 | 168.27 |
| MRET_1097 | uncharacterized protein | 39.21 | 76.47 |
| MRET_1098 | Fe-S cluster assembly protein DRE2 | 539.84 | 530.56 |
| MRET_1099 | multidrug resistance protein fnx1 | 213.46 | 97.64 |
| MRET_1100 | serine/threonine-protein kinase RIM15 | 133.39 | 63.94 |
| MRET_1101 | uncharacterized protein | 55.34 | 56.25 |
| MRET_1102 | phosphatidate cytidylyltransferase | 34.46 | 129.16 |
| MRET_1103 | serine/threonine-protein kinase | 58.51 | 34.73 |
| MRET_1104 | NADH dehydrogenase (ubiquinone) Fe-S protein 7 | 555.76 | 195.09 |
| MRET_1105 | type I protein arginine methyltransferase | 69.97 | 117.45 |
| MRET_1106 | golgi SNAP receptor complex member 2 | 47.48 | 60.77 |
| MRET_1107 | zinc finger CCHC domain protein 9 | 65.57 | 54.36 |

| | | | |
|---|---|---|---|
| MRET_1108 | tRNA-specific adenosine deaminase 1 | 30.17 | 33.56 |
| MRET_1109 | NAD-dependent deacetylase sirtuin 2 | 107.11 | 180.74 |
| MRET_1110 | uncharacterized protein | 351.08 | 206.85 |
| MRET_1111 | delta7-sterol 5-desaturase | 113.16 | 308.12 |
| MRET_1112 | sorting nexin-3/12 | 49.91 | 85.35 |
| MRET_1113 | large subunit ribosomal protein L7e | 162.95 | 598.57 |
| MRET_1114 | heat shock 70kDa protein 4 | 712.69 | 812.98 |
| MRET_1115 | conserved hypothetical protein | 6966.72 | 3923.27 |
| MRET_1116 | conserved hypothetical protein | 92.24 | 94.88 |
| MRET_1117 | transitional endoplasmic reticulum ATPase | 1374.72 | 1149.33 |
| MRET_1118 | IKS protein kinase | 127.95 | 124.11 |
| MRET_1119 | BRCA1-associated protein | 57.49 | 65.33 |
| MRET_1120 | dolichyl-phosphate mannosyltransferase polypeptide 2, regulatory subunit | 31.12 | 140.32 |
| MRET_1121 | uncharacterized protein | 25.9 | 42.82 |
| MRET_1122 | La domain protein | 23.45 | 46.43 |
| MRET_1123 | lysine methyltransferase | 19.17 | 69.17 |
| MRET_1124 | uncharacterized protein | 81.07 | 204.66 |
| MRET_1125 | HAT1-interacting factor 1 | 224.38 | 245.84 |
| MRET_1126 | adenylylsulfate kinase | 114.49 | 181.18 |
| MRET_1127 | FH domain protein | 23.82 | 41.23 |
| MRET_1128 | OHCU decarboxylase | 40.67 | 64.67 |
| MRET_1129 | cytochrome-b5 reductase | 146.02 | 84.15 |
| MRET_1130 | conserved oligomeric golgi complex subunit 4 | 116.47 | 75.25 |
| MRET_1131 | 3-hydroxy acid dehydrogenase/malonic semialdehyde reductase | 92.72 | 125.33 |
| MRET_1132 | spindle assembly associated Sfi1-like protein | 34.73 | 46.24 |
| MRET_1133 | pre-rRNA-processing protein TSR2 | 26.84 | 40.17 |
| MRET_1134 | programmed cell death 6-interacting protein | 138.99 | 44.21 |
| MRET_1135 | uncharacterized protein | 126.7 | 64.75 |
| MRET_1136 | ubiquitin-conjugating enzyme E2 G2 | 193.9 | 228.78 |
| MRET_1137 | ribosome assembly protein 4 | 41.44 | 63.44 |
| MRET_1138 | essential protein, constituent of 66S pre-ribosomal particles | 32.46 | 64.21 |
| MRET_1139 | DASH complex subunit DAD3 | 469.37 | 351.43 |
| MRET_1140 | protein of unknown function (DUF1769) | 357.46 | 199.47 |
| MRET_1141 | NAD-dependent histone deacetylase SIR2 | 37.35 | 35.08 |
| MRET_1142 | R3H domain protein | 235.56 | 113.91 |
| MRET_1143 | uncharacterized protein | 16.06 | 25.7 |
| MRET_1144 | ubiquitin-activating enzyme E1 C | 112.92 | 78.38 |

| | | | |
|---|---|---|---|
| MRET_1145 | deoxyhypusine synthase | 19.6 | 43.15 |
| MRET_1146 | phosphatidylinositol glycan, class K | 47.18 | 83.65 |
| MRET_1147 | V-type H+-transporting ATPase 21kDa proteolipid subunit | 287.76 | 428.99 |
| MRET_1148 | oleate-activated transcription factor | 27.32 | 54.93 |
| MRET_1149 | transcription elongation factor B, polypeptide 1 | 234.66 | 512.57 |
| MRET_1150 | protein STU1 | 4.24 | 33.65 |
| MRET_1151 | very-long-chain enoyl-CoA reductase | 148.36 | 183.69 |
| MRET_1152 | N-lysine methyltransferase SETD6 | 15.2 | 34.12 |
| MRET_1153 | zinc finger protein, C2H2 type | 25.66 | 32.3 |
| MRET_1154 | beta-glucan synthesis-associated protein KRE6 | 44.41 | 61.24 |
| MRET_1155 | beta-glucan synthesis-associated protein KRE6 | 14.19 | 36 |
| MRET_1156 | flap endonuclease-1 | 33.97 | 112.61 |
| MRET_1157 | uncharacterized protein | 32.67 | 31 |
| MRET_1158 | glutamate-5-semialdehyde dehydrogenase | 301.68 | 147.74 |
| MRET_1159 | acyl-CoA-dependent ceramide synthase | 82.21 | 106.07 |
| MRET_1160 | riboflavin aldehyde-forming enzyme | 153.3 | 124.41 |
| MRET_1161 | Rab geranylgeranyl transferase escort protein | 35.47 | 64.11 |
| MRET_1162 | tripeptidyl-peptidase II | 36.16 | 71.67 |
| MRET_1163 | SET and MYND domain protein | 25.25 | 19.21 |
| MRET_1164 | G1 S-specific cyclin Pcl5 | 4418.67 | 4701.9 |
| MRET_1165 | RTA1 domain protein | 157.96 | 412.85 |
| MRET_1166 | proliferating cell nuclear antigen | 395.75 | 946.27 |
| MRET_1167 | magnesium transporter | 16.84 | 41.56 |
| MRET_1168 | cell division control protein | 58.31 | 188 |
| MRET_1169 | cytochrome c oxidase subunit 7c | 164.68 | 169.42 |
| MRET_1170 | ribosome production factor 1 | 262.35 | 190.05 |
| MRET_1171 | nuclear pore complex protein Nup107 | 62.63 | 134.47 |
| MRET_1172 | transcription initiation factor TFIIB | 371.73 | 282.03 |
| MRET_1173 | zinc finger protein | 79.41 | 173.22 |
| MRET_1174 | histidine kinase | 22.77 | 33.44 |
| MRET_1175 | uncharacterized protein | 20.44 | 41.51 |
| MRET_1176 | small subunit ribosomal protein S7 | 46.39 | 92.21 |
| MRET_1177 | alkaline phosphatase D | 68.24 | 92.66 |
| MRET_1178 | glutamine amidotransferase/cyclase | 129.2 | 127.52 |
| MRET_1179 | secretory lipase | 396.83 | 221.83 |
| MRET_1180 | uncharacterized protein | 90.79 | 49.31 |
| MRET_1181 | 1-phosphatidylinositol-4-phosphate 5-kinase | 357.43 | 212.88 |

| | | | |
|---|---|---|---|
| MRET_1182 | DNA primase large subunit | 89.07 | 68.84 |
| MRET_1183 | U3 small nucleolar RNA-associated protein 6 | 236.58 | 150.41 |
| MRET_1184 | mitochondrial ATPase complex subunit ATP10 | 144.22 | 191.15 |
| MRET_1185 | Ras-related protein Rab-8A | 311.38 | 550.99 |
| MRET_1186 | uncharacterized protein | 29.85 | 50.78 |
| MRET_1187 | uncharacterized protein | 642.76 | 697.89 |
| MRET_1188 | vacuolar protein sorting-associated protein 11 | 39.71 | 32.14 |
| MRET_1189 | long-chain acyl-CoA synthetase | 92.28 | 65.9 |
| MRET_1190 | squalene monooxygenase | 41.87 | 102.06 |
| MRET_1191 | U3 small nucleolar RNA-associated protein 23 | 205.07 | 226.89 |
| MRET_1192 | leucine-carboxy methyltransferase | 95.35 | 60.62 |
| MRET_1193 | solute carrier family 25 (mitochondrial carnitine/acylcarnitine transporter), member 20/29 | 174.72 | 94.24 |
| MRET_1194 | uncharacterized protein | 336.21 | 167.59 |
| MRET_1195 | nuclear pore complex protein Nup155 | 51.17 | 35.35 |
| MRET_1196 | elongation factor Ts | 98.35 | 95.05 |
| MRET_1197 | Ras-related protein Rab-7A | 529.59 | 452.76 |
| MRET_1198 | Ras-related GTP-binding protein C/D | 361.92 | 194.97 |
| MRET_1199 | translation machinery-associated protein 22 | 291.51 | 195.34 |
| MRET_1200 | COMPASS component BRE2 | 31.87 | 31.62 |
| MRET_1201 | maintenance of mitochondrial morphology protein 1 | 63.66 | 38.02 |
| MRET_1202 | DnaJ chaperone Caj1 | 63.69 | 45.23 |
| MRET_1203 | NADPH-dependent beta-ketoacyl reductase | 517.92 | 314.92 |
| MRET_1204 | complex 1 protein (LYR family) | 22.08 | 32.23 |
| MRET_1205 | zinc finger protein, C2H2 type | 204.89 | 177.44 |
| MRET_1206 | phosphopantothenate---cysteine ligase (ATP) | 93.24 | 88.57 |
| MRET_1207 | DNA methyltransferase 1-associated protein 1 | 30.29 | 36.97 |
| MRET_1208 | uncharacterized protein | 21.51 | 24.99 |
| MRET_1209 | galactokinase | 227 | 173.15 |
| MRET_1210 | GINS complex subunit 4 | 85.16 | 91.37 |
| MRET_1211 | putative DNA replication factor C complex subunit Ctf8 | 129.13 | 74.28 |
| MRET_1212 | GATA zinc finger | 93.66 | 46.64 |
| MRET_1213 | transcription initiation factor TFIIA small subunit | 135.59 | 252.34 |
| MRET_1214 | centrin-3 | 126.7 | 211.08 |
| MRET_1215 | uncharacterized protein | 22.66 | 34.55 |
| MRET_1216 | mitochondrial export protein SOM1 | 85.68 | 98.34 |
| MRET_1217 | adenosylhomocysteinase | 114.14 | 193.4 |
| MRET_1218 | Xaa-Pro aminopeptidase | 240.43 | 394.24 |

| | | | |
|---|---|---|---|
| MRET_1219 | serine/threonine-protein kinase 24/25/MST4 | 24.45 | 27.62 |
| MRET_1220 | uncharacterized protein | 541.4 | 251.75 |
| MRET_1221 | uncharacterized protein | 344.15 | 318.23 |
| MRET_1222 | uncharacterized protein | 109.89 | 128.17 |
| MRET_1223 | N-acetyl-gamma-glutamyl-phosphate reductase/acetylglutamate kinase | 45.67 | 78.37 |
| MRET_1224 | ER-derived vesicles protein | 51.56 | 92.96 |
| MRET_1225 | phosphatidylinositol transfer protein | 1013.18 | 413.3 |
| MRET_1226 | serine/threonine-protein phosphatase 2A regulatory subunit A | 129.06 | 114.87 |
| MRET_1227 | DNA polymerase zeta | 75.47 | 42.36 |
| MRET_1228 | phosphatidylethanolamine N-methyltransferase | 36.73 | 62.91 |
| MRET_1229 | 2,3-bisphosphoglycerate-dependent phosphoglycerate mutase | 192.59 | 245.96 |
| MRET_1230 | zinc finger protein, GATA type | 45.38 | 120.3 |
| MRET_1231 | protein of unknown function (DUF2841) | 2.7 | 13.54 |
| MRET_1232 | ubiquitin-conjugating enzyme E2 N | 155.55 | 164.5 |
| MRET_1233 | Wiskott-Aldrich syndrome protein | 43.31 | 38.03 |
| MRET_1234 | elongation factor 1-alpha | 1299.6 | 1982.16 |
| MRET_1235 | RAT1-interacting protein | 25.74 | 36.86 |
| MRET_1236 | solute carrier family 30 (zinc transporter), member 1 | 1163.45 | 735.31 |
| MRET_1237 | uncharacterized protein | 38.56 | 45.36 |
| MRET_1238 | conserved hypothetical protein | 10.58 | 17.08 |
| MRET_1239 | transcription initiation factor TFIID subunit 7 | 113.47 | 90.25 |
| MRET_1240 | DNA-directed RNA polymerase III subunit RPC5 | 350.36 | 270.9 |
| MRET_1241 | nucleoside-diphosphate kinase | 468.7 | 361.18 |
| MRET_1242 | exosome complex exonuclease DIS3/RRP44 | 330.85 | 80.15 |
| MRET_1243 | sulfite reductase (NADPH) hemoprotein beta-component | 323.48 | 157.66 |
| MRET_1244 | mannose-1-phosphate guanylyltransferase | 43 | 84.08 |
| MRET_1245 | large subunit ribosomal protein L36 | 59.55 | 68.37 |
| MRET_1246 | Urb2/Npa2 family protein | 4.48 | 15.79 |
| MRET_1247 | transporter | 0 | 50.89 |
| MRET_1248 | transporter | 96.45 | 113.32 |
| MRET_1249 | SMR domain protein | 77.81 | 68.83 |
| MRET_1250 | mitotic cell cycle regulation protein | 214.85 | 146.28 |
| MRET_1251 | small subunit ribosomal protein S17 | 77.83 | 82.01 |
| MRET_1252 | uncharacterized protein | 27.02 | 76.33 |
| MRET_1253 | serine/threonine-protein kinase 16 | 86.15 | 35.18 |
| MRET_1254 | subunit of COMPASS (Set1C) | 45.51 | 26.45 |
| MRET_1255 | DUF543 domain protein | 165.9 | 180.86 |

| | | | |
|---|---|---|---|
| MRET_1256 | putative ATPase | 95.5 | 65.34 |
| MRET_1257 | E3 SUMO-protein ligase PIAS1 | 29.91 | 35.99 |
| MRET_1258 | SAGA complex subunit spt20 | 152.02 | 206.8 |
| MRET_1259 | conserved hypothetical protein | 251.05 | 227.94 |
| MRET_1260 | chromosome transmission fidelity protein 4 | 17.53 | 34.16 |
| MRET_1261 | phosphoserine aminotransferase | 263.65 | 184.11 |
| MRET_1262 | transcription initiation factor TFIIF subunit beta | 62.19 | 71.77 |
| MRET_1263 | zinc finger protein (RING finger) | 84.42 | 58.42 |
| MRET_1264 | heat shock 70kDa protein 1/2/6/8 | 3323.97 | 4615.66 |
| MRET_1265 | microtubule binding protein HOOK3 | 67.37 | 100.3 |
| MRET_1266 | uncharacterized protein | 31.77 | 30.36 |
| MRET_1267 | E3 ubiquitin-protein ligase HUWE1 | 252.12 | 148.25 |
| MRET_1268 | cullin 4 | 27.19 | 27.88 |
| MRET_1269 | biotin---protein ligase | 62.76 | 46.59 |
| MRET_1270 | 4-nitrophenyl phosphatase | 49.02 | 61.94 |
| MRET_1271 | ion transport protein | 210.78 | 311.66 |
| MRET_1272 | casein kinase I | 712.97 | 838.87 |
| MRET_1273 | transporter | 21.71 | 31.05 |
| MRET_1274 | U3 small nucleolar RNA-associated protein 11 | 18.39 | 69.61 |
| MRET_1275 | conserved hypothetical protein | 18.8 | 64.71 |
| MRET_1276 | 26S proteasome non-ATPase regulatory subunit 9 | 28.36 | 63.12 |
| MRET_1277 | uncharacterized protein | 463.83 | 251.49 |
| MRET_1278 | S-(hydroxymethyl)glutathione dehydrogenase/alcohol dehydrogenase | 1143.47 | 687.03 |
| MRET_1279 | carbohydrate kinase | 41.74 | 58.86 |
| MRET_1280 | ceramide glucosyltransferase | 37.85 | 47.17 |
| MRET_1281 | ubiquitin-conjugating enzyme E2 I | 54.6 | 156.33 |
| MRET_1282 | tropomyosin, fungi type | 98.5 | 261.95 |
| MRET_1283 | serine/threonine-protein kinase | 280.42 | 166.92 |
| MRET_1284 | PHD finger domain protein | 19.75 | 44.57 |
| MRET_1285 | uncharacterized protein | 43.62 | 93.06 |
| MRET_1286 | negative regulator of eIF2 kinase Gcn2p | 186.73 | 216.62 |
| MRET_1287 | FUN14 family protein | 1310.68 | 582.57 |
| MRET_1288 | fungal protein of unknown function (DUF2015) | 364.39 | 231.27 |
| MRET_1289 | trafficking protein particle complex subunit 12 | 32.95 | 41.29 |
| MRET_1290 | histone acetyltransferase 1 | 37.94 | 50.86 |
| MRET_1291 | type 2A phosphatase activator TIP41 | 163.54 | 165.12 |
| MRET_1292 | cell cycle checkpoint control protein RAD9A | 48.26 | 79.88 |

| | | | |
|---|---|---|---|
| MRET_1293 | p21-activated kinase 1 | 28.7 | 80.71 |
| MRET_1294 | cryptococcal mannosyltransferase 1 | 30.05 | 26.62 |
| MRET_1295 | conserved oligomeric golgi complex subunit 2 | 107.13 | 52.58 |
| MRET_1296 | UDP-N-acetylglucosamine/UDP-N-acetylgalactosamine diphosphorylase | 35.08 | 63.94 |
| MRET_1297 | uncharacterized protein | 30.87 | 62.53 |
| MRET_1298 | heat shock 70kDa protein 5 | 510.56 | 726.7 |
| MRET_1299 | peroxin-10 | 65.02 | 104.15 |
| MRET_1300 | cysteine-rich secretory protein | 2358.57 | 2077.02 |
| MRET_1301 | sphinganine C4-monooxygenase | 625.2 | 308.25 |
| MRET_1302 | osomolarity two-component system, response regulator SSK1 | 38.98 | 42.06 |
| MRET_1303 | nuclear control of ATPase protein 2 | 40.18 | 55.16 |
| MRET_1304 | glycylpeptide N-tetradecanoyltransferase | 69.16 | 106.35 |
| MRET_1305 | pre-mRNA-splicing factor RBM22/SLT11 | 100.65 | 106.2 |
| MRET_1306 | LYR motif containing 7 | 15.98 | 63.05 |
| MRET_1307 | E3 ubiquitin-protein ligase NEDD4 | 460.63 | 419.51 |
| MRET_1308 | integral membrane protein | 16.86 | 79.64 |
| MRET_1309 | histone deacetylase | 32.66 | 109.52 |
| MRET_1310 | chitin deacetylase | 44.1 | 295.46 |
| MRET_1311 | LETM1 and EF-hand domain protein 1, mitochondrial | 123.56 | 189.86 |
| MRET_1312 | ribosomal RNA-processing protein 12 | 33.75 | 42.41 |
| MRET_1313 | putative ferric reductase with similarity to Fre2p | 183.5 | 379.84 |
| MRET_1314 | phosphatidylinositol glycan, class T | 354.03 | 647.68 |
| MRET_1315 | protein ROT1 | 33.95 | 64.05 |
| MRET_1316 | uncharacterized protein | 15.2 | 24.12 |
| MRET_1317 | coatomer subunit alpha | 90.01 | 116.24 |
| MRET_1318 | mitochondrial ribosomal protein subunit | 34.01 | 81.89 |
| MRET_1319 | 6,7-dimethyl-8-ribityllumazine synthase | 69.75 | 172.03 |
| MRET_1320 | 3-hydroxyisobutyryl-CoA hydrolase | 130.67 | 116.96 |
| MRET_1321 | asparaginyl-tRNA synthetase | 101.18 | 119.83 |
| MRET_1322 | cell division cycle protein | 47.5 | 47.56 |
| MRET_1323 | nuclear cap-binding protein subunit 2 | 58.57 | 91.89 |
| MRET_1324 | nascent polypeptide-associated complex subunit beta | 31.52 | 89.69 |
| MRET_1325 | F-type H+-transporting ATP synthase subunit e | 351.12 | 443.32 |
| MRET_1326 | acetyl-CoA synthetase | 87.56 | 159.56 |
| MRET_1327 | replication factor A3 | 77.44 | 205.45 |
| MRET_1328 | sensitive to high expression protein 9, mitochondrial | 33.37 | 52.42 |
| MRET_1329 | elongation factor 3 | 73.77 | 112.78 |

| | | | |
|---|---|---|---|
| MRET_1330 | DNA excision repair protein ERCC-2 | 13.61 | 32.59 |
| MRET_1331 | uncharacterized protein | 24 | 30.35 |
| MRET_1332 | Rab guanine nucleotide exchange factor SEC2 | 41.3 | 52.23 |
| MRET_1333 | uncharacterized protein | 440.74 | 372.46 |
| MRET_1334 | 5-oxoprolinase (ATP-hydrolysing) | 354.34 | 369.37 |
| MRET_1335 | mitogen-activated protein kinase kinase kinase | 340.06 | 219.77 |
| MRET_1336 | manganese-transporting P-type ATPase | 34.88 | 67.86 |
| MRET_1337 | solute carrier family 25 (mitochondrial dicarboxylate transporter), member 10 | 15.58 | 35.61 |
| MRET_1338 | uncharacterized protein | 87.07 | 61.99 |
| MRET_1339 | Rab6A-GEF complex partner protein 1 | 13.46 | 20.79 |
| MRET_1340 | mitogen-activated protein kinase kinase kinase | 36.36 | 40.57 |
| MRET_1341 | uncharacterized protein | 50.61 | 68.38 |
| MRET_1342 | synaptobrevin homolog YKT6 | 193.71 | 381.41 |
| MRET_1343 | Vps51/Vps67 family protein | 308.34 | 150.12 |
| MRET_1344 | PHD finger domain protein | 34.51 | 43.28 |
| MRET_1345 | uncharacterized protein | 107.19 | 98.58 |
| MRET_1346 | uncharacterized protein | 13.21 | 83.01 |
| MRET_1347 | 2-oxoglutarate dehydrogenase E2 component (dihydrolipoamide succinyltransferase) | 1188.23 | 1063.41 |
| MRET_1348 | solute carrier family 25, member 38 | 18.72 | 31.21 |
| MRET_1349 | RasGEF | 6.95 | 15.58 |
| MRET_1350 | mitogen-activated protein kinase kinase | 28.87 | 62.36 |
| MRET_1351 | NADH dehydrogenase (ubiquinone) 1 alpha subcomplex subunit 6 | 64.16 | 76.14 |
| MRET_1352 | origin recognition complex subunit 2 | 24.83 | 25.46 |
| MRET_1353 | Integral ER membrane protein | 364.05 | 375.76 |
| MRET_1354 | uncharacterized protein | 113.63 | 110.08 |
| MRET_1355 | V-type H+-transporting ATPase subunit B | 185.6 | 262.8 |
| MRET_1356 | peptidyl-tRNA hydrolase ICT1 | 67.42 | 94.95 |
| MRET_1357 | large subunit ribosomal protein L28 | 30.11 | 100.59 |
| MRET_1358 | U3 small nucleolar ribonucleoprotein protein IMP4 | 324.42 | 158.36 |
| MRET_1359 | glycerol-3-phosphate dehydrogenase | 513.44 | 202.34 |
| MRET_1360 | cell cycle checkpoint protein | 38.95 | 33.86 |
| MRET_1361 | importin-7 | 16.71 | 24.45 |
| MRET_1362 | succinate dehydrogenase assembly factor 2 | 11.78 | 35.68 |
| MRET_1363 | large subunit ribosomal protein L37Ae | 65.54 | 366.7 |
| MRET_1364 | DnaJ homolog subfamily C member 19 | 526.01 | 348.2 |
| MRET_1365 | uncharacterized protein | 8.32 | 6.67 |
| MRET_1366 | WD domain, G-beta repeat protein | 34.12 | 30.09 |

| | | | |
|---|---|---|---|
| MRET_1367 | protein transport protein SEC23 | 119.42 | 157.12 |
| MRET_1368 | DEP domain protein 5 | 215.93 | 143.82 |
| MRET_1369 | Aur1-inositol phosphorylceramide synthase | 31.48 | 51.31 |
| MRET_1370 | uncharacterized protein | 257.68 | 289.74 |
| MRET_1371 | mitochondrial inner membrane protease subunit 1 | 20.84 | 35.15 |
| MRET_1372 | solute carrier family 25 (mitochondrial folate transporter), member 32 | 30.36 | 73.76 |
| MRET_1373 | PIN domain protein | 26.67 | 62.74 |
| MRET_1374 | ubiquitin-like protein 5 | 156.86 | 193.23 |
| MRET_1375 | translation initiation factor 4E | 582.48 | 267.23 |
| MRET_1376 | uncharacterized protein | 14.74 | 16.87 |
| MRET_1377 | histidinol-phosphatase (PHP family) | 98.59 | 76.2 |
| MRET_1378 | succinate dehydrogenase (ubiquinone) iron-sulfur subunit | 269.8 | 76.09 |
| MRET_1379 | DNA ligase 1 | 20.85 | 26.18 |
| MRET_1380 | mitochondrial inner membrane protein subunit 18 | 50.46 | 64.44 |
| MRET_1381 | uncharacterized protein | 14.13 | 19.42 |
| MRET_1382 | Spo7-like protein | 287.77 | 185.41 |
| MRET_1383 | recyclin-1 | 24.79 | 29.14 |
| MRET_1384 | cell division protein | 159.34 | 107.34 |
| MRET_1385 | ribosome biogenesis protein Sqt1 | 415.79 | 237.36 |
| MRET_1386 | ribose 5-phosphate isomerase A | 89.15 | 73.64 |
| MRET_1387 | 25S rRNA (adenine2142-N1)-methyltransferase | 38.27 | 45.98 |
| MRET_1388 | glutaredoxin 3 | 3293.49 | 2176.85 |
| MRET_1389 | autophagy-related protein 18 | 147.21 | 349.41 |
| MRET_1390 | ribosomal RNA-processing protein 9 | 27.25 | 45.4 |
| MRET_1391 | telomerase activating protein Est1 | 61.4 | 39.97 |
| MRET_1392 | COP9 signalosome complex subunit 1 | 84.39 | 44.79 |
| MRET_1393 | COP9 signalosome complex subunit 4 | 87.18 | 57.34 |
| MRET_1394 | Rwd-domain protein | 73.95 | 128.81 |
| MRET_1395 | molecular chaperone DnaK | 2405.91 | 2020.29 |
| MRET_1396 | PX domain protein | 105.43 | 142.69 |
| MRET_1397 | mitochondrial import receptor subunit TOM20 | 150.24 | 190.19 |
| MRET_1398 | choline/ethanolamine kinase | 13.26 | 29.34 |
| MRET_1399 | uncharacterized protein | 79.02 | 53.83 |
| MRET_1400 | poly(ADP-ribose) polymerase | 319.27 | 525.55 |
| MRET_1401 | smad nuclear-interacting protein 1 | 259.59 | 233.01 |
| MRET_1402 | tRNA ligase | 35.78 | 48.51 |
| MRET_1403 | multicopper oxidase | 71.64 | 52.88 |

| | | | |
|---|---|---|---|
| MRET_1404 | multicopper oxidase | 44.66 | 53.41 |
| MRET_1405 | phospholipase C | 48.85 | 98.49 |
| MRET_1406 | phospholipase C | 21.33 | 45.79 |
| MRET_1407 | phospholipase C | 155.7 | 123.21 |
| MRET_1408 | phospholipase C | 69 | 79.23 |
| MRET_1409 | uncharacterized protein | 110.46 | 161.79 |
| MRET_1410 | histone-lysine N-methyltransferase SETD1 | 69.34 | 205.94 |
| MRET_1411 | DDHD domain protein | 81.41 | 100.99 |
| MRET_1412 | fluoride exporter | 51.02 | 75.57 |
| MRET_1413 | uncharacterized protein | 19.74 | 36.14 |
| MRET_1414 | cell wall protein | 17.27 | 11.49 |
| MRET_1415 | uncharacterized protein | 26.89 | 26.11 |
| MRET_1416 | small subunit ribosomal protein S26e | 371.08 | 882.48 |
| MRET_1417 | cleavage and polyadenylation specificity factor subunit 1 | 40.37 | 58.25 |
| MRET_1418 | WW domain-binding protein 4 | 18.83 | 45.16 |
| MRET_1419 | transcription factor | 60.53 | 74.92 |
| MRET_1420 | tubulin alpha | 76.81 | 110.55 |
| MRET_1421 | solute carrier family 25 (mitochondrial phosphate transporter), member 23/24/25/41 | 91.19 | 158.71 |
| MRET_1422 | acyl-CoA oxidase | 1033.96 | 733.36 |
| MRET_1423 | lactoylglutathione lyase | 75.54 | 172.63 |
| MRET_1424 | bZIP transcription factor | 104.01 | 119.87 |
| MRET_1425 | coatomer subunit gamma | 63.25 | 79.13 |
| MRET_1426 | small subunit ribosomal protein S28e | 82.04 | 411.43 |
| MRET_1427 | peroxisomal membrane anchor protein PEX14p | 67.97 | 78.34 |
| MRET_1428 | DNA helicase INO80 | 77.9 | 55.44 |
| MRET_1429 | splicing factor 3B subunit 5 | 174.25 | 171.89 |
| MRET_1430 | 3-oxo-5-alpha-steroid 4-dehydrogenase 1 | 120.96 | 99.85 |
| MRET_1431 | CTP synthase | 46.85 | 74.07 |
| MRET_1432 | uncharacterized protein | 110.76 | 287.47 |
| MRET_1433 | zinc finger protein | 15.65 | 23.01 |
| MRET_1434 | uncharacterized protein | 28.93 | 49.62 |
| MRET_1435 | pre-mRNA-splicing factor CWC26 | 18.27 | 35.15 |
| MRET_1436 | transcription initiation factor TFIIIB component B'' | 69.37 | 104.14 |
| MRET_1437 | uncharacterized protein | 235.35 | 260.94 |
| MRET_1438 | translation machinery-associated protein 16 | 69.81 | 137.46 |
| MRET_1439 | conserved hypothetical protein | 29.86 | 19.05 |
| MRET_1440 | inositol-pentakisphosphate 2-kinase | 6 | 10.16 |

| | | | |
|---|---|---|---|
| MRET_1441 | SRP40, C-terminal domain protein | 20.88 | 32.05 |
| MRET_1442 | tRNASer (uridine44-2'-O)-methyltransferase | 13.65 | 19.72 |
| MRET_1443 | splicing factor 3B subunit 3 | 55.04 | 105.32 |
| MRET_1444 | RhoGEF domain protein | 20.21 | 42.05 |
| MRET_1445 | oxidation resistance protein 1 | 34.43 | 46.89 |
| MRET_1446 | uncharacterized protein | 60.75 | 51.55 |
| MRET_1447 | F-box domain protein | 62.4 | 77.9 |
| MRET_1448 | tubulin beta | 561.68 | 476.46 |
| MRET_1449 | aspartyl-tRNA synthetase | 50.54 | 85.72 |
| MRET_1450 | acylglycerol lipase | 85.73 | 119.58 |
| MRET_1451 | carnitine O-acetyltransferase | 184.12 | 156.48 |
| MRET_1452 | uncharacterized protein | 269.35 | 329.3 |
| MRET_1453 | protein kinase | 68.51 | 64.4 |
| MRET_1454 | heparinase II III family protein | 221.1 | 226.74 |
| MRET_1455 | AMP-binding enzyme | 19.96 | 28.91 |
| MRET_1456 | DNA-directed RNA polymerases I, II, and III subunit RPABC2 | 307.03 | 670.95 |
| MRET_1457 | zinc finger protein, C2H2 type | 225.18 | 198.57 |
| MRET_1458 | DNA mismatch repair protein MSH3 | 5.22 | 12.63 |
| MRET_1459 | geranylgeranyl transferase type-2 subunit beta | 158.02 | 181.79 |
| MRET_1460 | AHNAK nucleoprotein | 25 | 33.29 |
| MRET_1461 | pre-rRNA-processing protein TSR3 | 137.35 | 116.28 |
| MRET_1462 | isocitrate lyase | 620.96 | 392.67 |
| MRET_1463 | integral membrane protein (Ptm1) | 52.46 | 148.62 |
| MRET_1464 | SEL1 domain protein | 309.08 | 199.99 |
| MRET_1465 | peroxisomal biogenesis factor | 349.56 | 489.23 |
| MRET_1466 | translation initiation factor 3 subunit B | 88.05 | 179.16 |
| MRET_1467 | AdoMet-dependent methyltransferase SPB1 | 61.48 | 107.24 |
| MRET_1468 | large subunit ribosomal protein L22e | 160.63 | 987.68 |
| MRET_1469 | H/ACA ribonucleoprotein complex subunit 4 | 781.43 | 720.3 |
| MRET_1470 | cytochrome c oxidase subunit 11 | 150.02 | 311.2 |
| MRET_1471 | small subunit ribosomal protein S5e | 602.71 | 1230.58 |
| MRET_1472 | transducin (beta)-like 1 | 104.41 | 113.33 |
| MRET_1473 | rhomboid family protein | 319.11 | 161.52 |
| MRET_1474 | PPR repeat containing protein | 26.45 | 46.63 |
| MRET_1475 | mitochondrial 54S ribosomal protein YmL8 | 23.75 | 48.88 |
| MRET_1476 | NADH dehydrogenase (ubiquinone) 1 alpha/beta subcomplex 1 | 1421.6 | 945.26 |
| MRET_1477 | serine/threonine-protein kinase SCH9 | 345.09 | 519.26 |

| | | | |
|---|---|---|---|
| MRET_1478 | N-alpha-acetyltransferase 15/16, NatA auxiliary subunit | 18.03 | 34.59 |
| MRET_1479 | elongation factor 2 | 148.93 | 305.42 |
| MRET_1480 | prefoldin subunit 4 | 20.66 | 45.46 |
| MRET_1481 | superkiller protein 3 | 18.18 | 23.19 |
| MRET_1482 | UDP-N-acetylglucosamine--dolichyl-phosphate N-acetylglucosaminephosphotransferase | 22.36 | 33.79 |
| MRET_1483 | inosine triphosphate pyrophosphatase | 36.85 | 72.19 |
| MRET_1484 | NAD dependent epimerase dehydratase family protein | 876.25 | 322.07 |
| MRET_1485 | phosphatidylinositol 4-phosphatase | 114.19 | 322.69 |
| MRET_1486 | pre-mRNA-processing factor 19 | 78.87 | 156.59 |
| MRET_1487 | anthranilate synthase/indole-3-glycerol phosphate synthase/phosphoribosylanthranilate isomerase | 108.85 | 224.91 |
| MRET_1488 | actin-related protein 9 | 67.32 | 92.75 |
| MRET_1489 | phosphoacetylglucosamine mutase | 209.38 | 187.46 |
| MRET_1490 | mannose-P-dolichol utilization defect 1 | 106.43 | 124.33 |
| MRET_1491 | translation initiation factor 3 subunit H | 75.72 | 181.62 |
| MRET_1492 | zinc finger protein, C2H2 type | 18.23 | 33.55 |
| MRET_1493 | RNA polymerase II subunit A C-terminal domain phosphatase SSU72 | 194.05 | 260.45 |
| MRET_1494 | recombining binding protein suppressor of hairless | 108.74 | 83.14 |
| MRET_1495 | ribosomal biogenesis protein LAS1 | 65.08 | 94.69 |
| MRET_1496 | conserved hypothetical protein | 1979.36 | 988.17 |
| MRET_1497 | phosphoribosylaminoimidazole-succinocarboxamide synthase | 57.54 | 109.29 |
| MRET_1498 | kinesin family member 11 | 18.24 | 53.82 |
| MRET_1499 | COMPASS component SWD3 | 35 | 59.92 |
| MRET_1500 | DNA-directed RNA polymerase II subunit RPB4 | 108.09 | 144.12 |
| MRET_1501 | mitochondrial serine protease | 52.46 | 123.44 |
| MRET_1502 | kynurenine aminotransferase | 622.51 | 515.67 |
| MRET_1503 | sorting nexin | 334.23 | 423.41 |
| MRET_1504 | serine/threonine-protein phosphatase | 223.13 | 231.33 |
| MRET_1505 | translation initiation factor 5B | 131.87 | 144.47 |
| MRET_1506 | carbamoyl-phosphate synthase large subunit | 76.68 | 28.33 |
| MRET_1507 | alpha/beta-hydrolase lipase | 37.88 | 58.25 |
| MRET_1508 | uncharacterized protein | 475.91 | 370.77 |
| MRET_1509 | complement component 1 Q subcomponent-binding protein, mitochondrial | 124.42 | 150.9 |
| MRET_1510 | disulfide isomerase | 194.2 | 248.58 |
| MRET_1511 | rapamycin-insensitive companion of mTOR | 30.96 | 47.16 |
| MRET_1512 | cytochrome c oxidase assembly factor 1 | 72.43 | 166.12 |
| MRET_1513 | mannoprotein MP88 | 2080.55 | 1306.28 |
| MRET_1514 | small subunit ribosomal protein S16 | 111.89 | 158.49 |

| | | | |
|---|---|---|---|
| MRET_1515 | AHNAK nucleoprotein | 159.33 | 111.66 |
| MRET_1516 | conserved hypothetical protein | 27.93 | 44.57 |
| MRET_1517 | membrane magnesium transporter | 134.26 | 257.19 |
| MRET_1518 | actin beta/gamma 1 | 2269.4 | 1819.97 |
| MRET_1519 | nucleolar protein 56 | 80.71 | 178.67 |
| MRET_1520 | 20S proteasome subunit alpha 1 | 139.82 | 217.67 |
| MRET_1521 | RNA polymerase-associated protein CTR9 | 39.64 | 56.91 |
| MRET_1522 | 5'-phosphate synthase pdxT subunit | 98.75 | 97.83 |
| MRET_1523 | MT-A70 family protein | 34.93 | 35.63 |
| MRET_1524 | T-complex protein 1 subunit beta | 129.86 | 243.03 |
| MRET_1525 | cytochrome c oxidase subunit 4 | 351.97 | 299.24 |
| MRET_1526 | O-acetylhomoserine/O-acetylserine sulfhydrylase | 279.87 | 478.24 |
| MRET_1527 | cystathionine beta-lyase | 73.98 | 63.71 |
| MRET_1528 | RNA 3'-terminal phosphate cyclase-like protein | 24.31 | 42.67 |
| MRET_1529 | conserved hypothetical protein | 223.93 | 203.1 |
| MRET_1530 | microfibrillar-associated protein 1 | 168.63 | 118.87 |
| MRET_1531 | ADP-ribosylation factor | 72.11 | 67.81 |
| MRET_1532 | helicase SWR1 | 186.96 | 124.28 |
| MRET_1533 | F-type H+-transporting ATPase subunit epsilon | 238.09 | 334.99 |
| MRET_1534 | protein of unknown function (DUF423) | 1458.25 | 1234.35 |
| MRET_1535 | uncharacterized protein | 26.89 | 48.15 |
| MRET_1536 | TBC1 domain family, member 13 | 43.25 | 40.5 |
| MRET_1537 | LEM3 CDC50 family protein | 26.56 | 65.12 |
| MRET_1538 | GTP-binding protein 1 | 570.91 | 332.28 |
| MRET_1539 | tyrosine-protein phosphatase SIW14 | 164.66 | 368.14 |
| MRET_1540 | oxysterol-binding protein-related protein 8 | 212.33 | 485.76 |
| MRET_1541 | transcriptional co-repressor | 20.89 | 43.65 |
| MRET_1542 | pyridoxal 5'-phosphate synthase pdxS subunit | 158.1 | 225.91 |
| MRET_1543 | ubiquinol-cytochrome c reductase iron-sulfur subunit | 473.57 | 628.3 |
| MRET_1544 | FACT complex subunit SPT16 | 83.47 | 232.49 |
| MRET_1545 | GABA(A) receptor-associated protein | 79.23 | 301.88 |
| MRET_1546 | serine/threonine-protein phosphatase 4 regulatory subunit 1 | 29.2 | 56.78 |
| MRET_1547 | oligopeptide transporter | 43.79 | 95.71 |
| MRET_1548 | phosphodiesterase | 712.63 | 618.01 |
| MRET_1549 | glutamate synthase (NADPH/NADH) | 98.06 | 72.65 |
| MRET_1550 | U4/U6 small nuclear ribonucleoprotein PRP31 | 43.69 | 74.02 |
| MRET_1551 | adenylosuccinate synthase | 93.41 | 127.17 |

| | | | |
|---|---|---|---|
| MRET_1552 | succinate dehydrogenase (ubiquinone) membrane anchor subunit | 139.39 | 100.23 |
| MRET_1553 | DNA-directed RNA polymerase II subunit RPB9 | 148 | 465.75 |
| MRET_1554 | pyruvate dehydrogenase E1 component alpha subunit | 739.04 | 547.25 |
| MRET_1555 | protein HPT1 | 53.44 | 73.28 |
| MRET_1556 | DUF962 domain protein | 146.65 | 227.12 |
| MRET_1557 | trafficking protein particle complex subunit 13 | 16.96 | 18 |
| MRET_1558 | conserved hypothetical protein | 40.18 | 46.26 |
| MRET_1559 | E3 ubiquitin-protein ligase ZNF598 | 40.79 | 62.14 |
| MRET_1560 | DASH complex subunit DAD1 | 66.33 | 78.84 |
| MRET_1561 | tRNA (guanine10-N2)-methyltransferase | 159.09 | 172.51 |
| MRET_1562 | ATP dependent DNA ligase domain protein | 42.58 | 44.49 |
| MRET_1563 | uncharacterized protein | 20.91 | 25.91 |
| MRET_1564 | V-type H+-transporting ATPase subunit H | 92.85 | 95.61 |
| MRET_1565 | large subunit ribosomal protein L46 | 171.37 | 152.66 |
| MRET_1566 | 5'-AMP-activated protein kinase, regulatory gamma subunit | 60.46 | 90.36 |
| MRET_1567 | cytochrome b pre-mRNA-processing protein 3 | 1027.85 | 458.79 |
| MRET_1568 | phosphoribosylformimino-5-aminoimidazole carboxamide ribotide isomerase | 362.98 | 485.56 |
| MRET_1569 | nuclear pore complex protein Nup93 | 72.61 | 150.34 |
| MRET_1570 | serine/threonine-protein kinase SRPK3 | 429.14 | 305.31 |
| MRET_1571 | ubiquitin-conjugating enzyme | 83.74 | 290.5 |
| MRET_1572 | uncharacterized protein | 32.83 | 73.64 |
| MRET_1573 | 6-phosphogluconolactonase | 68.11 | 89.78 |
| MRET_1574 | threonyl-tRNA synthetase | 32.64 | 119.97 |
| MRET_1575 | component of the SMC5-SMC6 complex | 60.65 | 212.11 |
| MRET_1576 | exocyst complex component 6 | 47.32 | 80.77 |
| MRET_1577 | ATP-dependent RNA helicase DDX41 | 30.82 | 71.71 |
| MRET_1578 | ADP-ribosylation factor 6 | 45.81 | 73.84 |
| MRET_1579 | dynactin 1 | 33.72 | 42.38 |
| MRET_1580 | ATP-dependent Lon protease | 52.9 | 42.69 |
| MRET_1581 | glutamyl-tRNA synthetase | 47 | 43.72 |
| MRET_1582 | Ras-induced vulval development antagonist | 86.67 | 52.54 |
| MRET_1583 | translation initiation factor 5 | 67.7 | 137.91 |
| MRET_1584 | trafficking protein particle complex subunit 5 | 49.78 | 88.51 |
| MRET_1585 | serine/threonine-protein kinase/endoribonuclease IRE1 | 19.72 | 33.67 |
| MRET_1586 | arsenite/tail-anchored protein-transporting ATPase | 187.2 | 227.27 |
| MRET_1587 | N-terminal acetyltransferase B complex catalytic subunit | 45.5 | 65.48 |
| MRET_1588 | UV radiation resistance-associated protein | 12.38 | 39.26 |

| | | | |
|---|---|---|---|
| MRET_1589 | cell division control protein 45 | 18.25 | 56.65 |
| MRET_1590 | ATP-dependent helicase STH1/SNF2 | 1201.75 | 646.46 |
| MRET_1591 | universal stress protein | 1143.44 | 1287.65 |
| MRET_1592 | large subunit ribosomal protein L3e | 100.16 | 423.54 |
| MRET_1593 | PHD finger and BAH domain protein (Snt2) | 23.83 | 29.85 |
| MRET_1594 | 2OG-Fe(II) oxygenase superfamily | 11.13 | 49.2 |
| MRET_1595 | MFS sugar transporter | 25.09 | 27.59 |
| MRET_1596 | cyclin | 198.48 | 211.61 |
| MRET_1597 | zinc finger protein | 4.62 | 9.15 |
| MRET_1598 | transporter | 52.6 | 89.17 |
| MRET_1599 | splicing factor 3B subunit 4 | 18.4 | 39.13 |
| MRET_1600 | Mn2 homeostasis protein (Per1) | 30.35 | 85.35 |
| MRET_1601 | translation initiation factor 3 subunit J | 43.27 | 120.18 |
| MRET_1602 | isoleucyl-tRNA synthetase | 65.64 | 37.71 |
| MRET_1603 | mRNA (guanine-N7-)-methyltransferase | 15.26 | 25.92 |
| MRET_1604 | N-alpha-acetyltransferase 35, NatC auxiliary subunit | 11.32 | 10.04 |
| MRET_1605 | phosphatidylinositol glycan, class U | 15.87 | 44.28 |
| MRET_1606 | calcium binding protein 39 | 17 | 50.06 |
| MRET_1607 | AP-1 complex subunit gamma-1 | 25.55 | 66.69 |
| MRET_1608 | MFS family protein | 84.83 | 76.03 |
| MRET_1609 | regulator of nonsense transcripts 1 | 22.34 | 43.49 |
| MRET_1610 | molybdopterin binding domain protein | 2524.71 | 1738.06 |
| MRET_1611 | SH3 domain YSC84-like protein 1 | 214.16 | 268.61 |
| MRET_1612 | eukaryotic translation initiation factor 2-alpha kinase 4 | 85.29 | 101.17 |
| MRET_1613 | type IV protein arginine methyltransferase | 56.47 | 108.1 |
| MRET_1614 | nucleoporin p58/p45 | 52.69 | 139.72 |
| MRET_1615 | ATP-binding cassette, subfamily B (MDR/TAP), member 1 | 71.01 | 72.06 |
| MRET_1616 | uncharacterized protein | 35.32 | 54.25 |
| MRET_1617 | uncharacterized protein | 267.14 | 135.59 |
| MRET_1618 | cathepsin D | 9.35 | 24.63 |
| MRET_1619 | uncharacterized protein | 55.3 | 47.79 |
| MRET_1620 | electron transfer flavoprotein alpha subunit | 434.97 | 248.79 |
| MRET_1621 | NAD-dependent histone deacetylase SIR2 | 24.86 | 42.93 |
| MRET_1622 | Rdx family protein | 159.76 | 104.64 |
| MRET_1623 | uncharacterized protein | 563.34 | 350.92 |
| MRET_1624 | transcription initiation factor TFIID complex subunit 8 | 51.74 | 157.32 |
| MRET_1625 | Cdc25 family phosphatase | 80.44 | 102.96 |

| | | | |
|---|---|---|---|
| MRET_1626 | adiponectin receptor | 25.87 | 28.4 |
| MRET_1627 | protein phosphatase PTC2/3 | 61.69 | 250.26 |
| MRET_1628 | translation initiation factor 3 subunit I | 66.18 | 282.94 |
| MRET_1629 | pachytene checkpoint protein 2 | 20.61 | 59.95 |
| MRET_1630 | non-canonical poly(A) RNA polymerase PAPD5/7 | 54.4 | 45.95 |
| MRET_1631 | ATP-dependent RNA helicase DDX46/PRP5 | 119.74 | 73.16 |
| MRET_1632 | uncharacterized protein | 54.96 | 66.29 |
| MRET_1633 | protein KTI12 | 81.58 | 115.19 |
| MRET_1634 | bZIP transcription factor | 468.44 | 327.76 |
| MRET_1635 | protein SSH4 | 132.04 | 132.65 |
| MRET_1636 | serine/threonine-protein kinase 24/25/MST4 | 30.88 | 37.64 |
| MRET_1637 | uncharacterized protein | 118.63 | 81.34 |
| MRET_1638 | scaffold protein involved in the formation of early endocytic sites | 96.26 | 74.84 |
| MRET_1639 | oxidoreductase | 25.88 | 67.42 |
| MRET_1640 | 20S proteasome subunit beta 2 | 69.49 | 181.54 |
| MRET_1641 | SH3-binding, glutamic acid-rich protein | 10.95 | 46.44 |
| MRET_1642 | splicing factor 3B subunit 2 | 49.1 | 86.98 |
| MRET_1643 | translation initiation factor 3 subunit A | 30.31 | 60.03 |
| MRET_1644 | solute carrier family 41 | 15.76 | 33.4 |
| MRET_1645 | mitochondrial chaperone BCS1 | 24.64 | 61.12 |
| MRET_1646 | voltage-dependent calcium channel | 6.31 | 16.64 |
| MRET_1647 | tubulin gamma | 26.83 | 220.85 |
| MRET_1648 | rRNA 2'-O-methyltransferase fibrillarin | 649.92 | 668.67 |
| MRET_1649 | cytochrome-b5 reductase | 664.43 | 546.07 |
| MRET_1650 | predicted ATPase of the ABC class | 73.22 | 74.09 |
| MRET_1651 | chitin synthase | 29.04 | 45.69 |
| MRET_1652 | uncharacterized protein | 1106.44 | 491.19 |
| MRET_1653 | solute carrier family 35, member F5 | 147.53 | 182.37 |
| MRET_1654 | chromatin structure-remodeling complex subunit RSC1/2 | 20.95 | 37.76 |
| MRET_1655 | casein kinase II subunit alpha | 47.72 | 138.32 |
| MRET_1656 | tryptophanyl-tRNA synthetase | 59.38 | 105.86 |
| MRET_1657 | U6 snRNA-associated Sm-like protein LSm2 | 70.57 | 137.04 |
| MRET_1658 | avl9 protein | 19.3 | 49.04 |
| MRET_1659 | uncharacterized protein | 43.32 | 98.79 |
| MRET_1660 | DNA-directed RNA polymerases I, II, and III subunit RPABC1 | 162.05 | 458.79 |
| MRET_1661 | KH domain protein | 27.94 | 33.51 |
| MRET_1662 | RhoGAP | 11.93 | 14.11 |

| | | | |
|---|---|---|---|
| MRET_1663 | protein involved in microtubule-related processes | 50.29 | 73.13 |
| MRET_1664 | DUF6 domain protein | 19.23 | 49.16 |
| MRET_1665 | tRNA (uracil-5-)-methyltransferase | 25.46 | 79.95 |
| MRET_1666 | uncharacterized protein | 152.96 | 179.21 |
| MRET_1667 | uncharacterized protein | 76.43 | 153.87 |
| MRET_1668 | GINS complex subunit 2 | 38.09 | 70.28 |
| MRET_1669 | golgi membrane protein involved in vesicular trafficking and spindle migration | 151.73 | 181.21 |
| MRET_1670 | cytochrome c heme-lyase | 241.08 | 312.21 |
| MRET_1671 | DNA-directed RNA polymerase III subunit RPC3 | 180.56 | 294.38 |
| MRET_1672 | cell wall protein | 17.11 | 30.29 |
| MRET_1673 | heat shock transcription factor | 58.71 | 82.05 |
| MRET_1674 | transcription initiation factor TFIID TATA-box-binding protein | 1420.8 | 1227.88 |
| MRET_1675 | YidC/Oxa1 family membrane protein insertase | 91.88 | 156.76 |
| MRET_1676 | auxiliary protein of DNA polymerase delta | 14.63 | 41.16 |
| MRET_1677 | chromatin structure-remodeling complex subunit RSC9 | 55.28 | 82.36 |
| MRET_1678 | sorting nexin-1/2 | 139.67 | 224.3 |
| MRET_1679 | S-adenosylmethionine synthetase | 234.8 | 207.48 |
| MRET_1680 | uncharacterized protein | 8.17 | 43.22 |
| MRET_1681 | anaphase-promoting complex subunit 2 | 8.78 | 19.11 |
| MRET_1682 | Sec7 domain protein | 52.54 | 106.67 |
| MRET_1683 | protein STE50 | 142.12 | 91.99 |
| MRET_1684 | integral membrane protein | 35.73 | 98.04 |
| MRET_1685 | uncharacterized protein | 59.18 | 91.14 |
| MRET_1686 | mitochondrial intermembrane space import and assembly protein 40 | 2182.72 | 1368.25 |
| MRET_1687 | peroxin-7 | 224 | 229.71 |
| MRET_1688 | MFS phosphate transporter | 60.16 | 71.54 |
| MRET_1689 | cation-transporting P-type ATPase 13A3/4/5 | 38.27 | 60.75 |
| MRET_1690 | uncharacterized protein | 20.64 | 45.88 |
| MRET_1691 | uncharacterized protein | 8.56 | 29.27 |
| MRET_1692 | pericentrin-AKAP-450 domain of centrosomal targeting protein | 3.66 | 16.03 |
| MRET_1693 | uncharacterized protein | 22.17 | 27.3 |
| MRET_1694 | uncharacterized protein | 217.81 | 257.99 |
| MRET_1695 | ATP-dependent DNA helicase 2 subunit 2 | 39.68 | 71.17 |
| MRET_1696 | beta-1,4-mannosyltransferase | 11.86 | 16.62 |
| MRET_1697 | ubiquitin conjugation factor E4 B | 30.7 | 41.61 |
| MRET_1698 | translation initiation factor 2 subunit 1 | 44.72 | 145.31 |
| MRET_1699 | T-complex protein 1 subunit epsilon | 158.43 | 146.88 |

| | | | |
|---|---|---|---|
| MRET_1700 | peptidyl-prolyl isomerase domain and WD repeat protein 1 | 30.08 | 38.82 |
| MRET_1701 | mitochondrial carrier protein | 312.21 | 171.68 |
| MRET_1702 | uncharacterized protein | 16.13 | 18.81 |
| MRET_1703 | uncharacterized protein | 589.65 | 287.21 |
| MRET_1704 | Ca2+:H+ antiporter | 35.44 | 44.42 |
| MRET_1705 | aspartate-semialdehyde dehydrogenase | 77.4 | 126.35 |
| MRET_1706 | dihydroxy-acid dehydratase | 78.58 | 153.21 |
| MRET_1707 | phosphatidylserine decarboxylase | 30 | 49.59 |
| MRET_1708 | TRIAP1/MDM35 family protein | 87.45 | 154.29 |
| MRET_1709 | cytochrome c | 167.02 | 65.18 |
| MRET_1710 | GATA zinc finger | 13.34 | 22.38 |
| MRET_1711 | large subunit ribosomal protein L15e | 69.26 | 367.85 |
| MRET_1712 | large subunit ribosomal protein L5e | 53.7 | 311.17 |
| MRET_1713 | repressible acid phosphatase | 173.72 | 159.26 |
| MRET_1714 | uncharacterized protein | 29.36 | 89.18 |
| MRET_1715 | uncharacterized protein | 61.51 | 165.16 |
| MRET_1716 | NADH dehydrogenase (ubiquinone) 1 alpha subcomplex subunit 4 | 430.12 | 650.35 |
| MRET_1717 | delta14-sterol reductase | 208.51 | 284.39 |
| MRET_1718 | prenylcysteine oxidase/farnesylcysteine lyase | 34.68 | 47.3 |
| MRET_1719 | cofilin | 482.8 | 546.5 |
| MRET_1720 | Ras homolog, member A | 635.27 | 966.17 |
| MRET_1721 | guanosine-diphosphatase | 191.38 | 114.66 |
| MRET_1722 | derlin | 991.65 | 528.33 |
| MRET_1723 | m7GpppX diphosphatase | 20.96 | 52.41 |
| MRET_1724 | N-terminal domain of NEFA-interacting nuclear protein NIP30 | 44.78 | 64.44 |
| MRET_1725 | tubulin-specific chaperone D | 14.67 | 31.85 |
| MRET_1726 | osomolarity two-component system, phosphorelay intermediate protein YPD1 | 145.7 | 205.15 |
| MRET_1727 | ubiquitin carboxyl-terminal hydrolase MINDY-1/2 | 212.01 | 139.58 |
| MRET_1728 | proliferation-associated protein 1 | 90.9 | 113.18 |
| MRET_1729 | map microtubule affinity-regulating kinase | 51.45 | 36.61 |
| MRET_1730 | uncharacterized protein | 1130.38 | 688.4 |
| MRET_1731 | ATP-dependent Clp protease ATP-binding subunit ClpB | 1288.82 | 1368.03 |
| MRET_1732 | protein BCP1 | 23.24 | 42.19 |
| MRET_1733 | carboxy-terminal domain RNA polymerase II polypeptide A small phosphatase | 25.28 | 61.62 |
| MRET_1734 | p-loop containing nucleoside triphosphate hydrolase protein | 35.55 | 80.45 |
| MRET_1735 | mitochondrial intermembrane space protein | 2099.54 | 1621.16 |
| MRET_1736 | ribosome biogenesis protein BMS1 | 190.58 | 172.67 |

| | | | |
|---|---|---|---|
| MRET_1737 | cohesin loading factor subunit SCC2 | 52.85 | 77.18 |
| MRET_1738 | uncharacterized protein | 158.59 | 103.64 |
| MRET_1739 | membrane associated DnaJ chaperone | 11.65 | 46 |
| MRET_1740 | OPT oligopeptide transporter protein | 19.4 | 36.77 |
| MRET_1741 | uncharacterized protein | 19.98 | 58.21 |
| MRET_1742 | phosphatidyl synthase | 115.52 | 195.96 |
| MRET_1743 | integral membrane protein required for ER to golgi transport | 87.73 | 189.44 |
| MRET_1744 | uncharacterized protein | 10.9 | 38.51 |
| MRET_1745 | acyl-coenzyme A thioesterase 13 | 78.06 | 251.05 |
| MRET_1746 | ATP-dependent RNA helicase DHX37/DHR1 | 33 | 73.96 |
| MRET_1747 | V-type H+-transporting ATPase subunit d | 117.77 | 239.58 |
| MRET_1748 | histone deacetylase HOS3 | 352.37 | 343.41 |
| MRET_1749 | RIO kinase 1 | 10.68 | 29.3 |
| MRET_1750 | RhoGAP and Fes CIP4 domain protein | 12.18 | 35.31 |
| MRET_1751 | U2 small nuclear ribonucleoprotein B'' | 573.46 | 553.12 |
| MRET_1752 | small subunit ribosomal protein S19e | 112.13 | 648.62 |
| MRET_1753 | classical protein kinase C alpha type | 232.05 | 425.87 |
| MRET_1754 | uncharacterized protein | 56.7 | 163.63 |
| MRET_1755 | senataxin | 9.89 | 16.82 |
| MRET_1756 | tetratricopeptide repeat domain protein | 18.04 | 28.07 |
| MRET_1757 | DNA damage-binding protein 1 | 37.03 | 32.08 |
| MRET_1758 | uncharacterized protein | 18.78 | 24.77 |
| MRET_1759 | voltage-dependent anion channel protein 2 | 785.42 | 822.62 |
| MRET_1760 | mRNA stabilization protein | 1821.25 | 1545.5 |
| MRET_1761 | subunit 21 of mediator complex | 123.3 | 152.98 |
| MRET_1762 | helix-loop-helix DNA-binding domain protein | 138.7 | 239.65 |
| MRET_1763 | ATP-dependent DNA helicase Q1 | 9.22 | 14.31 |
| MRET_1764 | uncharacterized protein | 56.85 | 47.39 |
| MRET_1765 | N6-L-threonylcarbamoyladenine synthase | 388.71 | 314.72 |
| MRET_1766 | prolyl-tRNA synthetase | 100.52 | 91.58 |
| MRET_1767 | glycoside hydrolase family 16 protein | 206.22 | 120.99 |
| MRET_1768 | conserved hypothetical protein | 29.96 | 30.07 |
| MRET_1769 | NADH dehydrogenase (ubiquinone) 1 alpha subcomplex subunit 9 | 110.26 | 148.58 |
| MRET_1770 | WD repeat protein 22 | 86.21 | 64.96 |
| MRET_1771 | adaptin ear-binding coat-associated protein 1/2 | 257.32 | 174.5 |
| MRET_1772 | uncharacterized protein | 185.92 | 106.55 |
| MRET_1773 | oxidoreductase, short chain dehydrogenase reductase family | 22.67 | 33.41 |

| | | | |
|---|---|---|---|
| MRET_1774 | ribonuclease P/MRP protein subunit POP1 | 17.25 | 33.77 |
| MRET_1775 | conserved hypothetical protein | 149.08 | 61.88 |
| MRET_1776 | serine carboxypeptidase | 149.01 | 82.83 |
| MRET_1777 | uncharacterized protein | 292.48 | 218.75 |
| MRET_1778 | uncharacterized protein | 63.02 | 106.31 |
| MRET_1779 | uncharacterized protein | 37.33 | 34.56 |
| MRET_1780 | uncharacterized protein | 131 | 201.08 |
| MRET_1781 | tyrosyl-DNA phosphodiesterase 1 | 611.78 | 315.76 |
| MRET_1782 | transcription initiation factor TFIIE subunit beta | 92.71 | 83.08 |
| MRET_1783 | mitogen-activated protein kinase | 99.03 | 115.94 |
| MRET_1784 | PHD finger domain protein | 103.01 | 175.57 |
| MRET_1785 | casein kinase II subunit beta | 57.92 | 90.71 |
| MRET_1786 | nucleoporin NUP159 | 28.88 | 60.85 |
| MRET_1787 | tRNA-specific adenosine deaminase 3 | 21.62 | 59.83 |
| MRET_1788 | Na+/H+ antiporter | 44.86 | 132.22 |
| MRET_1789 | uncharacterized protein | 197.59 | 395.52 |
| MRET_1790 | mitochondrial import receptor subunit TOM22 | 151.42 | 320.43 |
| MRET_1791 | enhancer of polycomb-like protein | 855.45 | 547.04 |
| MRET_1792 | rRNA-processing protein | 19.31 | 53.97 |
| MRET_1793 | U3 small nucleolar RNA-associated protein 3 | 14.54 | 35.55 |
| MRET_1794 | actin related protein 2/3 complex, subunit 1A/1B | 1160.36 | 931.02 |
| MRET_1795 | phosphate transporter (Pho88) | 72.03 | 210.38 |
| MRET_1796 | lipid intermediate transporter | 34.19 | 98.17 |
| MRET_1797 | cullin 1 | 103.96 | 173.47 |
| MRET_1798 | elongation factor-2 kinase | 407.09 | 404.55 |
| MRET_1799 | Elongation factor-2 kinase | 714.93 | 779.13 |
| MRET_1800 | U4/U6 snRNA-associated-splicing factor PRP24 | 54.74 | 73.46 |
| MRET_1801 | DUF2373 domain protein | 9.91 | 33.86 |
| MRET_1802 | ATP-dependent RNA helicase DHX33 | 22.12 | 49.95 |
| MRET_1803 | ubiquitin-protein ligase E3 C | 93.34 | 56.42 |
| MRET_1804 | conserved hypothetical protein | 115.86 | 166.53 |
| MRET_1805 | mitochondrial Rho GTPase 1 | 14.5 | 27.98 |
| MRET_1806 | HD family hydrolase | 7.4 | 26.8 |
| MRET_1807 | sentrin-specific protease 7 | 11.21 | 28.66 |
| MRET_1808 | malate dehydrogenase | 400.57 | 376.11 |
| MRET_1809 | malate dehydrogenase | 953.98 | 867.9 |
| MRET_1810 | uncharacterized protein | 53.6 | 60.24 |

| | | | |
|---|---|---|---|
| MRET_1811 | DEAD/DEAH box helicase | 32.46 | 60.26 |
| MRET_1812 | pre-mRNA-processing factor 6 | 18.72 | 26.25 |
| MRET_1813 | DNA binding transcription factor | 78.68 | 101.62 |
| MRET_1814 | protein SCO1/2 | 848.53 | 796.83 |
| MRET_1815 | nulp1-pending protein | 34.83 | 53.92 |
| MRET_1816 | protoporphyrin/coproporphyrin ferrochelatase | 66.88 | 64.29 |
| MRET_1817 | exportin-T | 12 | 31.94 |
| MRET_1818 | uncharacterized protein | 82.31 | 46.61 |
| MRET_1819 | capping protein (actin filament) muscle Z-line, alpha | 175.89 | 79.95 |
| MRET_1820 | derlin | 62.75 | 88.92 |
| MRET_1821 | mitochondrial mRNA processing protein PET127 | 27.61 | 58.85 |
| MRET_1822 | large subunit ribosomal protein L6e | 320.63 | 840.62 |
| MRET_1823 | folylpolyglutamate synthase | 19.36 | 46.8 |
| MRET_1824 | mitochondrial translocator assembly and maintenance protein 41 | 44.47 | 130.82 |
| MRET_1825 | vesicle transport protein | 2412.99 | 2392.86 |
| MRET_1826 | GPI inositol-deacylase | 29.86 | 78.22 |
| MRET_1827 | SH3 domain YSC84-like protein 1 | 1234.93 | 555.07 |
| MRET_1828 | calcineurin-binding protein | 24.18 | 100.91 |
| MRET_1829 | large subunit ribosomal protein L36e | 186.02 | 482.81 |
| MRET_1830 | translation initiation factor 3 subunit E | 56.7 | 110.62 |
| MRET_1831 | NADH dehydrogenase (ubiquinone) 1 alpha subcomplex subunit 7 | 49.97 | 65.74 |
| MRET_1832 | DNA-directed RNA polymerase I subunit RPA49 | 167.09 | 115.92 |
| MRET_1833 | uncharacterized protein | 107.73 | 183.42 |
| MRET_1834 | ADP-ribosylation factor 1 | 1972.67 | 1642.32 |
| MRET_1835 | uncharacterized protein | 21.09 | 30.75 |
| MRET_1836 | uncharacterized protein | 80.33 | 115.18 |
| MRET_1837 | thioredoxin-like protein | 32.51 | 70.81 |
| MRET_1838 | uncharacterized protein | 13.46 | 19.9 |
| MRET_1839 | tRNA pseudouridine55 synthase | 41.06 | 46.99 |
| MRET_1840 | methionyl-tRNA formyltransferase | 96.21 | 71.95 |
| MRET_1841 | DUF500 domain protein | 147.33 | 157.64 |
| MRET_1842 | S-formylglutathione hydrolase | 247.1 | 181.4 |
| MRET_1843 | DnaJ domain protein | 104.41 | 135.88 |
| MRET_1844 | prenyl protein peptidase | 68.5 | 109.5 |
| MRET_1845 | uncharacterized protein | 1362 | 289.19 |
| MRET_1846 | 3'(2'), 5'-bisphosphate nucleotidase | 70.29 | 92.71 |
| MRET_1847 | nicotinate-nucleotide pyrophosphorylase (carboxylating) | 216.36 | 185.52 |

| | | | |
|---|---|---|---|
| MRET_1848 | AAA domain (dynein-related subfamily) | 182.39 | 107.5 |
| MRET_1849 | regulatory associated protein of mTOR | 189 | 83.64 |
| MRET_1850 | importin-9 | 95.23 | 68.71 |
| MRET_1851 | zinc finger protein, C3H1 type | 69.94 | 185.4 |
| MRET_1852 | uncharacterized protein | 52.85 | 162.31 |
| MRET_1853 | BolA-like protein 3 | 75.88 | 84.57 |
| MRET_1854 | cell division cycle 2-like protein | 6.62 | 25.1 |
| MRET_1855 | sterol-4alpha-carboxylate 3-dehydrogenase (decarboxylating) | 109.54 | 125.75 |
| MRET_1856 | zinc finger protein | 1092.6 | 1011.88 |
| MRET_1857 | diazepam-binding inhibitor (GABA receptor modulator, acyl-CoA-binding protein) | 1083.04 | 2122.52 |
| MRET_1858 | oligosaccharyltransferase complex subunit delta (ribophorin II) | 40.7 | 65.24 |
| MRET_1859 | transcriptional activator SPT8 | 109.25 | 82.96 |
| MRET_1860 | tRNAThr (cytosine32-N3)-methyltransferase | 56.57 | 44.23 |
| MRET_1861 | threonine aldolase | 240.19 | 95.53 |
| MRET_1862 | 26S proteasome regulatory subunit N2 | 80.12 | 91.24 |
| MRET_1863 | prefoldin subunit 2 | 62.38 | 141 |
| MRET_1864 | NADH-ubiquinone oxidoreductase 12 kda subunit | 155.36 | 348.73 |
| MRET_1865 | tRNA (guanine-N7-)-methyltransferase | 33.79 | 50.77 |
| MRET_1866 | translation initiation factor 3 subunit F | 221.26 | 276.93 |
| MRET_1867 | mitochondrial import receptor subunit TOM7 | 211.4 | 270.81 |
| MRET_1868 | N-alpha-acetyltransferase 40 | 36.59 | 43.9 |
| MRET_1869 | calcium/calmodulin-dependent protein kinase kinase 2 | 88.5 | 74.13 |
| MRET_1870 | putative transcription factor | 1208.63 | 774.1 |
| MRET_1871 | uncharacterized protein | 22.14 | 35.17 |
| MRET_1872 | cell growth-regulating nucleolar protein | 24.14 | 41.31 |
| MRET_1873 | G1 S-specific cyclin | 27.34 | 30.34 |
| MRET_1874 | large subunit ribosomal protein L30e | 1154.35 | 1174.89 |
| MRET_1875 | large subunit ribosomal protein L37e | 545.98 | 1400.66 |
| MRET_1876 | large subunit ribosomal protein L9e | 232.14 | 831.28 |
| MRET_1877 | pre-mRNA cleavage complex 2 protein Pcf11 | 166.96 | 197 |
| MRET_1878 | conserved oligomeric golgi complex subunit 3 | 30.33 | 45.63 |
| MRET_1879 | NADPH2:quinone reductase | 47.95 | 69.4 |
| MRET_1880 | aspartate kinase | 125.48 | 120.33 |
| MRET_1881 | exportin-2 (importin alpha re-exporter) | 163.25 | 389.81 |
| MRET_1882 | protein SSD1 | 63 | 54.86 |
| MRET_1883 | ATP-dependent RNA helicase DBP3 | 56.14 | 103.09 |
| MRET_1884 | WD repeat protein JIP5 | 24.81 | 40.16 |

| | | | |
|---|---|---|---|
| MRET_1885 | branched-chain amino acid aminotransferase | 148.03 | 106.06 |
| MRET_1886 | alpha/beta-hydrolase | 3.94 | 5.19 |
| MRET_1887 | small subunit ribosomal protein S29e | 155.46 | 660.89 |
| MRET_1888 | N-terminal acetyltransferase 2 | 41.59 | 154.59 |
| MRET_1889 | ATP-dependent bile acid permease | 78.16 | 89.82 |
| MRET_1890 | DNA mismatch repair protein | 16.61 | 38.56 |
| MRET_1891 | proteasome assembly chaperone 2 | 13.34 | 37.41 |
| MRET_1892 | ATP-dependent bile acid permease | 116.61 | 115.96 |
| MRET_1893 | CCR4-NOT transcription complex subunit 1 | 61.43 | 62.94 |
| MRET_1894 | protein involved in GPI anchor synthesis | 64.74 | 71.45 |
| MRET_1895 | calcium/calmodulin-dependent protein kinase I | 125.43 | 206.63 |
| MRET_1896 | histone-lysine N-methyltransferase, H3 lysine-79 specific | 53.42 | 104.02 |
| MRET_1897 | methylated-DNA-protein-cysteine methyltransferase related protein | 58.45 | 76.55 |
| MRET_1898 | uncharacterized protein | 695 | 460.33 |
| MRET_1899 | phenylalanyl-tRNA synthetase alpha chain | 68.82 | 76.55 |
| MRET_1900 | abhydrolase domain protein 12 | 83.77 | 161.85 |
| MRET_1901 | dehydrodolichyl diphosphate syntase complex subunit NUS1 | 10.7 | 19.82 |
| MRET_1902 | uncharacterized protein | 19.32 | 35.95 |
| MRET_1903 | GTP cyclohydrolase II | 51.31 | 43.47 |
| MRET_1904 | protein AATF/BFR2 | 101.99 | 143.54 |
| MRET_1905 | U4/U6 small nuclear ribonucleoprotein PRP3 | 123.88 | 121.71 |
| MRET_1906 | importin subunit alpha | 33.75 | 79.12 |
| MRET_1907 | ankyrin repeat domain protein | 11.68 | 20.77 |
| MRET_1908 | lipid-binding protein | 36.69 | 56.15 |
| MRET_1909 | template-activating factor I | 109.31 | 324.68 |
| MRET_1910 | uncharacterized protein | 433.02 | 796.67 |
| MRET_1911 | E3 ubiquitin-protein ligase HECTD2 | 277.07 | 383.53 |
| MRET_1912 | essential RNA-binding component of cleavage and polyadenylation factor | 62.47 | 50.3 |
| MRET_1913 | thioesterase | 116.26 | 245.13 |
| MRET_1914 | anaphase-promoting complex subunit 5 | 82.61 | 109.43 |
| MRET_1915 | phosphatase with a broad substrate specificity | 46.68 | 44.36 |
| MRET_1916 | uncharacterized protein | 472.33 | 230.57 |
| MRET_1917 | NADH-ubiquinone oxidoreductase | 713.04 | 1182.24 |
| MRET_1918 | cation efflux family protein | 44.14 | 117.42 |
| MRET_1919 | essential protein that forms a complex with Rli1p and Yae1p | 211.68 | 313.59 |
| MRET_1920 | uncharacterized protein | 36.29 | 100.56 |
| MRET_1921 | HUS1 checkpoint protein | 50.59 | 183.44 |

| | | | |
|---|---|---|---|
| MRET_1922 | oxidative stress survival svf1-like protein | 62.73 | 157.52 |
| MRET_1923 | protein transport protein SEC61 subunit beta | 151.54 | 272.61 |
| MRET_1924 | uncharacterized protein | 127.67 | 621.46 |
| MRET_1925 | acetyltransferase (GNAT) family | 17.11 | 99.61 |
| MRET_1926 | PPR repeat containing protein | 34.65 | 71.21 |
| MRET_1927 | uncharacterized protein | 361.25 | 195.13 |
| MRET_1928 | uncharacterized protein | 6.65 | 13.56 |
| MRET_1929 | phosphomannomutase | 57.68 | 94.17 |
| MRET_1930 | protein CWC21 | 297.31 | 309.76 |
| MRET_1931 | CCR4-NOT transcription complex subunit 9 | 31.81 | 51.72 |
| MRET_1932 | double-strand break repair protein MRE11 | 71.7 | 75.28 |
| MRET_1933 | phosphatidylglycerol phosphatidylinositol transfer protein | 22.69 | 36.4 |
| MRET_1934 | RIO kinase 2 | 83.38 | 111.92 |
| MRET_1935 | palmitoyltransferase ZDHHC9/14/18 | 82.45 | 145.53 |
| MRET_1936 | sister chromatid separation protein | 342.68 | 250.61 |
| MRET_1937 | uncharacterized protein | 49.51 | 64.72 |
| MRET_1938 | 20S proteasome subunit alpha 7 | 233 | 257.77 |
| MRET_1939 | U1 small nuclear ribonucleoprotein 70kDa | 65.29 | 77.6 |
| MRET_1940 | dolichol kinase | 85.19 | 74.83 |
| MRET_1941 | ESCRT-I complex subunit VPS28 | 72.7 | 77.24 |
| MRET_1942 | uncharacterized protein | 185.92 | 75.63 |
| MRET_1943 | tRNA-dihydrouridine synthase 1 | 108.3 | 74.32 |
| MRET_1944 | kinetochore protein Spc7/SPC105 | 24.47 | 50.79 |
| MRET_1945 | ATP-dependent RNA helicase DDX52/ROK1 | 98.17 | 70.76 |
| MRET_1946 | nucleolar MIF4G domain protein 1 | 46.07 | 51.06 |
| MRET_1947 | minor histocompatibility antigen H13 | 70.43 | 122.81 |
| MRET_1948 | ribulose-phosphate 3-epimerase | 463.64 | 417.47 |
| MRET_1949 | pre-mRNA-processing factor 17 | 251.73 | 176.01 |
| MRET_1950 | regulator of ribosome biosynthesis | 17.93 | 60.09 |
| MRET_1951 | interactor of little elongation complex ELL subunit 2 | 21.91 | 46.38 |
| MRET_1952 | ADP-ribosylation factor related protein 1 | 180.07 | 177.19 |
| MRET_1953 | NADH dehydrogenase (ubiquinone) Fe-S protein 1 | 296.57 | 121.47 |
| MRET_1954 | integral membrane protein | 50.82 | 79.26 |
| MRET_1955 | uridine kinase | 145.22 | 142.56 |
| MRET_1956 | serine/threonine-protein phosphatase PP1 catalytic subunit | 112.09 | 214.58 |
| MRET_1957 | CCR4-NOT complex subunit CAF16 | 32.41 | 45.76 |
| MRET_1958 | U3 small nucleolar RNA-associated protein 7 | 54.27 | 158.55 |

| | | | |
|---|---|---|---|
| MRET_1959 | glycosyl transferases group 1 | 1057.11 | 847.2 |
| MRET_1960 | WD domain, G-beta repeat protein | 112.72 | 72.32 |
| MRET_1961 | RNA-binding protein 39 | 39.29 | 41.2 |
| MRET_1962 | general transcription factor IIIA | 12.26 | 24.64 |
| MRET_1963 | guanyl-nucleotide exchange factor | 22.17 | 95.74 |
| MRET_1964 | uncharacterized protein | 1901.71 | 2027.31 |
| MRET_1965 | tubulin-tyrosine ligase family protein | 41.9 | 62.53 |
| MRET_1966 | uncharacterized protein | 33.49 | 85.52 |
| MRET_1967 | nucleolar protein 9 | 52.18 | 51.31 |
| MRET_1968 | argininosuccinate synthase | 156.19 | 191.06 |
| MRET_1969 | BolA-like protein 1 | 406.97 | 254.57 |
| MRET_1970 | trehalose 6-phosphate synthase | 39.94 | 106.28 |
| MRET_1971 | universal stress protein | 90.18 | 122.74 |
| MRET_1972 | uncharacterized protein | 34.14 | 17.7 |
| MRET_1973 | 17beta-estradiol 17-dehydrogenase | 71.99 | 86.01 |
| MRET_1974 | NAD+ synthase (glutamine-hydrolysing) | 32.72 | 35.46 |
| MRET_1975 | uncharacterized protein | 31.37 | 21.07 |
| MRET_1976 | uncharacterized protein | 47 | 35.25 |
| MRET_1977 | conserved hypothetical protein | 49.41 | 88.24 |
| MRET_1978 | cell polarity protein | 52.26 | 64.97 |
| MRET_1979 | replication factor C subunit 2/4 | 30.16 | 67.12 |
| MRET_1980 | uncharacterized protein | 43.33 | 43.78 |
| MRET_1981 | adiponectin receptor | 311.53 | 181.12 |
| MRET_1982 | pyridoxamine 5'-phosphate oxidase | 33.95 | 33.69 |
| MRET_1983 | mitochondrial protein | 105.34 | 159.96 |
| MRET_1984 | Ras-related GTP-binding protein A/B | 50.86 | 95.92 |
| MRET_1985 | uncharacterized protein | 38.24 | 60.6 |
| MRET_1986 | sphingolipid C9-methyltransferase | 56.61 | 62.27 |
| MRET_1987 | uncharacterized protein | 132.38 | 100.32 |
| MRET_1988 | peptidyl-prolyl cis-trans isomerase NIMA-interacting 1 | 113.12 | 165.08 |
| MRET_1989 | tRNA A64-2'-O-ribosylphosphate transferase | 13.69 | 22.88 |
| MRET_1990 | glycosyl hydrolase family 88 | 592.91 | 308.77 |
| MRET_1991 | isocitrate dehydrogenase IDP1 | 1676.11 | 1395.44 |
| MRET_1992 | mitochondrial ornithine carrier protein | 402.18 | 432.57 |
| MRET_1993 | conserved hypothetical protein | 161.77 | 117.74 |
| MRET_1994 | acid phosphatase | 1125.79 | 757.06 |
| MRET_1995 | ditrans,polycis-polyprenyl diphosphate synthase | 131.71 | 384.8 |

| | | | |
|---|---|---|---|
| MRET_1996 | uncharacterized protein | 30.86 | 40.91 |
| MRET_1997 | coenzyme Q-binding protein COQ10 | 20.97 | 44.74 |
| MRET_1998 | NADPH-dependent medium chain alcohol dehydrogenase | 44.73 | 125.5 |
| MRET_1999 | ubiquinol-cytochrome-c reductase complex subunit (QCR10) | 154.74 | 215.53 |
| MRET_2000 | carbonic anhydrase | 88.56 | 102.07 |
| MRET_2001 | uracil-DNA glycosylase | 87.6 | 112.96 |
| MRET_2002 | ubiquitin carboxyl-terminal hydrolase L3 | 107.94 | 59.55 |
| MRET_2003 | uncharacterized protein | 52.75 | 46.51 |
| MRET_2004 | PUA domain protein | 158.23 | 95.64 |
| MRET_2005 | prolyl oligopeptidase | 41.07 | 43.49 |
| MRET_2006 | large subunit GTPase 1 | 90.98 | 77.27 |
| MRET_2007 | U3 small nucleolar RNA-associated protein 25 | 19.54 | 25.55 |
| MRET_2008 | uncharacterized protein | 33.58 | 51.84 |
| MRET_2009 | SHO1 osmosensor | 974.17 | 1257.57 |
| MRET_2010 | cytochrome c oxidase subunit 5b | 431.9 | 655.05 |
| MRET_2011 | cytochrome b561 | 36.88 | 52.69 |
| MRET_2012 | large subunit ribosomal protein L49 | 41.05 | 112.1 |
| MRET_2013 | protein transport protein SEC61 subunit gamma and related proteins | 130.93 | 366.73 |
| MRET_2014 | large subunit ribosomal protein L35e | 171.78 | 682.47 |
| MRET_2015 | uncharacterized protein | 48.1 | 58.89 |
| MRET_2016 | Ras-related C3 botulinum toxin substrate 1 | 218.2 | 522.62 |
| MRET_2017 | uncharacterized protein | 39.15 | 62.92 |
| MRET_2018 | uncharacterized protein | 29.21 | 55.46 |
| MRET_2019 | nuclear protein localization protein 4 homolog | 423.39 | 300.81 |
| MRET_2020 | cardiolipin-specific phospholipase | 69.85 | 92.05 |
| MRET_2021 | recombination hotspot-binding protein | 39.12 | 476.12 |
| MRET_2022 | CCAAT-binding factor complex subunit | 98.84 | 186.93 |
| MRET_2023 | upstream activation factor subunit UAF30 | 166.8 | 351.19 |
| MRET_2024 | mitochondrial protein involved in assembly of succinate dehydrogenase | 122.65 | 138.73 |
| MRET_2025 | conserved hypothetical protein | 83.28 | 123.03 |
| MRET_2026 | prohibitin 1 | 1427.76 | 663.73 |
| MRET_2027 | uncharacterized protein | 1386.33 | 301.62 |
| MRET_2028 | uncharacterized protein | 248.11 | 160.55 |
| MRET_2029 | D-glycerate 3-kinase | 329.73 | 238.24 |
| MRET_2030 | ssDNA-binding protein essential for mitochondrial genome maintenance | 110.26 | 211.7 |
| MRET_2031 | sucrase/ferredoxin-like protein | 201.23 | 81.6 |
| MRET_2032 | 2-methoxy-6-polyprenyl-1,4-benzoquinol methylase | 67.63 | 106.45 |

| | | | |
|---|---|---|---|
| MRET_2033 | uncharacterized protein | 218.08 | 304.7 |
| MRET_2034 | inositol polyphosphate 5-phosphatase | 165.04 | 158.35 |
| MRET_2035 | uncharacterized protein | 310.22 | 227.54 |
| MRET_2036 | mitochondrial fission process protein 1 | 115.59 | 87.73 |
| MRET_2037 | thioredoxin | 13276.84 | 8853.33 |
| MRET_2038 | monooxygenase | 507.76 | 227.01 |
| MRET_2039 | cellular morphogenesis protein | 157.75 | 179.18 |
| MRET_2040 | Pescadillo homolog | 110.71 | 237.66 |
| MRET_2041 | small nuclear ribonucleoprotein | 314.17 | 232.29 |
| MRET_2042 | fumarate hydratase, class II | 634.43 | 262.16 |
| MRET_2043 | conserved hypothetical protein | 184.3 | 64.73 |
| MRET_2044 | 37S ribosomal protein RSM22 | 119.8 | 42.82 |
| MRET_2045 | protein DGCR14 | 12.06 | 14.26 |
| MRET_2046 | guanine nucleotide-binding protein G(I)/G(S)/G(T) subunit beta-1 | 36.23 | 48.33 |
| MRET_2047 | solute carrier family 24 (sodium potassium calcium exchanger), member 1 | 145.75 | 176.99 |
| MRET_2048 | thioesterase | 878.88 | 183.68 |
| MRET_2049 | ribosome assembly protein RRB1 | 108.46 | 205.97 |
| MRET_2050 | phosphoribosylamine--glycine ligase/phosphoribosylformylglycinamidine cyclo-ligase | 371.5 | 246.11 |
| MRET_2051 | transporter | 74.28 | 64.88 |
| MRET_2052 | uncharacterized protein | 40.99 | 38.88 |
| MRET_2053 | serine/threonine-protein kinase haspin | 44.71 | 54.98 |
| MRET_2054 | large subunit ribosomal protein L1 | 109.67 | 98.91 |
| MRET_2055 | SUR7/PalI family protein | 96.88 | 72.56 |
| MRET_2056 | uncharacterized protein | 44.73 | 37.16 |
| MRET_2057 | component of the EKC/KEOPS complex | 39.74 | 39.78 |
| MRET_2058 | adenine phosphoribosyltransferase | 50.99 | 84.87 |
| MRET_2059 | uncharacterized protein | 28.02 | 33.07 |
| MRET_2060 | uncharacterized protein | 33.84 | 48.95 |
| MRET_2061 | citrate lyase subunit beta-like protein | 254 | 143.44 |
| MRET_2062 | ATP-binding cassette, subfamily B (MDR/TAP), member 6 | 153.43 | 58.25 |
| MRET_2063 | uncharacterized protein | 197.48 | 283.64 |
| MRET_2064 | ER membrane protein SH3 | 822.13 | 614.89 |
| MRET_2065 | mitochondrial pyruvate dehydrogenase kinase | 77.47 | 102.99 |
| MRET_2066 | vesicle transport through interaction with t-SNAREs 1 | 1784.08 | 1122.44 |
| MRET_2067 | sensor protein CreC | 54.56 | 137.77 |
| MRET_2068 | IGR motif protein | 32.33 | 91.77 |
| MRET_2069 | uncharacterized protein | 22.26 | 70.2 |

| | | | |
|---|---|---|---|
| MRET_2070 | platelet-activating factor acetylhydrolase | 15.86 | 45.44 |
| MRET_2071 | ubiquitin-conjugating enzyme (huntingtin interacting protein 2) | 75.94 | 275.46 |
| MRET_2072 | oxalate---CoA ligase | 160.96 | 177.51 |
| MRET_2073 | ATP-dependent NAD(P)H-hydrate dehydratase | 74.18 | 91.58 |
| MRET_2074 | uncharacterized protein | 39.54 | 116.54 |
| MRET_2075 | uncharacterized protein | 130.56 | 168.69 |
| MRET_2076 | cytidine deaminase | 110.45 | 143.68 |
| MRET_2077 | 5-oxoprolinase (ATP-hydrolysing) | 246.73 | 152.85 |
| MRET_2078 | cytochrome c oxidase subunit 23 | 328.32 | 165.25 |
| MRET_2079 | WD repeat protein 61 | 179.91 | 134.14 |
| MRET_2080 | uncharacterized protein | 77.46 | 50.72 |
| MRET_2081 | solute carrier family 25 (mitochondrial carnitine/acylcarnitine transporter), member 20/29 | 167.66 | 91.61 |
| MRET_2082 | coatomer subunit epsilon | 53.95 | 74.11 |
| MRET_2083 | cytochrome c peroxidase | 725.57 | 282.01 |
| MRET_2084 | uncharacterized protein | 86.63 | 68.82 |
| MRET_2085 | Rab GDP dissociation inhibitor | 577.16 | 461.74 |
| MRET_2086 | adenylate kinase | 391.44 | 326.82 |
| MRET_2087 | large subunit ribosomal protein L22 | 71.84 | 109.11 |
| MRET_2088 | aldehyde dehydrogenase (NAD+) | 96.73 | 116.46 |
| MRET_2089 | peptidyl-prolyl cis-trans isomerase B (cyclophilin B) | 269.77 | 349.71 |
| MRET_2090 | RhoGAP | 15.13 | 31.48 |
| MRET_2091 | subunit of the heterohexameric cochaperone prefoldin complex | 52.95 | 113.14 |
| MRET_2092 | peptidyl-prolyl isomerase D | 129.96 | 131.21 |
| MRET_2093 | alanine-glyoxylate transaminase/serine-glyoxylate transaminase/serine-pyruvate transaminase | 221.97 | 156.8 |
| MRET_2094 | putative methyltransferase | 146.94 | 67.28 |
| MRET_2095 | actin related protein 2/3 complex, subunit 2 | 203.6 | 181.85 |
| MRET_2096 | ATP-binding protein involved in chromosome partitioning | 221.45 | 127.96 |
| MRET_2097 | succinate dehydrogenase (ubiquinone) flavoprotein subunit | 191.34 | 87.18 |
| MRET_2098 | altered inheritance of mitochondria protein 13 | 143.84 | 252.14 |
| MRET_2099 | zinc finger protein, C2HC5-type | 34.03 | 61.72 |
| MRET_2100 | F-box and leucine-rich repeat protein 10/11 | 78.97 | 62.23 |
| MRET_2101 | THO complex subunit 1 | 63.12 | 86.6 |
| MRET_2102 | RNA-binding protein 8A | 104.39 | 279.98 |
| MRET_2103 | small subunit ribosomal protein S27Ae | 4467.41 | 5795.3 |
| MRET_2104 | dual specificity kinase | 224.64 | 149.69 |
| MRET_2105 | DUF1713 domain protein | 230.68 | 153.07 |
| MRET_2106 | mitochondrial 54S ribosomal protein RML2 | 38.02 | 86.28 |

| | | | |
|---|---|---|---|
| MRET_2107 | S-phase kinase-associated protein 1 | 623.24 | 706.14 |
| MRET_2108 | SNF2 chromatin remodeling protein | 220.74 | 100.33 |
| MRET_2109 | NADH dehydrogenase (ubiquinone) 1 alpha subcomplex subunit 1 | 233.02 | 172.97 |
| MRET_2110 | mitochondrial import inner membrane translocase subunit TIM44 | 378.56 | 342.93 |
| MRET_2111 | CCCH zinc finger and SMR | 40.42 | 43.98 |
| MRET_2112 | RNA polymerase II-associated factor 1 | 84.32 | 121.45 |
| MRET_2113 | PAB-dependent poly(A)-specific ribonuclease subunit 2 | 81.83 | 61.21 |
| MRET_2114 | uncharacterized protein | 137.71 | 131.38 |
| MRET_2115 | SRP40, C-terminal domain protein | 21.67 | 36.65 |
| MRET_2116 | sugar transporter | 402.17 | 294.19 |
| MRET_2117 | proteasome activator subunit 4 | 48.39 | 39.54 |
| MRET_2118 | YdiU domain protein | 388.45 | 138.84 |
| MRET_2119 | phosphatidylinositol glycan, class M | 11.44 | 33.08 |
| MRET_2120 | magnesium transporter | 17.5 | 16.36 |
| MRET_2121 | SUR7/PalI family protein | 276.23 | 129.96 |
| MRET_2122 | uncharacterized protein | 99.72 | 68.07 |
| MRET_2123 | uncharacterized protein | 24825.52 | 20449.66 |
| MRET_2124 | acyl-CoA dehydrogenase | 460.21 | 397.13 |
| MRET_2125 | CBS PB1 domain protein | 52.19 | 75.57 |
| MRET_2126 | uncharacterized protein | 91.13 | 89.4 |
| MRET_2127 | phospholipid:diacylglycerol acyltransferase | 91.59 | 58.99 |
| MRET_2128 | diphthamide biosynthesis protein 4 | 36.2 | 44.75 |
| MRET_2129 | sentrin-specific protease 1 | 32.68 | 82.93 |
| MRET_2130 | regulatory factor Sgt1 | 51.36 | 96.97 |
| MRET_2131 | NADH dehydrogenase (ubiquinone) Fe-S protein 3 | 161.73 | 224.54 |
| MRET_2132 | pre-mRNA-splicing factor SPF27 | 110.7 | 165.78 |
| MRET_2133 | rRNA processing protein RRP15 | 82.59 | 176.19 |
| MRET_2134 | succinyl-CoA synthetase beta subunit | 289.66 | 367.09 |
| MRET_2135 | mitochondrial 37S ribosomal protein MRPS8 | 66.52 | 119.07 |
| MRET_2136 | S-adenosylmethionine-dependent methyltransferase | 77.79 | 174.59 |
| MRET_2137 | small subunit ribosomal protein S27e | 449.38 | 944.15 |
| MRET_2138 | large subunit ribosomal protein L19e | 298.07 | 906.54 |
| MRET_2139 | protein of unknown function (DUF2413) | 31.16 | 120.53 |
| MRET_2140 | uncharacterized protein | 48.79 | 369.47 |
| MRET_2141 | serine/threonine-protein phosphatase 2B catalytic subunit | 71.46 | 91.4 |
| MRET_2142 | 20S proteasome subunit beta 6 | 124.9 | 174.2 |
| MRET_2143 | Dsk2-ubiquitin-like protein | 199.44 | 204.79 |

| | | | |
|---|---|---|---|
| MRET_2144 | glycerol-3-phosphate phosphatase | 222.55 | 120.6 |
| MRET_2145 | glycerol-3-phosphate phosphatase | 27.7 | 72.37 |
| MRET_2146 | SNF1 kinase complex beta-subunit Gal83 | 32.6 | 60.21 |
| MRET_2147 | anaphase-promoting complex subunit 11 | 20.88 | 33.53 |
| MRET_2148 | kinesin family member C1 | 36.21 | 83.93 |
| MRET_2149 | nuclear segregation protein | 64.05 | 205.88 |
| MRET_2150 | CTD kinase subunit alpha | 68.2 | 77.35 |
| MRET_2151 | fungal Zn(2)-Cys(6) binuclear cluster domain protein | 875.7 | 452.31 |
| MRET_2152 | xeroderma pigmentosum group C-complementing protein | 22.92 | 29.92 |
| MRET_2153 | carnosine N-methyltransferase | 15.3 | 39.2 |
| MRET_2154 | cytomegalovirus gH-receptor family protein | 35.95 | 66.27 |
| MRET_2155 | uncharacterized protein | 66.64 | 70.88 |
| MRET_2156 | rRNA-processing protein EBP2 | 569.35 | 2751.51 |
| MRET_2157 | RNA polymerase-associated protein LEO1 | 46.81 | 67.08 |
| MRET_2158 | inverted formin | 254.64 | 297.81 |
| MRET_2159 | zinc finger protein | 76.69 | 50.45 |
| MRET_2160 | glucose-6-phosphate 1-dehydrogenase | 1176.04 | 852.85 |
| MRET_2161 | superoxide dismutase, Fe-Mn family | 967.52 | 989.01 |
| MRET_2162 | zinc finger protein (RING finger) | 36.84 | 100.17 |
| MRET_2163 | ATP-dependent RNA helicase UAP56/SUB2 | 133.72 | 233.55 |
| MRET_2164 | methyltransferase-like protein 13 | 28.83 | 36.13 |
| MRET_2165 | pre-mRNA-splicing factor spp2 | 32.04 | 53.96 |
| MRET_2166 | saccharopepsin | 1269.73 | 878.88 |
| MRET_2167 | uncharacterized protein | 523.07 | 548.62 |
| MRET_2168 | dynein light chain roadblock-type | 50.54 | 83.23 |
| MRET_2169 | mitotic spindle assembly checkpoint protein MAD1 | 136.61 | 110.87 |
| MRET_2170 | bZIP transcription factor | 1022.99 | 488.75 |
| MRET_2171 | tubulin-specific chaperone A | 100.35 | 55.5 |
| MRET_2172 | ribonuclease Z | 63.96 | 74.09 |
| MRET_2173 | serine/threonine-protein kinase PRP4 | 214.49 | 263.41 |
| MRET_2174 | ATP-dependent RNA helicase DDX5/DBP2 | 118.54 | 237.25 |
| MRET_2175 | DNA-(apurinic or apyrimidinic site) lyase | 40.16 | 36.38 |
| MRET_2176 | N6-L-threonylcarbamoyladenine synthase | 126.47 | 81.14 |
| MRET_2177 | uncharacterized protein | 176.89 | 160.42 |
| MRET_2178 | ribosomal protein L30p/L7e | 562.1 | 1519.89 |
| MRET_2179 | 4-hydroxybenzoate polyprenyltransferase, mitochondrial | 707.89 | 437.85 |
| MRET_2180 | 4-hydroxybenzoate polyprenyltransferase | 61.17 | 97.91 |

| | | | |
|---|---|---|---|
| MRET_2181 | uncharacterized protein | 236.76 | 265.11 |
| MRET_2182 | uncharacterized protein | 37.72 | 63.96 |
| MRET_2183 | uncharacterized protein | 38.54 | 50.8 |
| MRET_2184 | chitin deacetylase | 230.83 | 317.22 |
| MRET_2185 | ribosome biogenesis protein YTM1 | 105.51 | 104.26 |
| MRET_2186 | endothelin-converting enzyme | 290.08 | 146.16 |
| MRET_2187 | endothelin-converting enzyme | 43.64 | 76.19 |
| MRET_2188 | YagE family protein | 148.01 | 90.25 |
| MRET_2189 | coiled-coil domain protein | 123.54 | 82.63 |
| MRET_2190 | 4'-phosphopantetheinyl transferase | 68.28 | 46.29 |
| MRET_2191 | vacuolar protein sorting 55 superfamily | 55.22 | 97.45 |
| MRET_2192 | CCR4-NOT transcription complex subunit 4 | 407.34 | 460.17 |
| MRET_2193 | trafficking protein particle complex subunit 9 | 74.69 | 32.72 |
| MRET_2194 | uncharacterized protein | 111.75 | 172 |
| MRET_2195 | cysteine synthase A | 40.98 | 82.34 |
| MRET_2196 | helix-loop-helix DNA-binding domain protein | 42.62 | 96.84 |
| MRET_2197 | beta-glucan synthesis-associated protein KRE6 | 24.37 | 67 |
| MRET_2198 | AP complex subunit beta | 40.52 | 37.42 |
| MRET_2199 | U3 small nucleolar RNA-associated protein 15 | 68.22 | 101.17 |
| MRET_2200 | copper chaperone | 2525.18 | 1290.35 |
| MRET_2201 | small subunit ribosomal protein S21e | 108.98 | 399.62 |
| MRET_2202 | pumilio-family RNA binding repeat protein | 43.03 | 47.9 |
| MRET_2203 | TBC1 domain family member 15 | 28.1 | 25.8 |
| MRET_2204 | uncharacterized protein | 18.08 | 21.89 |
| MRET_2205 | 3-oxoacid CoA-transferase | 1014.8 | 652.94 |
| MRET_2206 | metal transporter CNNM | 410.62 | 148.97 |
| MRET_2207 | DnaJ homolog subfamily A member 2 | 486.51 | 535.44 |
| MRET_2208 | uncharacterized protein | 27.49 | 93.79 |
| MRET_2209 | mitochondrial 54S ribosomal protein YmL38 YmL34 | 30.39 | 77.03 |
| MRET_2210 | tRNA (guanine26-N2/guanine27-N2)-dimethyltransferase | 35.13 | 52.63 |
| MRET_2211 | L-ascorbic acid binding protein | 400.66 | 384.69 |
| MRET_2212 | peroxin-1 | 17.71 | 22.99 |
| MRET_2213 | mannosyl phosphorylinositol ceramide synthase SUR1 | 21.72 | 33 |
| MRET_2214 | metal resistance protein YCF1 | 22.58 | 30.83 |
| MRET_2215 | homoaconitase | 118.37 | 90.43 |
| MRET_2216 | suppressor of G2 allele of SKP1 | 95.79 | 83.71 |
| MRET_2217 | alpha-1,3-glucosyltransferase | 49.24 | 38.36 |

| | | | |
|---|---|---|---|
| MRET_2218 | Ras suppressor protein 1 | 66.06 | 59.25 |
| MRET_2219 | orotidine 5-phosphate decarboxylase | 91.05 | 98.72 |
| MRET_2220 | kinesin family member 5 | 38.1 | 73.24 |
| MRET_2221 | chorismate mutase | 17.94 | 11.71 |
| MRET_2222 | E3 ubiquitin-protein ligase listerin | 100.37 | 35.73 |
| MRET_2223 | DUF2346 domain protein | 206.23 | 267.85 |
| MRET_2224 | INO80 complex subunit C | 82.64 | 187.17 |
| MRET_2225 | endonuclease/exonuclease/phosphatase family | 465.51 | 425.28 |
| MRET_2226 | uncharacterized protein | 494.48 | 423.54 |
| MRET_2227 | uncharacterized protein | 632.99 | 437.95 |
| MRET_2228 | uncharacterized protein | 34.34 | 58.63 |
| MRET_2229 | uncharacterized protein | 12.91 | 18.58 |
| MRET_2230 | pyruvate decarboxylase | 366.35 | 179.11 |
| MRET_2231 | protein AIR1/2 | 32.15 | 57.01 |
| MRET_2232 | cell division control protein 6 | 15.25 | 27.31 |
| MRET_2233 | uncharacterized protein | 36.16 | 56.77 |
| MRET_2234 | mitochondrial division protein 1 | 23.28 | 49.89 |
| MRET_2235 | GET complex subunit GET2 | 115.97 | 130.87 |
| MRET_2236 | DNA topoisomerase II | 40.57 | 46.52 |
| MRET_2237 | 3-hydroxyisobutyryl-CoA hydrolase | 264.63 | 230.46 |
| MRET_2238 | uncharacterized protein | 63.34 | 73.74 |
| MRET_2239 | phosphatidylinositol glycan, class O | 138.79 | 94.89 |
| MRET_2240 | COP9 signalosome complex subunit 12 | 39.48 | 58.69 |
| MRET_2241 | DNA polymerase delta subunit 2 | 21.73 | 26.6 |
| MRET_2242 | tRNA-dihydrouridine synthase 3 | 75.99 | 68.1 |
| MRET_2243 | methionyl-tRNA synthetase | 90.84 | 155.81 |
| MRET_2244 | protein phosphatase inhibitor 2 (IPP-2) | 22.01 | 26.25 |
| MRET_2245 | lysophospholipid hydrolase | 19.86 | 24.12 |
| MRET_2246 | glutamine synthetase | 1161.99 | 1498.09 |
| MRET_2247 | protein phosphatase 4 regulatory subunit 3 | 126.89 | 65.03 |
| MRET_2248 | YEATS domain protein 4 | 31.57 | 46.83 |
| MRET_2249 | GDP-mannose transporter | 22.06 | 39.22 |
| MRET_2250 | cell division control protein 14 | 11.56 | 20.1 |
| MRET_2251 | small nuclear ribonucleoprotein E | 69.24 | 189.93 |
| MRET_2252 | protein MAK11 | 69.38 | 172.68 |
| MRET_2253 | transporter | 15.8 | 18.8 |
| MRET_2254 | ESCRT-II complex subunit VPS25 | 15.22 | 15.93 |

| | | | |
|---|---|---|---|
| MRET_2255 | uncharacterized protein | 38.54 | 39.94 |
| MRET_2256 | uracil phosphoribosyltransferase | 26.21 | 37.4 |
| MRET_2257 | LYR motif protein 4 | 536.04 | 528.74 |
| MRET_2258 | serine/threonine-protein kinase | 79.23 | 26.97 |
| MRET_2259 | zinc cluster transcription factor Rds2 | 26.77 | 64.04 |
| MRET_2260 | ornithine decarboxylase | 18.71 | 37.9 |
| MRET_2261 | Sds3-like protein | 46.62 | 70.69 |
| MRET_2262 | NADH dehydrogenase (ubiquinone) 1 alpha subcomplex subunit 5 | 200.13 | 274.76 |
| MRET_2263 | nitric oxide synthase-interacting protein | 129.87 | 139.46 |
| MRET_2264 | mitochondrial fission protein FIS1 | 628.03 | 440.89 |
| MRET_2265 | sorting nexin-4 | 246.91 | 190.61 |
| MRET_2266 | clathrin heavy chain | 148.06 | 125.18 |
| MRET_2267 | 26S proteasome regulatory subunit T4 | 102.45 | 133.44 |
| MRET_2268 | 1-phosphatidylinositol-3-phosphate 5-kinase | 56 | 39.33 |
| MRET_2269 | uncharacterized protein | 87.71 | 64.61 |
| MRET_2270 | ADP-ribosylation factor-binding protein GGA | 58.56 | 86.81 |
| MRET_2271 | uncharacterized protein | 19.52 | 38.43 |
| MRET_2272 | uncharacterized protein | 29.03 | 65.27 |
| MRET_2273 | arginyl-tRNA synthetase | 36.55 | 88.52 |
| MRET_2274 | nuclear GTP-binding protein | 190.06 | 262.52 |
| MRET_2275 | nuclear GTP-binding protein | 141.86 | 184.28 |
| MRET_2276 | viral A-type inclusion protein repeat protein | 5.26 | 48.14 |
| MRET_2277 | gamma-glutamyltranspeptidase/glutathione hydrolase | 166.4 | 71.52 |
| MRET_2278 | mitochondrial serine protease | 116.36 | 50.93 |
| MRET_2279 | T-complex protein 1 subunit eta | 208.02 | 195 |
| MRET_2280 | protein ATS1 | 348 | 74.28 |
| MRET_2281 | membrane associated protein | 192.42 | 60.38 |
| MRET_2282 | large subunit ribosomal protein L3 | 195.49 | 242.85 |
| MRET_2283 | cleavage and polyadenylation specificity factor subunit 2 | 37.83 | 31.09 |
| MRET_2284 | uncharacterized protein | 510.68 | 309.73 |
| MRET_2285 | PHD finger domain protein | 61.2 | 71.33 |
| MRET_2286 | U3 small nucleolar RNA-associated protein 10 | 100.9 | 47.71 |
| MRET_2287 | flap endonuclease-1 | 57.67 | 28.81 |
| MRET_2288 | uncharacterized protein | 27.01 | 30.95 |
| MRET_2289 | diphosphoinositol-polyphosphate diphosphatase | 80.1 | 80.21 |
| MRET_2290 | uncharacterized protein | 125.37 | 201.04 |
| MRET_2291 | uncharacterized protein | 32.43 | 29.17 |

| | | | |
|---|---|---|---|
| MRET_2292 | large subunit ribosomal protein L7Ae | 1816.55 | 2761.65 |
| MRET_2293 | putative stress-responsive nuclear envelope protein | 686.82 | 385.08 |
| MRET_2294 | DNA repair protein RAD50 | 40.76 | 65.34 |
| MRET_2295 | palmitoyltransferase ZDHHC2/15/20 | 754.64 | 354.34 |
| MRET_2296 | ubiquitin-like 1-activating enzyme E1 A | 584.77 | 296.24 |
| MRET_2297 | uncharacterized protein | 297.4 | 298.97 |
| MRET_2298 | RNA polymerase I-specific transcription initiation factor RRN7 | 74.5 | 55.47 |
| MRET_2299 | nucleoprotein TPR | 40.73 | 55.3 |
| MRET_2300 | tail-anchored protein insertion receptor | 35.25 | 156.2 |
| MRET_2301 | glutamine amidotransferase | 74.93 | 124.92 |
| MRET_2302 | conserved hypothetical protein | 273.37 | 378.43 |
| MRET_2303 | p24 family protein alpha | 67.4 | 120.08 |
| MRET_2304 | chloride channel 3/4/5 | 42.39 | 43.7 |
| MRET_2305 | mRNA export factor | 5.7 | 8.22 |
| MRET_2306 | dynein light intermediate chain 1, cytosolic | 34.24 | 43.29 |
| MRET_2307 | uncharacterized protein | 5.9 | 21.51 |
| MRET_2308 | PPR repeat containing protein | 64.42 | 48.51 |
| MRET_2309 | ATP-binding cassette, subfamily D (ALD), peroxisomal long-chain fatty acid import protein | 73.82 | 61.67 |
| MRET_2310 | CCR4-NOT transcriptional complex subunit CAF120 | 64.97 | 53.54 |
| MRET_2311 | gluconokinase | 9.29 | 5.48 |
| MRET_2312 | uncharacterized protein | 41.64 | 40.07 |
| MRET_2313 | fungal Zn(2)-Cys(6) binuclear cluster domain protein | 48.34 | 75.45 |
| MRET_2314 | nucleoporin POM152 | 88.02 | 51.29 |
| MRET_2315 | large subunit ribosomal protein L33 | 60.91 | 96.47 |
| MRET_2316 | uncharacterized protein | 107.41 | 119.71 |
| MRET_2317 | ribosomal protein L23 | 183.3 | 180.02 |
| MRET_2318 | CTD nuclear envelope phosphatase 1 | 23.29 | 23.47 |
| MRET_2319 | serine/threonine-protein kinase Chk1 | 114.78 | 75.84 |
| MRET_2320 | DUF2340 domain protein | 151.36 | 119.95 |
| MRET_2321 | sorting nexin | 48.99 | 34.03 |
| MRET_2322 | DNA excision repair protein ERCC-5 | 377.35 | 197.69 |
| MRET_2323 | NADH dehydrogenase (ubiquinone) 1 beta subcomplex subunit 8 | 198.83 | 159.31 |
| MRET_2324 | serine/threonine-protein phosphatase 2A catalytic subunit | 1217.26 | 1314.11 |
| MRET_2325 | DNA replication licensing factor MCM2 | 36.04 | 47.82 |
| MRET_2326 | serine/threonine-protein kinase | 29.52 | 42.43 |
| MRET_2327 | ubiquitin-conjugating enzyme E2 J2 | 87.77 | 164.96 |
| MRET_2328 | uncharacterized protein | 30.41 | 51.9 |

| | | | |
|---|---|---|---|
| MRET_2329 | ABC transporter | 131.71 | 195.86 |
| MRET_2330 | ABC transporter | 105.07 | 88.01 |
| MRET_2331 | syntaxin 6 | 21.38 | 29.89 |
| MRET_2332 | UDP-glucose:glycoprotein glucosyltransferase | 52.38 | 56.18 |
| MRET_2333 | SNARE associated golgi protein | 59.44 | 69.19 |
| MRET_2334 | transferase CAF17, mitochondrial | 117.61 | 143.36 |
| MRET_2335 | Bacterial low temperature requirement A protein (LtrA) | 39.15 | 92.37 |
| MRET_2336 | 5'-nucleotidase | 59.56 | 61.99 |
| MRET_2337 | HMG (high mobility group) box protein | 282.15 | 224.13 |
| MRET_2338 | glucose-6-phosphate isomerase | 405.45 | 288.99 |
| MRET_2339 | KH domain protein | 203.37 | 162.75 |
| MRET_2340 | uncharacterized protein | 15.01 | 22.85 |
| MRET_2341 | uncharacterized protein | 46.89 | 145.63 |
| MRET_2342 | uncharacterized protein | 110.4 | 286.67 |
| MRET_2343 | DNA polymerase epsilon subunit 1 | 62 | 45.49 |
| MRET_2344 | cytosolic Fe-S cluster assembly factor NBP35 | 323 | 277.88 |
| MRET_2345 | WD repeat and FYVE domain protein 3 | 117.46 | 47.14 |
| MRET_2346 | THO complex subunit 4 | 390.45 | 522.36 |
| MRET_2347 | ubiquinol-cytochrome c reductase subunit 8 | 385.47 | 533.64 |
| MRET_2348 | conserved hypothetical protein | 127.52 | 160 |
| MRET_2349 | mitochondrial NADH kinase | 45.96 | 97.91 |
| MRET_2350 | histone H3 | 3048.02 | 2446.42 |
| MRET_2351 | cell division cycle 20-like protein 1, cofactor of APC complex | 52.26 | 108.48 |
| MRET_2352 | GMP synthase (glutamine-hydrolysing) | 49.09 | 79.37 |
| MRET_2353 | cell cycle arrest protein BUB3 | 128.64 | 131.09 |
| MRET_2354 | cystathionine gamma-synthase | 78.03 | 103.42 |
| MRET_2355 | syntaxin 5 | 37.53 | 75.44 |
| MRET_2356 | succinate dehydrogenase (ubiquinone) cytochrome b560 subunit | 53.04 | 73.16 |
| MRET_2357 | gamma-tubulin complex component 2 | 31.32 | 75.13 |
| MRET_2358 | uroporphyrin-III C-methyltransferase | 25.83 | 59.66 |
| MRET_2359 | S-adenosylmethionine decarboxylase | 57.11 | 84.9 |
| MRET_2360 | zinc metalloprotease | 69.08 | 66.11 |
| MRET_2361 | putative MFS transporter, AGZA family, xanthine/uracil permease | 9.63 | 23.94 |
| MRET_2362 | uncharacterized protein | 145.23 | 276.68 |
| MRET_2363 | uncharacterized protein | 603.25 | 325.78 |
| MRET_2364 | reverse transcriptase | 635.32 | 207.81 |
| MRET_2365 | phospholipase C | 187.36 | 764.21 |

| | | | |
|---|---|---|---|
| MRET_2366 | aryl-alcohol dehydrogenase | 571.39 | 416.95 |
| MRET_2367 | uncharacterized protein | 604.42 | 354.04 |
| MRET_2368 | protein phosphatase methylesterase 1 | 237.84 | 93.25 |
| MRET_2369 | uncharacterized protein | 63.79 | 56.58 |
| MRET_2370 | uncharacterized protein | 59.44 | 40.03 |
| MRET_2371 | tRNA modification GTPase | 28.26 | 45.79 |
| MRET_2372 | uncharacterized protein | 56.86 | 46.6 |
| MRET_2373 | uncharacterized protein | 119.8 | 125.19 |
| MRET_2374 | asparagine | 46.34 | 26.12 |
| MRET_2375 | DUF757 domain protein | 135.25 | 213.88 |
| MRET_2376 | NADH-ubiquinone oxidoreductase subunit | 301.07 | 522.11 |
| MRET_2377 | multifunctional beta-oxidation protein | 526.27 | 312.25 |
| MRET_2378 | small nuclear ribonucleoprotein D2 | 21.65 | 116.23 |
| MRET_2379 | glutamyl-tRNA synthetase | 42.71 | 54.32 |
| MRET_2380 | dihydrofolate reductase | 20.65 | 27.3 |
| MRET_2381 | 2'-phosphotransferase | 31.26 | 35.94 |
| MRET_2382 | V-type H+-transporting ATPase subunit C | 128.94 | 131.56 |
| MRET_2383 | RNA-binding motif protein, X-linked 2 | 95.87 | 72.82 |
| MRET_2384 | protein phosphatase PTC6 | 37.08 | 32.85 |
| MRET_2385 | pre-rRNA processing protein Esf1 | 215.86 | 191.04 |
| MRET_2386 | transcription elongation factor SPT5 | 160.82 | 338.58 |
| MRET_2387 | elongation factor Tu | 296.28 | 341.9 |
| MRET_2388 | large subunit ribosomal protein L31e | 125.23 | 512.9 |
| MRET_2389 | SMN domain protein | 48.01 | 89.01 |
| MRET_2390 | alpha/beta-hydrolase | 50.45 | 50.3 |
| MRET_2391 | subunit of ATP-dependent Isw2p-Itc1p chromatin remodeling complex | 31.56 | 38.49 |
| MRET_2392 | U6 snRNA-associated Sm-like protein LSm5 | 672.31 | 654.36 |
| MRET_2393 | calcineurin-like phosphoesterase | 167.92 | 61.3 |
| MRET_2394 | ketol-acid reductoisomerase | 242.01 | 186 |
| MRET_2395 | homoaconitate hydratase | 109.95 | 91.69 |
| MRET_2396 | tRNA dimethylallyltransferase | 127.16 | 95.07 |
| MRET_2397 | leukotriene-A4 hydrolase | 67.71 | 47.89 |
| MRET_2398 | uncharacterized protein | 19.59 | 31.93 |
| MRET_2399 | uncharacterized protein | 27.22 | 24.96 |
| MRET_2400 | uncharacterized protein | 1927.28 | 1550.38 |
| MRET_2401 | pyruvate kinase | 531.03 | 278.7 |
| MRET_2402 | mediator of RNA polymerase II transcription subunit 5 | 9.69 | 12.1 |

| | | | |
|---|---|---|---|
| MRET_2403 | NADH dehydrogenase (ubiquinone) 1 alpha subcomplex subunit 5 | 155.89 | 112.62 |
| MRET_2404 | alpha-soluble NSF attachment protein | 100.21 | 109.13 |
| MRET_2405 | aspartyl-tRNA(Asn)/glutamyl-tRNA(Gln) amidotransferase subunit B | 131.92 | 69.45 |
| MRET_2406 | serine/threonine-protein phosphatase 2A activator | 19.39 | 23.75 |
| MRET_2407 | threonyl-tRNA synthetase | 35.05 | 59.46 |
| MRET_2408 | uncharacterized protein | 43.25 | 105.3 |
| MRET_2409 | AT-rich interactive domain protein | 21.17 | 31.5 |
| MRET_2410 | uncharacterized protein | 17.78 | 38.65 |
| MRET_2411 | farnesyl-diphosphate farnesyltransferase | 229.77 | 199.03 |
| MRET_2412 | protein KRI1 | 23.86 | 56.68 |
| MRET_2413 | U3 small nucleolar RNA-associated protein 20 | 34.14 | 32.11 |
| MRET_2414 | uncharacterized protein | 58.22 | 48.75 |
| MRET_2415 | SEL1 domain protein | 210.39 | 115.48 |
| MRET_2416 | ER membrane protein that plays a central role in ERAD | 57.97 | 47.9 |
| MRET_2417 | heat shock protein 70 homolog LHS1 | 95.58 | 94.56 |
| MRET_2418 | Ca2+-transporting ATPase | 35.13 | 37.37 |
| MRET_2419 | regulator of chromosome condensation (RCC1) repeat protein | 65.77 | 65.57 |
| MRET_2420 | ribonuclease P protein subunit POP4 | 168.49 | 118.86 |
| MRET_2421 | saccharopine dehydrogenase | 3682.84 | 2228.95 |
| MRET_2422 | T-complex protein 1 subunit theta | 77.61 | 124.82 |
| MRET_2423 | transportin-1 | 38.3 | 39.91 |
| MRET_2424 | xylulokinase | 165.22 | 95.5 |
| MRET_2425 | geranylgeranyl diphosphate synthase, type III | 148.66 | 70.87 |
| MRET_2426 | DNA-directed RNA polymerases I, II, and III subunit RPABC3 | 45.49 | 92.21 |
| MRET_2427 | TBC1 domain family member 2 | 240.47 | 155.44 |
| MRET_2428 | RNA exonuclease 4 | 21.3 | 16.01 |
| MRET_2429 | uncharacterized protein | 95.25 | 56.03 |
| MRET_2430 | phospholipid-transporting ATPase | 55.36 | 57.02 |
| MRET_2431 | uncharacterized protein | 72.16 | 41.8 |
| MRET_2432 | alkaline ceramidase | 186.06 | 123.3 |
| MRET_2433 | DUF850 domain protein | 61.41 | 84.39 |
| MRET_2434 | structural maintenance of chromosomes protein | 26.46 | 28.92 |
| MRET_2435 | uncharacterized protein | 22.04 | 35.61 |
| MRET_2436 | bZIP transcription factor | 333.42 | 238.02 |
| MRET_2437 | transcription initiation factor TFIID subunit 10 | 295.2 | 312.92 |
| MRET_2438 | uncharacterized protein | 25.37 | 26.37 |
| MRET_2439 | leucine-rich repeat protein | 44.83 | 58.04 |

| | | | |
|---|---|---|---|
| MRET_2440 | solute carrier family 39 (zinc transporter), member 9 | 454.94 | 210.46 |
| MRET_2441 | 2-nitropropane dioxygenase | 369.22 | 344.15 |
| MRET_2442 | ER lumen protein retaining receptor | 1261.74 | 343.69 |
| MRET_2443 | negative regulator of differentiation 1 | 116.16 | 70.76 |
| MRET_2444 | solute carrier family 25 (mitochondrial citrate transporter), member 1 | 242.9 | 125.88 |
| MRET_2445 | cell polarity protein | 57.63 | 38.24 |
| MRET_2446 | mitochondrial matrix iron chaperone | 42.17 | 34.49 |
| MRET_2447 | protein kinase A | 71.47 | 78.29 |
| MRET_2448 | cerevisin | 151.82 | 120.8 |
| MRET_2449 | asparaginyl-tRNA synthetase | 24.87 | 44.99 |
| MRET_2450 | essential protein required for maturation of 18S rRNA | 24.34 | 47.33 |
| MRET_2451 | transcriptional activator SPT7 | 154.53 | 157.38 |
| MRET_2452 | translation initiation factor 4G | 1168.35 | 1749.77 |
| MRET_2453 | uncharacterized protein | 90.24 | 29.57 |
| MRET_2454 | AHNAK nucleoprotein | 561 | 1233.22 |
| MRET_2455 | mitochondrial GTPase MTG1 | 287.74 | 332.45 |
| MRET_2456 | uncharacterized protein | 8.67 | 16.44 |
| MRET_2457 | pseudouridine synthase | 41.3 | 57.45 |
| MRET_2458 | uncharacterized protein | 140.42 | 112.53 |
| MRET_2459 | nuclear polyadenylated RNA-binding protein | 363.8 | 291.52 |
| MRET_2460 | U6 snRNA-associated Sm-like protein LSm4 | 57.71 | 169.08 |
| MRET_2461 | peptidyl-prolyl cis-trans isomerase | 92.18 | 65.88 |
| MRET_2462 | mediator of RNA polymerase II transcription subunit 17, fungi type | 125.9 | 84.13 |
| MRET_2463 | histidinol-phosphate aminotransferase | 21.42 | 46.16 |
| MRET_2464 | pre-mRNA-splicing factor CDC5/CEF1 | 179.82 | 159.03 |
| MRET_2465 | vacuolar protein sorting-associated protein | 356.82 | 221.97 |
| MRET_2466 | uncharacterized protein | 41.06 | 116.01 |
| MRET_2467 | calcium sensor Efh | 59.99 | 72.34 |
| MRET_2468 | nicotinate phosphoribosyltransferase | 528.14 | 487.1 |
| MRET_2469 | dynactin 5 | 805.55 | 425.13 |
| MRET_2470 | dynamin-binding protein | 433.17 | 183.42 |
| MRET_2471 | glucan 1,3-beta-glucosidase | 27.25 | 55.43 |
| MRET_2472 | Sad1 UNC domain protein | 12.79 | 25.62 |
| MRET_2473 | vacuole protein | 70.49 | 105.02 |
| MRET_2474 | potassium transport protein | 304.89 | 39.89 |
| MRET_2475 | PITH domain protein | 74.87 | 55.48 |
| MRET_2476 | SWI/SNF-related matrix-associated actin-dependent regulator of chromatin subfamily B member 1 | 82.56 | 52.36 |

| | | | |
|---|---|---|---|
| MRET_2477 | small subunit ribosomal protein S2 | 18.64 | 55.39 |
| MRET_2478 | heat shock 70kDa protein 1/2/6/8 | 104.41 | 292.78 |
| MRET_2479 | ATP-binding cassette, subfamily B (MDR/TAP), member 1 | 12 | 9.69 |
| MRET_2480 | cyclin | 74.61 | 8.3 |
| MRET_2481 | translational activator GCN1 | 10.43 | 17.68 |
| MRET_2482 | guanine nucleotide-binding protein G(i) subunit alpha | 38.13 | 64.59 |
| MRET_2483 | L-lactate dehydrogenase (cytochrome) | 134.75 | 221.67 |
| MRET_2484 | intracellular protein transport protein USO1 | 60.13 | 55.36 |
| MRET_2485 | uncharacterized protein | 19.56 | 26.18 |
| MRET_2486 | WD repeat protein 59 | 19.07 | 23.18 |
| MRET_2487 | uncharacterized protein | 24.58 | 63.45 |
| MRET_2488 | tyrosine-protein phosphatase 2/3 | 17.12 | 27.55 |
| MRET_2489 | lectin, mannose-binding 2 | 41.2 | 82.9 |
| MRET_2490 | oligoribonuclease | 15.4 | 18.63 |
| MRET_2491 | calpain-7 | 73.98 | 62.35 |
| MRET_2492 | 3-methyl-2-oxobutanoate hydroxymethyltransferase | 28.23 | 53.83 |
| MRET_2493 | ABC transporter | 78.86 | 106.47 |
| MRET_2494 | mannosyl-oligosaccharide alpha-1,3-glucosidase | 76.26 | 70.47 |
| MRET_2495 | translation initiation factor 2A | 188.59 | 150.1 |
| MRET_2496 | type I protein arginine methyltransferase | 169.73 | 159.54 |
| MRET_2497 | uncharacterized protein | 19.15 | 51.62 |
| MRET_2498 | uncharacterized protein | 19.97 | 42.7 |
| MRET_2499 | autophagy-related protein 101 | 111.9 | 39.58 |
| MRET_2500 | zinc metalloprotease | 94.64 | 59.67 |
| MRET_2501 | tRNA uridine 5-carboxymethylaminomethyl modification enzyme | 178.26 | 109.81 |
| MRET_2502 | GTP binding protein | 28.84 | 33.27 |
| MRET_2503 | uncharacterized protein | 26.55 | 29.35 |
| MRET_2504 | inositol phospholipid synthesis and fat-storage-inducing TM | 18.3 | 52.13 |
| MRET_2505 | Pin2-interacting protein X1 | 29.96 | 94.03 |
| MRET_2506 | farnesyl diphosphate synthase | 53.43 | 91.38 |
| MRET_2507 | omega-amidase | 225.57 | 123.83 |
| MRET_2508 | succinate dehydrogenase assembly factor 1 | 210.49 | 278.72 |
| MRET_2509 | mitochondrial inner membrane protease subunit 2 | 313.28 | 240.81 |
| MRET_2510 | endopolyphosphatase | 86.28 | 150.52 |
| MRET_2511 | isocitrate lyase | 488.65 | 188.01 |
| MRET_2512 | DNA-directed RNA polymerase III subunit RPC4 | 34.72 | 38.09 |
| MRET_2513 | conserved hypothetical protein | 807.66 | 202.3 |

| | | | |
|---|---|---|---|
| MRET_2514 | growth hormone-inducible transmembrane protein | 3309.3 | 1919.61 |
| MRET_2515 | profilin | 75.78 | 65.96 |
| MRET_2516 | triosephosphate isomerase (TIM) | 672.33 | 1189.74 |
| MRET_2517 | protein PET117 | 127.81 | 105.41 |
| MRET_2518 | uncharacterized protein | 40.14 | 93.03 |
| MRET_2519 | phosphoribosylaminoimidazolecarboxamide formyltransferase/IMP cyclohydrolase | 110.47 | 60.38 |
| MRET_2520 | vacuolar fusion protein MON1 | 13.32 | 11.46 |
| MRET_2521 | DUF890 domain protein | 8.63 | 10.63 |
| MRET_2522 | lipoyl synthase | 1136.01 | 849.79 |
| MRET_2523 | uncharacterized protein | 76.29 | 77.11 |
| MRET_2524 | mitochondrial alcohol dehydrogenase isozyme III | 3331.16 | 2583.11 |
| MRET_2525 | DUF202 domain protein | 257.77 | 330.93 |
| MRET_2526 | cytochrome c oxidase subunit 17 | 1256.64 | 786.61 |
| MRET_2527 | large subunit ribosomal protein LP1 | 148.21 | 665.83 |
| MRET_2528 | trehalase | 172.13 | 139.48 |
| MRET_2529 | uncharacterized protein | 185.8 | 75.27 |
| MRET_2530 | acetolactate synthase I/III small subunit | 203.58 | 246.23 |
| MRET_2531 | oxidoreductase | 582.34 | 436.44 |
| MRET_2532 | regulatory subunit of the type I protein phosphatase | 970.03 | 924.69 |
| MRET_2533 | AHNAK nucleoprotein | 123.51 | 188 |
| MRET_2534 | uncharacterized protein | 1705.5 | 1730.02 |
| MRET_2535 | DASH complex subunit DAD4 | 33.05 | 102.46 |
| MRET_2536 | ribosomal protein S21 | 20.7 | 63.6 |
| MRET_2537 | uncharacterized protein | 204.5 | 193.14 |
| MRET_2538 | protein ROT1 | 134.77 | 20.19 |
| MRET_2539 | amino-acid N-acetyltransferase | 52.31 | 35.44 |
| MRET_2540 | plasma membrane protein involved in remodeling GPI anchors | 53.42 | 37.74 |
| MRET_2541 | AP-1 complex subunit sigma 1/2 | 46.73 | 91.36 |
| MRET_2542 | uncharacterized protein | 42.77 | 64.12 |
| MRET_2543 | EKC/KEOPS complex subunit PCC1/LAGE3 | 219.04 | 224.02 |
| MRET_2544 | ATP-dependent DNA helicase PIF1 | 165.42 | 203.81 |
| MRET_2545 | translation initiation factor eIF-2B subunit beta | 115.6 | 110.46 |
| MRET_2546 | conserved hypothetical protein | 64.73 | 48.32 |
| MRET_2547 | origin recognition complex subunit 4 | 16.6 | 20.71 |
| MRET_2548 | elongator complex protein 4 | 42.87 | 59.18 |
| MRET_2549 | 26S proteasome regulatory subunit T6 | 165.58 | 169.8 |
| MRET_2550 | mitochondrial import receptor subunit TOM71 | 23.57 | 40.45 |

| | | | |
|---|---|---|---|
| MRET_2551 | tRNA-dihydrouridine synthase 1 | 10.36 | 22.85 |
| MRET_2552 | sphingosine-1-phosphate phosphohydrolase | 111.54 | 224.46 |
| MRET_2553 | NADH dehydrogenase (ubiquinone) 1 alpha subcomplex subunit 3 | 225.44 | 206.15 |
| MRET_2554 | peptidyl-prolyl isomerase H (cyclophilin H) | 90.25 | 94.68 |
| MRET_2555 | para-aminobenzoate (PABA) synthase | 58.14 | 116.46 |
| MRET_2556 | uncharacterized protein | 80.9 | 166.89 |
| MRET_2557 | uncharacterized protein | 2794.26 | 1774.81 |
| MRET_2558 | mitochondrial peroxiredoxin PRX1 | 145.33 | 92.55 |
| MRET_2559 | short-chain dehydrogenase reductase | 40.97 | 45.6 |
| MRET_2560 | calcium permeable stress-gated cation channel | 15.24 | 29.33 |
| MRET_2561 | ubiquitin related modifier 1 | 14.72 | 14.51 |
| MRET_2562 | metal transporter CNNM | 23.48 | 56.33 |
| MRET_2563 | uncharacterized protein | 79.9 | 146.03 |
| MRET_2564 | cytoplasmic tRNA 2-thiolation protein 1 | 116.57 | 93 |
| MRET_2565 | ubiquinone biosynthesis protein COQ4 | 78.48 | 81.52 |
| MRET_2566 | uncharacterized protein | 297.52 | 222.53 |
| MRET_2567 | cation diffusion facilitator | 128.74 | 87.02 |
| MRET_2568 | 3-isopropylmalate dehydrogenase | 101.89 | 118.49 |
| MRET_2569 | A transporter | 25.44 | 43.5 |
| MRET_2570 | thioesterase | 108.79 | 134.5 |
| MRET_2571 | transporter | 116.21 | 41.66 |
| MRET_2572 | U4/U6.U5 tri-snRNP-associated protein 2 | 38.74 | 51.91 |
| MRET_2573 | aspartyl aminopeptidase | 181.16 | 123.99 |
| MRET_2574 | zinc finger protein, GATA type | 59.77 | 35.19 |
| MRET_2575 | uncharacterized protein | 35.87 | 40.62 |
| MRET_2576 | exosome complex component CSL4 | 73.94 | 77.19 |
| MRET_2577 | protein LTV1 | 88.25 | 77.22 |
| MRET_2578 | uncharacterized protein | 39.93 | 139.49 |
| MRET_2579 | pyruvate dehydrogenase E2 component (dihydrolipoamide acetyltransferase) | 880.08 | 523.93 |
| MRET_2580 | RecQ-mediated genome instability protein 1 | 556.67 | 320.2 |
| MRET_2581 | proteophosphoglycan ppg4 | 497.89 | 251.5 |
| MRET_2582 | uncharacterized protein | 415.21 | 266.07 |
| MRET_2583 | uncharacterized protein | 17.37 | 19.32 |
| MRET_2584 | DNA-directed RNA polymerase II subunit RPB2 | 259.03 | 255.97 |
| MRET_2585 | pre-mRNA-splicing factor CWC25 | 7.18 | 18.24 |
| MRET_2586 | integral membrane protein (Ptm1) | 37.12 | 88.05 |
| MRET_2587 | aminoacyl tRNA synthase complex-interacting multifunctional protein 1 | 72.8 | 140.83 |

| | | | |
|---|---|---|---|
| MRET_2588 | zinc finger protein, C3H1 type | 35.74 | 46.7 |
| MRET_2589 | aminomethyltransferase | 56.91 | 99.2 |
| MRET_2590 | transporter | 53.74 | 37.49 |
| MRET_2591 | nuclear fragile X mental retardation-interacting protein 1 (NUFIP1) | 36.74 | 46.59 |
| MRET_2592 | large subunit ribosomal protein L4 | 35.27 | 137.61 |
| MRET_2593 | small subunit ribosomal protein S16e | 105.91 | 436.13 |
| MRET_2594 | SWI/SNF-related matrix-associated actin-dependent regulator of chromatin subfamily D | 90.35 | 104.88 |
| MRET_2595 | Ras-related protein Rab-5C | 91.13 | 191.46 |
| MRET_2596 | solute carrier family 25 (mitochondrial adenine nucleotide translocator), member 4/5/6/31 | 3307.35 | 2297.94 |
| MRET_2597 | UBX domain protein 1 | 966.28 | 683.79 |
| MRET_2598 | Tim17/Tim22/Tim23/Pmp24 family protein | 201.13 | 316.42 |
| MRET_2599 | aminopeptidase | 423.62 | 636.06 |
| MRET_2600 | aminopeptidase | 27.14 | 32.31 |
| MRET_2601 | uncharacterized protein | 6.58 | 10.43 |
| MRET_2602 | tRNA acetyltransferase TAN1 | 12.64 | 18.13 |
| MRET_2603 | 8-amino-7-oxononanoate synthase | 11.29 | 10.89 |
| MRET_2604 | Vam6/Vps39-like protein vacuolar protein sorting-associated protein 39 | 66.22 | 43.7 |
| MRET_2605 | uncharacterized protein | 202.17 | 131.96 |
| MRET_2606 | uncharacterized protein | 26.34 | 39.35 |
| MRET_2607 | RNA-binding protein | 75.73 | 84.21 |
| MRET_2608 | chitin synthase | 199.81 | 91.72 |
| MRET_2609 | SH3 domain protein | 9.49 | 41.85 |
| MRET_2610 | 4-aminobutyrate aminotransferase | 173.17 | 289.87 |
| MRET_2611 | FH domain protein | 17.63 | 36.81 |
| MRET_2612 | cullin 3 | 64.28 | 154.81 |
| MRET_2613 | serine/threonine-protein kinase MRCK | 783.41 | 630.66 |
| MRET_2614 | transcription factor | 140.98 | 151.29 |
| MRET_2615 | uncharacterized protein | 414.68 | 406.22 |
| MRET_2616 | Hsp70 nucleotide exchange factor FES1 | 66.72 | 104.11 |
| MRET_2617 | anaphase-promoting complex subunit 6 | 35 | 70.13 |
| MRET_2618 | A/G-specific adenine glycosylase | 31.74 | 70.26 |
| MRET_2619 | uncharacterized protein | 11588.3 | 2892.69 |
| MRET_2620 | AP-3 complex subunit sigma | 37.06 | 59.8 |
| MRET_2621 | ADP-ribosylation factor-like protein 2 | 49.15 | 121.71 |
| MRET_2622 | acyl-CoA dehydrogenase | 839.54 | 1062.57 |
| MRET_2623 | uncharacterized protein | 8.18 | 20.63 |
| MRET_2624 | exosome complex protein LRP1 | 8.94 | 15.23 |

| | | | |
|---|---|---|---|
| MRET_2625 | homeobox domain protein | 6.68 | 22.45 |
| MRET_2626 | homeobox domain protein | 15.29 | 10.97 |
| MRET_2627 | methionine permease | 31.22 | 88.54 |
| MRET_2628 | ATP-dependent RNA helicase DHX29 | 76.97 | 56.84 |
| MRET_2629 | INO80 complex subunit 1 | 42.87 | 59.49 |
| MRET_2630 | uncharacterized protein | 102.48 | 89.47 |
| MRET_2631 | DNA 5' AMP hydrolase involved in DNA repair | 24.15 | 84.53 |
| MRET_2632 | uncharacterized protein | 74.12 | 93.87 |
| MRET_2633 | N5-hydroxy-L-ornithine N5-transacylase | 80.9 | 170.79 |
| MRET_2634 | L-ornithine N5-monooxygenase | 92.99 | 176.26 |
| MRET_2635 | exocyst complex component 1 | 60.46 | 77.19 |
| MRET_2636 | DUF2305 domain protein | 40.46 | 60.57 |
| MRET_2637 | WD repeat protein | 47.04 | 30.93 |
| MRET_2638 | aspartate aminotransferase, mitochondrial | 205.68 | 337.46 |
| MRET_2639 | DNA polymerase mu | 46.9 | 107.19 |
| MRET_2640 | uncharacterized protein | 35.89 | 84.49 |
| MRET_2641 | uncharacterized protein | 45.01 | 132.76 |
| MRET_2642 | bacterial leucyl aminopeptidase | 51.32 | 189.19 |
| MRET_2643 | rRNA biogenesis protein RRP5 | 77.07 | 427.21 |
| MRET_2644 | Src like adaptor protein | 122.86 | 105.01 |
| MRET_2645 | uncharacterized protein | 26.51 | 58.74 |
| MRET_2646 | AN1-like zinc finger protein | 20.62 | 38.95 |
| MRET_2647 | antiviral helicase SKI2 | 124.11 | 136.85 |
| MRET_2648 | uncharacterized protein | 27.7 | 35.75 |
| MRET_2649 | phosphatidylglycerophosphatase GEP4 | 18.55 | 31.1 |
| MRET_2650 | transporter (MirC) | 19.14 | 106.54 |
| MRET_2651 | uncharacterized protein | 45.17 | 121.87 |
| MRET_2652 | plastin-1 | 43.79 | 86.83 |
| MRET_2653 | low-affinity vacuolar phosphate transporter | 110.77 | 90.39 |
| MRET_2654 | solute carrier family 25, member 39/40 | 266.37 | 160.5 |
| MRET_2655 | tRNA pseudouridine38/39 synthase | 198.94 | 142.26 |
| MRET_2656 | SET domain protein | 94.76 | 63.37 |
| MRET_2657 | peptidyl-prolyl isomerase E (cyclophilin E) | 205.16 | 192.12 |
| MRET_2658 | isopentenyl-diphosphate Delta-isomerase | 219.04 | 309.4 |
| MRET_2659 | uncharacterized protein | 17.57 | 145.16 |
| MRET_2660 | prephenate dehydratase | 15.08 | 44.16 |
| MRET_2661 | secreted aspartic endopeptidase | 73.69 | 100.25 |

| | | | |
|---|---|---|---|
| MRET_2662 | hydrolase, family 43 protein | 74.65 | 84.43 |
| MRET_2663 | GTPase KRas | 309.96 | 539.06 |
| MRET_2664 | nucleoside diphosphatase | 102.94 | 92.8 |
| MRET_2665 | exocyst complex component 3 | 13.65 | 17.18 |
| MRET_2666 | CCCH finger DNA binding protein | 6.57 | 46.29 |
| MRET_2667 | outer membrane protein TOM13 | 0.7 | 15.12 |
| MRET_2668 | uncharacterized protein | 5.28 | 78.32 |
| MRET_2669 | phosphoribosylaminoimidazole carboxylase | 256.96 | 126.61 |
| MRET_2670 | choline oxidase | 210.02 | 200.08 |
| MRET_2671 | uncharacterized protein | 262.74 | 179.37 |
| MRET_2672 | E3 ubiquitin-protein ligase MARCH6 | 34.71 | 48.45 |
| MRET_2673 | aromatic-L-amino-acid/L-tryptophan decarboxylase | 22.45 | 46.79 |
| MRET_2674 | nuclear pore complex protein Nup205 | 62.67 | 45.39 |
| MRET_2675 | uncharacterized protein | 9.35 | 14.38 |
| MRET_2676 | DUF2315 domain protein | 31.45 | 58.83 |
| MRET_2677 | DNA-directed RNA polymerase III subunit RPC7 | 127.46 | 126.34 |
| MRET_2678 | stress-induced-phosphoprotein 1 | 568.05 | 1032.79 |
| MRET_2679 | uncharacterized protein | 42.48 | 73.64 |
| MRET_2680 | cleavage stimulation factor subunit 3 | 24.99 | 52.69 |
| MRET_2681 | mitochondrial protein sorting (Msf1) | 119.93 | 108.1 |
| MRET_2682 | uncharacterized protein | 66.33 | 48.43 |
| MRET_2683 | uncharacterized protein | 32.16 | 70.54 |
| MRET_2684 | U3 small nucleolar RNA-associated protein 5 | 30.3 | 49.12 |
| MRET_2685 | negative regulator of the PHO system | 168.12 | 374.12 |
| MRET_2686 | cell division cycle 20, cofactor of APC complex | 42.23 | 186.18 |
| MRET_2687 | glucose transporter | 34.28 | 31.71 |
| MRET_2688 | ferredoxin-2, mitochondrial | 217.8 | 163.68 |
| MRET_2689 | DEAD/DEAH box helicase | 163.31 | 100.14 |
| MRET_2690 | beta-glucan synthesis-associated protein KRE6 | 61.31 | 120.33 |
| MRET_2691 | cellular nucleic acid-binding protein | 2038.08 | 1909.38 |
| MRET_2692 | signal recognition particle subunit SRP19 | 87.42 | 161.24 |
| MRET_2693 | prolyl-tRNA synthetase | 26.28 | 54 |
| MRET_2694 | seryl-tRNA synthetase | 54.48 | 103.31 |
| MRET_2695 | DnaJ homolog subfamily C member 2 | 60.62 | 102.55 |
| MRET_2696 | ribonuclease h-like protein | 6.71 | 24.83 |
| MRET_2697 | glycosyl hydrolases family 8 | 50.19 | 105.7 |
| MRET_2698 | small subunit ribosomal protein S17e | 77.56 | 326.45 |

| | | | |
|---|---|---|---|
| MRET_2699 | small subunit ribosomal protein S15Ae | 151.34 | 666.55 |
| MRET_2700 | THO complex subunit 7 | 66.56 | 157.68 |
| MRET_2701 | large subunit ribosomal protein L14e | 97.2 | 545.24 |
| MRET_2702 | inositol-hexakisphosphate 5-kinase | 100.48 | 78.84 |
| MRET_2703 | serine/threonine-protein kinase | 84.62 | 80.78 |
| MRET_2704 | uncharacterized protein | 44.97 | 61.59 |
| MRET_2705 | uncharacterized protein | 119.63 | 77.6 |
| MRET_2706 | mitochondrial-processing peptidase subunit alpha | 427.14 | 363.6 |
| MRET_2707 | PHD finger domain protein | 10.05 | 22.44 |
| MRET_2708 | uncharacterized protein | 11.48 | 40.35 |
| MRET_2709 | ubiquinol-cytochrome c reductase subunit 9 | 262.25 | 371.17 |
| MRET_2710 | tether containing UBX domain for GLUT4 | 25.99 | 59.35 |
| MRET_2711 | synaptojanin | 193.01 | 85.4 |
| MRET_2712 | NADH dehydrogenase | 249.57 | 192.47 |
| MRET_2713 | amino acid transporter | 93.31 | 67.48 |
| MRET_2714 | oxidoreductase | 2411.13 | 708.94 |
| MRET_2715 | arp2 3 complex 34 kda subunit | 25.68 | 26.08 |
| MRET_2716 | uncharacterized protein | 121.72 | 108.02 |
| MRET_2717 | heat shock factor-binding protein 1 | 127.36 | 130.45 |
| MRET_2718 | CORD and CS domain protein | 647.58 | 405.81 |
| MRET_2719 | uncharacterized protein | 50.58 | 81.1 |
| MRET_2720 | COMPASS component SWD2 | 22.06 | 44.34 |
| MRET_2721 | enoyl-CoA hydratase/isomerase family | 88.97 | 68.9 |
| MRET_2722 | enoyl-CoA hydratase/isomerase family | 145.67 | 103.03 |
| MRET_2723 | leucine-rich repeat protein | 219.27 | 140.71 |
| MRET_2724 | uncharacterized protein | 46.78 | 50.58 |
| MRET_2725 | large subunit ribosomal protein LP0 | 333.49 | 763.31 |
| MRET_2726 | uncharacterized protein | 330.48 | 175.01 |
| MRET_2727 | glutamate decarboxylase | 140.98 | 187.07 |
| MRET_2728 | DNA-(apurinic or apyrimidinic site) lyase | 32.85 | 71.79 |
| MRET_2729 | chromosome transmission fidelity protein 18 | 28.17 | 33.13 |
| MRET_2730 | U5 small nuclear ribonucleoprotein component | 36.91 | 41.75 |
| MRET_2731 | solute carrier family 45, member 1/2/4 | 8.89 | 23.09 |
| MRET_2732 | GTP-binding protein | 26.94 | 92.86 |
| MRET_2733 | uncharacterized protein | 29.76 | 57.58 |
| MRET_2734 | bloom syndrome protein | 102.33 | 74.64 |
| MRET_2735 | 6-phosphogluconate dehydrogenase | 2015.57 | 1584.63 |

| | | | |
|---|---|---|---|
| MRET_2736 | small glutamine-rich tetratricopeptide repeat protein alpha | 117.45 | 182.75 |
| MRET_2737 | sterol-sensing domain of SREBP cleavage-activation protein | 117.93 | 109.51 |
| MRET_2738 | cohesin complex subunit SCC1 | 95.06 | 42.93 |
| MRET_2739 | solute carrier family 38 (sodium-coupled neutral amino acid transporter), member 11 | 70.77 | 65.34 |
| MRET_2740 | GTPase-activating protein | 22.73 | 32.06 |
| MRET_2741 | F-box protein 9 | 15.51 | 18.71 |
| MRET_2742 | transcription factor | 248.67 | 174.88 |
| MRET_2743 | actin cytoskeleton-regulatory complex protein SLA1 | 279.54 | 209.26 |
| MRET_2744 | START domain protein | 249.32 | 150.37 |
| MRET_2745 | protein MAK16 | 48.22 | 99.77 |
| MRET_2746 | TKL protein kinase | 123.99 | 130.1 |
| MRET_2747 | uncharacterized protein | 4997.78 | 1717.44 |
| MRET_2748 | uncharacterized protein | 130.65 | 121.74 |
| MRET_2749 | uncharacterized protein | 467.99 | 196.26 |
| MRET_2750 | cyclin-dependent protein kinase regulator Pho80 | 86.69 | 244.13 |
| MRET_2751 | pre-mRNA-splicing factor 18 | 157.07 | 146.82 |
| MRET_2752 | cytochrome c oxidase subunit 6b | 1422.35 | 1176.63 |
| MRET_2753 | tRNA (uracil-5-)-methyltransferase TRM9 | 38.06 | 65.8 |
| MRET_2754 | chitin binding peritrophin-A domain protein | 538.27 | 304.38 |
| MRET_2755 | diacylglycerol kinase catalytic domain protein | 176.73 | 95.16 |
| MRET_2756 | conserved hypothetical protein | 60.66 | 39.89 |
| MRET_2757 | uncharacterized protein | 336.18 | 268.73 |
| MRET_2758 | T-complex protein 1 subunit delta | 46.81 | 81.68 |
| MRET_2759 | ubiquitin fusion degradation protein 1 | 525.48 | 499.91 |
| MRET_2760 | demethylmenaquinone methyltransferase | 703.23 | 358.15 |
| MRET_2761 | Csr1-phosphatidylinositol transfer protein | 143.59 | 145.57 |
| MRET_2762 | zinc finger protein, C2H2 type | 525.59 | 442.72 |
| MRET_2763 | L-methionine (R)-S-oxide reductase | 32.62 | 83.87 |
| MRET_2764 | homoserine O-acetyltransferase | 59.45 | 151.52 |
| MRET_2765 | histidyl-tRNA synthetase | 174.39 | 157.1 |
| MRET_2766 | membrane fusion protein Use1 | 24.37 | 49.07 |
| MRET_2767 | 2-iminobutanoate/2-iminopropanoate deaminase | 170.86 | 183.1 |
| MRET_2768 | RhoGEF domain protein | 60.07 | 55.92 |
| MRET_2769 | NADH dehydrogenase (ubiquinone) Fe-S protein 5 | 335.61 | 299.85 |
| MRET_2770 | phosphatidylinositol glycan, class Q | 46.79 | 35.08 |
| MRET_2771 | Ras-related protein Rab-6A | 182.32 | 252.41 |
| MRET_2772 | seipin | 35.39 | 41.01 |

| | | | |
|---|---|---|---|
| MRET_2773 | immunoglobulin-binding protein 1 | 32.03 | 48.72 |
| MRET_2774 | sphingosine kinase | 86.53 | 51.78 |
| MRET_2775 | protein disulfide-isomerase A6 | 357.31 | 274.3 |
| MRET_2776 | uncharacterized protein | 7.91 | 13.38 |
| MRET_2777 | urease | 25.92 | 38.5 |
| MRET_2778 | ubiquitin-like protein Nedd8 | 33.46 | 53.35 |
| MRET_2779 | peroxiredoxin Q/BCP | 77.81 | 60.7 |
| MRET_2780 | ESCRT-I complex subunit TSG101 | 90.77 | 60.46 |
| MRET_2781 | ATP-dependent RNA helicase MRH4, mitochondrial | 36.38 | 22.17 |
| MRET_2782 | exopolyphosphatase | 228.63 | 189.58 |
| MRET_2783 | tyrosine-protein phosphatase SIW14 | 195.12 | 122.67 |
| MRET_2784 | YagE family protein | 222.93 | 128.79 |
| MRET_2785 | vacuolar protein sorting-associated protein 54 | 46.4 | 39.24 |
| MRET_2786 | BTB domain and ankyrin repeat protein | 15.98 | 37.68 |
| MRET_2787 | uncharacterized protein | 26.76 | 60.16 |
| MRET_2788 | Obg-like ATPase 1 | 129.17 | 432.44 |
| MRET_2789 | uncharacterized protein | 1939.8 | 1255.73 |
| MRET_2790 | ribonuclease H2 subunit A | 56.95 | 74.65 |
| MRET_2791 | exosome complex exonuclease RRP6 | 72.53 | 97.58 |
| MRET_2792 | cellulase (glycosyl hydrolase family 5) | 131.54 | 176.95 |
| MRET_2793 | cellulase (glycosyl hydrolase family 5) | 140.67 | 135.47 |
| MRET_2794 | ATP-dependent RNA helicase | 135.25 | 179.84 |
| MRET_2795 | splicing factor 3A subunit 3 | 46.71 | 90.56 |
| MRET_2796 | transcription regulator staf-5 like protein | 582.94 | 452.36 |
| MRET_2797 | uncharacterized protein | 62.92 | 164.76 |
| MRET_2798 | ribonuclease T2 | 298.02 | 231.64 |
| MRET_2799 | integrase | 23.21 | 29.58 |
| MRET_2800 | DNA-directed RNA polymerase I subunit RPA1 | 199.55 | 124.88 |
| MRET_2801 | tRNA threonylcarbamoyladenosine dehydratase | 189.07 | 206.85 |
| MRET_2802 | 6-phosphofructo-2-kinase/fructose-2,6-biphosphatase 2 | 52.5 | 71.08 |
| MRET_2803 | uncharacterized protein | 23.86 | 38.27 |
| MRET_2804 | methionyl aminopeptidase | 73.18 | 101.64 |
| MRET_2805 | ubiquitin-conjugating enzyme E2 M | 164.75 | 52.03 |
| MRET_2806 | thiosulfate/3-mercaptopyruvate sulfurtransferase | 1054.74 | 454.2 |
| MRET_2807 | arrestin-related trafficking adapter 3/6 | 154.74 | 300.4 |
| MRET_2808 | nuclear pore complex protein Nup188 | 37.2 | 38.5 |
| MRET_2809 | conserved oligomeric golgi complex subunit 6 | 135.57 | 107.12 |

| | | | |
|---|---|---|---|
| MRET_2810 | uncharacterized protein | 838.47 | 491.26 |
| MRET_2811 | mitochondrial genome maintenance protein MGR2 | 793.17 | 659.51 |
| MRET_2812 | methyltransferase-like protein 6 | 362.39 | 341.88 |
| MRET_2813 | nuclear pore complex protein Nup133 | 95.9 | 55.96 |
| MRET_2814 | protein of unknown function (DUF1682) | 87.83 | 95.51 |
| MRET_2815 | ubiquitin carboxyl-terminal hydrolase 7 | 435.79 | 204.16 |
| MRET_2816 | chalcone-flavanone isomerase | 1430.4 | 589.45 |
| MRET_2817 | glucosamine-phosphate N-acetyltransferase | 59.54 | 48.72 |
| MRET_2818 | alpha/beta-hydrolase | 41 | 59.99 |
| MRET_2819 | uncharacterized protein | 47.23 | 70.65 |
| MRET_2820 | uncharacterized protein | 93.05 | 126.81 |
| MRET_2821 | Taurine dioxygenase | 40.91 | 57.21 |
| MRET_2822 | eukaryotic aspartyl protease | 108.27 | 129.89 |
| MRET_2823 | uncharacterized protein | 106.66 | 199.96 |
| MRET_2824 | uncharacterized protein | 463.44 | 1914.71 |
| MRET_2825 | secreted aspartic endopeptidase | 144.85 | 197.09 |
| MRET_2826 | lipase precursor-like protein | 6.28 | 12.66 |
| MRET_2827 | eukaryotic aspartyl protease | 0 | 0 |
| MRET_2828 | uncharacterized protein | 21.27 | 109.47 |
| MRET_2829 | uncharacterized protein | 23.94 | 55.73 |
| MRET_2830 | MFS monocarboxylate transporter | 10.01 | 46.96 |
| MRET_2831 | phenylalanyl-tRNA synthetase beta chain | 43.7 | 98.22 |
| MRET_2832 | electron-transferring-flavoprotein dehydrogenase | 124.35 | 57.61 |
| MRET_2833 | Rare lipoprotein A (RlpA)-like double-psi beta-barrel | 68.51 | 82.44 |
| MRET_2834 | prefoldin beta subunit | 20.35 | 69.22 |
| MRET_2835 | rabenosyn-5 | 23.4 | 46.41 |
| MRET_2836 | carboxyl methyltransferase | 8.64 | 15.74 |
| MRET_2837 | 5-methyltetrahydropteroyltriglutamate--homocysteine methyltransferase | 432.86 | 862.01 |
| MRET_2838 | uncharacterized protein | 162.77 | 1805.05 |
| MRET_2839 | uncharacterized protein | 633.64 | 1069.7 |
| MRET_2840 | protein YOP1 | 117.85 | 55.01 |
| MRET_2841 | protein-tyrosine phosphatase | 18.63 | 18.48 |
| MRET_2842 | exocyst complex component 7 | 18.17 | 23.68 |
| MRET_2843 | uncharacterized protein | 75.48 | 135.31 |
| MRET_2844 | verprolin, proline-rich actin-associated protein | 417.27 | 213.43 |
| MRET_2845 | translation initiation factor 4E | 80.17 | 159.83 |
| MRET_2846 | serine/threonine-protein kinase | 140.98 | 210.71 |

| | | | |
|---|---|---|---|
| MRET_2847 | porphobilinogen synthase | 521.68 | 556.63 |
| MRET_2848 | protein FET5 | 70.46 | 137.76 |
| MRET_2849 | protein CWC15 | 62.7 | 118.79 |
| MRET_2850 | PCI domain 2 protein | 20.26 | 47.8 |
| MRET_2851 | tRNA (guanine9-N1)-methyltransferase | 76.22 | 108.01 |
| MRET_2852 | serine/threonine-protein kinase | 193.7 | 319.95 |
| MRET_2853 | glycine dehydrogenase | 124.91 | 212.67 |
| MRET_2854 | hexokinase | 793.74 | 626.67 |
| MRET_2855 | UDP-galactopyranose mutase | 123.88 | 235.02 |
| MRET_2856 | large subunit ribosomal protein L44e | 573.36 | 959.43 |
| MRET_2857 | serine/threonine-protein kinase BUR1 | 105.12 | 118.08 |
| MRET_2858 | pre-mRNA-processing factor SLU7 | 513.84 | 468.91 |
| MRET_2859 | ribosome recycling factor | 827.58 | 319.47 |
| MRET_2860 | transcription initiation factor TFIID subunit 9B | 141.14 | 251.54 |
| MRET_2861 | coatomer subunit delta | 107.93 | 241.74 |
| MRET_2862 | 20S proteasome subunit beta 4 | 186.89 | 349.25 |
| MRET_2863 | kinetochore protein Mis12/MTW1 | 8.22 | 27.42 |
| MRET_2864 | splicing factor 1 | 61.01 | 81.71 |
| MRET_2865 | uncharacterized protein | 124.9 | 375.47 |
| MRET_2866 | diacylglycerol kinase (CTP) | 57.23 | 227.85 |
| MRET_2867 | uncharacterized protein | 15.09 | 55.84 |
| MRET_2868 | pre-mRNA-splicing factor clf1 | 10.17 | 34.2 |
| MRET_2869 | copper ion binding protein | 21.73 | 46.4 |
| MRET_2870 | eukaryotic sulfide quinone oxidoreductase | 434.16 | 319.71 |
| MRET_2871 | transporter | 32.88 | 58.36 |
| MRET_2872 | mitochondrial genome maintenance protein MGM101 | 27.86 | 74.95 |
| MRET_2873 | ATP-dependent RNA helicase DDX47/RRP3 | 18.65 | 34.89 |
| MRET_2874 | serine/threonine-protein kinase | 585.13 | 364.52 |
| MRET_2875 | mitochondrial inner membrane | 195.41 | 171.17 |
| MRET_2876 | amino acid permease | 71.08 | 52.39 |
| MRET_2877 | uncharacterized protein | 273.02 | 236.81 |
| MRET_2878 | uncharacterized protein | 157.66 | 66.85 |
| MRET_2879 | PH domain protein | 27.44 | 55.05 |
| MRET_2880 | small subunit ribosomal protein S12 | 117.38 | 140.27 |
| MRET_2881 | uncharacterized protein | 2.68 | 10.53 |
| MRET_2882 | elongation factor 1-beta | 206.68 | 375.24 |
| MRET_2883 | PHD finger domain protein | 97.07 | 136 |

| | | | |
|---|---|---|---|
| MRET_2884 | peptidyl-prolyl cis-trans isomerase | 1546.97 | 1538.58 |
| MRET_2885 | nuclear transport factor | 229.76 | 432.88 |
| MRET_2886 | aspartic-type endopeptidase CTSD | 2613.11 | 7402.24 |
| MRET_2887 | uncharacterized protein | 62.17 | 154.12 |
| MRET_2888 | transcription initiation factor TFIID subunit 2 | 520.79 | 193.75 |
| MRET_2889 | mediator of RNA polymerase II transcription subunit 7 | 56.12 | 115.12 |
| MRET_2890 | DNA replication ATP-dependent helicase DNA2 | 13.06 | 11.02 |
| MRET_2891 | amino acid transporter | 47.02 | 42.26 |
| MRET_2892 | amino acid transporter | 21.81 | 95.37 |
| MRET_2893 | amino acid transporter | 39.11 | 56.97 |
| MRET_2894 | amino acid transporter | 319.66 | 190.61 |
| MRET_2895 | SNW domain protein 1 | 54.29 | 461.32 |
| MRET_2896 | DNA polymerase eta | 0.14 | 43.88 |
| MRET_2897 | coproporphyrinogen III oxidase | 398.55 | 278.6 |
| MRET_2898 | conserved hypothetical protein | 67.32 | 28.43 |
| MRET_2899 | 3-phosphoinositide dependent protein kinase-1 | 65.99 | 56.83 |
| MRET_2900 | AHNAK nucleoprotein | 54.91 | 78.58 |
| MRET_2901 | HIV Tat-specific factor 1 | 264.36 | 372.57 |
| MRET_2902 | hexaprenyl-diphosphate synthase | 172.41 | 167.2 |
| MRET_2903 | peptidyl-tRNA hydrolase, PTH1 family | 37.56 | 16.98 |
| MRET_2904 | PAB1 binding protein | 166.37 | 174.45 |
| MRET_2905 | uncharacterized protein | 122.14 | 112.02 |
| MRET_2906 | mRNA splicing protein | 19.42 | 33.57 |
| MRET_2907 | kynureninase | 36.76 | 56.41 |
| MRET_2908 | D-tyrosyl-tRNA(Tyr) deacylase | 11.34 | 30.79 |
| MRET_2909 | RAD51-like protein 2 | 138.24 | 135.73 |
| MRET_2910 | myo-inositol-1(or 4)-monophosphatase | 78.37 | 78.92 |
| MRET_2911 | nuclear pore complex protein Nup54 | 42.53 | 53.42 |
| MRET_2912 | serine dehydratase beta chain | 198.16 | 148.34 |
| MRET_2913 | uncharacterized protein | 40.06 | 51.11 |
| MRET_2914 | dehydrogenase | 149.17 | 115.87 |
| MRET_2915 | uncharacterized protein | 175.29 | 110.05 |
| MRET_2916 | chromatin assembly factor 1 subunit B | 204.59 | 118.08 |
| MRET_2917 | UBX domain protein | 75.56 | 92.34 |
| MRET_2918 | cell division control protein | 77.53 | 46.74 |
| MRET_2919 | RNA exonuclease NGL2 | 15.09 | 27.53 |
| MRET_2920 | tRNA (cytidine32/guanosine34-2'-O)-methyltransferase | 13.24 | 23.52 |

| | | | |
|---|---|---|---|
| MRET_2921 | tRNA (adenine57-N1/adenine58-N1)-methyltransferase catalytic subunit | 42.27 | 62.05 |
| MRET_2922 | ribosome biogenesis protein MAK21 | 32.2 | 135.09 |
| MRET_2923 | transcription elongation factor S-II | 38.38 | 131.99 |
| MRET_2924 | signal peptidase complex subunit 1 | 108.81 | 151.24 |
| MRET_2925 | DnaJ domain protein | 23.81 | 55.43 |
| MRET_2926 | amyloid beta precursor protein binding protein 1 | 40.41 | 79.68 |
| MRET_2927 | nucleolar pre-ribosomal-associated protein 1 | 163.13 | 55.05 |
| MRET_2928 | uncharacterized protein | 75.34 | 45.67 |
| MRET_2929 | ubiquitin carboxyl-terminal hydrolase 22/27/51 | 101.04 | 82.71 |
| MRET_2930 | protein of unknown function (DUF2424) | 128.04 | 124.43 |
| MRET_2931 | phosphatidylinositol glycan, class C | 222.37 | 180.82 |
| MRET_2932 | zinc finger protein, C2H2 type | 1196.26 | 702.43 |
| MRET_2933 | monomeric glyoxalase I | 453.37 | 339.15 |
| MRET_2934 | tyrosyl-tRNA synthetase | 55.77 | 108.25 |
| MRET_2935 | endosomal cargo receptor (Erp3) | 287.12 | 402.26 |
| MRET_2936 | solute carrier family 29 (equilibrative nucleoside transporter), member 1/2/3 | 46.75 | 60.97 |
| MRET_2937 | NADH-ubiquinone oxidoreductase 9.5 kDa subunit | 74.09 | 143.82 |
| MRET_2938 | uncharacterized protein | 31.7 | 107.33 |
| MRET_2939 | uncharacterized protein | 51.9 | 85.53 |
| MRET_2940 | protein transport protein SEC13 | 195.38 | 344.28 |
| MRET_2941 | pre-mRNA-splicing factor ATP-dependent RNA helicase DHX15/PRP43 | 195.33 | 223.22 |
| MRET_2942 | nuclear pore complex protein Nup85 | 89.62 | 131.11 |
| MRET_2943 | large subunit ribosomal protein L23e | 143.81 | 477.54 |
| MRET_2944 | GTP cyclohydrolase IA | 40.82 | 109.2 |
| MRET_2945 | cytochrome b5 | 1411.17 | 776.87 |
| MRET_2946 | serine/threonine-protein kinase KIN1/2 | 166 | 150 |
| MRET_2947 | T-complex protein 1 subunit zeta | 56.81 | 94.22 |
| MRET_2948 | WD repeat protein 48 | 53.55 | 70.96 |
| MRET_2949 | actin cytoskeleton-regulatory complex protein PAN1 | 413.25 | 182.08 |
| MRET_2950 | non-classical export protein 1 | 136.25 | 164.13 |
| MRET_2951 | chitin synthase | 9.76 | 30.31 |
| MRET_2952 | phosphatase domain, paladin 1 | 285.58 | 96.9 |
| MRET_2953 | protein of unknown function (DUF1749) | 371.18 | 312.18 |
| MRET_2954 | large subunit ribosomal protein L17e | 249.66 | 258.58 |
| MRET_2955 | vesicle-associated membrane protein 4 | 944.45 | 790.66 |
| MRET_2956 | 2-oxoglutarate dehydrogenase E1 component | 2324.26 | 1051.01 |
| MRET_2957 | glutamate dehydrogenase | 26.99 | 41.11 |

| | | | |
|---|---|---|---|
| MRET_2958 | nucleolar protein 4 | 43.06 | 50.65 |
| MRET_2959 | uncharacterized protein | 559.95 | 432.4 |
| MRET_2960 | chromatin modification-related protein EAF6 | 328.91 | 220.31 |
| MRET_2961 | CTD kinase subunit gamma | 45.48 | 109.05 |
| MRET_2962 | cytosolic iron-sulfur protein assembly protein 1 | 42.35 | 80.74 |
| MRET_2963 | ubiquinone biosynthesis protein COQ9 | 49.05 | 121.98 |
| MRET_2964 | Ca2+-transporting ATPase | 304.95 | 215.51 |
| MRET_2965 | uncharacterized protein | 169.12 | 236.03 |
| MRET_2966 | fungal specific transcription factor domain protein | 95 | 89.38 |
| MRET_2967 | signal recognition particle subunit SRP68 | 25.87 | 47.23 |
| MRET_2968 | septum formation protein | 19.68 | 61.45 |
| MRET_2969 | small subunit ribosomal protein S23 | 21.8 | 66.84 |
| MRET_2970 | D-3-phosphoglycerate dehydrogenase/2-oxoglutarate reductase | 11.21 | 51.08 |
| MRET_2971 | cell division control protein 11 | 81.11 | 242.89 |
| MRET_2972 | chaperone | 27.39 | 72.1 |
| MRET_2973 | uncharacterized protein | 149.6 | 184.83 |
| MRET_2974 | uncharacterized protein | 769.63 | 947.77 |
| MRET_2975 | NADH-dependent flavin oxidoreductase | 1468.25 | 3602.99 |
| MRET_2976 | uncharacterized protein | 66.05 | 78.51 |
| MRET_2977 | parafibromin | 87.95 | 130.13 |
| MRET_2978 | histone H3-like centromeric protein A | 123.82 | 122.48 |
| MRET_2979 | GTI1/PAC2 family transcription factor | 79.7 | 53.47 |
| MRET_2980 | uncharacterized protein | 70.34 | 129.17 |
| MRET_2981 | mitochondrial ribosomal subunit S27 | 35.73 | 146.08 |
| MRET_2982 | chorismate synthase | 71.25 | 140.33 |
| MRET_2983 | ADP-ribosylation factor GTPase-activating protein 2/3 | 96.77 | 118.91 |
| MRET_2984 | CCR4-NOT transcription complex subunit 2 | 50.2 | 90.47 |
| MRET_2985 | uncharacterized protein | 350.14 | 348.46 |
| MRET_2986 | serum/glucocorticoid-regulated kinase 2 | 169.75 | 149.07 |
| MRET_2987 | DNA-directed RNA polymerases I and III subunit RPAC2 | 92.31 | 204.03 |
| MRET_2988 | quinone oxidoreductase | 309.76 | 243.51 |
| MRET_2989 | phosphoglucomutase | 34 | 40.24 |
| MRET_2990 | tRNA-splicing endonuclease subunit Sen34 | 37.55 | 58.55 |
| MRET_2991 | phosphatidylinositol 3-kinase | 6.11 | 13.94 |
| MRET_2992 | alpha-1,2-mannosyltransferase | 22.87 | 44.78 |
| MRET_2993 | electron transfer flavoprotein beta subunit | 144.31 | 181.1 |
| MRET_2994 | DNA polymerase alpha-associated DNA helicase A | 54.85 | 75.24 |

| | | | |
|---|---|---|---|
| MRET_2995 | neuronal calcium sensor 1 | 978.01 | 882.24 |
| MRET_2996 | anthranilate phosphoribosyltransferase | 15.97 | 36.51 |
| MRET_2997 | secretory pathway protein Ssp120 | 668.97 | 511.93 |
| MRET_2998 | ATP-binding cassette, subfamily B (MDR/TAP), member 1 | 220.68 | 204.45 |
| MRET_2999 | phosphoribosylglycinamide formyltransferase | 42.79 | 58.2 |
| MRET_3000 | mitofusin 2 | 1326.63 | 840.83 |
| MRET_3001 | uncharacterized protein | 122.51 | 189.12 |
| MRET_3002 | large subunit ribosomal protein L23Ae | 116.86 | 469.52 |
| MRET_3003 | uncharacterized protein | 51.91 | 74.5 |
| MRET_3004 | transcription factor | 30.89 | 41.85 |
| MRET_3005 | activating transcription factor 7 interacting protein | 8.05 | 43.24 |
| MRET_3006 | exosome complex component RRP42 | 53.14 | 67.95 |
| MRET_3007 | 3-isopropylmalate dehydratase | 34.23 | 47.58 |
| MRET_3008 | V-type H+-transporting ATPase 16kDa proteolipid subunit | 407.47 | 435.6 |
| MRET_3009 | mitochondrial-processing peptidase subunit beta | 2522.49 | 1644.49 |
| MRET_3010 | zinc finger protein, C3HC4 type (RING finger) | 99.62 | 79.07 |
| MRET_3011 | mediator of RNA polymerase II transcription subunit 6 | 16.05 | 21.15 |
| MRET_3012 | uncharacterized protein | 142 | 85.7 |
| MRET_3013 | uncharacterized protein | 59.99 | 63.87 |
| MRET_3014 | signal peptidase I | 66.92 | 108.75 |
| MRET_3015 | uncharacterized protein | 37.71 | 73.51 |
| MRET_3016 | poly(A) polymerase | 105.89 | 143.57 |
| MRET_3017 | S2P endopeptidase | 32.02 | 18.33 |
| MRET_3018 | epsin | 181.85 | 357.32 |
| MRET_3019 | 20S proteasome subunit alpha 5 | 157.48 | 170.61 |
| MRET_3020 | trafficking protein particle complex subunit 10 | 27.2 | 31.35 |
| MRET_3021 | RNA polymerase II-associated protein 1 | 53.08 | 37.04 |
| MRET_3022 | transcription initiation factor TFIID subunit 13 | 44.66 | 54.09 |
| MRET_3023 | glutathione reductase (NADPH) | 912.18 | 443.07 |
| MRET_3024 | Indole-diterpene biosynthesis protein PaxU | 76.98 | 101.53 |
| MRET_3025 | WD repeat protein | 31.66 | 56.69 |
| MRET_3026 | dephospho-CoA kinase | 348.92 | 235.48 |
| MRET_3027 | transcription initiation factor TFIID subunit 12 | 70.69 | 145.22 |
| MRET_3028 | release factor glutamine methyltransferase | 68.9 | 144.93 |
| MRET_3029 | U3 small nucleolar RNA-associated protein 19 | 79.84 | 122.81 |
| MRET_3030 | uncharacterized protein | 76.58 | 78.71 |
| MRET_3031 | U3 small nucleolar RNA-associated protein 18 | 81.09 | 107.25 |

| | | | |
|---|---|---|---|
| MRET_3032 | hydroxymethylbilane synthase | 56.35 | 65.76 |
| MRET_3033 | serine/threonine-protein kinase ATR | 10.74 | 17.5 |
| MRET_3034 | oxysterol-binding protein-related protein 9/10/11 | 81.03 | 191.54 |
| MRET_3035 | U3 small nucleolar RNA-associated protein 12 | 83.75 | 67.66 |
| MRET_3036 | tubulin-tyrosine ligase family protein | 156.07 | 147.94 |
| MRET_3037 | small subunit ribosomal protein S8e | 453.31 | 911.33 |
| MRET_3038 | bZIP transcription factor | 1373.51 | 1070.23 |
| MRET_3039 | GTPase-activating protein | 46.11 | 100.35 |
| MRET_3040 | oxysterol-binding protein-related protein 3/6/7 | 303.43 | 522 |
| MRET_3041 | cellular morphogenesis regulator DopA | 20.66 | 37.34 |
| MRET_3042 | exocyst complex component 8 | 122.17 | 71.83 |
| MRET_3043 | signal transduction protein Syg1 | 29.45 | 29.78 |
| MRET_3044 | small subunit ribosomal protein S15 | 72.03 | 119.32 |
| MRET_3045 | ubiquitin-protein ligase E3 D | 180.52 | 135.79 |
| MRET_3046 | protein SHQ1 | 31.17 | 51.93 |
| MRET_3047 | WD repeat protein | 331.13 | 192.24 |
| MRET_3048 | chitinase | 195.24 | 157.42 |
| MRET_3049 | putative integral membrane protein that interacts with Rpp0p | 21.71 | 37.56 |
| MRET_3050 | signal recognition particle subunit SRP14 | 67.33 | 118.69 |
| MRET_3051 | auxin efflux carrier | 69.47 | 81.72 |
| MRET_3052 | AP-2 complex-associated kinase | 23.83 | 28.71 |
| MRET_3053 | ribosomal large subunit biogenesis | 55.5 | 62.54 |
| MRET_3054 | structure-specific endonuclease subunit SLX1 | 64.2 | 37.5 |
| MRET_3055 | conserved hypothetical protein | 573.17 | 608.66 |
| MRET_3056 | ER membrane protein complex subunit 2 | 116.45 | 207.93 |
| MRET_3057 | serine/threonine-protein phosphatase PP1 catalytic subunit | 603.52 | 1150.59 |
| MRET_3058 | Wwm1-ww domain containing protein interacting with metacaspase | 1436.23 | 903.16 |
| MRET_3059 | polyribonucleotide 5'-hydroxyl-kinase | 55.25 | 96.07 |
| MRET_3060 | CD2 antigen cytoplasmic tail-binding protein 2 | 10.26 | 21.54 |
| MRET_3061 | NADH dehydrogenase (ubiquinone) 1 alpha subcomplex subunit 2 | 1084.54 | 839.88 |
| MRET_3062 | transketolase | 391.69 | 286.53 |
| MRET_3063 | 2-isopropylmalate synthase | 53.49 | 67.32 |
| MRET_3064 | uncharacterized protein | 300.37 | 247.39 |
| MRET_3065 | SET domain protein | 68.99 | 151.92 |
| MRET_3066 | peroxin-19 | 152.17 | 195.67 |
| MRET_3067 | palmitoyltransferase ZDHHC6 | 104.69 | 79.01 |
| MRET_3068 | myosin I | 104.98 | 69.15 |

| | | | |
|---|---|---|---|
| MRET_3069 | acyl-protein thioesterase responsible for depalmitoylation of Gpa1p | 61.26 | 85.04 |
| MRET_3070 | fungal Zn(2)-Cys(6) binuclear cluster domain protein | 17.56 | 39.37 |
| MRET_3071 | uncharacterized protein | 97.69 | 198.72 |
| MRET_3072 | large subunit ribosomal protein L34e | 223.91 | 715.03 |
| MRET_3073 | large subunit ribosomal protein L6 | 57.73 | 120.35 |
| MRET_3074 | E3 ubiquitin-protein ligase synoviolin | 666.67 | 397.87 |
| MRET_3075 | NAD+ kinase | 59.04 | 58.9 |
| MRET_3076 | zinc finger protein | 485.5 | 444.11 |
| MRET_3077 | uncharacterized protein | 195.75 | 216.27 |
| MRET_3078 | uncharacterized protein | 35.91 | 38.11 |
| MRET_3079 | uncharacterized protein | 76.93 | 228.8 |
| MRET_3080 | phosducin family | 81.55 | 220.27 |
| MRET_3081 | uncharacterized protein | 118.51 | 305.45 |
| MRET_3082 | glucan synthesis regulatory protein | 158.07 | 109.71 |
| MRET_3083 | mannosyl-oligosaccharide glucosidase | 98.45 | 86.65 |
| MRET_3084 | dynamitin | 15.06 | 13.82 |
| MRET_3085 | RNA-binding protein | 22.14 | 39.31 |
| MRET_3086 | RAM signalling pathway protein domain protein | 47.61 | 22.83 |
| MRET_3087 | transcription elongation factor SPT6 | 33.56 | 56.24 |
| MRET_3088 | phospholipase C | 18.99 | 81.7 |
| MRET_3089 | histone-lysine N-methyltransferase SUV420H | 335 | 169.95 |
| MRET_3090 | 26S proteasome regulatory subunit T3 | 81.79 | 98.07 |
| MRET_3091 | pyrroline-5-carboxylate reductase | 56 | 63.58 |
| MRET_3092 | uncharacterized protein | 12.45 | 16.49 |
| MRET_3093 | endosomal peripheral membrane protein | 20.45 | 21.52 |
| MRET_3094 | cytochrome p450 | 41.65 | 73.71 |
| MRET_3095 | Yqey-like protein | 33.49 | 27.89 |
| MRET_3096 | F-box and WD-40 domain protein MET30 | 517.83 | 552.99 |
| MRET_3097 | large subunit ribosomal protein L26e | 104.79 | 344.67 |
| MRET_3098 | GTP-binding protein SAR1 | 290.34 | 606.68 |
| MRET_3099 | glycogen synthase kinase 3 beta | 390.87 | 415.51 |
| MRET_3100 | urease accessory protein | 88.89 | 157.35 |
| MRET_3101 | U3 small nucleolar RNA-associated protein 24 | 183.66 | 311.2 |
| MRET_3102 | translation initiation factor 1 | 520.92 | 1159.73 |
| MRET_3103 | conserved hypothetical protein | 79.45 | 107.62 |
| MRET_3104 | uncharacterized protein | 11.64 | 49.07 |
| MRET_3105 | alpha-1,3/alpha-1,6-mannosyltransferase | 87.1 | 51.87 |

| | | | |
|---|---|---|---|
| MRET_3106 | mitochondrial fusion and transport protein UGO1 | 156.98 | 124.04 |
| MRET_3107 | multisite-specific tRNA:(cytosine-C5)-methyltransferase | 242.1 | 140.9 |
| MRET_3108 | molecular chaperone DnaJ | 1416.81 | 1074.07 |
| MRET_3109 | low molecular weight phosphotyrosine protein phosphatase | 57.67 | 130.41 |
| MRET_3110 | Sec1 family domain protein 1 | 42.55 | 77.89 |
| MRET_3111 | acyl-CoA thioesterase 8 | 99.37 | 121.88 |
| MRET_3112 | JmjC domain, hydroxylase | 216.05 | 125.84 |
| MRET_3113 | GYF domain protein | 25.3 | 29.34 |
| MRET_3114 | histone deacetylase 1/2 | 78.97 | 147.79 |
| MRET_3115 | V-type H+-transporting ATPase subunit F | 103.82 | 156.31 |
| MRET_3116 | ubiquitin C | 5358.25 | 4367.52 |
| MRET_3117 | peptidyl-tRNA hydrolase, PTH2 family | 72.98 | 158.13 |
| MRET_3118 | mRNA-decapping enzyme subunit 2 | 96.44 | 161.16 |
| MRET_3119 | biotin synthase | 428.49 | 389.08 |
| MRET_3120 | uncharacterized protein | 200.61 | 109.2 |
| MRET_3121 | dynein heavy chain 1, cytosolic | 59.68 | 40.69 |
| MRET_3122 | component of the septin ring that is required for cytokinesis | 63.99 | 166.09 |
| MRET_3123 | U2 small nuclear ribonucleoprotein A' | 11.01 | 39.76 |
| MRET_3124 | dynein heavy chain 1, cytosolic | 19.57 | 54.74 |
| MRET_3125 | uncharacterized protein | 25.84 | 40.36 |
| MRET_3126 | CCAAT-binding transcription factor subunit HAPB | 186.89 | 239.14 |
| MRET_3127 | vacuolar protein sorting-associated protein 13A/C | 285.54 | 274.52 |
| MRET_3128 | large subunit ribosomal protein L47 | 49.49 | 126.66 |
| MRET_3129 | metal iron transporter | 40.13 | 85.97 |
| MRET_3130 | AHNAK nucleoprotein | 20.12 | 82.91 |
| MRET_3131 | polyketide synthase | 54.84 | 143.57 |
| MRET_3132 | nuclear pore complex protein Nup98-Nup96 | 92.24 | 78.39 |
| MRET_3133 | uncharacterized protein | 361.99 | 196.27 |
| MRET_3134 | molecular chaperone HscB | 69.92 | 71.18 |
| MRET_3135 | protein phosphatase PTC7 | 519 | 404.68 |
| MRET_3136 | Kelch repeats protein | 69.87 | 148.57 |
| MRET_3137 | endoplasmic reticulum-golgi intermediate compartment protein 2 | 316.34 | 365.35 |
| MRET_3138 | U6 snRNA-associated Sm-like protein LSm7 | 83.81 | 110.69 |
| MRET_3139 | protein transport protein DSL1/ZW10 | 36.39 | 42.44 |
| MRET_3140 | serine/threonine-protein phosphatase 2A regulatory subunit B | 66.95 | 132.42 |
| MRET_3141 | prolyl oligopeptidase | 78.21 | 60.37 |
| MRET_3142 | solute carrier family 31 (copper transporter), member 1 | 525.48 | 213.66 |

| | | | |
|---|---|---|---|
| MRET_3143 | F-box and WD-40 domain protein CDC4 | 38.09 | 33.51 |
| MRET_3144 | DnaJ homolog subfamily C member 7 | 18.38 | 47.92 |
| MRET_3145 | tRNA pseudouridine38-40 synthase | 88.2 | 85.57 |
| MRET_3146 | RNA polymerase-associated protein RTF1 | 43.25 | 46.75 |
| MRET_3147 | actin-related protein 2 | 740.44 | 686.24 |
| MRET_3148 | aarF domain kinase | 75.61 | 143.49 |
| MRET_3149 | cytochrome c oxidase subunit 5a | 550.33 | 601.09 |
| MRET_3150 | E3 ubiquitin-protein ligase SHPRH | 91.54 | 45.99 |
| MRET_3151 | Myb-like DNA-binding domain protein | 154.66 | 122.56 |
| MRET_3152 | RNA recognition motif domain protein | 8153.67 | 2236.21 |
| MRET_3153 | Ras-related protein Rab-5C | 715.52 | 417.66 |
| MRET_3154 | pantoate--beta-alanine ligase | 156.69 | 117.81 |
| MRET_3155 | methionyl aminopeptidase | 286.73 | 364.1 |
| MRET_3156 | cytochrome c oxidase subunit 7 | 197.36 | 254.07 |
| MRET_3157 | oxidoreductase which may be involved in DNA replication (By similarity) | 49.3 | 53.5 |
| MRET_3158 | elongation factor 1-gamma | 236.18 | 385.39 |
| MRET_3159 | 20S proteasome subunit alpha 3 | 81.7 | 139.94 |
| MRET_3160 | small nuclear ribonucleoprotein D3 | 59.73 | 170.59 |
| MRET_3161 | high-mobility group non-histone chromatin protein | 313.58 | 523.01 |
| MRET_3162 | solute carrier family 25 (mitochondrial citrate transporter), member 1 | 19.19 | 32.81 |
| MRET_3163 | chloride channel | 103.98 | 60.28 |
| MRET_3164 | multiple RNA-binding domain protein 1 | 153.87 | 143.65 |
| MRET_3165 | mediator of RNA polymerase II transcription subunit 10 | 46.97 | 38.65 |
| MRET_3166 | 26S proteasome regulatory subunit N3 | 51.56 | 69.19 |
| MRET_3167 | uncharacterized protein | 163.42 | 219.26 |
| MRET_3168 | cysteine and glycine-rich protein | 133.18 | 102.74 |
| MRET_3169 | choline-phosphate cytidylyltransferase | 8.35 | 54.63 |
| MRET_3170 | NADH dehydrogenase (ubiquinone) 1 alpha subcomplex subunit 6 | 175.38 | 278.94 |
| MRET_3171 | serine/threonine-protein phosphatase 6 catalytic subunit | 33.66 | 72.84 |
| MRET_3172 | uncharacterized protein | 59.76 | 63.54 |
| MRET_3173 | dihydrolipoamide dehydrogenase | 3758.82 | 1658.61 |
| MRET_3174 | mRNA m6A methyltransferase | 158.68 | 95.44 |
| MRET_3175 | BAP31 domain protein | 877.06 | 402.62 |
| MRET_3176 | DUF89 domain protein | 270.97 | 174.99 |
| MRET_3177 | NGG1 interacting factor | 253.75 | 167.62 |
| MRET_3178 | DNA-directed RNA polymerase III subunit RPC2 | 92.82 | 38.81 |
| MRET_3179 | protein of unknown function (DUF3128) | 392.04 | 880.58 |

| | | | |
|---|---|---|---|
| MRET_3180 | Mago binding protein | 51.1 | 174.85 |
| MRET_3181 | large subunit ribosomal protein L28e | 96.17 | 339.27 |
| MRET_3182 | prefoldin subunit | 15.94 | 35.16 |
| MRET_3183 | translation initiation factor IF-2 | 101.89 | 58.23 |
| MRET_3184 | ubiquitin-conjugating enzyme E2 G1 | 224.49 | 185.5 |
| MRET_3185 | CCR4-NOT transcription complex subunit 7/8 | 45.64 | 80.95 |
| MRET_3186 | 26S proteasome regulatory subunit N10 | 89.38 | 155.17 |
| MRET_3187 | pheromone a factor receptor | 124.4 | 0 |
| MRET_3188 | exosome complex component RRP41 | 64.09 | 117.05 |
| MRET_3189 | glyceraldehyde 3-phosphate dehydrogenase | 3328.35 | 1924.7 |
| MRET_3190 | cell division control protein 24 | 194.92 | 115.98 |
| MRET_3191 | protein disulfide-isomerase A1 | 1107.8 | 802.57 |
| MRET_3192 | L-gulonolactone oxidase | 140.36 | 66.72 |
| MRET_3193 | 26S proteasome regulatory subunit N9 | 30.29 | 37.78 |
| MRET_3194 | phosphatidylethanolamine/phosphatidyl-N-methylethanolamine N-methyltransferase | 142.41 | 100.96 |
| MRET_3195 | N-acetylglucosaminylphosphatidylinositol deacetylase | 116.07 | 94.41 |
| MRET_3196 | serine/threonine-protein kinase | 262.99 | 238.08 |
| MRET_3197 | uncharacterized protein | 62.21 | 73.64 |
| MRET_3198 | cytochrome c oxidase subunit | 496.95 | 230.4 |
| MRET_3199 | Fes/CIP4, and EFC/F-BAR homology domain protein | 209.74 | 98.39 |
| MRET_3200 | p38 MAP kinase | 223.81 | 50.52 |
| MRET_3201 | calcium channel MID1 | 111.29 | 65.23 |
| MRET_3202 | uncharacterized protein | 112.79 | 64.35 |
| MRET_3203 | uncharacterized protein | 264.53 | 154.55 |
| MRET_3204 | oligosaccharide translocation protein RFT1 | 61.22 | 61.13 |
| MRET_3205 | C2 domain protein | 80.2 | 45.64 |
| MRET_3206 | RNA-binding protein Musashi | 2719.81 | 2276.76 |
| MRET_3207 | DNA-directed RNA polymerase II subunit RPB11 | 151.98 | 203.47 |
| MRET_3208 | peroxin-14 | 39.32 | 51.84 |
| MRET_3209 | vacuolar protein sorting-associated protein 45 | 18.72 | 15.63 |
| MRET_3210 | amino acid transporter | 181 | 91.32 |
| MRET_3211 | solute carrier family 25 (mitochondrial 2-oxodicarboxylate transporter), member 21 | 573.47 | 369.39 |
| MRET_3212 | serine/threonine-protein kinase | 56.52 | 26.09 |
| MRET_3213 | cytoplasmic tRNA 2-thiolation protein 2 | 82.47 | 43.2 |
| MRET_3214 | PAB-dependent poly(A)-specific ribonuclease subunit 3 | 104.4 | 43.88 |
| MRET_3215 | terminal uridylyltransferase | 31.37 | 20.96 |
| MRET_3216 | uncharacterized protein | 532.6 | 334.67 |

| | | | |
|---|---|---|---|
| MRET_3217 | oxalate---CoA ligase | 45.61 | 13.66 |
| MRET_3218 | dual specificity phosphatase | 379.5 | 185.96 |
| MRET_3219 | CUE domain protein | 138.52 | 137.04 |
| MRET_3220 | transcription factor | 140.27 | 139.06 |
| MRET_3221 | ubiquitin carboxyl-terminal hydrolase 14 | 102.34 | 139.53 |
| MRET_3222 | nucleolar complex protein 2 | 129.05 | 118.6 |
| MRET_3223 | WD repeat protein | 212.46 | 147.71 |
| MRET_3224 | drebrin-like protein | 418.19 | 376.54 |
| MRET_3225 | conserved hypothetical protein | 56.23 | 63.12 |
| MRET_3226 | uncharacterized protein | 6.13 | 4.69 |
| MRET_3227 | uncharacterized protein | 21.18 | 27.65 |
| MRET_3228 | chromatin modification-related protein YNG2 | 531.78 | 272.96 |
| MRET_3229 | conserved hypothetical protein | 158.14 | 122.19 |
| MRET_3230 | FAD binding domain protein | 25.3 | 42.73 |
| MRET_3231 | purine nucleoside permease | 5793.57 | 8519.2 |
| MRET_3232 | ADP-ribose pyrophosphatase | 38.24 | 101.1 |
| MRET_3233 | sterol 14-demethylase | 117.97 | 335.74 |
| MRET_3234 | homoserine dehydrogenase | 113.29 | 118.33 |
| MRET_3235 | uncharacterized protein | 270.8 | 130.93 |
| MRET_3236 | 20S proteasome subunit beta 5 | 172.1 | 154.34 |
| MRET_3237 | polyphosphoinositide phosphatase | 81.84 | 44.04 |
| MRET_3238 | delta24(24(1))-sterol reductase | 62.92 | 103.13 |
| MRET_3239 | large subunit ribosomal protein L7/L12 | 69.65 | 208.52 |
| MRET_3240 | uncharacterized protein | 15.4 | 24.21 |
| MRET_3241 | Cu+-exporting ATPase | 93.22 | 50.75 |
| MRET_3242 | uncharacterized protein | 123.89 | 62.31 |
| MRET_3243 | uncharacterized protein | 83.8 | 118.7 |
| MRET_3244 | MSF1 domain protein | 7.13 | 10.51 |
| MRET_3245 | signal recognition particle subunit SRP54 | 153.2 | 118.23 |
| MRET_3246 | phosphatidylinositol glycan, class V | 69.75 | 56.5 |
| MRET_3247 | protein SYS1 | 31.8 | 40.81 |
| MRET_3248 | AP-2 complex subunit sigma-1 | 37.59 | 43.91 |
| MRET_3249 | required for respiratory growth protein 9, mitochondrial | 550.05 | 316.44 |
| MRET_3250 | NADH-ubiquinone oxidoreductase 21 kDa subunit | 182.63 | 290.83 |
| MRET_3251 | translocation protein SEC72 | 20.2 | 52.5 |
| MRET_3252 | ATP synthase mitochondrial F1 complex assembly factor 2 | 73.16 | 72.53 |
| MRET_3253 | integral peroxisomal membrane peroxin | 163.98 | 94.78 |

| | | | |
|---|---|---|---|
| MRET_3254 | phosphatidylinositol glycan, class S | 36.92 | 42.17 |
| MRET_3255 | uncharacterized protein | 2.55 | 8.61 |
| MRET_3256 | uncharacterized protein | 97.23 | 162.53 |
| MRET_3257 | stress responsive A/B barrel domain protein | 655.6 | 805.93 |
| MRET_3258 | gem associated protein 2 | 26.56 | 50.02 |
| MRET_3259 | ribosomal RNA-processing protein 1 | 292.02 | 231.13 |
| MRET_3260 | solute carrier family 25 (mitochondrial S-adenosylmethionine transporter), member 26 | 180.03 | 89.98 |
| MRET_3261 | aspartyl-tRNA(Asn)/glutamyl-tRNA(Gln) amidotransferase subunit A | 47.26 | 34.48 |
| MRET_3262 | SDE2 telomere maintenance homolog | 367.33 | 119.71 |
| MRET_3263 | uncharacterized protein | 116.09 | 37.77 |
| MRET_3264 | tetratricopeptide repeat domain protein | 849.94 | 944.06 |
| MRET_3265 | TFIIH basal transcription factor complex TTD-A subunit | 122.71 | 93.87 |
| MRET_3266 | thiamine pyrophosphokinase | 77.23 | 62.82 |
| MRET_3267 | Cys-Gly metallodipeptidase DUG1 | 82.15 | 114.46 |
| MRET_3268 | proline-rich, actin-associated protein Vrp1 | 28.35 | 24.57 |
| MRET_3269 | dolichyldiphosphatase | 65.68 | 69.39 |
| MRET_3270 | membrane associated protein-like protein | 76.32 | 49.4 |
| MRET_3271 | solute carrier family 25 (mitochondrial carrier protein), member 16 | 77.71 | 111.28 |
| MRET_3272 | DNA helicase II/ATP-dependent DNA helicase PcrA | 12.74 | 14.42 |
| MRET_3273 | glycerol-3-phosphate O-acyltransferase/dihydroxyacetone phosphate acyltransferase | 85.9 | 80.07 |
| MRET_3274 | acylphosphatase | 874.3 | 558.17 |
| MRET_3275 | DASH complex subunit DAM1 | 133.97 | 74.75 |
| MRET_3276 | uncharacterized protein | 268.07 | 215.35 |
| MRET_3277 | uncharacterized protein | 54.89 | 43.3 |
| MRET_3278 | short-chain dehydrogenase | 61.28 | 50.49 |
| MRET_3279 | uncharacterized protein | 33.7 | 32.31 |
| MRET_3280 | protoheme IX farnesyltransferase, mitochondrial | 196.39 | 117.78 |
| MRET_3281 | CDK-activating kinase assembly factor MAT1 | 21.82 | 28.06 |
| MRET_3282 | uncharacterized protein | 328.44 | 164.25 |
| MRET_3283 | uncharacterized protein | 250.85 | 251.87 |
| MRET_3284 | coronin-1B/1C/6 | 710.35 | 413.67 |
| MRET_3285 | uncharacterized protein | 37.45 | 37.86 |
| MRET_3286 | diphthine methyl ester synthase | 41.81 | 55.72 |
| MRET_3287 | G patch domain protein 1 | 100.24 | 42.94 |
| MRET_3288 | uncharacterized protein | 199.83 | 82.88 |
| MRET_3289 | UPF0160 domain protein MYG1 | 100.14 | 106.72 |
| MRET_3290 | zinc finger protein | 278.93 | 289.34 |

| | | | |
|---|---|---|---|
| MRET_3291 | urease accessory protein | 7.03 | 9.64 |
| MRET_3292 | MFS transporter | 24.66 | 38.87 |
| MRET_3293 | vacuole protein | 97.57 | 71.66 |
| MRET_3294 | uncharacterized protein | 23.78 | 36.99 |
| MRET_3295 | mitotic spindle assembly checkpoint protein MAD2B | 36.06 | 72.38 |
| MRET_3296 | uncharacterized protein | 33.11 | 96.28 |
| MRET_3297 | ubiquitin thioesterase protein OTUB1 | 52.29 | 143.26 |
| MRET_3298 | DNA-directed RNA polymerases I and III subunit RPAC1 | 33.7 | 68.76 |
| MRET_3299 | minichromosome maintenance protein 10 | 8.14 | 11.26 |
| MRET_3300 | protein phosphatase type 1 complex subunit Hex2 Reg1 | 56.5 | 81.01 |
| MRET_3301 | conserved hypothetical protein | 92.72 | 26.24 |
| MRET_3302 | ATP-dependent RNA helicase DDX56/DBP9 | 116.97 | 59.51 |
| MRET_3303 | transmembrane component | 16.54 | 6.05 |
| MRET_3304 | exopolyphosphatase | 60.17 | 34.43 |
| MRET_3305 | peroxin-5 | 358.67 | 194.69 |
| MRET_3306 | peptidyl-prolyl cis-trans isomerase-like 2 | 97.29 | 52.39 |
| MRET_3307 | uncharacterized protein | 43.67 | 43.38 |
| MRET_3308 | beta receptor associated protein 1 | 87.01 | 39.64 |
| MRET_3309 | tRNA nucleotidyltransferase | 74.2 | 40.24 |
| MRET_3310 | mitogen-activated protein kinase 1/3 | 73.87 | 66.27 |
| MRET_3311 | solute carrier family 35 (UDP-galactose transporter), member B1 | 149.66 | 137.95 |
| MRET_3312 | small subunit ribosomal protein S29 | 376.86 | 236.98 |
| MRET_3313 | maintenance of ploidy protein MOB2 | 28.8 | 31.52 |
| MRET_3314 | uncharacterized protein | 52.09 | 71.19 |
| MRET_3315 | peptidyl-prolyl cis-trans isomerase-like 4 | 19.31 | 26.08 |
| MRET_3316 | elongation factor G | 124.51 | 94.93 |
| MRET_3317 | DUF866 domain protein | 127.47 | 69.77 |
| MRET_3318 | CDP-diacylglycerol---serine O-phosphatidyltransferase | 672.46 | 311.59 |
| MRET_3319 | VHS domain protein | 109.53 | 126.12 |
| MRET_3320 | serine/threonine-protein phosphatase 4 catalytic subunit | 117.81 | 81.26 |
| MRET_3321 | (R)-2-hydroxyglutarate---pyruvate transhydrogenase | 78.99 | 99.49 |
| MRET_3322 | mitochondrial hypoxia responsive domain protein | 3876.77 | 3141.9 |
| MRET_3323 | phosphodiesterase | 388.64 | 302.04 |
| MRET_3324 | nucleolar protein 53 | 40.17 | 77.64 |
| MRET_3325 | oligosaccharyltransferase complex subunit gamma | 94.22 | 89.62 |
| MRET_3326 | uncharacterized protein | 512.7 | 270.23 |
| MRET_3327 | mitochondrial 54S ribosomal protein YmL47 | 212.34 | 274.34 |

| | | | |
|---|---|---|---|
| MRET_3328 | golgi vesicular membrane trafficking protein | 23.95 | 18.2 |
| MRET_3329 | orotate phosphoribosyltransferase | 11.6 | 34.03 |
| MRET_3330 | NACHT domain protein | 82.45 | 44.09 |
| MRET_3331 | N-alpha-acetyltransferase 38, NatC auxiliary subunit | 353.88 | 215.4 |
| MRET_3332 | HIT family protein 1 | 385.85 | 307.79 |
| MRET_3333 | caffeine-induced death protein 2 | 204.66 | 182.54 |
| MRET_3334 | glycerol-3-phosphate dehydrogenase (NAD+) | 181.53 | 189.71 |
| MRET_3335 | uncharacterized protein | 208.36 | 180.56 |
| MRET_3336 | etoposide-induced 2.4 mRNA | 18.62 | 31.4 |
| MRET_3337 | uncharacterized protein | 0.34 | 345.83 |
| MRET_3338 | nudix family hydrolase | 607.85 | 449.15 |
| MRET_3339 | U1 small nuclear ribonucleoprotein C | 40.48 | 47.56 |
| MRET_3340 | ATP-dependent RNA helicase DDX49/DBP8 | 35.68 | 45.33 |
| MRET_3341 | cell wall biogenesis protein | 87.42 | 68.83 |
| MRET_3342 | acetylornithine aminotransferase | 236.94 | 164.27 |
| MRET_3343 | MOB kinase activator 1 | 342.38 | 1279.53 |
| MRET_3344 | acid phosphatase | 212.61 | 108.54 |
| MRET_3345 | amidophosphoribosyltransferase | 46.45 | 56.95 |
| MRET_3346 | solute carrier family 25, member 33/36 | 276.87 | 195.54 |
| MRET_3347 | tyrosine-protein phosphatase OCA1 | 129.94 | 159.77 |
| MRET_3348 | aldo-keto reductase | 148.92 | 168.93 |
| MRET_3349 | chitin synthase | 114.1 | 84.37 |
| MRET_3350 | DASH complex subunit ASK1 | 307.58 | 228.65 |
| MRET_3351 | OTU domain protein 6 | 155.54 | 156.34 |
| MRET_3352 | chromatin modification-related protein | 202.73 | 153.93 |
| MRET_3353 | uncharacterized protein | 29.65 | 64.44 |
| MRET_3354 | dynactin 6 | 14.67 | 28.54 |
| MRET_3355 | uncharacterized protein | 2.14 | 6.78 |
| MRET_3356 | phospholipase C | 122.21 | 125.22 |
| MRET_3357 | multidrug transporter | 38.86 | 81.36 |
| MRET_3358 | uncharacterized protein | 61.13 | 69 |
| MRET_3359 | uncharacterized protein | 362.24 | 154.27 |
| MRET_3360 | uncharacterized protein | 16.84 | 15.59 |
| MRET_3361 | uncharacterized protein | 108.59 | 216.82 |
| MRET_3362 | uncharacterized protein | 26.13 | 53.97 |
| MRET_3363 | sporulation protein RMD1 | 79.59 | 134.04 |
| MRET_3364 | GTP-binding protein | 27 | 61.7 |

| | | | |
|---|---|---|---|
| MRET_3365 | alpha-1,3-mannosyltransferase | 57.71 | 101.41 |
| MRET_3366 | DUF218 domain protein | 28.74 | 69.55 |
| MRET_3367 | transmembrane protein 33 | 226.79 | 328.95 |
| MRET_3368 | RAD51-like protein | 15.16 | 33.49 |
| MRET_3369 | DNA-directed RNA polymerase II subunit RPB1 | 1034.89 | 979.64 |
| MRET_3370 | ubiquitin binding protein | 101.48 | 179.29 |
| MRET_3371 | conserved hypothetical protein | 19.79 | 50.51 |
| MRET_3372 | uncharacterized protein | 17.79 | 43.16 |
| MRET_3373 | uncharacterized protein | 157.61 | 150.78 |
| MRET_3374 | protein of unknown function (DUF2424) | 71.15 | 188.01 |
| MRET_3375 | iron-sulfur cluster assembly protein ISA2 | 2513.78 | 1477.24 |
| MRET_3376 | nascent polypeptide-associated complex subunit alpha | 82.53 | 185.83 |
| MRET_3377 | 20S proteasome subunit alpha 4 | 268.5 | 524.37 |
| MRET_3378 | SNARE associated golgi protein | 27.55 | 129.72 |
| MRET_3379 | pumilio-family RNA binding repeat protein | 81.41 | 104.07 |
| MRET_3380 | exosome complex component RRP43 | 35.99 | 60.13 |
| MRET_3381 | phosphomevalonate kinase | 15.82 | 21.34 |
| MRET_3382 | ATP-binding cassette, subfamily F, member 3 | 243.56 | 165.98 |
| MRET_3383 | ATP-binding cassette, subfamily F, member 3 | 96.81 | 53.77 |
| MRET_3384 | uncharacterized protein | 0 | 0 |
| MRET_3385 | lactate 2-monooxygenase | 200.37 | 177.16 |
| MRET_3386 | ATP-binding cassette, subfamily F, member 3 | 91.72 | 49.9 |
| MRET_3387 | uncharacterized protein | 0 | 0 |
| MRET_3388 | lactate 2-monooxygenase | 199.41 | 177.8 |
| MRET_3389 | transcription initiation factor TFIIH subunit 1 | 42.84 | 54.13 |
| MRET_3390 | cell division cycle protein 37 | 37.08 | 40.94 |
| MRET_3391 | protein LSM14 | 1148.69 | 1391.15 |
| MRET_3392 | fungal Zn(2)-Cys(6) binuclear cluster domain protein | 98.18 | 128.59 |
| MRET_3393 | vacuolar protein sorting-associated protein 29 | 93.99 | 199.22 |
| MRET_3394 | uncharacterized protein | 81.67 | 120.45 |
| MRET_3395 | lipoyl(octanoyl) transferase | 15.63 | 32.31 |
| MRET_3396 | protein of unknown function (DUF3712) | 213.37 | 410.13 |
| MRET_3397 | actin-related protein 5 | 117.36 | 205.25 |
| MRET_3398 | myo-inositol-1-phosphate synthase | 645.95 | 439.24 |
| MRET_3399 | PLP dependent protein | 50.19 | 95.25 |
| MRET_3400 | ribosome maturation protein SDO1 | 1209.59 | 933.46 |
| MRET_3401 | sulfate permease, SulP family | 124.6 | 121.61 |

| | | | |
|---|---|---|---|
| MRET_3402 | uncharacterized protein | 29.12 | 73.29 |
| MRET_3403 | translation initiation factor 4A | 1339.27 | 1213.3 |
| MRET_3404 | plasma membrane sulfite pump involved in sulfite metabolism | 24.82 | 41.28 |
| MRET_3405 | dolichyl-phosphate-mannose-protein mannosyltransferase | 22.73 | 40.46 |
| MRET_3406 | target of rapamycin complex subunit LST8 | 26.76 | 61.55 |
| MRET_3407 | large subunit ribosomal protein L44 | 49.62 | 149.24 |
| MRET_3408 | signal recognition particle receptor subunit alpha | 44.54 | 109.97 |
| MRET_3409 | uncharacterized protein | 26.37 | 73.68 |
| MRET_3410 | solute carrier family 9 (sodium/hydrogen exchanger), member 6/7 | 42.83 | 101.49 |
| MRET_3411 | proline iminopeptidase | 65.83 | 127.4 |
| MRET_3412 | SWI/SNF related-matrix-associated actin-dependent regulator of chromatin subfamily C | 75.55 | 104.63 |
| MRET_3413 | uncharacterized protein | 526.47 | 499.82 |
| MRET_3414 | Rho GTPase activating protein | 159.4 | 83.55 |
| MRET_3415 | glycoside hydrolase family 5 protein | 11.58 | 224.57 |
| MRET_3416 | uncharacterized protein | 11.36 | 32.46 |
| MRET_3417 | cellulase (glycosyl hydrolase family 5) | 68.97 | 124.94 |
| MRET_3418 | uncharacterized protein | 5.83 | 18.75 |
| MRET_3419 | pre-mRNA-splicing factor SYF1 | 14.57 | 37.09 |
| MRET_3420 | ubiquitin carboxyl-terminal hydrolase 10 | 9.94 | 26.93 |
| MRET_3421 | R3H domain protein | 46.95 | 95.49 |
| MRET_3422 | WD repeat protein involved in ribosome biogenesis | 19.54 | 69.17 |
| MRET_3423 | transcription factor SPN1 | 54.05 | 112.88 |
| MRET_3424 | small subunit ribosomal protein S9e | 145.13 | 398.23 |
| MRET_3425 | large subunit ribosomal protein L21e | 500.38 | 797.6 |
| MRET_3426 | carbohydrate esterase family 9 protein | 194.95 | 125.35 |
| MRET_3427 | ankyrin repeat domain protein | 100.04 | 169.96 |
| MRET_3428 | Sds3-like protein | 43.32 | 61.87 |
| MRET_3429 | pyruvate dehydrogenase kinase 2/3/4 | 63.52 | 133.16 |
| MRET_3430 | F-type H+-transporting ATPase subunit b | 956.73 | 1311.37 |
| MRET_3431 | cyclin-dependent kinase 7 | 77.95 | 78.97 |
| MRET_3432 | Rab5 GDP/GTP exchange factor | 22.14 | 41.87 |
| MRET_3433 | nucleolar protein 12 | 68.98 | 81.52 |
| MRET_3434 | hydroxymethylglutaryl-CoA synthase | 40.57 | 147.25 |
| MRET_3435 | uncharacterized protein | 21.66 | 26.75 |
| MRET_3436 | AAA family ATPase | 185.05 | 124.44 |
| MRET_3437 | ubiquinone biosynthesis monooxygenase Coq6 | 69.84 | 24.15 |
| MRET_3438 | DNA repair/transcription protein MET18/MMS19 | 77.16 | 31.82 |

| | | | |
|---|---|---|---|
| MRET_3439 | essential RNA-binding component of cleavage and polyadenylation factor | 114.58 | 98.55 |
| MRET_3440 | DUF602 domain protein | 620.12 | 531.24 |
| MRET_3441 | DNA-directed RNA polymerase II subunit RPB7 | 656.36 | 670.25 |
| MRET_3442 | serine/threonine-protein kinase TTK/MPS1 | 54.82 | 83.31 |
| MRET_3443 | trehalose 6-phosphate synthase complex regulatory subunit | 217.94 | 62.06 |
| MRET_3444 | actin related protein 2/3 complex, subunit 5 | 74.08 | 92.87 |
| MRET_3445 | serine/threonine-protein kinase CLA4 | 206.45 | 157.1 |
| MRET_3446 | splicing factor U2AF 65 kDa subunit | 319.78 | 162.61 |
| MRET_3447 | pre-mRNA-processing factor 40 | 420.49 | 334.03 |
| MRET_3448 | heat shock protein | 55.58 | 97.16 |
| MRET_3449 | guanylate kinase | 26.5 | 44.59 |
| MRET_3450 | uncharacterized protein | 97.11 | 84.53 |
| MRET_3451 | cyclin-dependent kinase | 296.89 | 373.31 |
| MRET_3452 | peroxisomal membrane protein 4 | 375.11 | 438.75 |
| MRET_3453 | uncharacterized protein | 116.69 | 155.42 |
| MRET_3454 | pseudouridine 5'-phosphatase | 381.62 | 528.04 |
| MRET_3455 | charged multivesicular body protein 5 | 211.13 | 354.13 |
| MRET_3456 | bridging integrator 3 | 155.46 | 162.27 |
| MRET_3457 | serine/threonine-protein phosphatase 5 | 21.3 | 48.91 |
| MRET_3458 | ribosomal prokaryotic L21 protein | 97.48 | 107.33 |
| MRET_3459 | F-type H+-transporting ATPase subunit alpha | 1004.66 | 1063.28 |
| MRET_3460 | dynamin 1-like protein | 505.68 | 421.12 |
| MRET_3461 | U3 small nucleolar ribonucleoprotein protein LCP5 | 15.61 | 44.23 |
| MRET_3462 | sterol O-acyltransferase | 251.92 | 148.95 |
| MRET_3463 | solute carrier family 35 (UDP-xylose/UDP-N-acetylglucosamine transporter), member B4 | 44.99 | 43.36 |
| MRET_3464 | ubiquitin carboxyl-terminal hydrolase 25 | 8.65 | 18.78 |
| MRET_3465 | guanine nucleotide exchange factor LTE1 | 8.81 | 17.11 |
| MRET_3466 | ribonuclease P/MRP protein subunit POP3 | 10.26 | 24.03 |
| MRET_3467 | transcription factor | 33.51 | 38.19 |
| MRET_3468 | sortilin | 38.33 | 37.83 |
| MRET_3469 | uncharacterized protein | 17.83 | 28.54 |
| MRET_3470 | uncharacterized protein | 9.49 | 22.09 |
| MRET_3471 | tRNA-dihydrouridine synthase 4 | 13.5 | 39.86 |
| MRET_3472 | histone chaperone ASF1 | 47.21 | 102.92 |
| MRET_3473 | sterol 24-C-methyltransferase | 229.68 | 647.67 |
| MRET_3474 | U6 snRNA-associated Sm-like protein LSm8 | 50.45 | 97.36 |
| MRET_3475 | dihydroorotate dehydrogenase | 68.1 | 80.79 |

| | | | |
|---|---|---|---|
| MRET_3476 | zinc finger protein | 150.54 | 116 |
| MRET_3477 | structural maintenance of chromosomes protein | 129.98 | 75.55 |
| MRET_3478 | Prm1-pheromone-regulated multispanning membrane protein | 106.39 | 111.17 |
| MRET_3479 | RNA-binding protein with serine-rich domain 1-like protein | 31.04 | 60.24 |
| MRET_3480 | protein of unknown function (DUF2418) | 44.47 | 63.75 |
| MRET_3481 | uncharacterized protein | 86.49 | 64.37 |
| MRET_3482 | cyclin-dependent kinase regulatory subunit CKS1 | 420.21 | 402.79 |
| MRET_3483 | aldo/keto reductase family | 2666.3 | 2843.5 |
| MRET_3484 | aldo/keto reductase family | 657.43 | 668.51 |
| MRET_3485 | aldo/keto reductase family | 67.11 | 102.29 |
| MRET_3486 | mitochondrial protein FMP25 | 33.61 | 62.03 |
| MRET_3487 | glycyl-tRNA synthetase | 56.87 | 82.05 |
| MRET_3488 | uncharacterized protein | 52.87 | 51.91 |
| MRET_3489 | H/ACA ribonucleoprotein complex subunit 3 | 59.3 | 142.02 |
| MRET_3490 | p24 family protein delta-1 | 293.98 | 263.32 |
| MRET_3491 | E3 ubiquitin-protein ligase UBR7 | 162.04 | 173.26 |
| MRET_3492 | uncharacterized protein | 417.52 | 237.32 |
| MRET_3493 | uncharacterized protein | 23.38 | 36.43 |
| MRET_3494 | fanconi-associated nuclease 1 | 61.05 | 58.58 |
| MRET_3495 | myosin heavy chain | 68.94 | 142.68 |
| MRET_3496 | histone acetyltransferase | 20.09 | 51.63 |
| MRET_3497 | CBF NF-Y family transcription factor | 43.41 | 97.9 |
| MRET_3498 | platelet-activating factor acetylhydrolase IB subunit alpha | 19.9 | 58.18 |
| MRET_3499 | kinesin family member 4/21/27 | 36.1 | 43.05 |
| MRET_3500 | 3-deoxy-7-phosphoheptulonate synthase | 54.57 | 61.46 |
| MRET_3501 | negative cofactor 2 | 43.98 | 123.99 |
| MRET_3502 | methylenetetrahydrofolate dehydrogenase (NADP+) | 71.55 | 191.01 |
| MRET_3503 | RNA-binding protein | 60.6 | 87.92 |
| MRET_3504 | uncharacterized protein | 12.51 | 21.32 |
| MRET_3505 | F-type H+-transporting ATPase subunit beta | 1016.06 | 842.65 |
| MRET_3506 | septin 3/9/12 | 220.38 | 385.74 |
| MRET_3507 | secretory carrier-associated membrane protein | 73.55 | 96.86 |
| MRET_3508 | component of the RSC chromatin remodeling complex | 65.48 | 59.66 |
| MRET_3509 | secretory pathway protein Sec39 | 127.13 | 68.96 |
| MRET_3510 | betaine lipid synthase | 74.12 | 92.01 |
| MRET_3511 | AP-3 complex subunit delta | 21.4 | 43.17 |
| MRET_3512 | glutamine amidotransferase | 303.23 | 305.21 |

| | | | |
|---|---|---|---|
| MRET_3513 | uncharacterized protein | 93.71 | 70.33 |
| MRET_3514 | exosome complex component RRP46 | 132.03 | 126.89 |
| MRET_3515 | mitochondrial import inner membrane translocase subunit TIM54 | 31.14 | 55.88 |
| MRET_3516 | tRNA (uracil-5-)-methyltransferase TRM9 | 52.36 | 83.09 |
| MRET_3517 | 5'-methylthioadenosine phosphorylase | 114.3 | 172.94 |
| MRET_3518 | mRNA capping protein | 14.71 | 48.37 |
| MRET_3519 | catalytic protein kinase domain protein | 104.06 | 141.27 |
| MRET_3520 | DNA mismatch repair protein MSH6 | 96.22 | 57.06 |
| MRET_3521 | serine/threonine-protein kinase mTOR | 54.95 | 47.08 |
| MRET_3522 | phospholipase A2 activating protein | 74.28 | 116.45 |
| MRET_3523 | PQ loop repeat protein | 74.97 | 107.75 |
| MRET_3524 | endonuclease G, mitochondrial | 81.53 | 149.15 |
| MRET_3525 | ornithine--oxo-acid transaminase | 65.42 | 103.03 |
| MRET_3526 | heme steroid binding protein | 115.28 | 117.74 |
| MRET_3527 | H/ACA ribonucleoprotein complex subunit 2 | 62.73 | 176.35 |
| MRET_3528 | tryptophanyl-tRNA synthetase | 16.63 | 26.84 |
| MRET_3529 | peroxin-13 | 257.25 | 284.37 |
| MRET_3530 | large subunit ribosomal protein L10e | 6188.26 | 5243.83 |
| MRET_3531 | uncharacterized protein | 49.12 | 97.58 |
| MRET_3532 | replication factor A1 | 557.09 | 339.98 |
| MRET_3533 | phospholipid binding protein | 91.88 | 119.41 |
| MRET_3534 | vacuolar protein sorting-associated protein IST1 | 6.78 | 11.72 |
| MRET_3535 | DNA primase small subunit | 224.34 | 150.71 |
| MRET_3536 | CAP-Gly domain protein | 351.72 | 167.66 |
| MRET_3537 | glutamate decarboxylase | 192.78 | 140.33 |
| MRET_3538 | serine/threonine-protein kinase | 77.75 | 39.86 |
| MRET_3539 | breast cancer 2 susceptibility protein | 250.63 | 232.04 |
| MRET_3540 | kinase phosphorylation protein | 635.53 | 353.41 |
| MRET_3541 | glucosamine---fructose-6-phosphate aminotransferase | 170.88 | 286.62 |
| MRET_3542 | chromosome transmission fidelity protein 1 | 16.78 | 31.94 |
| MRET_3543 | phosphoglycerate mutase family protein | 71.23 | 106.75 |
| MRET_3544 | golgi matrix protein | 23.64 | 47.47 |
| MRET_3545 | small subunit ribosomal protein S20e | 82.22 | 442.92 |
| MRET_3546 | large subunit ribosomal protein L27Ae | 433.23 | 1111.11 |
| MRET_3547 | ariadne-1 | 203.42 | 148.31 |
| MRET_3548 | 3-hydroxyacyl-CoA dehydrogenase | 378.68 | 516.93 |
| MRET_3549 | dolichol-phosphate mannosyltransferase | 106.16 | 142.42 |

| | | | |
|---|---|---|---|
| MRET_3550 | WD repeat and SOF domain protein 1 | 106.93 | 100.61 |
| MRET_3551 | conserved hypothetical protein | 42.95 | 28.36 |
| MRET_3552 | ribose-phosphate pyrophosphokinase | 98.3 | 143.71 |
| MRET_3553 | small nuclear ribonucleoprotein D1 | 264.59 | 293.57 |
| MRET_3554 | exonuclease 1 | 44.54 | 88.36 |
| MRET_3555 | HCNGP-like protein | 47.94 | 96.49 |
| MRET_3556 | uncharacterized protein | 61.92 | 101.07 |
| MRET_3557 | UDP-glucose 6-dehydrogenase | 697.58 | 596.83 |
| MRET_3558 | conserved hypothetical protein | 223.83 | 163.13 |
| MRET_3559 | ribosome biogenesis protein UTP30 | 27.52 | 76.78 |
| MRET_3560 | ER membrane DUF1077 domain protein | 16.58 | 47.47 |
| MRET_3561 | cytochrome c oxidase subunit 20 | 34.27 | 74.44 |
| MRET_3562 | pyridine nucleotide-disulphide oxidoreductase | 265.73 | 230.63 |
| MRET_3563 | small subunit ribosomal protein S24e | 348.81 | 837.01 |
| MRET_3564 | U6 snRNA-associated Sm-like protein LSm3 | 62.6 | 112.13 |
| MRET_3565 | large subunit ribosomal protein L5 | 44.24 | 61.78 |
| MRET_3566 | DNA damage-responsive protein | 14.42 | 22.64 |
| MRET_3567 | vacuolar protein sorting-associated protein 41 | 81.82 | 50.64 |
| MRET_3568 | NADH dehydrogenase (ubiquinone) 1 alpha subcomplex subunit 13 | 54.65 | 80.08 |
| MRET_3569 | nucleolar protein 14 | 55.47 | 95.24 |
| MRET_3570 | DCN1-like protein 1/2 | 267.22 | 262.02 |
| MRET_3571 | small nuclear ribonucleoprotein F | 204.76 | 298.77 |
| MRET_3572 | solute carrier family 45, member 1/2/4 | 69.45 | 79.75 |
| MRET_3573 | 14-3-3 protein epsilon | 2900.7 | 2104.63 |
| MRET_3574 | AP-2 complex subunit alpha | 9.4 | 16.09 |
| MRET_3575 | RNA polymerase I-specific transcription initiation factor RRN3 | 382.49 | 236.35 |
| MRET_3576 | GTP-binding nuclear protein Ran | 339.28 | 459.93 |
| MRET_3577 | DNA-directed RNA polymerase I subunit RPA43 | 32.62 | 81.88 |
| MRET_3578 | thioredoxin | 65.71 | 104.54 |
| MRET_3579 | negative regulation of cAMP metabolic process | 6.23 | 13.98 |
| MRET_3580 | putative methyltransferase | 45.74 | 30.35 |
| MRET_3581 | uncharacterized protein | 84.37 | 88.97 |
| MRET_3582 | mRNA m6A methyltransferase | 39.59 | 98.64 |
| MRET_3583 | uncharacterized protein | 29.59 | 32.89 |
| MRET_3584 | uncharacterized protein | 26.63 | 55.89 |
| MRET_3585 | F-box and WD-40 domain protein 1/11 | 86.28 | 247.06 |
| MRET_3586 | Pal1 cell morphology protein | 232.92 | 260.84 |

| | | | |
|---|---|---|---|
| MRET_3587 | lipid transfer protein | 1354.37 | 1324.07 |
| MRET_3588 | collagen | 100.8 | 206.58 |
| MRET_3589 | putative sensor/transporter protein involved in cell wall biogenesis | 80.24 | 98.82 |
| MRET_3590 | eukaryotic mitochondrial regulator protein | 2183.62 | 829.5 |
| MRET_3591 | uncharacterized protein | 127.83 | 111.54 |
| MRET_3592 | cortical actin cytoskeleton protein asp1 | 98.39 | 57.11 |
| MRET_3593 | LETM1-like protein | 117.02 | 247.36 |
| MRET_3594 | U6 snRNA-associated Sm-like protein LSm1 | 86.11 | 223.22 |
| MRET_3595 | AdoMet-dependent methyltransferase | 64.94 | 138.74 |
| MRET_3596 | WD40 repeat-like protein | 10.18 | 20.23 |
| MRET_3597 | trafficking protein particle complex subunit 8 | 19.91 | 32.04 |
| MRET_3598 | integral peroxisomal membrane peroxin | 20.66 | 11.12 |
| MRET_3599 | nitric oxide dioxygenase | 380.1 | 54.74 |
| MRET_3600 | nucleolin | 30.8 | 72.79 |
| MRET_3601 | DNA-directed RNA polymerase III subunit RPC1 | 112.06 | 74.33 |
| MRET_3602 | protein unc-45 | 27.67 | 27.3 |
| MRET_3603 | ubiquitin-activating enzyme E1 | 777.99 | 481.84 |
| MRET_3604 | uncharacterized protein | 187.29 | 145.71 |
| MRET_3605 | SRP40, C-terminal domain protein | 277.84 | 1225.93 |
| MRET_3606 | mediator of RNA polymerase II transcription subunit 13 | 94.7 | 50.21 |
| MRET_3607 | uncharacterized protein | 50.74 | 46.43 |
| MRET_3608 | kinesin-like protein 8 | 20.81 | 18.47 |
| MRET_3609 | uncharacterized protein | 28.82 | 17.82 |
| MRET_3610 | homoisocitrate dehydrogenase | 13.62 | 37.26 |
| MRET_3611 | elongation factor | 735.06 | 507 |
| MRET_3612 | elongation factor 1 alpha-like protein | 782.99 | 641.97 |
| MRET_3613 | Myb-like DNA-binding domain protein | 151.36 | 52.89 |
| MRET_3614 | trafficking protein particle complex subunit 2 | 59.18 | 101.26 |
| MRET_3615 | uncharacterized protein | 252.15 | 391.99 |
| MRET_3616 | protein kinase C substrate 80K-H | 27.02 | 51.12 |
| MRET_3617 | lysophosphatidate acyltransferase | 77.1 | 99.2 |
| MRET_3618 | uncharacterized protein | 56.56 | 63.38 |
| MRET_3619 | TBC domain protein | 635.64 | 721.5 |
| MRET_3620 | Rab5-interacting protein (Rab5ip) | 363.5 | 459.85 |
| MRET_3621 | COMPASS component SWD1 | 141.9 | 136.55 |
| MRET_3622 | ATP-dependent Clp protease ATP-binding subunit ClpB | 1660.48 | 1980.09 |
| MRET_3623 | transcription factor | 45.03 | 43.39 |

| | | | |
|---|---|---|---|
| MRET_3624 | RNA recognition motif domain protein | 283.87 | 274.05 |
| MRET_3625 | aldehyde dehydrogenase (NAD+) | 319.44 | 217.66 |
| MRET_3626 | serine/threonine-protein kinase | 181.96 | 142.43 |
| MRET_3627 | phosphoserine phosphatase | 49.92 | 52.59 |
| MRET_3628 | 20S proteasome subunit alpha 2 | 104.09 | 198.85 |
| MRET_3629 | ribosome biogenesis protein ENP2 | 34.88 | 47.36 |
| MRET_3630 | large subunit ribosomal protein L11 | 13.25 | 93.23 |
| MRET_3631 | sorbose reductase | 794.79 | 709.34 |
| MRET_3632 | iron-sulfur cluster assembly enzyme ISCU, mitochondrial | 1888.51 | 1866.17 |
| MRET_3633 | charged multivesicular body protein 6 | 77.69 | 150.91 |
| MRET_3634 | ribosomal RNA-processing protein 36 | 38.86 | 52.61 |
| MRET_3635 | U3 small nucleolar RNA-associated protein 14 | 28.17 | 54.56 |
| MRET_3636 | 60S ribosome subunit biogenesis protein NIP7 | 37.84 | 59 |
| MRET_3637 | AHNAK nucleoprotein | 181.16 | 140.21 |
| MRET_3638 | ribosomal RNA-processing protein 8 | 64.61 | 59.21 |
| MRET_3639 | brefeldin A-inhibited guanine nucleotide-exchange protein | 45.59 | 43.34 |
| MRET_3640 | translation initiation factor 6 | 323.04 | 251.76 |
| MRET_3641 | mitochondrial import inner membrane translocase subunit TIM9 | 49.23 | 227.11 |
| MRET_3642 | uncharacterized protein | 16.47 | 62.4 |
| MRET_3643 | THO complex subunit 2 | 43.39 | 67.79 |
| MRET_3644 | DNA-directed RNA polymerases I, II, and III subunit RPABC4 | 279.76 | 262.89 |
| MRET_3645 | AP-3 complex subunit mu | 39.44 | 58.75 |
| MRET_3646 | ubiquitin metalloprotease fusion protein | 201.68 | 204.07 |
| MRET_3647 | glycosyl hydrolase catalytic core | 116.73 | 508.43 |
| MRET_3648 | bud emergence protein 1 | 13.75 | 41.05 |
| MRET_3649 | uncharacterized protein | 50.41 | 105.81 |
| MRET_3650 | proton-dependent oligopeptide transporter, POT family | 46.5 | 70.8 |
| MRET_3651 | uncharacterized protein | 198.1 | 225.89 |
| MRET_3652 | guanyl nucleotide binding protein | 55 | 106.45 |
| MRET_3653 | mitochondrial 37S ribosomal protein NAM9 | 19.03 | 65.94 |
| MRET_3654 | S-adenosylmethionine synthetase | 203.53 | 260.59 |
| MRET_3655 | uncharacterized protein | 24.57 | 36.64 |
| MRET_3656 | sulfite reductase (NADPH) flavoprotein alpha-component | 71.56 | 78.05 |
| MRET_3657 | small subunit ribosomal protein S7e | 37.05 | 230.71 |
| MRET_3658 | DUF1768 domain protein | 24.45 | 111.84 |
| MRET_3659 | mitochondrial alcohol dehydrogenase isozyme III | 26.56 | 64.86 |
| MRET_3660 | oligosaccharyltransferase complex subunit epsilon | 63.42 | 86.3 |

| | | | |
|---|---|---|---|
| MRET_3661 | peptide chain release factor subunit 3 | 454.25 | 387.88 |
| MRET_3662 | conserved hypothetical protein | 90.44 | 102.18 |
| MRET_3663 | Rho GDP-dissociation inhibitor | 820.72 | 532.63 |
| MRET_3664 | coupling of ubiquitin conjugation to ER degradation protein 1 | 252.88 | 231.35 |
| MRET_3665 | NADH dehydrogenase (ubiquinone) 1 beta subcomplex subunit 9 | 679.78 | 768.1 |
| MRET_3666 | flavin reductase domain protein | 441.21 | 311.35 |
| MRET_3667 | trafficking protein particle complex subunit 2 | 77.18 | 129.1 |
| MRET_3668 | uncharacterized protein | 244.6 | 207.78 |
| MRET_3669 | fungal protein of unknown function (DUF1748) | 349.2 | 164.12 |
| MRET_3670 | serine/threonine-protein kinase ULK2 | 1303.73 | 1080.01 |
| MRET_3671 | protein-serine/threonine kinase | 18.46 | 33.01 |
| MRET_3672 | U4/U6.U5 tri-snRNP-associated protein 1 | 45.17 | 65.43 |
| MRET_3673 | RhoGAP | 45.12 | 86.45 |
| MRET_3674 | uncharacterized protein | 179.09 | 105.15 |
| MRET_3675 | CCR4-NOT transcription complex subunit 3 | 73.11 | 96.05 |
| MRET_3676 | large subunit ribosomal protein L19 | 30.02 | 25.17 |
| MRET_3677 | small subunit ribosomal protein S14 | 74.45 | 47.41 |
| MRET_3678 | 5-aminolevulinate synthase | 706.74 | 224.22 |
| MRET_3679 | integral membrane protein | 134.27 | 143.98 |
| MRET_3680 | alpha/beta-hydrolase | 98.73 | 62.95 |
| MRET_3681 | uncharacterized protein | 38.19 | 47.31 |
| MRET_3682 | cytochrome c oxidase assembly factor 5 | 274.21 | 559.27 |
| MRET_3683 | conserved serine/proline-rich protein | 65.22 | 68.68 |
| MRET_3684 | uncharacterized protein | 95.43 | 71.5 |
| MRET_3685 | Ras GTPase-activating-like protein IQGAP2/3 | 15.16 | 30.4 |
| MRET_3686 | coatomer subunit zeta | 477.87 | 832.31 |
| MRET_3687 | TBC1 domain family member 20 | 219.68 | 134.8 |
| MRET_3688 | Ras homolog enriched in brain | 44.51 | 80.8 |
| MRET_3689 | D-xylulose reductase | 791.41 | 429.95 |
| MRET_3690 | uncharacterized protein | 24.8 | 30.58 |
| MRET_3691 | gamma-tubulin complex component 3 | 93.22 | 54.03 |
| MRET_3692 | endosome-associated ubiquitin isopeptidase (AmsH) | 31.84 | 22.49 |
| MRET_3693 | paxillin | 608.56 | 309.34 |
| MRET_3694 | uncharacterized protein | 35.54 | 70.82 |
| MRET_3695 | DNA replication licensing factor MCM7 | 117.15 | 103.26 |
| MRET_3696 | nucleoporin NDC1 | 123.38 | 58.54 |
| MRET_3697 | superoxide dismutase, Fe-Mn family | 54.87 | 63.68 |

| | | | |
|---|---|---|---|
| MRET_3698 | SNARE associated golgi protein | 112.79 | 204.68 |
| MRET_3699 | DUF431 domain protein | 32.48 | 29.72 |
| MRET_3700 | DUF775 domain protein | 253.03 | 125.95 |
| MRET_3701 | ribokinase | 31.11 | 33.49 |
| MRET_3702 | general repressor of transcription | 20.69 | 23.31 |
| MRET_3703 | threonine synthase | 97.07 | 92.38 |
| MRET_3704 | riboflavin kinase | 71.93 | 54.81 |
| MRET_3705 | uncharacterized protein | 91.16 | 54.19 |
| MRET_3706 | RNA-binding protein | 150.48 | 186.42 |
| MRET_3707 | chromatin structure-remodeling complex subunit SFH1 | 52.08 | 81.4 |
| MRET_3708 | small subunit ribosomal protein S3Ae | 89.75 | 597.14 |
| MRET_3709 | protein of unknown function (DUF788) | 135.59 | 12.06 |
| MRET_3710 | transcription factor | 76.43 | 35.41 |
| MRET_3711 | RF-1 domain protein | 562.04 | 375.09 |
| MRET_3712 | AMMECR1 family protein | 13.91 | 8.48 |
| MRET_3713 | phospholipid-binding protein that interacts with both Ypt7p and Vps33p | 46.85 | 75.76 |
| MRET_3714 | myosin regulatory light chain cdc4 | 660.44 | 867.74 |
| MRET_3715 | small subunit ribosomal protein S25e | 257.44 | 880.06 |
| MRET_3716 | 40S ribosomal protein S25 | 47.67 | 122.99 |
| MRET_3717 | F-type H+-transporting ATPase subunit d | 513.03 | 696.03 |
| MRET_3718 | cAMP-dependent protein kinase regulator | 135.21 | 161.27 |
| MRET_3719 | aspartyl-tRNA synthetase | 64.57 | 51.07 |
| MRET_3720 | Tol-Pal system protein YbgF | 900.92 | 1263.11 |
| MRET_3721 | conserved hypothetical protein | 57.14 | 138.34 |
| MRET_3722 | uncharacterized protein | 535.17 | 421.74 |
| MRET_3723 | Shwachman-Bodian-Diamond syndrome (SBDS) protein | 2936.2 | 2704.36 |
| MRET_3724 | pleiotropic regulator 1 | 458.68 | 445.47 |
| MRET_3725 | N-acetylated-alpha-linked acidic dipeptidase | 255.06 | 164.59 |
| MRET_3726 | 20S proteasome subunit alpha 6 | 424.76 | 381.12 |
| MRET_3727 | signal recognition particle receptor beta subunit | 9.24 | 25.71 |
| MRET_3728 | ribosome biogenesis protein ERB1 | 61.98 | 75.43 |
| MRET_3729 | oxidoreductase | 104.06 | 59.75 |
| MRET_3730 | transporter | 91.06 | 94.54 |
| MRET_3731 | charged multivesicular body protein 4 | 67.84 | 181.44 |
| MRET_3732 | transporter | 167.28 | 234.12 |
| MRET_3733 | phosphatidylinositol 4-kinase B | 19.82 | 25.64 |
| MRET_3734 | transformation/transcription domain-associated protein | 291.64 | 231.65 |

| | | | |
|---|---|---|---|
| MRET_3735 | DNA-binding protein | 17.02 | 19.98 |
| MRET_3736 | transporter | 36.21 | 2.92 |
| MRET_3737 | Chs5-Arf1p-binding protein BUD7/BCH1 | 16.38 | 34.68 |
| MRET_3738 | solute carrier family 25 (mitochondrial carnitine/acylcarnitine transporter), member 20/29 | 22.93 | 88.54 |
| MRET_3739 | DnaJ homolog subfamily B member 12 | 63.95 | 424.58 |
| MRET_3740 | DNA-directed RNA polymerase I subunit RPA12 | 15.05 | 23.65 |
| MRET_3741 | hydroxyacylglutathione hydrolase | 179.66 | 0.41 |
| MRET_3742 | short-chain dehydrogenase | 541.52 | 985.68 |
| MRET_3743 | protein HIRA/HIR1 | 11.4 | 13.8 |
| MRET_3744 | uncharacterized protein | 21.02 | 27.59 |
| MRET_3745 | histone acetyltransferase | 110.59 | 85.1 |
| MRET_3746 | mitochondrial distribution and morphology protein 34 | 263.66 | 173.01 |
| MRET_3747 | TBC domain protein | 16.91 | 38.55 |
| MRET_3748 | TBC domain protein | 20.64 | 44.06 |
| MRET_3749 | cyclin | 832.34 | 728.39 |
| MRET_3750 | uncharacterized protein | 50.18 | 41.86 |
| MRET_3751 | magnesium transporter | 80.54 | 87.85 |
| MRET_3752 | transaldolase | 622.57 | 655.62 |
| MRET_3753 | mediator of RNA polymerase II transcription subunit 11 | 608.22 | 553.08 |
| MRET_3754 | N-acetyltransferase 10 | 23.35 | 20.46 |
| MRET_3755 | type I protein arginine methyltransferase | 11.53 | 20.7 |
| MRET_3756 | Bromodomain associated protein | 35.08 | 50.22 |
| MRET_3757 | NADH dehydrogenase (ubiquinone) Fe-S protein 2 | 266.18 | 308.05 |
| MRET_3758 | endoplasmic reticulum-golgi intermediate compartment protein 3 | 120.21 | 111.71 |
| MRET_3759 | response regulator receiver domain protein | 32.85 | 44.29 |
| MRET_3760 | tubulin-specific chaperone B | 67.39 | 37.36 |
| MRET_3761 | GINS complex subunit 1 | 36.75 | 35.29 |
| MRET_3762 | uncharacterized protein | 318.34 | 180.24 |
| MRET_3763 | sister chromatid cohesion protein DCC1 | 15.03 | 39.11 |
| MRET_3764 | solute carrier family 24 (sodium/potassium/calcium exchanger), member 6 | 21.42 | 29.66 |
| MRET_3765 | triacylglycerol lipase | 44.78 | 15.81 |
| MRET_3766 | serine/threonine-protein phosphatase | 119.36 | 42.21 |
| MRET_3767 | uncharacterized protein | 122.84 | 264.28 |
| MRET_3768 | uncharacterized protein | 62.7 | 102.48 |
| MRET_3769 | uncharacterized protein | 257.39 | 94.23 |
| MRET_3770 | eukaryotic aspartyl protease | 9.66 | 82.85 |
| MRET_3771 | deoxyribodipyrimidine photo-lyase | 56.57 | 75.84 |

| | | | |
|---|---|---|---|
| MRET_3772 | lipase precursor-like protein | 28.34 | 60.43 |
| MRET_3773 | MFS family protein | 93.43 | 170.23 |
| MRET_3774 | multidrug transporter of the major facilitator superfamily | 205.69 | 243.56 |
| MRET_3775 | multidrug transporter of the major facilitator superfamily | 464.06 | 247.73 |
| MRET_3776 | glutathione S-transferase | 737.76 | 285.68 |
| MRET_3777 | mitochondrial intermediate peptidase | 154.81 | 107.85 |
| MRET_3778 | phenylalanyl-tRNA synthetase alpha chain | 74.24 | 87.24 |
| MRET_3779 | U4/U6 small nuclear ribonucleoprotein PRP4 | 127.49 | 118.25 |
| MRET_3780 | large subunit ribosomal protein L24e | 141.13 | 175.83 |
| MRET_3781 | glutaredoxin domain protein | 68.77 | 143.21 |
| MRET_3782 | signal recognition particle subunit SRP9 | 13.51 | 41.5 |
| MRET_3783 | uncharacterized protein | 72.31 | 96.77 |
| MRET_3784 | FYVE, RhoGEF and PH domain protein | 13.29 | 34.66 |
| MRET_3785 | Mus7/MMS22 family protein | 26.04 | 20.97 |
| MRET_3786 | sphingomyelin phosphodiesterase | 158.93 | 145.26 |
| MRET_3787 | proteasomal ATPase-associated factor 1 | 379.67 | 143.7 |
| MRET_3788 | transporter (MirC) | 167.3 | 128.2 |
| MRET_3789 | RNA binding effector protein Scp160 | 79.41 | 93.7 |
| MRET_3790 | DUF907 domain protein | 357.35 | 208.1 |
| MRET_3791 | nicotinamidase | 858.1 | 558.17 |
| MRET_3792 | protein phosphatase PTC7 | 44.41 | 47.18 |
| MRET_3793 | component of the NuA4 histone acetyltransferase complex | 304.59 | 91.85 |
| MRET_3794 | uncharacterized protein | 174.49 | 111.88 |
| MRET_3795 | uncharacterized protein | 42.13 | 61.78 |
| MRET_3796 | NADH dehydrogenase (ubiquinone) Fe-S protein 4 | 107.8 | 185.6 |
| MRET_3797 | exosome complex component RRP40 | 46.77 | 48.13 |
| MRET_3798 | vacuolar protein sorting-associated protein 53 | 128.82 | 128.08 |
| MRET_3799 | nucleosome assembly protein 1-like 1 | 303.91 | 427.19 |
| MRET_3800 | DNA polymerase kappa | 45.81 | 50.8 |
| MRET_3801 | pH-response regulator protein palC | 37.51 | 47.08 |
| MRET_3802 | F-type H+-transporting ATPase subunit f | 466.17 | 602.31 |
| MRET_3803 | uncharacterized protein | 92.33 | 387.38 |
| MRET_3804 | small subunit ribosomal protein SAe | 226.46 | 595.76 |
| MRET_3805 | histone acetyltransferase HTATIP | 288.25 | 257.32 |
| MRET_3806 | splicing factor 3B subunit 1 | 255.7 | 164.1 |
| MRET_3807 | U3 small nucleolar RNA-associated protein 13 | 32.4 | 46.74 |
| MRET_3808 | uncharacterized protein | 318.66 | 508.13 |

| | | | |
|---|---|---|---|
| MRET_3809 | telomerase reverse transcriptase | 154.61 | 123.39 |
| MRET_3810 | PRA1 family protein 1 | 1039.89 | 699.23 |
| MRET_3811 | mitochondrial carrier protein | 1079.09 | 563 |
| MRET_3812 | uncharacterized protein | 197.33 | 248.13 |
| MRET_3813 | mitochondrial carrier protein | 62.83 | 55.94 |
| MRET_3814 | uncharacterized protein | 55.52 | 69.65 |
| MRET_3815 | component of nuclear aminoacylation-dependent tRNA export pathway | 55.84 | 68.85 |
| MRET_3816 | mediator of RNA polymerase II transcription subunit 14 | 8.48 | 16.32 |
| MRET_3817 | maltose acetyltransferase | 56.02 | 176.89 |
| MRET_3818 | N-terminal acetyltransferase B complex non-catalytic subunit | 29.07 | 32.68 |
| MRET_3819 | crossover junction endonuclease EME1 | 47.2 | 50.9 |
| MRET_3820 | phosphatidylinositol glycan, class B | 37.88 | 46.42 |
| MRET_3821 | mitogen-activated protein kinase kinase | 187.33 | 145.41 |
| MRET_3822 | autophagy-related protein 13 | 187.94 | 139.91 |
| MRET_3823 | peroxin-16 | 272.84 | 204.89 |
| MRET_3824 | methyltransferase | 63.62 | 56.99 |
| MRET_3825 | lanosterol synthase | 522.06 | 140.99 |
| MRET_3826 | transmembrane protein 167 | 35.55 | 34.78 |
| MRET_3827 | 26S proteasome non-ATPase regulatory subunit 10 | 41.07 | 50.33 |
| MRET_3828 | transcription initiation factor TFIID subunit 11 | 69.05 | 89.71 |
| MRET_3829 | saccharopine dehydrogenase | 40.26 | 50.39 |
| MRET_3830 | tubulin beta | 183.81 | 211.89 |
| MRET_3831 | filamentation protein (Rhf1) | 144.58 | 81.18 |
| MRET_3832 | carbon catabolite-derepressing protein kinase | 60.81 | 56.95 |
| MRET_3833 | Ca2+:H+ antiporter | 46.04 | 45.11 |
| MRET_3834 | epsin | 80.65 | 178.7 |
| MRET_3835 | ammonium transporter, Amt family | 38.69 | 36.78 |
| MRET_3836 | tRNA pseudouridine13 synthase | 56.28 | 50.51 |
| MRET_3837 | Ras-related protein Rab-18 | 35.69 | 35.49 |
| MRET_3838 | UPF0172 domain protein | 95.28 | 98.47 |
| MRET_3839 | adrenodoxin-NADP+ reductase | 18.12 | 29.57 |
| MRET_3840 | splicing factor 3A subunit 2 | 22.93 | 45.49 |
| MRET_3841 | cell division control protein 12 | 405.39 | 428.38 |
| MRET_3842 | DNA repair and recombination protein RAD52 | 25.18 | 90.5 |
| MRET_3843 | vesicle transport protein SEC22 | 78.64 | 148.87 |
| MRET_3844 | general stress response protein Whi2 | 196.06 | 465.98 |
| MRET_3845 | uncharacterized protein | 124.58 | 151.34 |

| | | | |
|---|---|---|---|
| MRET_3846 | small subunit ribosomal protein S10 | 77.95 | 130.6 |
| MRET_3847 | phospholipase A2 | 32.74 | 32.13 |
| MRET_3848 | uncharacterized protein | 471.74 | 226.03 |
| MRET_3849 | transcription initiation protein SPT3 | 91.38 | 83.73 |
| MRET_3850 | FHA domain protein | 163.93 | 141.21 |
| MRET_3851 | elongation factor G | 78.1 | 110.93 |
| MRET_3852 | STE24 endopeptidase | 53.83 | 43.06 |
| MRET_3853 | STE24 endopeptidase | 554.19 | 320.16 |
| MRET_3854 | DNA ligase 1 | 48.12 | 55.42 |
| MRET_3855 | structural maintenance of chromosomes protein | 55.62 | 47.07 |
| MRET_3856 | solute carrier family 25 (mitochondrial aspartate/glutamate transporter), member 12/13 | 71.38 | 75.53 |
| MRET_3857 | chaperonin GroEL | 1715.35 | 2307.74 |
| MRET_3858 | chaperonin GroES | 1531.6 | 1049.56 |
| MRET_3859 | putative mago nashi protein, exon junction complex | 49.29 | 156.57 |
| MRET_3860 | N-glycosylation protein | 53.9 | 66.35 |
| MRET_3861 | uncharacterized protein | 174.73 | 147.33 |
| MRET_3862 | uncharacterized protein | 13.14 | 11.82 |
| MRET_3863 | mitochondrial import receptor subunit TOM70 | 385.12 | 271.72 |
| MRET_3864 | small ubiquitin-related modifier | 450.91 | 527.23 |
| MRET_3865 | ESF2/ABP1 family protein | 82.84 | 84.7 |
| MRET_3866 | diphthamide biosynthesis protein 3 | 112.93 | 68.93 |
| MRET_3867 | DUF1708 domain protein | 188.2 | 122.06 |
| MRET_3868 | putative subunit of the 90S preribosome processome complex | 182.48 | 159.56 |
| MRET_3869 | Dr1-associated corepressor | 218.33 | 255.77 |
| MRET_3870 | carbamoyl-phosphate synthase/aspartate carbamoyltransferase | 309.47 | 100.35 |
| MRET_3871 | uncharacterized protein | 25.56 | 22.81 |
| MRET_3872 | BAR domain protein | 92.1 | 99.08 |
| MRET_3873 | mortality factor 4-like protein 1 | 38.17 | 60.57 |
| MRET_3874 | uncharacterized protein | 99 | 136.18 |
| MRET_3875 | mitochondrial import inner membrane translocase subunit TIM13 | 25.15 | 44.59 |
| MRET_3876 | uncharacterized protein | 20.64 | 27.96 |
| MRET_3877 | Bromodomain associated protein | 18.73 | 29.83 |
| MRET_3878 | tRNA (guanine37-N1)-methyltransferase | 15.8 | 24.39 |
| MRET_3879 | uncharacterized protein | 638.03 | 361.09 |
| MRET_3880 | cation efflux family protein | 45.18 | 27.71 |
| MRET_3881 | 20S proteasome subunit beta 3 | 56.12 | 112.92 |
| MRET_3882 | ATPase family AAA domain protein | 194.94 | 141.06 |

| | | | |
|---|---|---|---|
| MRET_3883 | ATP phosphoribosyltransferase | 149.68 | 189.52 |
| MRET_3884 | transcription initiation factor TFIIIB 90 kDa subunit | 68.01 | 66.13 |
| MRET_3885 | CobW domain protein | 51.51 | 61.9 |
| MRET_3886 | DUF202 domain protein | 15.56 | 16.1 |
| MRET_3887 | SH3 domain protein | 28.75 | 82.09 |
| MRET_3888 | ribonuclease P/MRP protein subunit POP5 | 62.32 | 226.72 |
| MRET_3889 | Mob1/phocein family | 178.99 | 451.02 |
| MRET_3890 | zinc finger protein, C3HC4 type (RING finger) | 2102.8 | 1445.2 |
| MRET_3891 | vacuolar protein-sorting protein BRO1 | 367.43 | 142.88 |
| MRET_3892 | condensin complex subunit 3 | 18.7 | 57.18 |
| MRET_3893 | lipoate---protein ligase | 327.96 | 239.04 |
| MRET_3894 | chitin synthase | 54.89 | 56.44 |
| MRET_3895 | uncharacterized protein | 438.62 | 218.55 |
| MRET_3896 | uncharacterized protein | 97.86 | 52.61 |
| MRET_3897 | charged multivesicular body protein 1 | 52.74 | 107.07 |
| MRET_3898 | protein of unknown function (DUF2416) | 9.78 | 10.59 |
| MRET_3899 | mitochondrial distribution and morphology protein 31 | 96.71 | 55.31 |
| MRET_3900 | centromere protein k | 30.39 | 31.78 |
| MRET_3901 | peroxisomal carrier protein | 133.6 | 86.09 |
| MRET_3902 | UTP--glucose-1-phosphate uridylyltransferase | 91.4 | 103.98 |
| MRET_3903 | capping protein (actin filament) muscle Z-line, beta | 35.53 | 63.44 |
| MRET_3904 | protein phosphatase 1 regulatory subunit 7 | 179.63 | 118.23 |
| MRET_3905 | large subunit ribosomal protein L41 | 14.09 | 57.99 |
| MRET_3906 | ATP-binding cassette, subfamily F, member 2 | 86.03 | 98.29 |
| MRET_3907 | nuclear transcription Y subunit beta | 62.07 | 170.1 |
| MRET_3908 | protein cornichon | 72.07 | 266.9 |
| MRET_3909 | structural maintenance of chromosomes protein | 26.93 | 46.41 |
| MRET_3910 | uncharacterized protein | 793.52 | 299.27 |
| MRET_3911 | large subunit ribosomal protein L13 | 169.43 | 181.38 |
| MRET_3912 | clathrin light chain | 82.14 | 141.73 |
| MRET_3913 | DNA-binding protein | 30.43 | 41.42 |
| MRET_3914 | NAD binding dehydrogenase family protein | 79.55 | 93.4 |
| MRET_3915 | cofilin/tropomyosin-type actin-binding protein | 80.28 | 70.46 |
| MRET_3916 | NAD binding dehydrogenase family protein | 45.62 | 120.18 |
| MRET_3917 | conserved hypothetical protein | 265.68 | 406.32 |
| MRET_3918 | RNA-binding protein NOB1 | 80.17 | 129.75 |
| MRET_3919 | DnaJ domain protein | 164.54 | 95.01 |

| | | | |
|---|---|---|---|
| MRET_3920 | polarized growth protein | 104.94 | 188.58 |
| MRET_3921 | metacaspase-1 | 570.95 | 359.4 |
| MRET_3922 | tubulin-specific chaperone C | 20.18 | 18.41 |
| MRET_3923 | ubiquitin carboxyl-terminal hydrolase 36/42 | 41.58 | 39.09 |
| MRET_3924 | vacuole morphology and inheritance protein 14 | 46.69 | 44.02 |
| MRET_3925 | beta-catenin-like protein 1 | 58.21 | 49.32 |
| MRET_3926 | poly(A) RNA-binding protein | 190.07 | 86.62 |
| MRET_3927 | uncharacterized protein | 441.86 | 775.25 |
| MRET_3928 | nicotinamide-nucleotide adenylyltransferase | 68.22 | 81.9 |
| MRET_3929 | pre-mRNA-splicing factor ISY1 | 19.45 | 55.48 |
| MRET_3930 | large subunit ribosomal protein L10Ae | 223.92 | 623.84 |
| MRET_3931 | small subunit ribosomal protein S11e | 114 | 526.47 |
| MRET_3932 | small subunit ribosomal protein S12e | 312.84 | 1012.46 |
| MRET_3933 | histone deacetylase complex subunit SAP18 | 388.11 | 402.4 |
| MRET_3934 | uncharacterized protein | 82.87 | 44.3 |
| MRET_3935 | uncharacterized protein | 118.17 | 65.79 |
| MRET_3936 | tubulin-specific chaperone E | 40.41 | 22.55 |
| MRET_3937 | uncharacterized protein | 98.36 | 45.68 |
| MRET_3938 | ribosome assembly protein 1 | 170.69 | 119.12 |
| MRET_3939 | Prp8 binding protein | 94.8 | 89.9 |
| MRET_3940 | vacuolar protein sorting-associated protein | 70.01 | 18.83 |
| MRET_3941 | lysophospholipid acyltransferase | 46.08 | 28.32 |
| MRET_3942 | palmitoyl-protein thioesterase | 25.56 | 24.08 |
| MRET_3943 | phosphatidylglycerol phospholipase C | 118.67 | 136.16 |
| MRET_3944 | vaculolar membrane protein | 63.49 | 63.01 |
| MRET_3945 | uncharacterized protein | 279.07 | 139.73 |
| MRET_3946 | peptidyl-prolyl cis-trans isomerase-like 3 | 227.22 | 173.05 |
| MRET_3947 | vacuolar transporter chaperone 1 | 207.47 | 235.79 |
| MRET_3948 | modifier of rudimentary (Mod(r)) protein | 120.62 | 79.4 |
| MRET_3949 | Myb-like DNA-binding domain protein | 56.55 | 70.99 |
| MRET_3950 | coatomer subunit beta' | 86.71 | 90.13 |
| MRET_3951 | F-box and leucine-rich repeat protein GRR1 | 167.33 | 118.01 |
| MRET_3952 | uncharacterized protein | 70.38 | 68.21 |
| MRET_3953 | uncharacterized protein | 450.9 | 289.28 |
| MRET_3954 | transcription initiation factor TFIID subunit 5 | 30.34 | 45.86 |
| MRET_3955 | serine palmitoyltransferase | 154.98 | 193.31 |
| MRET_3956 | ubiquitin-conjugating enzyme E2 A | 73.69 | 174.87 |

| | | | |
|---|---|---|---|
| MRET_3957 | uncharacterized protein | 245.13 | 451.5 |
| MRET_3958 | trimethylguanosine synthase | 95.93 | 245.21 |
| MRET_3959 | exosome complex component MTR3 | 83.97 | 139.43 |
| MRET_3960 | glycoside hydrolase family 16 protein | 5786.22 | 8077.37 |
| MRET_3961 | DNA repair and recombination protein RAD54 and RAD54-like protein | 445.52 | 261.43 |
| MRET_3962 | transcription initiation factor TFIIH subunit 4 | 47.02 | 78.39 |
| MRET_3963 | acyl-CoA dehydrogenase | 580.43 | 442.35 |
| MRET_3964 | putative phosphomutase | 75.19 | 94.9 |
| MRET_3965 | uncharacterized protein | 76.33 | 112.37 |
| MRET_3966 | phospholipid-translocating ATPase | 55.99 | 78.04 |
| MRET_3967 | SUN domain protein (Adg3) | 233.03 | 195.78 |
| MRET_3968 | uncharacterized protein | 1579.5 | 1337.47 |
| MRET_3969 | hexosyltransferase | 80.6 | 61.69 |
| MRET_3970 | inner centromere protein | 19.69 | 51.89 |
| MRET_3971 | SNARE complex subunit Vam7 | 34.74 | 54.64 |
| MRET_3972 | general repressor of transcription | 231.1 | 218.15 |
| MRET_3973 | serine/arginine repetitive matrix protein 1 | 436.45 | 266.42 |
| MRET_3974 | ribose-phosphate pyrophosphokinase | 74.29 | 125.47 |
| MRET_3975 | ATP-dependent RNA helicase DDX55/SPB4 | 59.04 | 25.67 |
| MRET_3976 | GRAM domain protein | 57.75 | 44.79 |
| MRET_3977 | AN1-like zinc finger protein | 4.82 | 7.31 |
| MRET_3978 | component of the endoplasmic reticulum- associated degradation (ERAD) pathway | 13.72 | 10.66 |
| MRET_3979 | golgi phosphoprotein 3 | 35.73 | 59.34 |
| MRET_3980 | uncharacterized protein | 7.35 | 12.48 |
| MRET_3981 | U3 small nucleolar RNA-associated protein 22 | 10.96 | 17.8 |
| MRET_3982 | ankyrin repeat domain protein | 33.5 | 46.65 |
| MRET_3983 | uncharacterized protein | 389.46 | 154.07 |
| MRET_3984 | uncharacterized protein | 162.15 | 185.18 |
| MRET_3985 | acetolactate synthase I/II/III large subunit | 17.65 | 31.4 |
| MRET_3986 | cyclin | 134.26 | 90.69 |
| MRET_3987 | tuftelin-interacting protein 11 | 12.21 | 15.43 |
| MRET_3988 | uncharacterized protein | 65.9 | 72.2 |
| MRET_3989 | establishment of cell polarity | 31.5 | 31.53 |
| MRET_3990 | separase | 260.68 | 61.41 |
| MRET_3991 | uncharacterized protein | 47.37 | 29.44 |
| MRET_3992 | conserved hypothetical protein | 37.6 | 47.02 |
| MRET_3993 | protein transport protein SEC24 | 123.84 | 166.27 |

| | | | |
|---|---|---|---|
| MRET_3994 | GDSL-like lipase/acylhydrolase | 38.95 | 63.05 |
| MRET_3995 | Got1 family protein | 127.7 | 245.43 |
| MRET_3996 | cyclin-dependent kinase 8/11 | 34.42 | 77.43 |
| MRET_3997 | translation initiation factor eIF-2B subunit alpha | 58.95 | 71.69 |
| MRET_3998 | ATP-dependent RNA helicase DDX27 | 31.29 | 102.68 |
| MRET_3999 | FHA domain protein | 919.9 | 544.84 |
| MRET_4000 | ribosomal RNA assembly protein | 1250.88 | 302.64 |
| MRET_4001 | SNF2 family helicase | 19.74 | 48 |
| MRET_4002 | geranylgeranyl transferase type-1 subunit beta | 46.71 | 96.51 |
| MRET_4003 | ATP-binding cassette, subfamily E, member 1 | 28.11 | 68.17 |
| MRET_4004 | uncharacterized protein | 42.33 | 581.74 |
| MRET_4005 | RING-14 protein | 88.32 | 152.12 |
| MRET_4006 | coatomer subunit beta | 77.99 | 98.43 |
| MRET_4007 | peptide chain release factor 1 | 75.48 | 43.78 |
| MRET_4008 | SAGA-associated factor 29 | 45.37 | 81.04 |
| MRET_4009 | uncharacterized protein | 628 | 486.92 |
| MRET_4010 | uncharacterized protein | 294.5 | 142.51 |
| MRET_4011 | metallo-beta-lactamase domain protein | 103.02 | 61.55 |
| MRET_4012 | enolase | 502.85 | 648.62 |
| MRET_4013 | insulysin | 225.48 | 211.52 |
| MRET_4014 | lariat debranching enzyme | 177.62 | 96.25 |
| MRET_4015 | fatty acid desaturase | 87.64 | 86.37 |
| MRET_4016 | cytochrome b5-like heme/steroid binding domain protein | 132.06 | 154.91 |
| MRET_4017 | uncharacterized protein | 29.64 | 59.39 |
| MRET_4018 | H/ACA ribonucleoprotein complex non-core subunit NAF1 | 50.67 | 50.76 |
| MRET_4019 | malate synthase | 853.05 | 591.97 |
| MRET_4020 | SIT4-associating protein SAP185/190 | 29.52 | 70.43 |
| MRET_4021 | acetyl-CoA acyltransferase 2 | 387.28 | 239.39 |
| MRET_4022 | conserved oligomeric golgi complex subunit 5 | 68.36 | 136.02 |
| MRET_4023 | uncharacterized protein | 69.9 | 166.35 |
| MRET_4024 | uncharacterized protein | 70.53 | 69.87 |
| MRET_4025 | hydroxymethylpyrimidine/phosphomethylpyrimidine kinase/thiaminase | 46.38 | 50.28 |
| MRET_4026 | AP-3 complex subunit beta | 10.35 | 16.65 |
| MRET_4027 | essential nuclear protein 1 | 51.41 | 59.44 |
| MRET_4028 | molecular chaperone HtpG | 3458.52 | 5200.28 |
| MRET_4029 | CCR4-NOT transcription complex subunit 6 | 132.46 | 240.35 |
| MRET_4030 | uncharacterized protein | 16.67 | 28.69 |

| | | | |
|---|---|---|---|
| MRET_4031 | chitin synthase | 79.39 | 41.27 |
| MRET_4032 | lipase | 1216.78 | 653 |
| MRET_4033 | nitrosoguanidine resistance protein | 457.74 | 103.87 |
| MRET_4034 | structural maintenance of chromosomes protein | 45.04 | 70.95 |
| MRET_4035 | conserved oligomeric golgi complex subunit 8 | 11.05 | 34.11 |
| MRET_4036 | uncharacterized protein | 62.6 | 86.71 |
| MRET_4037 | uncharacterized protein | 85.82 | 306.72 |
| MRET_4038 | trehalose 6-phosphate synthase/phosphatase | 58.89 | 168 |
| MRET_4039 | mannosyl-oligosaccharide alpha-1,2-mannosidase | 53.71 | 85.56 |
| MRET_4040 | AP-2 complex subunit mu-1 | 21.69 | 35.21 |
| MRET_4041 | tRNA-splicing endonuclease subunit Sen54 | 30.46 | 39.58 |
| MRET_4042 | calcium permeable stress-gated cation channel | 67.75 | 89.27 |
| MRET_4043 | nucleolar protein 6 | 87.05 | 132.95 |
| MRET_4044 | uncharacterized protein | 186.98 | 213.55 |
| MRET_4045 | oxidoreductase | 80.39 | 79.6 |
| MRET_4046 | cytosol aminopeptidase | 138.51 | 127.88 |
| MRET_4047 | DNA excision repair protein ERCC-4 | 9.31 | 34.1 |
| MRET_4048 | uncharacterized protein | 3536.03 | 2038 |
| MRET_4049 | uncharacterized protein | 434.26 | 223.35 |
| MRET_4050 | MFS transporter, DHA1 family, multidrug resistance protein | 227.07 | 120.03 |
| MRET_4051 | secreted protein | 178.86 | 208.13 |
| MRET_4052 | DNA replication licensing factor MCM6 | 17.45 | 28.01 |
| MRET_4053 | uncharacterized protein | 61.98 | 35.8 |
| MRET_4054 | ribosomal RNA-processing protein 7 | 72.07 | 89.16 |
| MRET_4055 | small subunit ribosomal protein S35 | 46.29 | 61.05 |
| MRET_4056 | putative methyltransferase | 131.53 | 97.3 |
| MRET_4057 | mitochondrial import inner membrane translocase subunit TIM50 | 257.28 | 225.77 |
| MRET_4058 | acetylornithine aminotransferase | 142.8 | 128.87 |
| MRET_4059 | glyoxylate reductase | 243.46 | 161.44 |
| MRET_4060 | uncharacterized protein | 29.3 | 18.13 |
| MRET_4061 | peptidyl-tRNA hydrolase domain 1 | 70.09 | 92.05 |
| MRET_4062 | trafficking protein particle complex subunit 4 | 47.57 | 92.58 |
| MRET_4063 | translation initiation factor 3 subunit M | 24.18 | 83.04 |
| MRET_4064 | actin related protein 2/3 complex, subunit 3 | 103.4 | 101.21 |
| MRET_4065 | ribosome biogenesis ATPase | 102.13 | 153.14 |
| MRET_4066 | tRNA-specific adenosine deaminase 2 | 108.05 | 223.4 |
| MRET_4067 | arginase | 57.1 | 115.35 |

| | | | |
|---|---|---|---|
| MRET_4068 | cytochrome b5 | 221.79 | 95.21 |
| MRET_4069 | glutathione S-transferase | 2143.6 | 1569.59 |
| MRET_4070 | glycoside hydrolase family 55 protein | 60.82 | 108.62 |
| MRET_4071 | conserved hypothetical protein | 53.78 | 96.55 |
| MRET_4072 | DUF159 domain protein | 24.13 | 102.91 |
| MRET_4073 | MFS sugar transporter | 124.13 | 144.08 |
| MRET_4074 | 23S rRNA (uridine2552-2'-O)-methyltransferase | 132.27 | 262.42 |
| MRET_4075 | protein-lysine N-methyltransferase EEF2KMT | 26.04 | 48.48 |
| MRET_4076 | DnaJ homolog subfamily A member 5 | 7.51 | 18.41 |
| MRET_4077 | uncharacterized protein | 19.96 | 44.77 |
| MRET_4078 | NADH dehydrogenase (ubiquinone) 1 alpha subcomplex subunit 8 | 174.64 | 297.12 |
| MRET_4079 | uncharacterized protein | 82.55 | 152.95 |
| MRET_4080 | uncharacterized protein | 95.89 | 72.92 |
| MRET_4081 | succinate-semialdehyde dehydrogenase/glutarate-semialdehyde dehydrogenase | 408.18 | 300.84 |
| MRET_4082 | PA domain protein | 69.64 | 129.33 |
| MRET_4083 | Sec14 cytosolic factor | 322.01 | 417.66 |
| MRET_4084 | N-alpha-acetyltransferase 10/11 | 120.85 | 456.63 |
| MRET_4085 | uncharacterized protein | 60.02 | 140.56 |
| MRET_4086 | nucleolar GTP-binding protein | 29.07 | 124.65 |
| MRET_4087 | uncharacterized protein | 192.1 | 194.16 |
| MRET_4088 | zinc finger protein | 330.82 | 239.98 |
| MRET_4089 | 1-pyrroline-5-carboxylate dehydrogenase | 32.05 | 40.32 |
| MRET_4090 | trafficking protein particle complex subunit 3 | 81.24 | 163.68 |
| MRET_4091 | axial budding pattern protein 2 | 15.95 | 31.59 |
| MRET_4092 | nucleoporin GLE1 | 12.51 | 26.64 |
| MRET_4093 | N-glycosylase/DNA lyase | 55.48 | 102.24 |
| MRET_4094 | SNF2 family helicase ATPase | 58.74 | 160.59 |
| MRET_4095 | uncharacterized protein | 218.07 | 223.11 |
| MRET_4096 | secreted aspartic endopeptidase | 19.15 | 50.7 |
| MRET_4097 | uncharacterized protein | 404.85 | 852.34 |
| MRET_4098 | secretory lipase | 198.23 | 173.22 |
| MRET_4099 | secretory lipase | 40.7 | 71.02 |
| MRET_4100 | Jumonji domain protein | 293.53 | 159.42 |
| MRET_4101 | protein RER1 | 53.77 | 81.62 |
| MRET_4102 | ubiquitin carboxyl-terminal hydrolase 25 | 52.17 | 34.32 |
| MRET_4103 | HEAT repeat protein | 723.48 | 61.05 |
| MRET_4104 | NLR family carD domain protein 3 | 60.45 | 41.08 |

| | | | |
|---|---|---|---|
| MRET_4105 | mannosyl-oligosaccharide alpha-1,2-mannosidase | 18.47 | 23.83 |
| MRET_4106 | pyrimidine and pyridine-specific 5'-nucleotidase | 149.14 | 142.2 |
| MRET_4107 | beta-1,4-N-acetylglucosaminyltransferase | 17.65 | 74.07 |
| MRET_4108 | GTPase activating protein | 178.59 | 247.68 |
| MRET_4109 | alkyl hydroperoxide reductase 1 | 1981.97 | 1280.57 |
| MRET_4110 | uncharacterized protein | 262.45 | 507.35 |
| MRET_4111 | ISM6 protein | 59.26 | 308.86 |
| MRET_4112 | DnaJ homolog subfamily C member 11 | 35.3 | 63.66 |
| MRET_4113 | phosphoadenosine phosphosulfate reductase | 103.87 | 473.06 |
| MRET_4114 | large subunit ribosomal protein L4e | 234.29 | 685.34 |
| MRET_4115 | carboxylesterase family | 62.01 | 77.16 |
| MRET_4116 | thioredoxin reductase | 1846.87 | 1048.86 |
| MRET_4117 | aconitate hydratase | 154.06 | 43.42 |
| MRET_4118 | aconitase | 76.72 | 228.04 |
| MRET_4119 | Hsp90 binding co-chaperone (Sba1) | 1038.11 | 1757.99 |
| MRET_4120 | nucleolar protein 15 | 188.06 | 428.88 |
| MRET_4121 | type II protein arginine methyltransferase | 24.04 | 44.36 |
| MRET_4122 | uncharacterized protein | 12.55 | 30.9 |
| MRET_4123 | N-alpha-acetyltransferase 50 | 25.66 | 74.12 |
| MRET_4124 | actin-related protein 3 | 372.26 | 460.75 |
| MRET_4125 | phosphoinositide-3-kinase, regulatory subunit 4 | 49.01 | 38.18 |
| MRET_4126 | nucleolar protein 16 | 262.27 | 382.81 |
| MRET_4127 | retrotransposon | 77.12 | 421.52 |
| MRET_4128 | FAD synthetase | 30.6 | 60.56 |
| MRET_4129 | 5'-3' exoribonuclease 1 | 36.95 | 50.82 |
| MRET_4130 | ribosome biogenesis protein NSA1 | 69.56 | 43.62 |
| MRET_4131 | cullin-associated NEDD8-dissociated protein 1 | 136.91 | 33.44 |
| MRET_4132 | homoserine O-acetyltransferase | 275.09 | 89.48 |
| MRET_4133 | uncharacterized protein | 35.57 | 29.73 |
| MRET_4134 | transcription elongation regulator 1 | 22.15 | 26.14 |
| MRET_4135 | uncharacterized protein | 72.55 | 44.62 |
| MRET_4136 | vacuolar protein 8 | 339.1 | 352.85 |
| MRET_4137 | cytochrome c oxidase subunit 6a | 493.32 | 396.85 |
| MRET_4138 | uncharacterized protein | 684 | 310.27 |
| MRET_4139 | short-chain dehydrogenase reductase | 359.67 | 438.94 |
| MRET_4140 | Sec20 domain protein | 87.6 | 69.85 |
| MRET_4141 | Rab6A-GEF complex partner protein 2 | 69.69 | 69.34 |

| | | | |
|---|---|---|---|
| MRET_4142 | replication factor C subunit 1 | 57.91 | 83.41 |
| MRET_4143 | member of the PUF protein family | 78.85 | 95.75 |
| MRET_4144 | alpha/beta-hydrolase lipase | 553.97 | 498.9 |
| MRET_4145 | zinc finger protein, C2H2 type | 553.04 | 597.12 |
| MRET_4146 | bromodomain factor 1 | 258.67 | 241.28 |
| MRET_4147 | trafficking protein particle complex subunit 6 | 99.16 | 64.35 |
| MRET_4148 | E3 ubiquitin-protein ligase UBR1 | 80.68 | 77.6 |
| MRET_4149 | E3 ubiquitin-protein ligase HUWE1 | 88.76 | 146.61 |
| MRET_4150 | MAPEG family protein | 1796.92 | 796.92 |
| MRET_4151 | serine/threonine-protein kinase | 70.41 | 187.8 |
| MRET_4152 | kinetochore protein Spc25, fungi type | 26.92 | 96.32 |
| MRET_4153 | aarF domain kinase | 30.93 | 73.19 |
| MRET_4154 | MFS multidrug transporter | 45.92 | 42.1 |
| MRET_4155 | conserved hypothetical protein | 297.15 | 164.41 |
| MRET_4156 | delta3,5-delta2,4-dienoyl-CoA isomerase | 105.02 | 95.56 |
| MRET_4157 | elongator complex protein 1 | 166.8 | 51.82 |
| MRET_4158 | valyl-tRNA synthetase | 27.23 | 60.75 |
| MRET_4159 | histone H1/5 | 22.53 | 403.97 |
| MRET_4160 | regulator of chromosome condensation | 84.57 | 369.04 |
| MRET_4161 | peptidyl-prolyl cis-trans isomerase | 2389.52 | 1762.06 |
| MRET_4162 | regulator of Ty1 transposition protein 109 | 84.37 | 56.62 |
| MRET_4163 | DNA mismatch repair protein | 31.47 | 21.34 |
| MRET_4164 | uncharacterized protein | 51.31 | 46.37 |
| MRET_4165 | phosphatidate phosphatase LPIN | 244.83 | 144.31 |
| MRET_4166 | uncharacterized protein | 431.71 | 196.3 |
| MRET_4167 | unfolded protein response protein Orm1 | 1798.7 | 1241.77 |
| MRET_4168 | pre-rRNA-processing protein TSR1 | 49.83 | 107.09 |
| MRET_4169 | 26S proteasome regulatory subunit N8 | 71.26 | 120.02 |
| MRET_4170 | uncharacterized protein | 84.04 | 63.6 |
| MRET_4171 | RNA-binding protein PNO1 | 22.19 | 40.05 |
| MRET_4172 | protein YOP1 | 294.51 | 505.92 |
| MRET_4173 | molecular chaperone GrpE | 464.85 | 665.07 |
| MRET_4174 | polyadenylate-binding protein | 817.1 | 702.74 |
| MRET_4175 | nucleolar complex protein 3 | 39.06 | 44.4 |
| MRET_4176 | carboxypeptidase D | 62.94 | 67.41 |
| MRET_4177 | uncharacterized protein | 69.27 | 57.12 |
| MRET_4178 | NFU1 iron-sulfur cluster scaffold homolog, mitochondrial | 1756.25 | 758.99 |

| | | | |
|---|---|---|---|
| MRET_4179 | DNA-directed RNA polymerase I subunit RPA2 | 154.81 | 101.35 |
| MRET_4180 | protein transport protein YIF1 | 65.63 | 72.66 |
| MRET_4181 | translation initiation factor eIF1A | 102.54 | 163.27 |
| MRET_4182 | paired amphipathic helix protein Sin3a | 83.46 | 90.42 |
| MRET_4183 | dolichyl-phosphate beta-glucosyltransferase | 65.5 | 73.83 |
| MRET_4184 | uncharacterized protein | 61.69 | 99.83 |
| MRET_4185 | uncharacterized protein | 95.21 | 329.55 |
| MRET_4186 | serine/threonine-protein kinase | 38.45 | 138.7 |
| MRET_4187 | histone H2A | 1047.07 | 1245.13 |
| MRET_4188 | WD domain, G-beta repeat protein | 122.06 | 59.07 |
| MRET_4189 | uncharacterized protein | 311.09 | 153.96 |
| MRET_4190 | mitochondrial import receptor subunit TOM40 | 100.51 | 141.68 |
| MRET_4191 | PHD finger domain protein | 175.78 | 151.71 |
| MRET_4192 | osomolarity two-component system, sensor histidine kinase NIK1 | 67.58 | 76.44 |
| MRET_4193 | helix-loop-helix DNA-binding domain protein | 25.31 | 62.02 |
| MRET_4194 | DNA topoisomerase III | 48.77 | 134.34 |
| MRET_4195 | diphthine methyl ester acylhydrolase | 32.86 | 103.15 |
| MRET_4196 | meiotic recombination protein SPO11 | 43.41 | 79.31 |
| MRET_4197 | diacylglycerol diphosphate phosphatase/phosphatidate phosphatase | 54.37 | 199.99 |
| MRET_4198 | mitochondrial ABC transporter ATM | 54.44 | 455.95 |
| MRET_4199 | nitrogen regulatory protein | 35.89 | 215.07 |
| MRET_4200 | ADP-ribosylation factor-like protein 1 | 29.52 | 76.06 |
| MRET_4201 | COPII coat assembly protein SEC16 | 347.57 | 242.5 |
| MRET_4202 | peptide chain release factor subunit 1 | 42.71 | 145.25 |
| MRET_4203 | large subunit ribosomal protein L43 | 42.9 | 105.3 |
| MRET_4204 | cytokinesis protein | 15.18 | 23.36 |
| MRET_4205 | V-type H+-transporting ATPase subunit D | 95.69 | 212.87 |
| MRET_4206 | protein FAM32A | 28.46 | 67.01 |
| MRET_4207 | UMP-CMP kinase | 47.7 | 96.39 |
| MRET_4208 | protein BTN | 17.18 | 19.46 |
| MRET_4209 | charged multivesicular body protein 2A | 16.11 | 36.99 |
| MRET_4210 | mitogen-activated protein kinase kinase 2 | 61.79 | 74.11 |
| MRET_4211 | cytochrome c heme-lyase | 79.85 | 126.59 |
| MRET_4212 | arabinose-5-phosphate isomerase | 89.58 | 121.3 |
| MRET_4213 | YTH domain family protein | 45.75 | 43.75 |
| MRET_4214 | cytoskeleton-associated protein 5 | 50.93 | 41.53 |
| MRET_4215 | permease | 108.91 | 81.68 |

| | | | |
|---|---|---|---|
| MRET_4216 | homocitrate synthase | 1238.74 | 560.06 |
| MRET_4217 | uncharacterized protein | 67.22 | 129.28 |
| MRET_4218 | U4/U6.U5 tri-snRNP-associated protein 3 | 66.29 | 124.57 |
| MRET_4219 | uncharacterized protein | 17.81 | 22.67 |
| MRET_4220 | tRNA-dihydrouridine synthase 2 | 10.86 | 20.7 |
| MRET_4221 | pentafunctional AROM polypeptide | 163.26 | 80.01 |
| MRET_4222 | ATP-dependent permease | 303.8 | 178.88 |
| MRET_4223 | cardiolipin synthase | 90.65 | 58.3 |
| MRET_4224 | translation initiation factor eIF-2B subunit gamma | 39.41 | 51.64 |
| MRET_4225 | regulator of nonsense transcripts 2 | 74.79 | 66.43 |
| MRET_4226 | heterogeneous nuclear rnp K-like protein | 107.65 | 182.72 |
| MRET_4227 | small plasma membrane protein | 64.05 | 65.61 |
| MRET_4228 | diacylglycerol acyltransferase family | 73.31 | 156.39 |
| MRET_4229 | diacylglycerol acyltransferase family | 74.69 | 131.1 |
| MRET_4230 | cytochrome p450 | 325.54 | 380.36 |
| MRET_4231 | dUTP pyrophosphatase | 205.96 | 167.32 |
| MRET_4232 | putative methyltransferase | 319.05 | 276.53 |
| MRET_4233 | peptidyl-prolyl cis-trans isomerase SDCCAG10 | 75.17 | 93.51 |
| MRET_4234 | RalA-binding protein 1 | 17.6 | 21.43 |
| MRET_4235 | Ran-binding protein 3 | 75.35 | 134.1 |
| MRET_4236 | universal stress protein | 82.92 | 87.45 |
| MRET_4237 | ATP-dependent Lon protease | 610.4 | 335.39 |
| MRET_4238 | DNA polymerase delta subunit 1 | 79.7 | 84.54 |
| MRET_4239 | 25S rRNA (cytosine2870-C5)-methyltransferase | 120.05 | 432.51 |
| MRET_4240 | RHO1 GDP-GTP exchange protein 1/2 | 168.82 | 73.04 |
| MRET_4241 | calcium permeable stress-gated cation channel | 166.87 | 87.15 |
| MRET_4242 | NAD(P)H-hydrate epimerase | 458.23 | 214.34 |
| MRET_4243 | flavin-binding monooxygenase-like protein | 107.29 | 113.31 |
| MRET_4244 | putative vacuolar membrane transporter for cationic amino acids | 446.98 | 356.35 |
| MRET_4245 | amyloid beta (A4) precursor protein-binding, family B, member 1 interacting protein | 122.35 | 89.98 |
| MRET_4246 | DNA polymerase gamma 1 | 40.44 | 33.87 |
| MRET_4247 | nuclear envelope organization | 52.51 | 39.89 |
| MRET_4248 | tyrosine phosphatase family | 104.45 | 110.29 |
| MRET_4249 | SUN domain protein 1/2 | 40.84 | 34.76 |
| MRET_4250 | nucleoside-diphosphate kinase | 76.87 | 220.38 |
| MRET_4251 | L-glyceraldehyde reductase | 781.92 | 609.94 |
| MRET_4252 | ER membrane protein SH3 | 160.55 | 113.44 |

| | | | |
|---|---|---|---|
| MRET_4253 | zinc finger protein | 572.39 | 265.04 |
| MRET_4254 | DNA excision repair protein ERCC-3 | 21.3 | 35.1 |
| MRET_4255 | dynactin 4 | 30.55 | 37.32 |
| MRET_4256 | actin-interacting protein | 95.08 | 120.41 |
| MRET_4257 | DUF500 domain protein | 22.37 | 58.31 |
| MRET_4258 | Rho guanyl nucleotide exchange factor | 32.66 | 39.81 |
| MRET_4259 | chitin biosynthesis protein CHS5 | 33.42 | 49.81 |
| MRET_4260 | Vps51/Vps67 family protein | 39.62 | 44.11 |
| MRET_4261 | transcription initiation factor TFIIF subunit alpha | 244.03 | 126.16 |
| MRET_4262 | protein involved in negative regulation of iron regulon transcription | 34.93 | 33.77 |
| MRET_4263 | TPR repeat protein | 26.89 | 34.75 |
| MRET_4264 | cell division control protein 7 | 71.87 | 61.49 |
| MRET_4265 | uncharacterized protein | 72.84 | 89.57 |
| MRET_4266 | TP53 regulating kinase and related kinases | 92.65 | 95.1 |
| MRET_4267 | chromatin modification-related protein | 253.66 | 238.26 |
| MRET_4268 | DNA mismatch repair protein MSH2 | 144.95 | 141.34 |
| MRET_4269 | uncharacterized protein | 105.04 | 65.39 |
| MRET_4270 | protein transport protein SEC9 | 36.21 | 56.68 |
| MRET_4271 | mitochondrial inner membrane protease ATP23 | 17.23 | 39.59 |
| MRET_4272 | large subunit ribosomal protein L13e | 234.19 | 742.56 |
| MRET_4273 | Ras GTPase-activating-like protein IQGAP2/3 | 22.02 | 93.43 |
| MRET_4274 | serine/threonine-protein kinase | 58.63 | 56.36 |
| MRET_4275 | AT rich DNA binding protein | 44.42 | 39.17 |
| MRET_4276 | uncharacterized protein | 70.1 | 83.62 |
| MRET_4277 | sphingomyelin phosphodiesterase | 112.02 | 87.03 |
| MRET_4278 | 2-phosphoxylose phosphatase | 271.65 | 148.85 |
| MRET_4279 | aryl-alcohol dehydrogenase | 1125.54 | 592.91 |
| MRET_4280 | ion channel regulatory protein UNC-93 | 62.31 | 17.27 |
| MRET_4281 | eukaryotic aspartyl protease | 40.11 | 40.44 |
| MRET_4282 | eukaryotic aspartyl protease | 15.78 | 22.7 |
| MRET_4283 | uncharacterized protein | 378.51 | 466.49 |
| MRET_4284 | glucose oxidase | 28.66 | 41.79 |
| MRET_4285 | DnaJ-related protein SCJ1 | 49.3 | 67.38 |
| MRET_4286 | eukaryotic aspartyl protease | 55.49 | 78.14 |
| MRET_4287 | cathepsin D | 158.62 | 125.26 |
| MRET_4288 | mitochondrial import inner membrane translocase subunit TIM10 | 92.08 | 217.95 |
| MRET_4289 | small subunit ribosomal protein S10e | 39.22 | 162.48 |

| | | | |
|---|---|---|---|
| MRET_4290 | protein of unknown function (DUF2034) | 85.73 | 65.22 |
| MRET_4291 | DASH complex subunit DAD2 | 198.28 | 131.64 |
| MRET_4292 | uncharacterized protein | 269.68 | 165.36 |
| MRET_4293 | SAGA-associated factor 73 | 148.5 | 65.43 |
| MRET_4294 | ATP-dependent RNA helicase DDX35 | 89.63 | 49.69 |
| MRET_4295 | charged multivesicular body protein 7 | 76.9 | 44.67 |
| MRET_4296 | v-SNARE component of the vacuolar SNARE complex | 202.45 | 109 |
| MRET_4297 | aspartyl-tRNA synthetase | 60.21 | 39.84 |
| MRET_4298 | 2-dehydropantoate 2-reductase | 105.33 | 103.07 |
| MRET_4299 | translation machinery associated TMA7 | 458.55 | 139.61 |
| MRET_4300 | transcription factor | 33.05 | 64.66 |
| MRET_4301 | methionyl-tRNA synthetase | 97.48 | 93.42 |
| MRET_4302 | uncharacterized protein | 175.57 | 107.29 |
| MRET_4303 | sorting and assembly machinery component 37 | 169.25 | 94.27 |
| MRET_4304 | uncharacterized protein | 138.62 | 109.49 |
| MRET_4305 | short-chain dehydrogenase | 88.82 | 139.13 |
| MRET_4306 | NADH dehydrogenase (ubiquinone) flavoprotein 2 | 1404.71 | 1125.35 |
| MRET_4307 | uncharacterized protein | 28.34 | 36.74 |
| MRET_4308 | uncharacterized protein | 341.93 | 229.31 |
| MRET_4309 | fungal protein of unknown function (DUF1748) | 140 | 94.24 |
| MRET_4310 | ribonuclease H2 subunit B | 24.67 | 24.72 |
| MRET_4311 | uncharacterized protein | 201.75 | 166.74 |
| MRET_4312 | NADH dehydrogenase (ubiquinone) Fe-S protein 8 | 232.19 | 344.07 |
| MRET_4313 | anaphase-promoting complex subunit 3 | 21.21 | 39.89 |
| MRET_4314 | cytosolic Fe-S cluster assembly factor CFD1 | 155.08 | 85.43 |
| MRET_4315 | adenylate kinase | 163.51 | 139.09 |
| MRET_4316 | DNA repair protein REV1 | 146.35 | 64.84 |
| MRET_4317 | mitofilin | 414.72 | 261.77 |
| MRET_4318 | 3,4-dihydroxy 2-butanone 4-phosphate synthase | 1217.5 | 526.9 |
| MRET_4319 | V-type H+-transporting ATPase subunit A | 461.45 | 320.42 |
| MRET_4320 | 26S proteasome regulatory subunit T1 | 373.28 | 212.9 |
| MRET_4321 | coiled-coil domain protein 75 | 44.06 | 46.44 |
| MRET_4322 | acetyl-CoA C-acetyltransferase | 112.86 | 178.49 |
| MRET_4323 | ubiquitin-conjugating enzyme E2 variant | 245.03 | 210.15 |
| MRET_4324 | exosome complex component RRP4 | 195.35 | 87.62 |
| MRET_4325 | pre-mRNA-splicing factor CWC22 | 407.81 | 106.65 |
| MRET_4326 | DNA repair protein RAD7 | 209.85 | 82.74 |

| | | | |
|---|---|---|---|
| MRET_4327 | actin-related protein 10 | 180.88 | 99.36 |
| MRET_4328 | pre-mRNA-splicing factor SYF2 | 651 | 598.31 |
| MRET_4329 | regulator of nonsense transcripts 3 | 154.19 | 80.46 |
| MRET_4330 | ubiquitin-protein ligase involved in ER-associated protein degradation | 298.27 | 174.54 |
| MRET_4331 | proline-rich receptor-like protein kinase | 50.44 | 51.63 |
| MRET_4332 | uncharacterized protein | 48.58 | 52.21 |
| MRET_4333 | lipase esterase family protein | 47.39 | 52.3 |
| MRET_4334 | cytochrome-b5 reductase | 41.27 | 21.18 |
| MRET_4335 | saccharopine dehydrogenase (NAD+, L-lysine forming) | 261.73 | 125.93 |
| MRET_4336 | Ran-interacting Mog1 protein | 29.52 | 54.38 |
| MRET_4337 | uncharacterized protein | 43.33 | 35.4 |
| MRET_4338 | pre-rRNA-processing protein IPI3 | 108.47 | 107.39 |
| MRET_4339 | asparagine synthase (glutamine-hydrolysing) | 196.07 | 106.16 |
| MRET_4340 | mitochondrial import inner membrane translocase subunit TIM16 | 847.41 | 770.43 |
| MRET_4341 | OmpA-like domain protein | 79.61 | 85.64 |
| MRET_4342 | spinocerebellar ataxia type 10 protein domain protein | 81.15 | 94.28 |
| MRET_4343 | protein SIP5 | 265.98 | 131.89 |
| MRET_4344 | cell division control protein 42 | 195.5 | 325.92 |
| MRET_4345 | uncharacterized protein | 132.14 | 136.89 |
| MRET_4346 | centromeric DNA binding protein | 87.03 | 65.62 |
| MRET_4347 | conserved hypothetical protein | 38.9 | 31.59 |
| MRET_4348 | integrin alpha FG-GAP repeat containing protein 1 | 317.16 | 143.76 |
| MRET_4349 | V-type H+-transporting ATPase subunit a | 319.97 | 279.17 |
| MRET_4350 | adenylate kinase | 19.27 | 23.26 |
| MRET_4351 | uncharacterized protein | 11.58 | 16.57 |
| MRET_4352 | splicing factor 3A subunit 1 | 140.11 | 61.1 |
| MRET_4353 | anthranilate synthase component I | 66.76 | 55.05 |
| MRET_4354 | uncharacterized protein | 50.63 | 203.02 |
| MRET_4355 | uncharacterized protein | 772.19 | 371.64 |
| MRET_4356 | lipase | 124.99 | 91.15 |
| MRET_4357 | glycine cleavage system H protein | 206.25 | 350.5 |
| MRET_4358 | DNA-dependent metalloprotease WSS1 | 158.33 | 207.94 |
| MRET_4359 | cohesin complex subunit SA-1/2 | 27.24 | 29.97 |
| MRET_4360 | AHNAK nucleoprotein | 53.76 | 51.2 |
| MRET_4361 | uncharacterized protein | 118.86 | 233.93 |
| MRET_4362 | ubiquitin-conjugating enzyme E2 L3 | 212.47 | 232.63 |
| MRET_4363 | uncharacterized protein | 96.09 | 145.06 |

| | | | |
|---|---|---|---|
| MRET_4364 | tRNA(His) guanylyltransferase | 89.35 | 109.54 |
| MRET_4365 | peroxin-12 | 97.04 | 65.67 |
| MRET_4366 | ubiquitin carboxyl-terminal hydrolase 5/13 | 136.15 | 84.91 |
| MRET_4367 | adenylyl cyclase-associated protein | 72.95 | 48.47 |
| MRET_4368 | kinetochore protein NDC80 | 63.44 | 83.28 |
| MRET_4369 | ribonucleoside-diphosphate reductase subunit M2 | 1100.78 | 683.8 |
| MRET_4370 | YL1 nuclear protein | 222.25 | 82.52 |
| MRET_4371 | diacylglycerol acyltransferase family | 106.78 | 115.38 |
| MRET_4372 | BRCT domain protein | 43.79 | 26.77 |
| MRET_4373 | S-adenosylmethionine-dependent methyltransferase | 27.95 | 19.11 |
| MRET_4374 | periodic tryptophan protein 1 | 35.92 | 45.41 |
| MRET_4375 | ribosome production factor 2 | 49.01 | 57.42 |
| MRET_4376 | membrane-associating domain protein | 95.24 | 200.11 |
| MRET_4377 | ribonucleoside-diphosphate reductase subunit M1 | 851.17 | 408.65 |
| MRET_4378 | uncharacterized protein | 236.35 | 144.83 |
| MRET_4379 | uncharacterized protein | 69.33 | 61.58 |
| MRET_4380 | DUF833 domain protein | 31.89 | 28.26 |
| MRET_4381 | DNA-directed RNA polymerase III subunit RPC11 | 652.44 | 256.08 |
| MRET_4382 | uncharacterized protein | 666.62 | 440.35 |
| MRET_4383 | structure-specific recognition protein 1 | 140.82 | 146.8 |
| MRET_4384 | nicotinamide/nicotinate riboside kinase | 31.13 | 55.49 |
| MRET_4385 | mitochondrial 37S ribosomal protein RSM19 | 76.5 | 146.3 |
| MRET_4386 | NAD+ diphosphatase | 51.5 | 135.85 |
| MRET_4387 | pre-rRNA-processing protein IPI1 | 35.1 | 23.64 |
| MRET_4388 | MFS family protein | 149.76 | 43.87 |
| MRET_4389 | septicolysin | 999.28 | 1085.39 |
| MRET_4390 | eukaryotic aspartyl protease | 8.43 | 28.01 |
