## Supplementary material for "A novel virus alters gene expression and vacuolar morphology in *Malassezia* cells and induces a TLR3-mediated inflammatory immune response": Table S2

**Table S2. Primers used in this study**

| Primer | Sequence (5’-3’) |
|---|---|
| Tagged dN$_6$_1 | CCTGAATTCGGATCCTCCNNNNNN |
| Tagged oligo_1 | CCTGAATTCGGATCCTCC |
| Linkage1_V40 | CACACCTGACGAGTTGTATGACA |
| Linkage2_V40 | GCTGGCTGTTAAAGTCATCATAATCG |
| V40L_SP1 | CGAAATGCTCAATCGGGATGAAGC |
| V40L_SP2 | GTCCATTCCATCGAGGGAAGTT |
| V40L_SP3 | CTCACTACACAATGAGGCCATCG |
| V40L_SP4-3 | CCTCGAGTGCCAGAACTGACAT |
| V40L_SP5-2 | CATCGGCAGAGTGGACTCATGAC |
| V40L_SP6 | CTCACTGCACAACGGTGATGATA |
| V40L_SP7 | GAACAGAGTCTGAGGCGCCTA |
| V40L_SP8 | GGACTACGTCAAAGAACGCTTCAA |
| V40L_SP9 | CGGTATCGAGACTGACGAGAGAA |
| MrV40L_CP_F1 | CTCGGTAGGGTTGATGAAACCTGT |
| MrV40L_CP_R1 | CGCTTGAGGAAGTTTTCATACGAG |
| MrV40L_CP_F2 | GTGATTTCGATCTAGCCTACCACA |
| MrV40L_CP_R2 | ATTCCTCGACGCCGCCAGGACT |
| MrV40L_CP_F3 | CTCGTATGAAAACTTCCTCAAGCG |
| MrV24L_CP_F2 | GGCACAGGAAAGTATAAGCCTG |
| MrV24L_CP_R2 | ATTCCTCGACGCCGCCAGGACT |
| MrV79L_CP_F1 | GACGTATAAGCAGCATTTGGG |
| MrV79L_CP_R1 | CGGTGAGTCCCAACTGATCCA |
| MrV40L_RDRP_F1 | GATCACGCAACAGCATCATGTTCATG |
| MrV40L_RDRP_R1 | CTCAGTATGCCAATCCACAGTGTC |
| MrV40L_RDRP_F2 | GCCACCACTAGCACAAAATATG |
| MrV40L_RDRP_R2 | TATCATCACCGTTGTGCAGTGAG |
| MrV40L_RDRP_F4 | CCACTAGCACAAAATATGAGTGG |
| MrV50L_RDRP_F1 | CGCATCCACTAGTACAAAGTATG |
| MrV50L_RDRP_R3 | TCGTCACCATTGTGCAACGAG |
| MrV24L_RDRP_F1 | GGGTCTGGGATGTTACACGCAA |
| MrV24L_RDRP_R1 | CATCTTACTTAACTGTGCGCGC |
| V40S_SP1 | GAATAGCTGCGAAGAGTCAAGCA |
| V40S_SP2 | CATCGAATACTCGCTCCACAGC |
| V40S_SP3-2 | GGGCTGAAAGTTGTCATAAGACC |
| V40S_SP4 | CTGCATTTGCTTAGGTGAACATGG |
| V40S_SP5 | CAGAGACGTATGCCAGTATGAA |
| V40S_SP6 | CAGAGACCGGGTTCTGATAGT |

| | |
|---|---|
| MrV40S_F1 | GCACTATCAGAACCCGGTCTCT |
| MrV40S_F2 | CCTGCCACTGTGTTGTCAAGCAA |
| MrV40S_F3 | GCTGTGGAGCGAGTATTCGATG |
| MrV40S_F5 | GGTCTTATGACAACTTTCAGCCC |
| MrV40S_R1 | GCACTATCAGAACCCGGTCTCT |
| MrV40S_ORF_F1 | ATGAAGATATTTGACTACTTTAG |
| MrV40L.CP.F_BamHI | GATGGTGGATCCTCGTTTACGTTATTTG ATCAATTGACAGGTCC |
| MrV40L.CP.R_HindIII | GATTGGTAAGCTTTTAGTGTTCCGCCGG TGCACCAT |
| MrV40S.F_BamHI | GTTGGGGATCCATGAAGATATTTGACTA CTTTAGC |
| MrV40S.R_XbaI | CCTGCTCTAGAATCAGTTACGAATTGCA ATCCAACT |
