## Supplementary material for "A novel virus alters gene expression and vacuolar morphology in *Malassezia* cells and induces a TLR3-mediated inflammatory immune response": Table S3

**Table S3. Primers used in Q-RT PCR**

| Target | | Sequence (5′-3′) |
|---|---|---|
| MRET_0131 | Forward | GGAGCATAAGCATGCACTTTTG |
| | Reverse | CCCATGCCCGCCATT |
| MRET_2499 | Forward | GCTCGGCCTGCTCCAA |
| | Reverse | CGGCATCGATACGCCTTT |
| MRET_3200 | Forward | TCCATCTTTGGGACCGTGTT |
| | Reverse | GCCCACAGGCTGCAAGTT |
| MRET_1953 | Forward | TCGATGCGCTACGGTAAAGA |
| | Reverse | CACGCTTTCCAGTCGTCTCA |
| MRET_2956 | Forward | TCCCATTGAGGGCATTCTG |
| | Reverse | CCACCGCCCTCTTCGTT |
| MRET_0230 | Forward | TGAACAAGGCGCCTGCTAA |
| | Reverse | CCGTTGCGCTCAGCAAA |
| MRET_1468 | Forward | TGATGCCGACGCTTTCG |
| | Reverse | GGACGACCATCGACCTTGAT |
| MRET_1518 | Forward | CCTTCCTTGCCCTCTTCTCAT |
| | Reverse | AGCGACGACAGGGACAATG |
| TLR3 | Forward | TTGCGTTGCGAAGTGAAGAA |
| | Reverse | TCAGTTGGGCGTTGTTCAAG |
| TLR7 | Forward | ATATCCCAGAGGCCCATGTG |
| | Reverse | ACACACATTGGCTTTGGACC |
| TLR8 | Forward | TCCTCCCTGCAAACCAAGAT |
| | Reverse | AAAACAGGACAGCTGCAGTG |
| TLR9 | Forward | CCTGAAGTCTGTACCCCGTT |
| | Reverse | TCTGGGCTCAATGGTCATGT |

| | | |
|---|---|---|
| β-actin | Forward | CCTCTATGCCAACACAGTGC |
| | Reverse | CCTGCTTGCTGATCCACATC |
