## Supplementary material for "A novel virus alters gene expression and vacuolar morphology in *Malassezia* cells and induces a TLR3-mediated inflammatory immune response": Figure S1

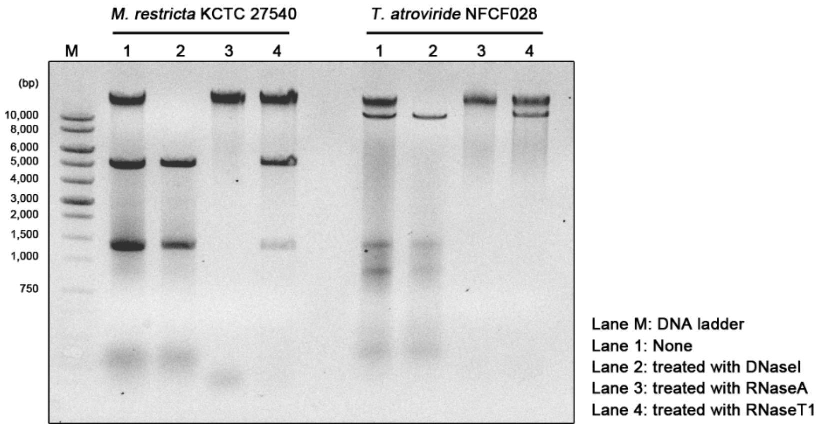


**Fig. S1. Extrachromosomal dsRNA segments in *Malassezia restricta* KCTC 27540 and *T. atroviride* NFCF028.** Nucleic acids from the strains were separated on a 0.7% agarose gel. Lane 1, total nucleic acids; lane 2, total nucleic acids treated with DNase I; lane 3, total nucleic acids treated with RNase A; lane 4, total nucleic acids treated with RNase T1.
