## Supplementary material for "A novel virus alters gene expression and vacuolar morphology in *Malassezia* cells and induces a TLR3-mediated inflammatory immune response": Figure S2

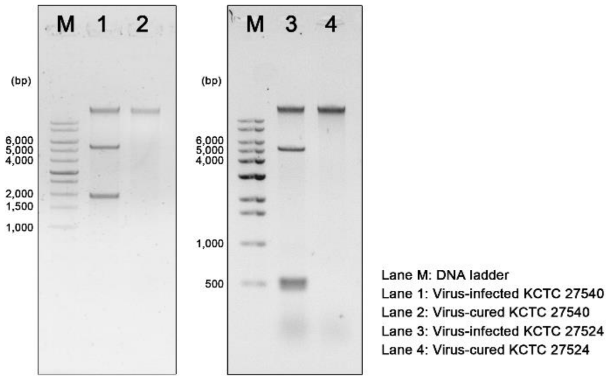


**Fig. S2. Confirmation of the virus-cured *Malassezia restricta* strain**. Total nucleic acids from the virus-infected and the virus-cured *M. restricta* KCTC 27540 and KCTC 27524 strains were extracted and treated with RNase T1. The dsRNA viral segments were not detected in the virus-cured *M. restricta* KCTC 27540 and KCTC 27524 strains (lane 2 and lane 4).
