## Supplementary material for "A novel virus alters gene expression and vacuolar morphology in *Malassezia* cells and induces a TLR3-mediated inflammatory immune response": Figure S3

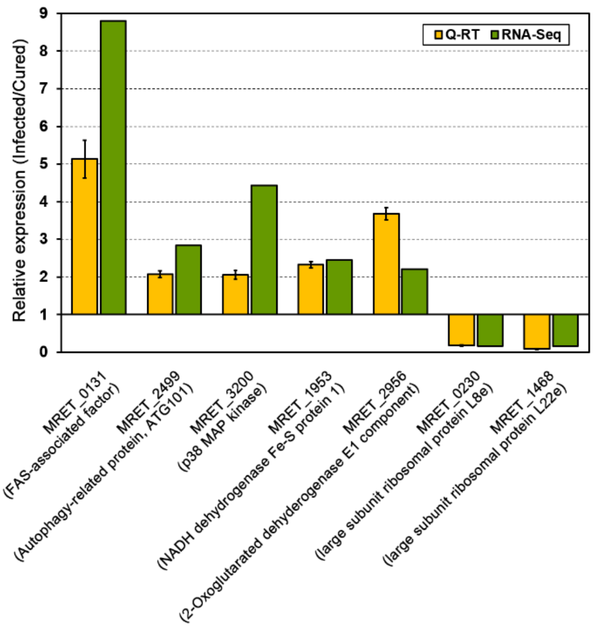


**Fig. S3. Validation of the differential expression in transcriptome analysis.** Differential expression of selected genes was confirmed by Q-RT PCR. MRET_0131 (FAS-associated factor); MRET_2499 (Autophagy-related protein, ATG101); MRET_3200 (p38 MAP kinase); MRET_1953 (NADH dehydrogenase Fe-S protein 1); MRET_2956 (2-Oxoglutarated dehyderogenase E1 component); MRET_0230 (large subunit ribosomal protein L8e); MRET_1468 (large subunit ribosomal protein L22e).
