## Supplementary material for "A novel virus alters gene expression and vacuolar morphology in *Malassezia* cells and induces a TLR3-mediated inflammatory immune response": Figure S4

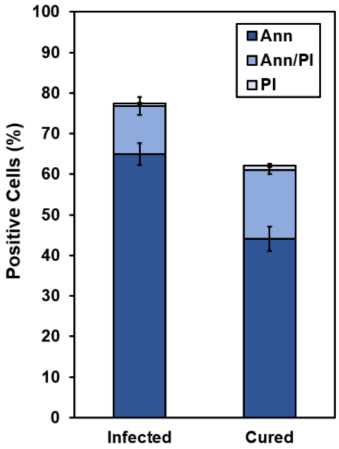


**Fig. S4. Evaluation of apoptosis in *Malassezia restricta* strains.** Virus-infected and virus-cured *M. restricta* KCTC 27540 cells were stained with Annexin V and propidium iodide (PI) to evaluate apoptosis. Annexin V binds to phosphatidylserine (PS), which translocate from the inner leaflet of the plasma membrane to the outer leaflet upon apoptosis while the plasma membrane integrity is maintained (1). PI is permeable when cell membrane is ruptured by damage or upon cell death (2). Overall apoptosis was quantified by Annexin V(Ann), and early and late apoptotic events were quantified by Annexin V(Ann)/PI and PI, respectively.


1. Martin S, Reutelingsperger C, McGahon AJ, Rader JA, Van Schie R, LaFace DM, Green DR. 1995. Early redistribution of plasma membrane phosphatidylserine is a general feature of apoptosis regardless of the initiating stimulus: inhibition by overexpression of Bcl-2 and Abl. The Journal of experimental medicine 182:1545-1556.

2. Carmona-Gutierrez D, Bauer MA, Zimmermann A, Aguilera A, Austriaco N, Ayscough K, Balzan R, Bar-Nun S, Barrientos A, Belenky P. 2018. Guidelines and recommendations on yeast cell death nomenclature. Microbial Cell 5:4
