## Supplementary material for "A novel virus alters gene expression and vacuolar morphology in *Malassezia* cells and induces a TLR3-mediated inflammatory immune response": Figure S5

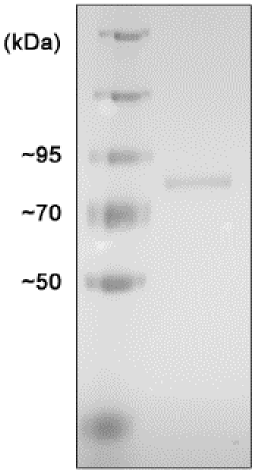


**Fig. S5. Purification of the MrV40 capsid protein**. The capsid protein encoded by ORF1 of MrV40 was heterologously expressed in *E. coli*, purified using a His-tag column, and used to analyze the expression of TLRs and cytokines in BMDCs.
